# An open-source FACS automation system for high-throughput cell biology

**DOI:** 10.1101/2023.03.24.534165

**Authors:** Diane M. Wiener, Emily Huynh, Ilakkiyan Jeyakumar, Sophie Bax, Samia Sama, Joana P. Cabrera, Verina Todorova, Madhuri Vangipuram, Shivanshi Vaid, Fumitaka Otsuka, Yoshitsugu Sakai, Manuel D. Leonetti, Rafael Gómez-Sjöberg

## Abstract

Recent advances in gene editing are enabling the engineering of cells with an unprecedented level of scale. To capitalize on this opportunity, new methods are needed to accelerate the different steps required to manufacture and handle engineered cells. Here, we describe the development of an integrated software and hardware platform to automate Fluorescence-Activated Cell Sorting (FACS), a central step for the selection of cells displaying desired molecular attributes. Sorting large numbers of samples is laborious, and, to date, no automated system exists to sequentially manage FACS samples, likely owing to the need to tailor sorting conditions (“gating”) to each individual sample. Our platform is built around a commercial instrument and integrates the handling and transfer of samples to and from the instrument, autonomous control of the instrument’s software, and the algorithmic generation of sorting gates, resulting in walkaway functionality. Automation eliminates operator errors, standardizes gating conditions by eliminating operator-to-operator variations, and reduces hands-on labor by 93%. Moreover, our strategy for automating the operation of a commercial instrument control software in the absence of an Application Program Interface (API) exemplifies a universal solution for other instruments that lack an API. Our software and hardware designs are fully open-source and include step-by-step build documentation to contribute to a growing open ecosystem of tools for high-throughput cell biology.

## Introduction

Rapid advances in CRISPR/Cas and other genome editing technologies are transforming our ability to engineer the properties of cells. The ability to deploy these technologies at scale offers unprecedented opportunities for both research and clinical applications. For example, the large-scale generation of fluorescently-labeled cell libraries is enabling the mapping of sub-cellular localization and molecular interactions of all individual proteins encoded in the human genome [1,2]. For clinical applications, high-throughput cell engineering enables the identification of efficient design rules for immune cell therapies [3] and the systematic optimization of therapeutic delivery vectors [4,5]. To capitalize on these opportunities, new methods are needed to automate and accelerate the different steps required to manufacture and handle large numbers of engineered cells.

A key step is the selection of engineered cells with specific molecular properties from a heterogeneous edited pool (for example, triaging cells that have received a therapeutic payload from cells who have not) [6–9]. Fluorescent-Activated Cell Sorting (FACS) is a widely used method for cell selection [6,10,11]. Traditional FACS operates by flowing the cells single-file past a set of illumination lasers and light detectors that analyze each cell’s size and fluorescence intensity in multiple wavelengths [6,12]. The fluorescence signals can come from stains added to the cells (dyes and/or fluorescent antibodies) or from proteins that have been genetically modified to carry a fluorescent tag. For sorting, the cells are encapsulated in a stream of droplets (one cell per droplet), and any droplet containing a cell with desired characteristics is electrically charged and routed to a destination container by a large electric field [9,13]. Despite its widespread usage, to our knowledge, no instrumentation exists that fully automates sample handling and sorting, which greatly limits the ability to sequentially handle large numbers of samples. This contrasts with other workflows where automation has revolutionized basic and applied biological research practices [14–20]. For example, liquid handling equipment can assist in cell maintenance and treatment, enabling screening campaigns on the order of 10,000 to 100,000 samples per day [21–23]. Commercial flow cytometers have also been developed with robotic operation to allow high-throughput primary phenotypic screening [21–24], but remain limited to applications where samples are analyzed (profiled, see Fig 1) but not sorted.

**Fig 1.**
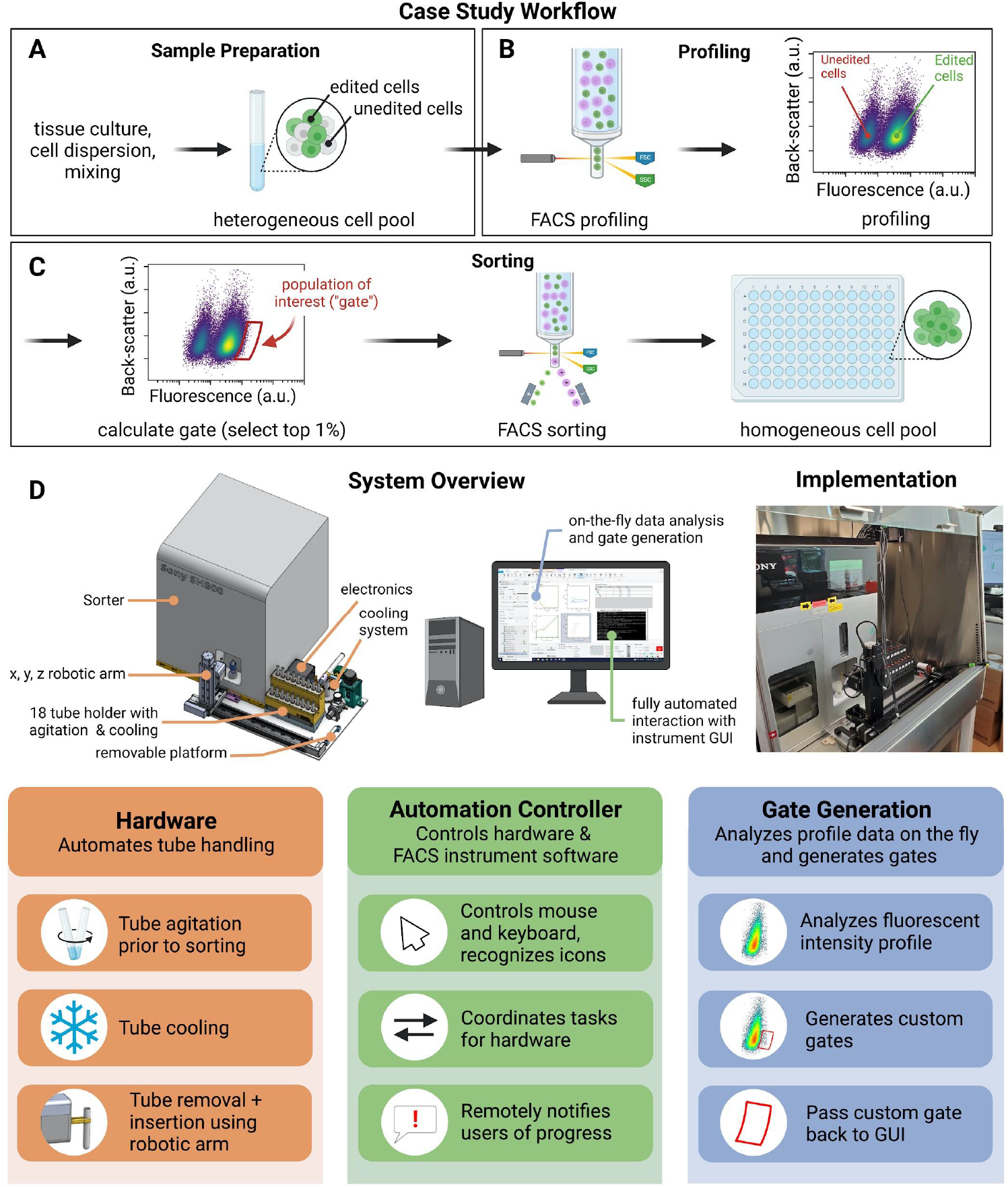
Automated sampling and control system for FACS streamlines high-throughput cell sorting workflows. (A-C) FACS automation case study and workflow. (A) Upstream sample preparation creates a library of engineered cell populations CRISPR-edited to express fluorescently tagged protein-coding genes. The cell pool contains a heterogeneous mixture of unedited (non-fluorescent) cells and successfully edited (fluorescent) cells. (B) The fluorescence signature of a representative sample of the cells is profiled via the cell sorter. (C) The profile is analyzed to specify and select the sub-population of interest within a sort gate. The cell sorter then selects those cells for collection and recovery, creating a homogenous population of successfully edited cells. (D) Design summary of the automated cell sorting system. A computer-aided design (CAD) rendering of the automated system and the real implementation highlight the details of the system (top left). The sample tubes are contained in a cooled tube housing, which agitates the samples to keep them in suspension. A robotic arm shuttles the samples between the tube housing and the cell sorter. Custom software controls the keyboard and mouse of the computer in order to interact with the sorter GUI. An algorithm analyzes the profiled cell data from (B) to generate a custom gate for each cell sample.

Here, we describe the development of an open-source software and hardware platform for FACS automation. Our platform provides walk-away automation by combining a commercial FACS instrument with a robotic arm for sample handling and custom software to autonomously control the sorting process. In particular, our software automatically generates sorting “gates” (rules for selecting the cells of interest) by analyzing the specific profile of each sample, so that libraries of samples with different properties can be sorted without operator intervention. In addition, our system communicates with the users through a messaging application to provide alerts when errors arise and when tasks are completed. This FACS automation platform reduces hands-on effort by 93%, eliminates operator error, and decreases variability in selection of the desired cell populations by applying a consistent gating algorithm.

A key aspect of our software strategy is that it controls a commercial instrument in the absence of a dedicated application programming interface (API). Our software layer operates the FACS instrument’s proprietary graphical user interface (GUI) using the open-source PyAutoGUI library [25] to programmatically control the mouse and keyboard as if a human was using them. This strategy exemplifies a universal solution for the control of other instruments for which a programmable automation interface might not be readily available. Finally, our hardware and software designs are made fully open-source and include detailed documentation to facilitate replication and re-use of components. We aim to contribute to a growing open ecosystem of tools for high-throughput cell biology.

## Results

### Case Study Design and Workflow Description

Here, we describe the development of a FACS automation platform by focusing on a specific application as a case study. OpenCell is a large-scale effort to map the localization and interactions of proteins within the human cell [1]. To this end, we use CRISPR editing to endogenously tag protein-coding genes of interest with a mNeonGreen fluorescent reporter (S1 Fig). Specifically, each engineered cell population contains a single gene tagged with the fluorescent reporter. This CRISPR engineering workflow creates an arrayed library of engineered populations expressing individual fluorescently-tagged genes, all using the same parental HEK293T cell line (S1 FIg). For each gene target, FACS is used to isolate successfully edited cells from a heterogeneous cell pool (Fig 1 A-C). This workflow encompasses the key steps common to all FACS experiments: (1) a cell suspension is loaded into the instrument (Fig 1A); (2) the fluorescence distribution of single cells is profiled from a representative fraction of that sample (Fig 1B); (3) this profile is analyzed to define a fluorescent sub-population of interest (“gating”, in our case the top 1% brightest cells, Fig 1C, left); (4) this sub-population is isolated by cell sorting into a new vessel for subsequent cell culture (Fig 1C, right).

For our high-throughput applications, libraries of fluorescently edited cells are generated in an arrayed format, typically containing 96 separate gene targets. To maximize cell viability, sorting is performed in 8x batches of 12 samples, with the requirement that each batch be processed in less than one hour outside of a tissue culture incubator (S1 Fig). The goal of the present study was to create an automation system to minimize the time and effort for researchers and enhance the scalability of our experiments. All work was performed on a SH800S FACS instrument from SONY Biotechnology. Because our specific application exemplifies general FACS operation, the concepts developed for the automation strategy described here will be applicable to other selection workflows and to other FACS instruments.

### Automation Design and Requirements

Figure 1D summarizes the different components of our automation system. A tube holder (detailed in S2 Fig) keeps up to 18 cell samples cooled (typically close to 4°C) by using cold air flow, and in suspension by vortexing, all controlled by an Arduino-based electronic unit. A robotic arm equipped with a gripper transports sample tubes between the holder and the FACS instrument. In addition, bespoke software interfaces with the sorter GUI to profile, gate, and sort the cells from each tube. With these components, the user deposits the sample tubes into the system, inputs startup information including a worklist of samples, starts the automation program, and then walks away. The overall operation can be viewed in the S1 Video.

Several key requirements guided our design. The most important is to process 12 samples within one hour with minimal user intervention, from the moment the user deposits the samples in the instrument to when all 12 samples are sorted. This time constraint maximizes cell viability and enables sorting a full set of 96 samples in a single workday. Additional specifications imposed on the system ensure sample viability and push-button operation, including: (1) maintaining the samples close to 4°C; (2) keeping the cells well-mixed in suspension; (3) automatically shuttling input samples to and from the sorter; (4) automatically controlling the commercial FACS GUI; (5) generating a custom sorting gate for each sample; and (6) communicating status updates to the operators. Finally, because the sorter is a shared resource, the automation hardware cannot interfere with the normal, manual operation of the instrument, and must reside within a biosafety cabinet where the instrument is housed. To minimize any possible interference with manual operation and instrument maintenance, all added hardware is mounted on a removable metal platform that can be easily detached (Fig 1D, left).

### Automation Software in the Absence of an API

An API is a mainstay in lab automation in order to connect and control instruments with robotics and scheduling software. However, many commercial instruments lack an API and are controlled by software designed to be interacted with only by a human operator via a GUI. While no API exists to control the SH800S instrument, we took advantage of Python’s PyAutoGUI module [25] to mimic the way a human operator would drive software operation. PyAutoGUI enables the software to programmatically read the computer screen and control the mouse and keyboard, automating all mouse clicks and text entries into the GUI.

### Automation Control Software Operation and User Communication

Overall, our bespoke control software written in Python coordinates all operations needed to seamlessly process each sample. A central strategy of our control software is to use PyAutoGUI to find and click on particular icons in the instrument’s commercial GUI, change parameters within the GUI, and enter text, such as sample names or file paths. A configuration file read by the software stores metadata that include reference images for the icons to be located, location offsets used for multiple dropdown menus present in the GUI, as well as key user-defined parameters such as the number of cells to be profiled or sorted, and the target gate percentage (by default the top 1% brightest cells). Figure 2A shows examples of specific icons that our software searches for on the computer screen to perform an action. For example, the control software looks for the blue tube icon (Fig 2A, bottom left) to perform a click on the neighboring text and rename the sample according to a user-provided worklist. For each sample, ∼60 interactions with the sorter’s commercial GUI are required to complete the profile, gate setting, and sort. A detailed and illustrated step-by-step list of all these interactions is provided in the S6 File FACS Automation Controller Software.

**Fig 2.**
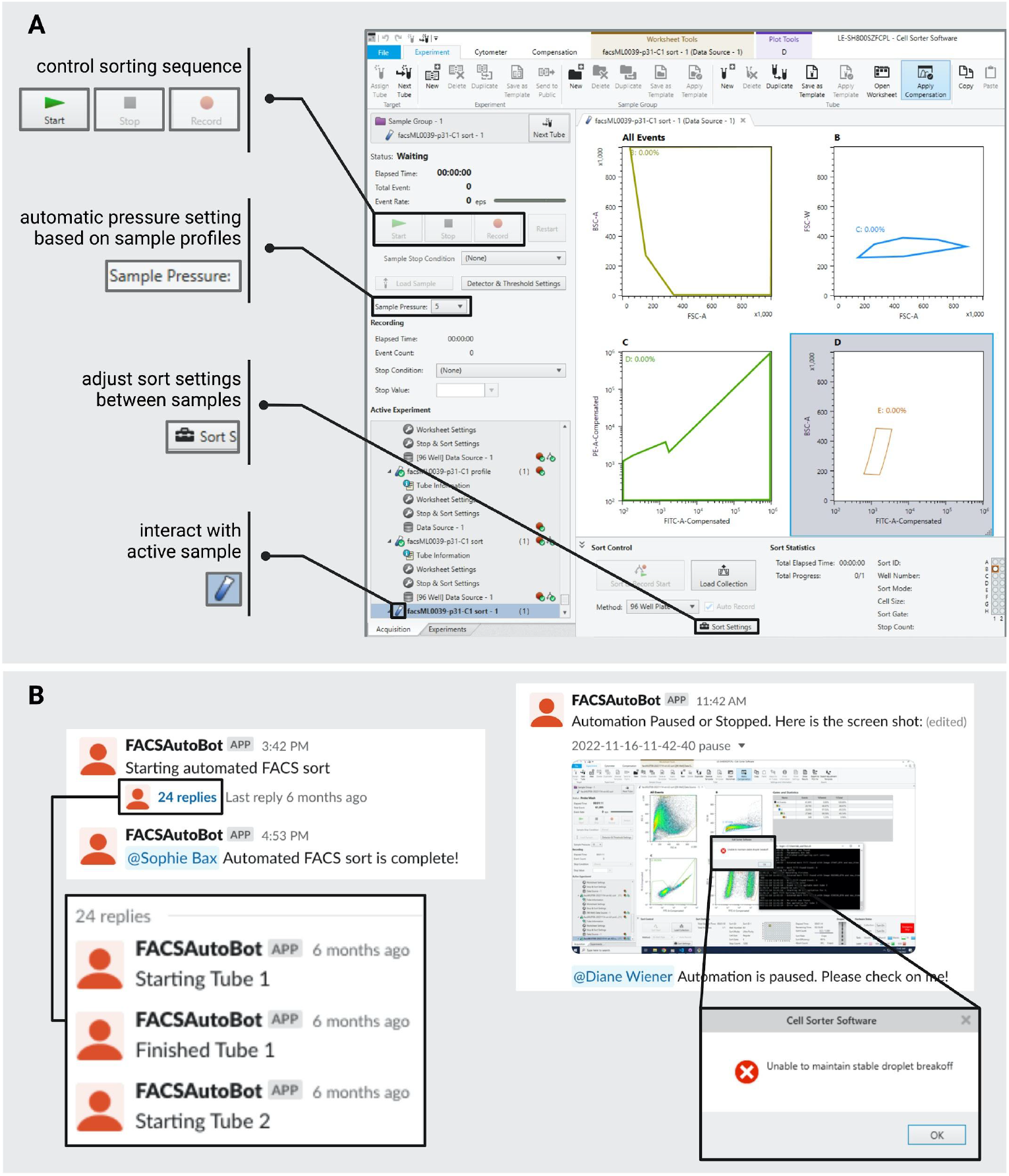
Bespoke software controls the sorter GUI and reports on instrument status to the user. (A) The automation software manipulates the sorter GUI using the PyAutoGUI Python package. The software identifies icons within the sorter GUI to operate the cell sorter (examples shown on the left). Additionally, the software enters texts and modifies the sorting settings between samples. (B) The software reports through instant messaging (Slack) on the status of a sorting run. It is capable of detecting errors with the sorter instrument and notifying the user of the problem (right panel).

A key feature of our control software is the ability to monitor the instrument’s status. This enables the software to change sorting parameters to account for sample-to-sample variation whilst maintaining the target sort time per sample to maximize sort efficiency. For example, our control software estimates the cell concentration of individual samples by reading the rate of cells that are measured by the instrument (e.p.s.: events per second) during profiling, and automatically adjusts the instrument’s sample pressure during sorting to maintain a desired flow and event rate. In addition, continuous monitoring of the instrument’s screen allows our software to identify issues or errors in real-time, which appear as pop-up windows in the GUI.

Finally, the control software communicates in real time with the human operator via instant messaging to provide updates about the run status (for example, when a given sample has finished sorting, Fig 2B left panel) and alert to any errors (Fig 2B right panel). To facilitate troubleshooting, any error notification is accompanied by an image of the instrument screen for rapid diagnosis (Fig 2B right panel). This dynamic layer of communication makes remote monitoring possible and ensures walk-away capability. At the end of a sorting run, all data are automatically exported to a backup storage location and a summary PDF report is generated, including figures of algorithmically-drawn gates (see below), so that the operator can easily examine run performance (S4 File).

### Automated Definition of Sorting Gates

A central requirement for sequential sorting of different samples is to tailor sorting gates to the specific properties of each sample. In our case study application (S1 Fig), we are generating arrayed libraries of fluorescently tagged engineered cell populations in which each sample corresponds to the tagging of a different gene from the human genome. Because different genes are expressed at different levels in the cells, and because the efficiency of CRISPR editing can vary between different genes, each sample will have a unique fluorescence signature (variations in fluorescence intensity or number of edited cells). Consequently, each sample will require a different sorting gate (see examples in Fig 3A).

**Fig 3.**
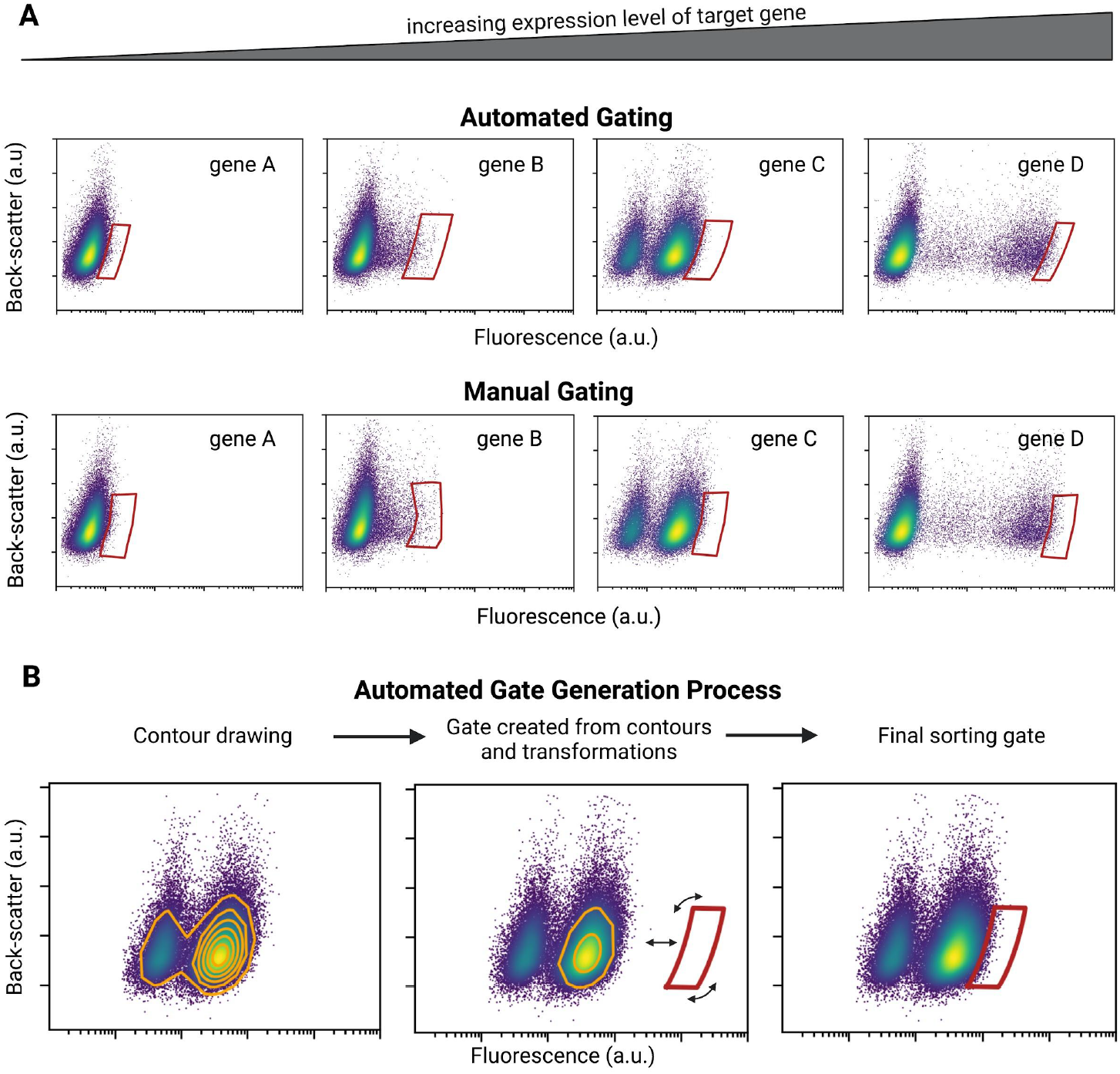
Automated gating strategy. (A) Examples of sorting gates corresponding to the fluorescent tagging of genes with increasing expression level (left to right). As the expression of the fluorescently-tagged gene increases, the gate for the desired engineered population must shift such that only the brightest cells within the sample are collected based on the specific gating algorithm. The same cell samples were sorted both automatically and manually, confirming that the gating algorithm generates results consistent with manual sorting. (B) Automated gating strategy applied in the case study. Contour lines from the fluorescent signal from the cell sample profile (see Fig. 1B) are used to define the shape and vertical position of the gate. The gate is translated and rotated such that it includes a user-defined percentage of the fluorescent population (in our case, top 1%). Alternative gating methods can be programmed into the automation control software.

In our automated workflow, just like in traditional FACS operation, the profiling step (Fig 1B) measures the fluorescence signature of each sample. Gates can then be drawn based on a user-defined selection strategy (in our case, we aim to sort the top 1% brightest cells from a plot of cell fluorescence vs back-scatter, Fig 1C). In manual FACS operation, complex gate shapes are created by manually drawing closed polygons on the plot. Following this principle, our gating algorithm approximates the calculated gate shape as a concave polygon defined by up to 60 vertices, which constitutes the final sorting gate. To define sample-specific gates, our control software first exports the raw profiling data to a local file through GUI interaction, which is then read by a bespoke Python algorithm for gate generation (Fig 3B). To tailor the shape of the gate to the profile of each sample, the gating algorithm starts by generating contour lines from the fluorescence vs back-scatter distribution (Fig 3B). For our application, we used the 80% contour line to set the height of the final gate (the range of the acceptable back-scatter signal), while the 25% contour is used to define the shape of the gate’s sides (the range of the desired fluorescence signal). The gate is then translated and rotated until it encompasses the fraction of the cell population specified by the user (in our case 1%) (Fig 3B). The goal of this strategy was to reproduce as close as possible the gates that an operator would draw manually (see Fig 3A for a comparison and S3 Fig for gate overlays comparing automation-vs manually-drawn gates from the sorts of the same engineered population). One advantage of using the automated method is that it eliminates any variation or bias between operators. It is no longer dependent on the skill and expertise of the researcher to properly draw the final gate. The gating algorithm, regardless of user, will invariably draw the same gate. In addition, the selection rules in our gating strategy can be easily modified to accommodate different sorting needs (for example, other cell types with different fluorescent properties.) For example, different shapes of gates could be used by modifying the *create_gate*.*py* class in the GitHub repo (https://github.com/czbiohub-sf/2023-facs-automation-pub).

After gate generation, the coordinates of the gate vertices are transferred into the GUI using a command line tool provided by SONY Biotechnology called *GateVertexTool*.*exe*, which is being made available for non-profit use (see Materials and Methods). This transfer step could alternatively be automated by drawing the gate with the mouse, through PyAutoGUI’s mouse-click control.

### Speed optimization

We sought to optimize the speed at which our automation system operates to both enable the highest throughput for our experiments and maximize the viability of cells during sorting by minimizing the time cells spend outside a tissue culture incubator. Since all sorting steps are performed within the FACS instrument and are the same in manual and automated workflows, no difference in cell viability is expected between these workflows to the extent their overall speed (and therefore the time that cells spend outside an incubator) is equivalent [26,27]. Our specific goal was to be able to process twelve samples in about an hour, which is the amount of time to manually sort the samples.

The time spent processing each sample can be broken into five distinct categories (Fig 4A): (1) profiling; (2) sorting; (3) hardware movement; (4) algorithmic gate calculation and (5) automated GUI control (icon searching, mouse clicks, etc.). The time spent profiling and sorting are application-specific: the user defines how many cells should be measured during profiling and how many cells should be sorted. Because the sorter operates at a set flow rate, the speed of these steps is dependent on the initial concentration of cells in a given sample (more concentrated samples will be processed quicker). For our high-throughput applications, we sort cells from small-scale input cultures (0.8 million cells per culture in 12-well plate format, S1 Fig), leading to relatively low starting cell concentrations (∼2×10^6^ cells/mL). Consequently, the profiling and sorting steps take the longest in our workflow (Fig 4). In contrast, the time spent mixing samples, moving samples with the robot arm, creating the gate and controlling the GUI are mostly sample-independent and were optimized to be as short as possible. For sample transport, we found the maximal speed achievable by the robotic arm that still ensured sample integrity (up to 1 m/s). For gate creation, we systematically compared the CPU run time for different code implementations to maximize computational speed. For GUI interactions, we used the highest possible “mouse-click” cadence that still ensured a robust operation. Overall, the automation system can process each sample in an average of 328 seconds (∼5.5 minutes). Because 75% of the total time is spent profiling and sorting (Fig 4A), it could be significantly shorter for samples with much higher cell concentrations.

**Fig 4.**
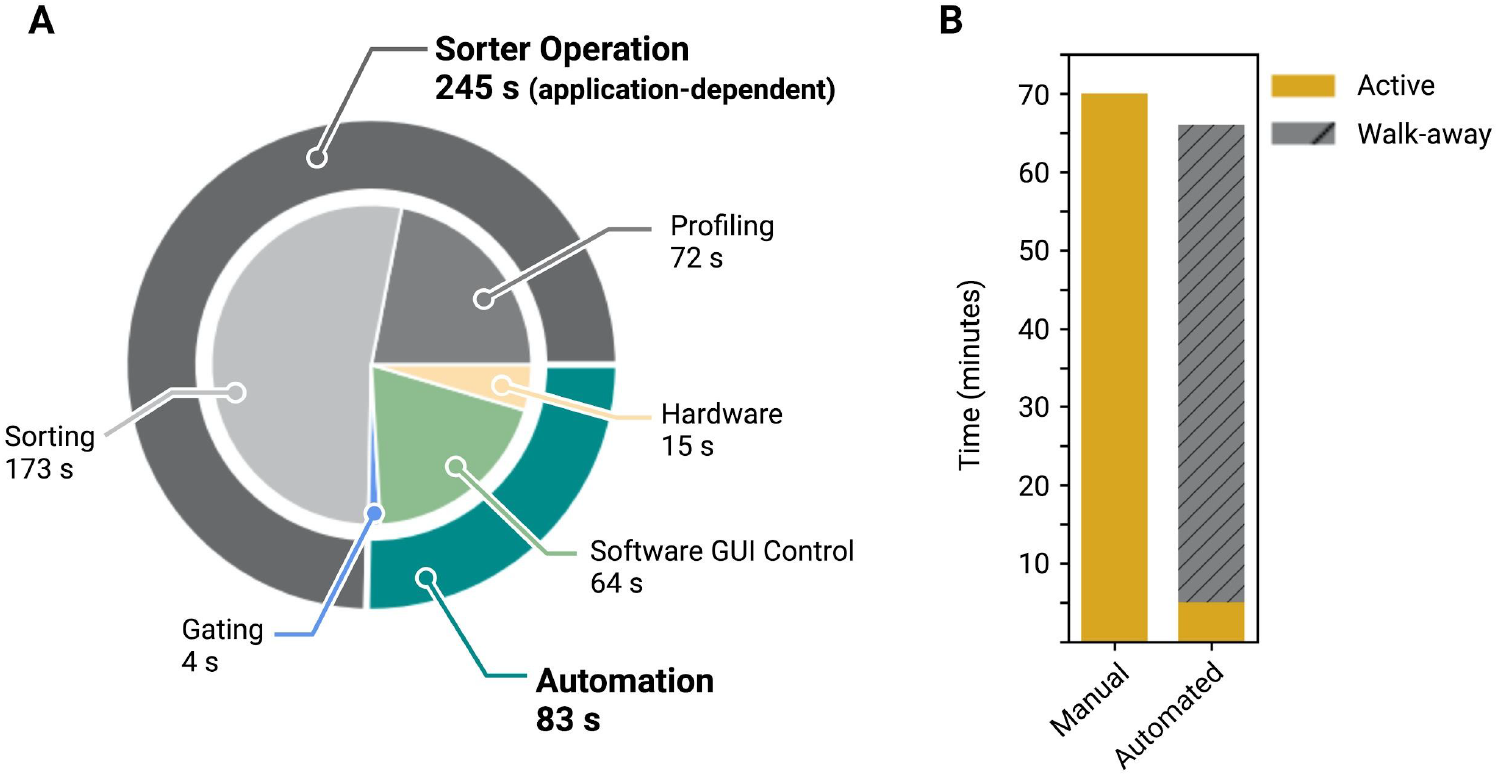
Sorting time is nominally similar between automated and manual sorting, but automation enables walk-away functionality. (A) The time spent for each process to sort a sample is divided into five steps. For the case study defined in Fig. 1A and S1 Fig, a complete sort takes on average 5.5 minutes, with 75% of the time spent profiling and sorting. The remainder of the time is spent transporting the samples, defining the gate for the sample, and setting up the sorter for each sample. (B) The same set of 12 samples were sorted with the automated system and manually. The total amount of time was comparable, but the user hands-on time for the automated system was reduced by 93%.

Our automation platform was designed to minimize the total user hands-on time. We performed an experiment comparing the amount of time that an expert user spent sorting 12 samples manually versus the amount of time that same user spent running the automation system for the same samples. Whereas the *total* run time for 12 samples was nearly equivalent: 66 minutes for the automation vs. 70 minutes for a manual run (Fig 4B), the actual *hands-on* time required from the user was significantly decreased. In manual operation, the user needed to operate the instrument for the entire 70 minutes, constantly monitoring the status of each step to be ready for the next one. In automated operation, the user spent less than 5 minutes setting up the run, and then walked away (Fig 4B). This amounts to a 93% reduction in hands-on time. While this does not amount to an increased throughput as defined by the total processing time per sample [28], the large reduction in active user time has significant impact in terms of practical throughput, which is mainly limited by operator fatigue. In our workflows, we found that a single user could handle 96 samples in one day using the automated system, compared to four users needed over two days for manual operation. The practical throughput gained with the automated system is via a significant reduction in labor. It is not possible to reduce the absolute throughput of samples handled by the FACS instrument as that is solely dependent on the sample cell density.

## Discussion

Laboratory automation is critical to increasing the throughput, accuracy, and efficiency of experiments [17–19,21]. Automation standardizes procedures and minimizes human error and variation between operators. Automation also reduces the hands-on time required from researchers, allowing them to focus on data analysis and interpretation. Another advantage is that the skill level required to operate instrumentation is reduced because complex decisions (for example, how to define sorting gates) can be made algorithmically. Overall, this results in increased robustness, reproducibility and data generation throughput, while also reducing the cost and time required.

Manual FACS is a “textbook” use-case for lab automation: a stable workflow, a high volume of samples for processing, and repetitive work with a high chance of user error. However, no end-to-end automated commercial systems exist to date. Here we described the development of a fully integrated hardware and software platform that reduces the hands-on user time by 93%, and in our experience, limits errors, and user-to-user variations, and generates equivalent results to manually sorted samples. From the speed optimization rests (Fig 4) comparing the time to sort 12 samples manually or with the automation, we observe that the manual and automated gates contain nearly equivalent populations (see S3 Fig). Furthermore, the automated gate generation is deterministic, producing the same gate for the same cell population, whereas manually drawn gates are inherently qualitative with variation between users.

A key aspect of our design is that the system is built on top of a commercial instrument (in our case, a SH800S sorter from SONY Biotechnology). To maximize flexibility, all hardware is mounted on a removable, stand-alone platform. In addition, we developed our automation software in the absence of an available API. Constrained to using the commercial GUI developed for the FACS instrument, our strategy leverages Python’s PyAutoGUI package to (1) read the information from the computer screen, (2) locate icons, menus and text boxes to be interacted with and (3) programmatically control the mouse clicks and keyboard strokes to enable end-to-end automation. This strategy could be universally applied to automate FACS sorters from different vendors, as well as other instruments beyond FACS sorters. PyAutoGUI provides a means to automate any repetitive tasks within the commercial GUI, from sorting control to data management and export. While we envision that the broad adoption of laboratory automation will encourage device companies to provide end-user APIs in the future, tools such as PyAutoGUI provide flexibility in software development for device management. One limitation of automation in the absence of an API is in error handling from the instrument. With a seamlessly integrated API, errors would immediately be flagged and the system could automatically handle some of them more directly. To mitigate this limitation, our software watches for any error popup window appearing on the screen and messages the user when an error is detected, which we found to be a suitable strategy. Such direct messaging to the user dynamically informs them on run status and notifies them of errors for greater walk-away operation.

Twin platforms have been built to automate two separate FACS instruments in our laboratory. After processing over 1,000 production samples to date (see S4 File for a representative subset of 480 algorithmically-gated sorted engineered populations), the automation system has proven to be robust, efficient at handling common errors (such as clogs), and transformative in terms of throughput and user experience (saving over 376 hours of hands-on user time). We have observed high rates of cell survival and recovery post-sort, equivalent to manual sorting (which is expected, since all sorting steps remain the same and the overall time that cells spend outside of an incubator is unchanged).

The automated FACS instrument provides a flexible solution for cell sorting in high-throughput applications. We demonstrated the utility of the system with a library of engineered cell populations. The instrument can complete sorting campaigns from libraries of different cell lines, provided that the gating strategy can be quantitatively prescribed with the cell sample profile data and implemented in a gating algorithm. Furthermore, the bespoke software can be modified to control other FACS instruments extending the functionality of the system to other selection workflows and sorters.

To enable other laboratories to reproduce our automation system, or to re-use any parts of our hardware or software designs, we provide extensive documentation in the Materials and Methods and Supplementary Information sections, including complete CAD models, a bill of materials, and a step-by-step build guide. All computer code is available on Github at: https://github.com/czbiohub-sf/2023-facs-automation-pub. Our goal is to provide fully open-source materials for FACS automation and to contribute to the growing community ecosystem of laboratory automation to achieve high-throughput cell biology.

## Materials and Methods

We include as supplementary information a fully-detailed build guide, design files and schematics, and bill of materials such that the automation system may be replicated. The total cost of the system is ∼$15,000, where the majority is the price of the robotic arm ($10,500) and machining ($3,500). The Standard Operating Procedure (SOP), software screen recording, and video of operation are included in the supplement as well. In addition, a brief summary of the design and methods is below.

### Mechanical Design

The Sony SH800S FACS is housed inside a Baker biosafety cabinet, specifically designed for this instrument. The interior space of the biosafety cabinet places hard constraints on the design of the automation system. We maximized the available space by moving the FACS to one side of the cabinet, which required creating a custom rear exhaust port, redirecting the FACS exhaust tubing, and creating a shorter side door to access the fluidics compartment.

The main hardware component of the automation system is the robotic arm that shuttles the sample tubes between the tube housing and the FACS. The robotic arm is composed of three motorized linear axis stages for XYZ motion, and a motorized gripper, all from Zaber Technologies (Vancouver, Canada). (The Supplemental Build Guide details the assembly of the robotic arm.) The linear stages and grippers have slip / stall detection, which is imperative to limit serious damage due to collisions. A single controller unit provides both power and communication to all of the linear stages and gripper. The automation software makes use of the Zaber Python motion library, which provides functions to move the stages and gripper.

The design of the gripper claws, which grab the sample tubes, is defined by the space constraints and the ability to tolerate high variability in location of the sample tubes when grabbing them. The claws were machined from aluminum (Protolabs), which allowed them to be thin and decreased the necessary space between tubes in the sample tube housing, thereby increasing the total number of sample tubes in the housing.

The space constraints also defined the design of the sample tube holder. The holder (S2 Fig) can accommodate 18 separate samples, in two rows of 9, and provides mixing and cooling.

Each sample tube is held in an off-axis socket attached to a DC motor, and gently pinched by a gasket situated ∼20 mm up from the bottom of the tube, to induce vortex mixing in the liquid by fast rotation. Each off-axis tube holder socket is funnel-shaped to ensure proper tube insertion even when the robotic arm holds the tube slightly off-axis. The rotational position of each off-axis socket is read by an optical encoder (S2 Fig), so that rotation always stops at the same angle and tubes are always presented to the robotic arm in the same orientation, for robust and reproducible gripping by the robot arm. The sample tube housing is 3D printed in Tough PLA material (MatterHackers) with an interstitial fill pattern to insulate the housing and help regulate the temperature.

The tube socket design is sensitive to the sample tube diameter (tubes from different brands show small variations in diameter). The inner diameter of the tube socket cups needs to have a fit tolerance such that the tube can be easily inserted and removed while still holding the tube at the base when the socket cup rotates. We machined the tube socket cups from teflon to decrease friction and wear on the sample tubes. We secured the tube socket cup to the DC motor shaft using a machine screw that threads into a square nut recessed in a slot in the tube socket cup and locks against the flat face of the motor shaft. Finally, encoders report the tube socket cup position so that when the motors stop the sample tube is at a precise orientation for the gripper claws to grab it. We mounted reflective strips to the tube socket cups to signal to the optical encoders when to stop the motors. This reduced the possibility of interference, slipping, or dropping tubes.

The samples must be kept cold (between 4 and 6°C) until they are sorted. To reduce the complexity of the system, we chose to use a vortex cold air gun (cat# 680, Vortec) to provide a flow of cold air through the tube housing. The vortex gun only requires a supply of filtered, dry compressed air to operate. The air pressure is regulated so that the temperature inside of the tube housing is within the acceptable range. A solenoid valve opens the air supply only during operation of the automation. Thermistors monitor the temperature inside of the tube housing, and the user is alerted by the software if the temperature is outside of the acceptable range. All of the tubing carrying the cold air to the sample tube housing is wrapped in insulation to minimize thermal losses.

All of the automation hardware components mount to a custom aluminum plate, which is secured to the FACS instrument so that there is no change in the relative positions of any of the locations for the robotic arm. We added a 3D printed funnel to the FACS instrument sample holder to assist the gripper claws in properly placing the sample tubes into the FACS sample holder. Two custom clamps attach to the locking nuts on the two front feet of the FACS and connect to a bar spanning between them. The automation platform aligns to the bar with a dowel pin and is affixed with a bracket and screws. It is easy to remove and reinstall the automation platform by disconnecting the bracket from the spanning bar and the coupling to the compressed air in ∼5 minutes. The dowel pin locator maintains the position of the platform relative to the FACS when it is reinstalled. Furthermore, the positional accuracy of the system is stable over time, not requiring any realignment.

### Electronics Design

Custom electronics, built around an Arduino Mega microcontroller board (cat# A08000, Digikey), drive many of the hardware subsystems. The electronics control (1) DC motors that spin the tube sockets for sample mixing, (2) socket encoders that stop the sockets at a precise angular position, (3) a solenoid valve that turns on and off the compressed air supply for cooling the tube housing, and (4) the temperature sensors that monitor the tube housing temperature.

The DC motors are 100:1 brushed 12V DC gear motors (cat# 3038, Pololu), chosen for their durability, power requirements, and metal shaft for connection to the tube sockets. The motors are driven by H-bridges (cat# MAX14870ETC+CT-ND, Digikey) that receive a global digital pulse width modulated (PWM) signal to control their speed, and digital signals to set the rotation direction and to turn each motor on and off. Each DC motor is paired with an optical encoder (cat# 516-2467-1-ND, Digikey) that generates a digital signal when a reflective strip on the tube socket is right in front of it. The Arduino reads when the encoder signal triggers and stops the tube socket motors. The optical encoders were chosen for their Arduino-compatible digital outputs and the small sensing distance (2 mm), which is optimal for the spacing between the tube socket cups and the mounting area for the sensor. The location of the reflective strip on the tube socket that triggers the encoders was experimentally determined by measuring the time lag between the encoder signal and the socket rotation stopping at the desired position. During normal mixing, the PWM signal that drives the motors is set to 75% duty cycle, and is reduced to 40% when the rotation is about to stop at the position detected by the optical encoder. This 75% duty cycle was selected experimentally to ensure good mixing of the cell suspension.

The solenoid valve (cat# 8262H0222, Asco Red Hat) that turns on and off the compressed air for the cooling vortex gun was chosen for its voltage requirement (12V), being normally closed (e.g., supplying power to the solenoid opens the valve), and exceeding the minimum flow requirements. As mentioned above, thermistors are used to monitor the temperature inside the tube housing. Each row of tubes in the housing has three thermistors placed along the length of the housing. Thermistors were chosen over thermocouples because of their comparatively simple readout circuit and lower hardware overhead, and because temperature readouts were expected to be within a narrow range without rapid temperature changes.

The main printed circuit board (PCB) for the system contains the H-bridge motor drivers, a transistor to drive the compressed air solenoid valve, and circuitry for the thermistor readout. Two breakout PCBs manage the optical encoders, nine encoders per PCB. The breakout PCBs are fastened inside of the tube housing spaced such that the optical encoders align with the location of the tube socket cups.

### Hardware Control

The hardware (mechanical and electrical) is controlled with a wrapper Python class that manages all of the high-level function calls to the hardware, and converts them into properly sequenced low-level function calls to the Zaber robotic arm and to the Arduino. A configuration file, which is specific for each installation of the automation, contains the communications port for both of the peripherals, the (x, y, z) location information for each position visited by the Zaber robotic arm, the accepted temperature range within the tube housing, and lag times for the motor driver stopping algorithm.

A lower-level Python class manages the Zaber robotic arm to move the arm to specific locations and to open and close the gripper. Another class manages the sending of commands to the Arduino microcontroller to control the motors, read the orientation of the tube, measure the thermistor temperatures, and toggle the state of the solenoid valve.

To maximize the speed of the automation, the travel of the robotic arms is minimized and all of the hardware operations are performed in parallel with other processes as much as possible. After careful optimization, the hardware movements add trivial amounts of time to the overall sorting procedure.

### Systems Integration and Error Handling

The complete automation system is driven by a Python wrapper class that integrates and coordinates all of the function calls for the control of the FACS GUI, automated gating, and hardware. It schedules all events based on a user-provided worklist of samples, which is a file that prescribes the sample names, location in the tube housing, and the destination well location for the sorted cells. Upon start up, the experiment name is automatically detected and recorded by the software, and the operator is asked whether the initial gates need to be drawn from a control sample. After this point the automation takes control for the remainder of the sorting process for the batch of samples defined in the worklist.

The software connects via an API to a Slack messaging workspace and posts notifications about the automation to an instrument-specific channel. Updates are provided on the status of the processing and completion of each sample. If an error occurs with the automation system (including the FACS), a mention to the user is posted to the Slack channel with information relevant to that error.

As much as possible, errors are handled automatically at the level in the software where they arise, but some errors may need to be elevated to the user. These include errors with the Zaber robotic arm, with the sorter instrument itself, and with control of the sorter GUI. Rarely, the Zaber robotic arm may fail to move to its intended location, stall, or crash if an obstruction is in the path of travel. These situations require that the Zaber motor controller be reset, e.g. needing to power cycle the controller and restart the automation. When any of these types of errors is encountered, the integration software stops and alerts the operators of a hardware error that needs their attention. For any errors that cannot be handled by the software and require user intervention, the integration software pauses the system and alerts the Slack channel notifying the user of the problem. Popup warnings from the FACS GUI occur when issues are detected within the sorter, such as sheath fluid level problems or empty sample detection. On startup, the integration software starts a thread to run in parallel with the main program to alert the user when popup warnings occur. The user must clear the error with the instrument prior to restarting the automation. Rarely, errors might occur with the automated control of the instrument GUI, such as if the GUI is much slower to refresh than normal, an icon isn’t found, or a previously stored location to return to is no longer valid. In these cases the user can simply perform the missing step. When an error occurs that requires user intervention, the user must decide whether the state of the automation system and the sorter are able to be resumed or if the automation needs to be restarted. If it is not possible to resume running a sample after certain errors with the FACS system are encountered, sorting of that sample needs to be repeated. In most cases, it is possible to recover from errors and resume the run without having to abort. To help with error handling, the integration software includes a thread that listens for keyboard hotkey presses. The keyboard listener provides three options for user intervention during a run: pause, resume, and stop. These hotkeys allow the user to address any problems and either resume or fully stop the automation.

### Sample Preparation

HEK-293T cells (ATCC CRL-3216) were cultured in D10 medium: {DMEM high-glucose medium (Gibco, cat. #11965118) supplemented with 10% fetal bovine serum (Omega Scientific, cat. #FB-11), 2mM glutamine (Gibco, cat. #25030081), and penicillin and streptomycin (Gibco, cat. #15140163)}. All cell lines were maintained at 37°C and 5% CO2 and routinely tested for the absence of mycoplasma. For endogenous gene tagging with fluorescent reporters, CRISPR/Cas9 methods were used to introduce split-mNeonGreen tag sequences into genomic open-reading frames as described in (Cho et al. Science 2022) [1].

For high-throughput cell sorting, cells were grown in 12-well plate format until ∼90% confluent (∼0.8M cells per well). Prior to sorting, cells were washed with divalent-free D-PBS (500μL/well, Gibco, cat, #14190144) and dissociated with 0.25% trypsin-EDTA for 5 min at 37°C (300μL/well, Gibco, cat. #25200056). Trypsin was quenched by the addition of 600μL D10 medium per well, cultures dissociated into single-cells by pipetting, and transferred to 1.5mL microcentrifuge tubes. Cells were spun down by centrifugation at 500xg for 5 min (room temperature), and cell pellets were resuspended in FACS resuspension medium (500μL/sample): {divalent-free HBSS (Gibco cat. #14175095) supplemented with 25mM Hepes (Gibco cat. #15630080) and 1% fetal bovine serum (Omega Scientific, cat. #FB-11)}. Cell resuspension samples were strained into 12×75mm test tubes to remove any clumps (35μm mesh size, Corning cat. #352235). Tubes were kept on ice before being decapped and loaded onto the FACS autosampler to start the sort.

In our routine protocol, 1,200 cells from each sample were sorted into a 96-well rescue plate containing 180μL D20 recovery medium per well: {DMEM high-glucose medium (Gibco, cat. #11965118) supplemented with 20% fetal bovine serum (Omega Scientific, cat. #FB-11), 2mM glutamine (Gibco, cat. #25030081), and penicillin and streptomycin (Gibco, cat. #15140163)}. Samples are typically sorted in batches of 12 (to minimize the time spent outside a tissue culture incubator), and rescued cells are placed back into a tissue culture incubator immediately after sort (37°C, 5% CO2).

### Construction

A build guide with instructions, a bill-of-materials, construction photos, and design files are provided as Supporting Information, in addition to the brief summary below.

#### Electronics

Both PCBs (main and encoder breakout) were designed in KiCad and externally fabricated by OSH Park (Portland, Oregon). The surface mount components were soldered onto the boards with a commercial reflow oven (LPKF ProtoFLow S N2, Garbsen, Germany) using solder paste (Chip-Quik Inc. SMD291SNL, SMDLTLFP10T5) applied with a stainless steel stencil (OSH Stencil, Provo, Utah) using a manual solder printer (Neoden FP2636, Ringwood, New Jersey). Details of the PCB layout and bill of materials are found in the Supporting Information.

#### Mechanics

All CAD assemblies and parts were designed in Onshape. The Onshape CAD document is available at the following link: https://cad.onshape.com/documents/c1a3ab256e8df7a71b82db8c/w/ab65c37920d47d4a2484a277/e/421207c6e308bccf3dd4e5b1?configuration=default&renderMode=0&uiState=640b930bf24bcf207edf60a6.

All of the aluminum parts were externally machined by Protolabs and Xometry. The teflon tube sockets were externally machined by Hubs. The 3D printed parts were printed in-house on an Ultimaker S5 3D printer in black Tough PLA with PVA support material. The adapter funnel on the Sony sample holder was printed in-house on a Formlabs Form3 SLA printer from durable resin material. The durable resin was washed and cured per the manufacturer’s recommendations. Silicone gaskets inside of the tube housing were cut in-house on an Universal Laser Systems PLS6.15D laser cutter.

#### Software

All software was written in Python 3.7.

Requests to access the GateVertexTool may be made to SONY Biotechnology via the following URL: https://go.sonybiotechnology.com/gate-vertex.html.

All other source code is available at the following repository: https://github.com/czbiohub-sf/2023-facs-automation-pub.

## Supporting information

Build Guide

Video of the system in operation

Hardware Bill of Materials

Electronics Bill of Materials

Example Summary Report of Sorted Samples

Automation Standard Operation Procedure

Automation Controller Software Instructions

## Acknowledgements

We thank Marco Brun for preliminary discussions and connection to Sony Biotechnology; Rinku Jana and Don Hesler at Sony Biotechnology for championing our collaboration; and Sandy Schmid and Greg Courville for feedback on the manuscript. Figure illustrations were created with BioRender.com.

## Supporting information

**S1 Fig. A CRISPR engineering workflow creates a library of fluorescently tagged protein-coding genes**.

CRISPR/Cas methods are used to tag endogenous protein-coding genes with a mNeonGreen fluorescent reporter. Corresponding cell pools contain a heterogeneous mixture of unedited (non-fluorescent) cells and cells with the successful edit. The gene edits are performed in a 96 well plate for high-throughput library generation. Prior to sorting, each cell population is expanded into 12-well plates and batches of 12 samples are processed for sorting with the FACS automation platform. In our case study, we aim to isolate 1,200 cells from the top 1% of the brightest fluorescent cell population for each sample.

**S2 Fig. The tube housing stores, cools, and shakes the samples during the automation run**.

Samples are stored in a tube housing prior to starting the automation. Up to 18 samples can be housed at a time. The tube housing is cooled with a stream of cold air generated by a vortex tube, maintaining the sample temperature close to 4°C. Each sample sits inside of a tube socket attached to a DC motor. The motor rotates the tube off-axis such that it is shaken to maintain the cells in suspension. Optical encoders ensure that the sample tubes are presented to the robotic gripper in a controlled orientation.

**S3 Fig. Overlays of the gates drawn for the same 12 samples sorted both by hand and with the automation show similar gated populations**.

Twelve samples were sorted both manually and with the automated system to compare the total time and active time to sort (Fig 4). In this figure, each graph represents the fluorescence profile from a cell population where a different gene was targeted (and will therefore display different fluorescent properties). The gates drawn by the algorithm (red) are overlaid with the gates drawn manually by the user (orange) on the final population of fluorescent cells. The gated populations are similar. Importantly, the automated gating consistently draws the same gate from the distribution of cells, whereas the manual gates can vary from user-to-user and day-to-day.

**S1 Video. The overall operation of the FACS automation system is shown in the movie**.

**S1 File. The FACS automation system build guide outlines the steps to reproduce the hardware system**.

**S2 File. The FACS Hardware BOM lists all of the parts required to reproduce the hardware system**.

**S3 File. The FACS Electronics BOM lists all of the electronic components required for the PCBs and connections**.

**S4 File. Summary PDF reports of five-96 well plate sorts (480 samples)**.

**S5 File. The FACS Automation SOP details the Standard Operating Procedure followed by users of the system**.

**S6 File. The FACS Automation Controller Software defines each step in the software control of the system**.

