## Supplementary material for "An open-source FACS automation system for high-throughput cell biology": Build Guide

### FACS Automation Build Guide

This guide serves as a manual for manufacture and assembly of the hardware for the FACS Automation platform.

| Authors | Date of Last Revision | Onshape Version (Top-level Assembly) |
| --- | --- | --- |
| Diane M. Wiener<br>Emily Huynh | 03/08/2023 | 1-0015 V1.1 Published Version |

#### Top-level assembly (1-0015 FACS Assembly)

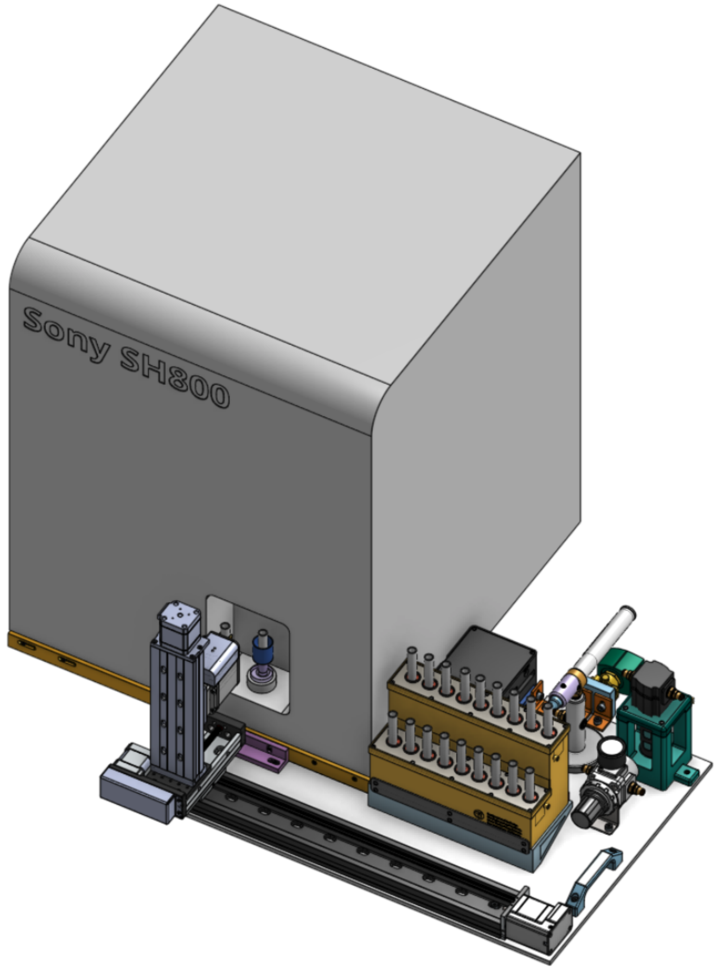

### Sub-assembly 2-0126 Sony Cell Sorter and Adapter

#### Sony SH800

The Sony SH800 is situated inside of a Baker biosafety cabinet that was specifically designed for the Sony SH800. In standard (not automated) operation, the Sony SH800 is centered inside of the cabinet. However, in order to install the automation platform, the Sony SH800 must be moved to the rear left corner of the cabinet. This requires two modifications to the instrument:

1. The factory-installed fluidics maintenance door needs to be removed from the internal hinges. Keep the screws. The door must be replaced with 3-0613 Sony Sorter Acrylic Fluidics Maintenance Door, a shorter door that can freely rotate open and clear the access panel of the biosafety hood.
2. The factory-installed HEPA filter plate needs to be removed. Keep the screws. The plate needs to be replaced with 3-0757 Sony Vacuum Plate Replacement, a plate that has a shorter exhaust port with a rotated orientation for the exhaust plumbing to properly fit with the Sony in the back left corner.

#### 3-0613 Sony Sorter Acrylic Fluidics Maintenance Door

1. Laser cut a piece of clear acrylic (2 ft x 1 ft x 0.177”) (LxWxT) with the hole pattern and outline.
    a. For the in-house laser cutter (specified below), export the face as a DXF, load into the laser cutter software, and cut with the standard power settings for acrylic (4.5 mm thick, both lasers on, 100% power, 7.2% speed).
2. Install the door on the hinges with a set of M4x16mm screws, washers, and nuts.
3. The door should fully open providing easy access to the fluidics.

#### 3-0757 Sony Vacuum Plate Replacement

1. This part was machined in aluminum by an outside vendor.
2. Unscrew the front panel from the biosafety cabinet and remove it.
3. Roll the base plate forward to easily access the back of the Sony Sorter.
4. Remove the exhaust tubing connected to the factory exhaust plate.
5. Unscrew the factory exhaust plate.
6. Mount 3-0757 Sony vacuum plate replacement with the exhaust port pointing to the top right of the biosafety cabinet using the same screws.
7. Replace the exhaust tubing.
8. Roll the base plate back and replace the front panel of the biosafety cabinet, and screw it in place.

#### Sony LE-U3M004 SYX SH800 Sample Holder

In addition, the Sony LE-U3M004 SYX SH800 Sample Holder (5 mL) needs to be shortened for the Zaber gripper robotic arm to insert the tubes in the Sony SH800.

1. Use a saw to cut the top 14 mm of the sample holder off.
2. Sand the surface smooth.

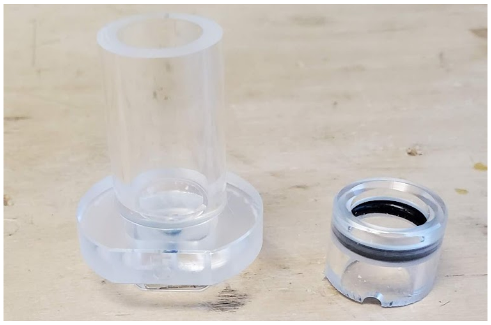

##### 3-0630 Sony Adapter Funnel

The adapter funnel is used to allow for some flexibility in positional accuracy of the Zaber stage in depositing a sample tube in the Sony sample holder.

1. This part is 3D printed in Durable resin with a Formlabs Form3 3D printer, and washed and cured following the Formlabs recommendations.
2. After removing the support material, sand the dimpled surfaces smooth.
3. Press the funnel onto the modified Sony LE-U3M004 SYX SH800 Sample Holder. It should press on tightly with manual pressure, and still be able to be removed, if needed. It is not necessary to remove the funnel to use the Sony SH800 in manual operation.

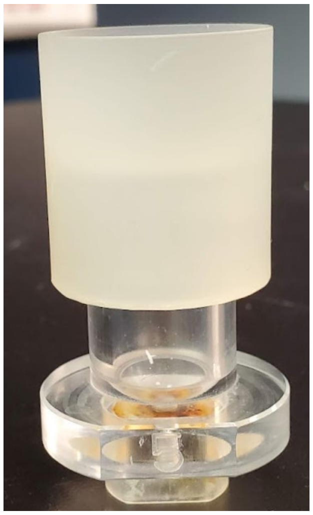

#### Sub-assembly 2-0059 FACS Zaber Stage

##### Zaber SO36459 3-axis stage

The Zaber stage consists of three linear axis stages with a gripper. The stages are mounted with the factory-provided ¼” x 20 screws. The Zaber LSQ075B-T4-ENG2690 is mounted with the center slots onto the center of the stage plate of the Zaber LSQ450X-E01T4A-ENG2710.

The LSQ150B-T4A is mounted onto the center of the stage plate of the LSQ075B. The Zaber X-GLP-E03 is mounted onto an AP-ENG3208 bracket that is fastened to the center of the LSQ150B-T4A.

1. Remove the back plate of the X-GLP-E03 gripper by unscrewing the four screws.

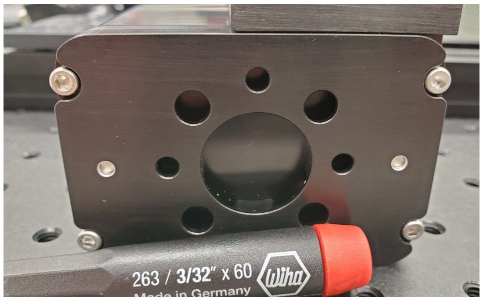

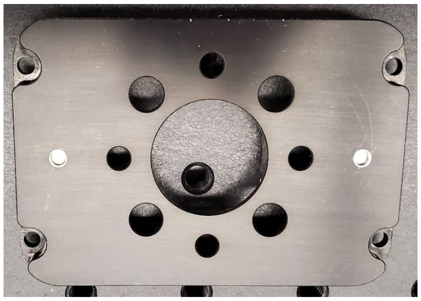

2. Attach the back plate of the X-GLP-E03 to the LSQ150B-T4A with the provided ¼” x 20 screws and with the AP-ENG3208 between them.

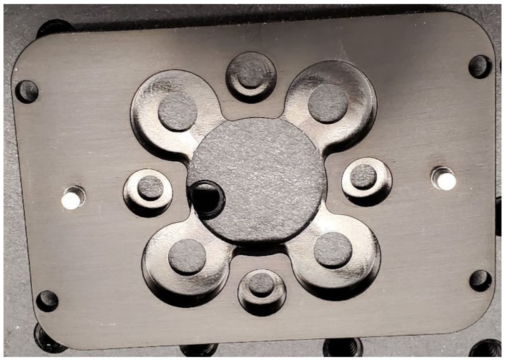
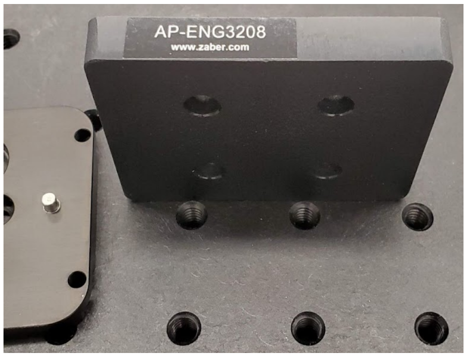

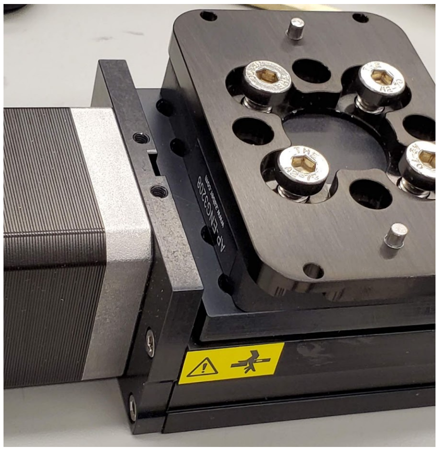

3. The pins on the back plate of the X-GLP-E03 insert into the holes on the main X-GLP-E03 gripper.
4. Using the screws that were removed previously, attach the main X-GLP-E03 to the LSQ150B-T4A. It is easiest to do this with the LSQ150B-T4A stage in the horizontal position.

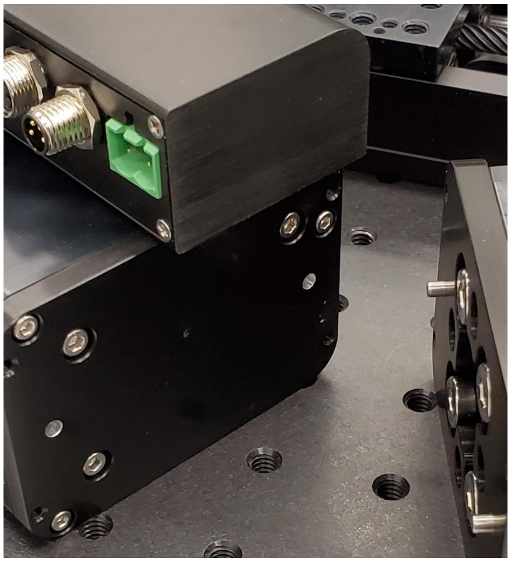
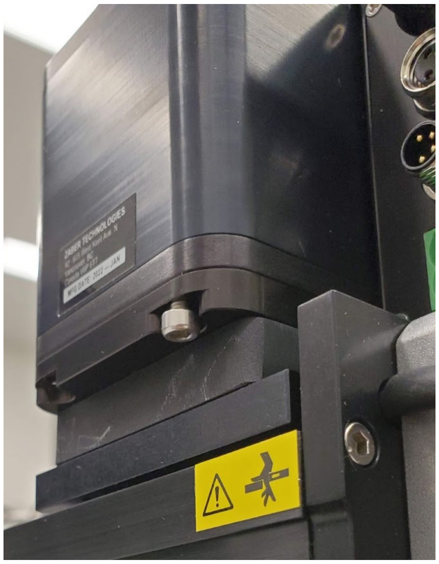

5. Attach the LSQ150B-T4A to the LSQ075B-T4-ENG2690 stage with the ¼” x 20 screws to the center of the LSQ075B-T4-ENG2690 stage.

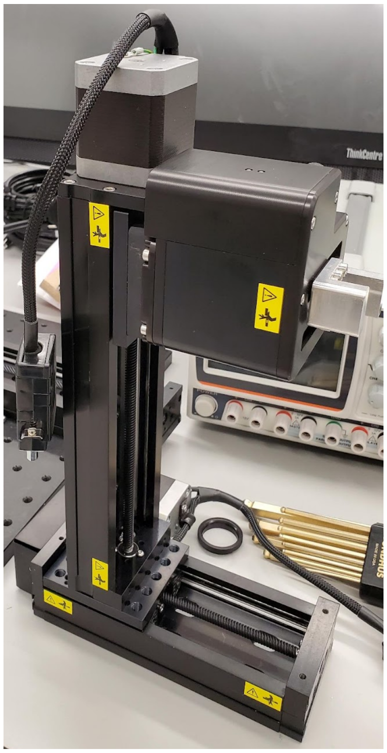
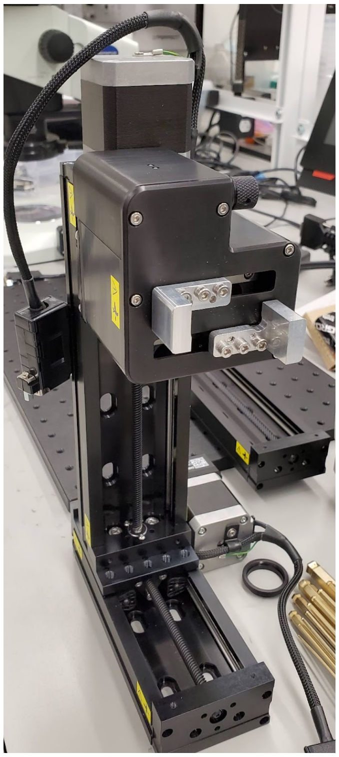

6. Attach the LSQ075B-T4-ENG2690 to the LSQ450X-E01T4A-ENG2710 stage with the low profile ¼” x 20 screws to the center of the LSQ450X-E01T4A-ENG2710 stage.

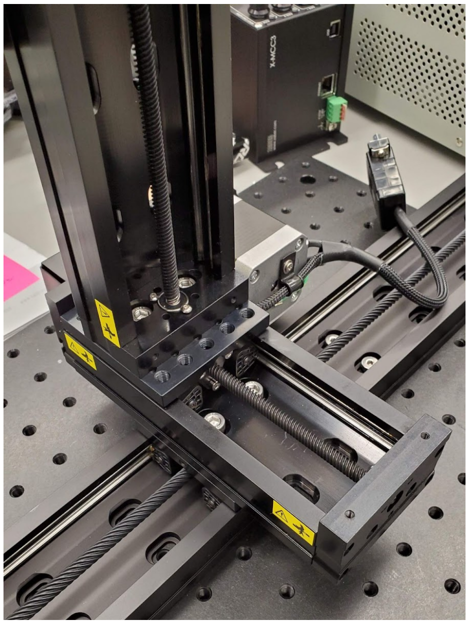

#### 3-0297-TBO Top Claw Blunted Off-axis & 3-0297-BBO Bottom Claw Blunted Off-axis

1. The claws are machined from aluminum by an outside vendor.
2. Cut heat shrink tubing to fit around the claw.
3. Use a hot air gun to shrink the tubing onto each claw.
4. Apply several coats of Pliobond 25 rubber cement to the heat shrink tubing on the claw.
5. Remove the blanks from the X-GLP-E03 and attach the 3-0297-TBO Top Claw Blunted Off-axis and 3-0297-BBO Bottom Claw Blunted Off-axis to the X-GLP-E03 in their place.

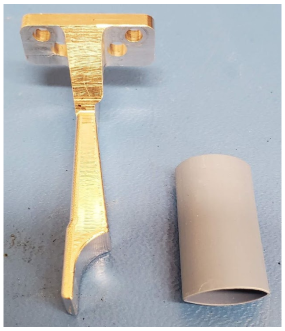
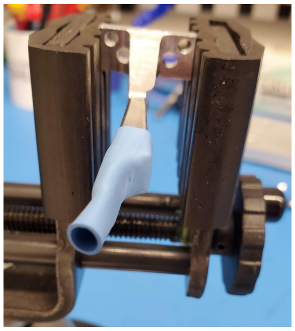
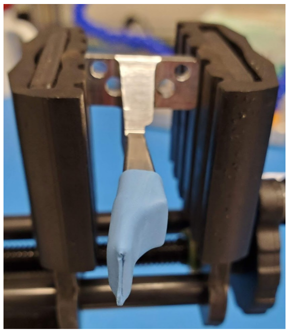
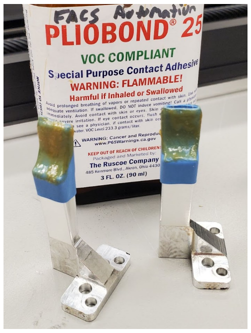

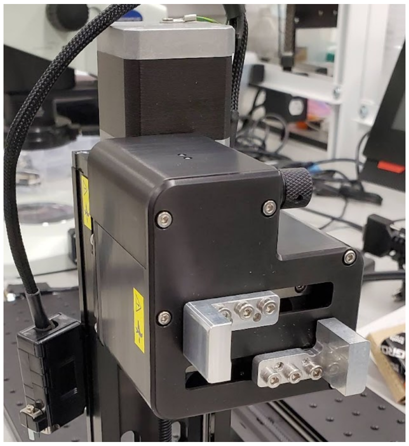

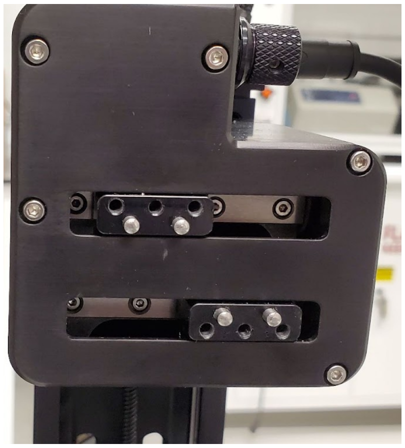

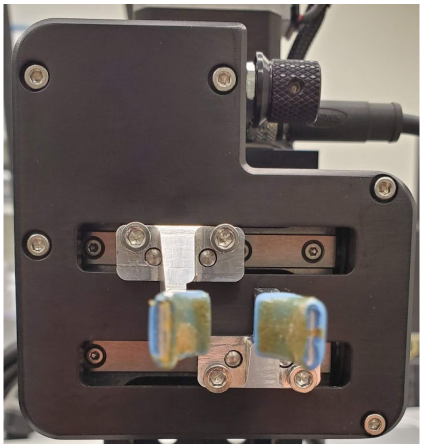

#### Sub-assembly 5-0004 & 5-0007 FACS PCBs

Both boards are fabricated by OSH Park, and 4mil stainless steel stencils are cut from OSHStencil.

##### 5-0004 FACS Automation PCB

The board is double-sided.

1. Align the corresponding stencil to the side with the logo and apply high-temperature solder paste (SMD291SNL).
2. Populate surface mount components using the schematic or electronics BOM.
3. Use a reflow oven to reflow the components using the soldering profile recommended by the SMD291SNL data sheet.
4. Visually inspect the board for proper reflow.
5. For the back side, align the corresponding stencil to the side without the logo and apply low-temperature paste (SMDLTLFP).
6. Populate surface mount components using the schematic or electronics BOM.
7. Use a reflow oven to reflow the components using the soldering profile recommended by the SMDLTLPF manufacturer.
8. Visually inspect the board for proper reflow.
9. Populate remaining through-hole components by hand using a standard soldering iron setup.
10. For cabling connecting the PCB to electronics in the tube housing, use a 26 connector ribbon cable (Digikey PN 3M157833-1-ND).
11. Pull back the shielding jacket about 8 inches, and splice out the last wire in the cable.
12. Cut the singular wire flush to where the jacket is, and cut the shielding jacket on the other side to be flush with the gray ribbon cable.
13. Pull the ribbon cable back into the shielding jacket until there is an equal amount of ribbon cable exposed on both sides of the shielding jacket.

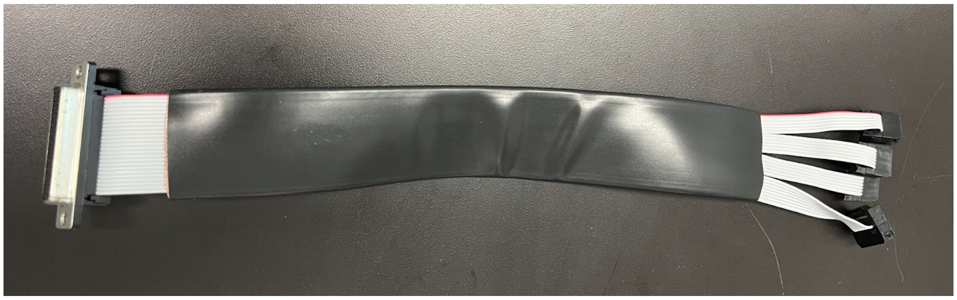

14. Using a vise, clamp a 25 pin female DB-25 with IDC termination (2057-DB25-SF-M1-ND, Digikey) onto the side of the cable with the last wire cut off. Ensure that labeled pin 1 of the DB-25 corresponds to pin 1 on the ribbon cable.

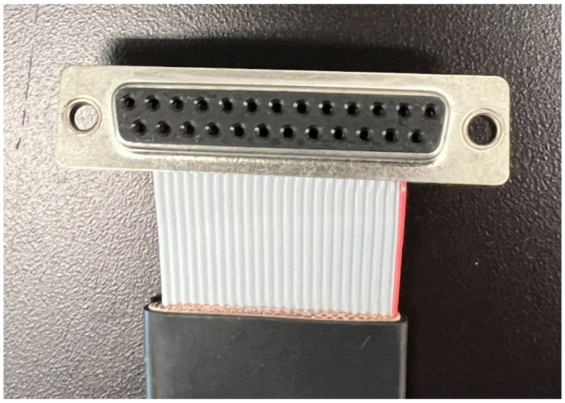

15. On the other side, splice the 26-pin connector in the following order, starting from pin 1: 3x groups of 6, and 1 group of 8.
16. For each of the 3 groups of 6 wires, clamp a 6-pin IDC terminated header (732-5443-ND, Digikey) with the header polarities seen in the image below.
17. For the remaining group, clamp an 8-pin IDC terminated header (732-5444-ND, Digikey) with the polarities seen in image below. The 8-pin header can be seen next to the white plastic housings.

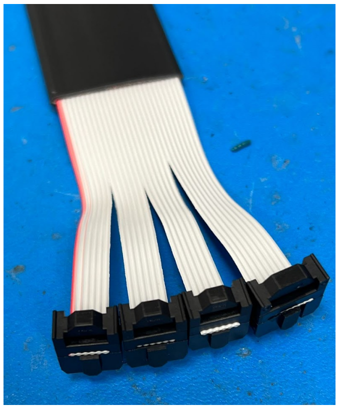 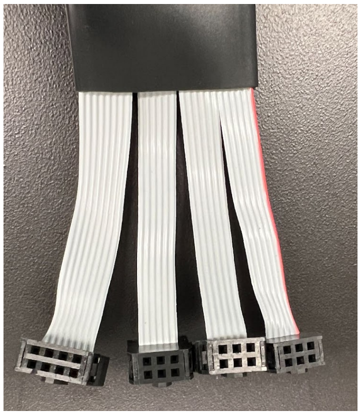 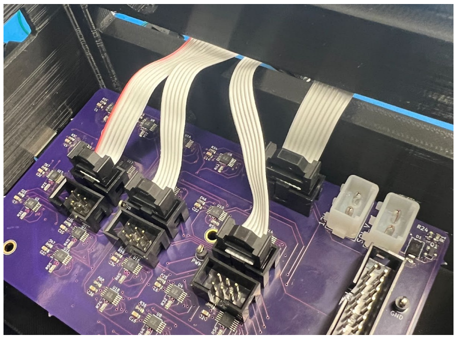

18. Repeat steps 10-17 on another 26-pin ribbon cable.
19. For the last ribbon cable, remove the wiring from the sleeve and splice off 16 connectors. Discard the remaining 11 connectors. With the 16 position ribbon cable, splice off the

very last connector entirely. The ribbon cable can be placed back into its sleeve, or remain as is.

20. Using a vise, clamp a 15 pin female DB-15 with IDC termination (AFR15B-ND, Digikey) onto the side of the cable with the last wire position snipped off. Ensure that labeled pin 1 of the DB-15 corresponds to pin 1 on the ribbon cable.

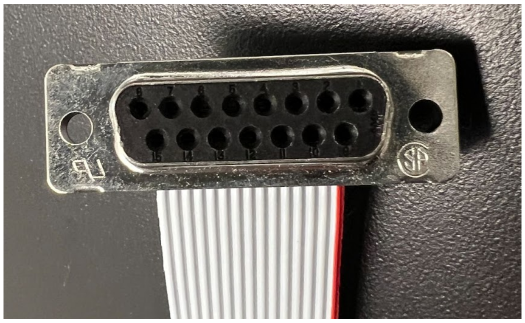

21. On the other side of the cable, clamp a 2x8 IDC terminated header (S9288-ND, Digikey), ensuring that the polarity follows the pin 1 notations on the ribbon cable and the header. The image below shows how all the ribbon cables should be connected to the PCB after assembly.

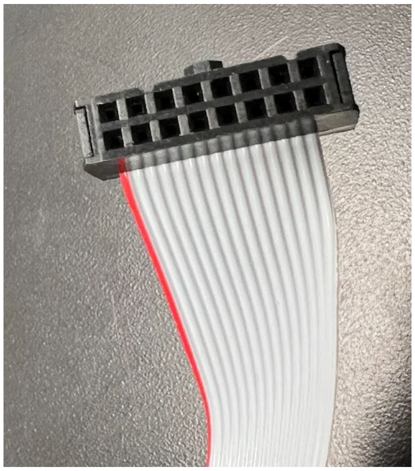

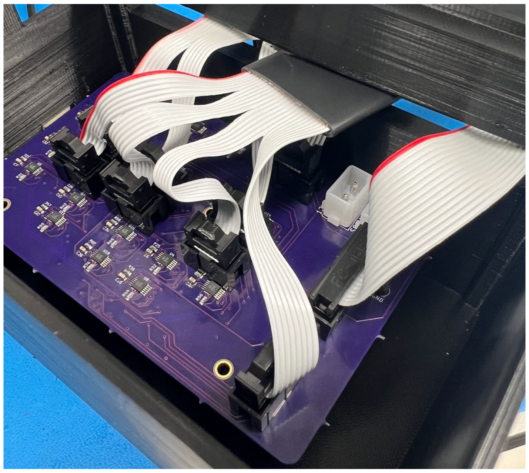

22. To assemble the connectors for power, cut about 8 inches each of red and black 18 AWG wire. Strip off 0.2" of insulation on both sides. Solder the red wire to the center of the barrel jack connector, and black to the pin on the outer side.

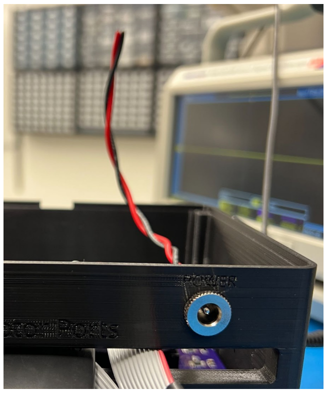

23. Add 0.25" of heat shrink to the exposed solder joints and apply heat.

24. Twist the wires together, and use a crimp tool to crimp the exposed ends of each wire (A25555-ND, Digikey).
25. Push the wires into the housing (A1411-ND, Digikey). With the polarizing key facing you, the red wire should be pushed into the left side, and the black wire into the right side.

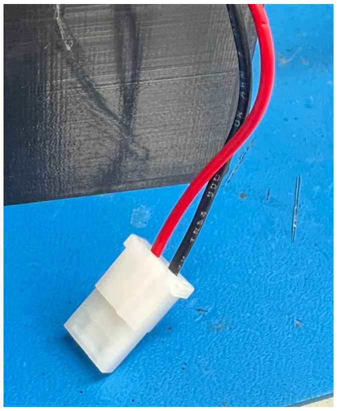

26. To assemble the connectors for the solenoid valve, cut off about 12 inches of the green wire from the solenoid valve (the neutral wire). Strip off about 0.25” of insulation from both of the remaining yellow wires.
27. Twist the solenoid valve wires together, and use a crimp tool to crimp the exposed ends of each wire (A25555-ND, Digikey).
28. Push the wires into the housing (A1411-ND, Digikey). The order does not matter.
29. See the final cabling internals of the electronics box below.

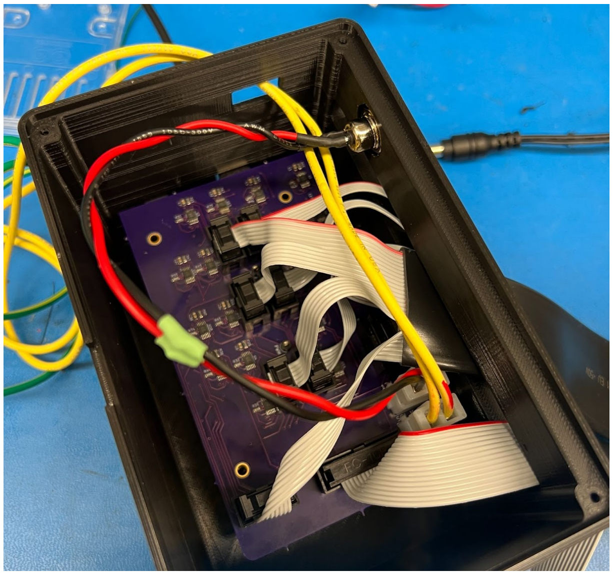

#### 5-0007 FACS Photointerrupter Breakout Board

The board is single-sided.

1. Align the stencil to the side with the logo and apply high temperature solder paste (SMD291SNL).
2. Populate surface mount components using the schematic or electronics BOM.

3. Use a reflow oven to reflow the components using the soldering profile recommended by the SMD291SNL manufacturer.
4. Visually inspect the board for proper reflow.
5. Cut the following quantities of wire: 8 inches each of black and red 26 AWG wire, 8 inches of 5x green and 4x blue 26 AWG wire.
6. On both ends of the wires, strip off about 0.2” of insulation. Solder the red wire onto the +5V pad, the black wire onto the GND pad, and alternate green and blue wires for the nine ENC pads.

### Sub-assembly 2-0119 FACS Automation Alignment Plate

#### 3-0312 Mounting plate

The plate is machined from an aluminum plate and the alignment dowel pins are press fit by an outside vendor.

#### 3-0313 Clamp mounting block

The block is machined from aluminum and the alignment dowel pin is press fit by an outside vendor.

#### 3-0628 L-bracket clamp to plate

The bracket is machined from aluminum by an outside vendor.

#### 3-1116 Tube housing attachment bracket

Laser cut 2 copies of the bracket from 5.75 mm thick acetal. When cutting with the in-house laser cutter, use the standard laser cutter settings for acetal.

#### 3-1168 Tube housing front plate

Laser cut from 1/16” acrylic. When cutting with the in-house laser cutter, use the standard laser cutter settings for acrylic.

#### 2-0120 FACS Sony Clamp

Two copies are needed to attach to the two front feet of the Sony sorter.

##### 3-0310 Primary clamp finger

Two copies are machined in aluminum by an outside vendor.

##### 3-0311 Secondary clamp

Two copies are machined in aluminum by an outside vendor.

#### 3-0574-18T FACS Tube Holder Adapter

1. Print the FACS Tube Holder Adapter with the top face on the build plate.
2. Print in Tough PLA with PVA support material with the Ultimaker S5.
3. Wash in the water bath to dissolve the PVA support material.

#### 2-0056 Tube Holder

##### 3-0281 Motor Plate

1. Print two copies of the motor plate with the base on the build plate.
2. Print in Tough PLA without support material with the Ultimaker S5.

##### 3-0283 Test Tube Guide

1. Print two copies of the test tube guide with the top on the build plate.
2. Print in Tough PLA without support material with the Ultimaker S5.

##### 3-0285 Top Plate

1. Print two copies of the top plate with the base on the build plate.
2. Print in Tough PLA without support material with the Ultimaker S5.

##### 3-0282-18T Housing

1. Print the tube housing with the base on the build plate.
2. Print in Tough PLA with PVA support material with the Ultimaker S5. The infill is 20% to create an insulation layer for containing the cold air environment inside.
3. Wash in the water bath to dissolve the PVA support material.

##### 3-0712 Rear Access Plate

1. Print the rear access plate with the back on the build plate.
2. Print in Tough PLA without support material with the Ultimaker S5. The infill is 20% to create an insulation layer for containing the cold air environment inside.

##### 3-0284 Rubber Insert

1. Laser cut 2 copies of the rubber insert gasket from 1/32” thick, 50A durometer silicone rubber sheet with crisscross texture (McMaster-Carr, 5777T11).
    a. Cutting with the in-house laser cutter, use the standard laser settings, 100% power, 89% speed, 0.8 mm thickness.

##### 3-0713 Rear Gasket

1. Laser cut the rear gasket from 1 mm thick smooth, 50A durometer silicone rubber gasket material (McMaster-Carr, 3788T22).
    a. Cutting with the in-house laser cutter, use the standard laser settings, 100% power, 89% speed, 1 mm thickness.

#### 3-0614 Motor Encoder PCB Spacer

1. Laser cut 8 copies of the spacer from 4.9 mm thick acetal.
    a. When cutting with the in-house laser cutter, use the standard laser cutter settings for acetal.

#### 2-0055 Motor Socket Assembly

##### 3-0280 Test Tube Socket

1. Eighteen copies are machined in Teflon by an outside vendor.
2. Insert the M3 square nut into the slot and secure it with the M3x6mm pan head screw and M3 washer. See the hardware BOM for specific hardware used in-house.
3. Cut a 5 mm x 25 mm strip of aluminum foil.
4. Apply Pliobond 25 rubber cement to the raised surface between the fiducial marks of the tube socket and to the matte finish side of the aluminum foil.
5. Allow the rubber cement to dry.
6. Press the aluminum foil to the tube socket.
7. Use a razor blade to cut the excess aluminum foil along the edges of the fiducial marks.
8. Repeat for all 18 tube sockets.

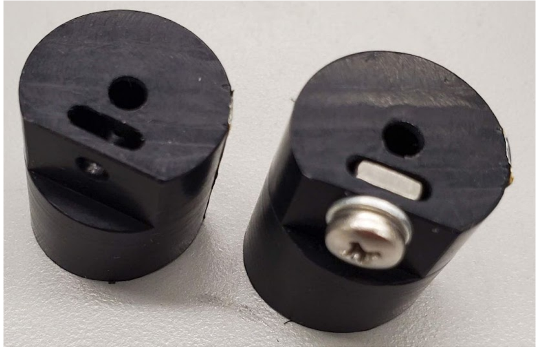
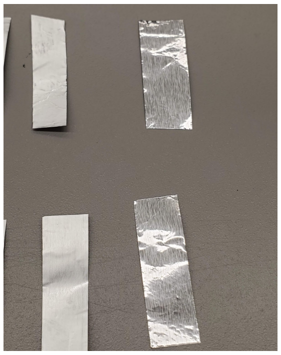
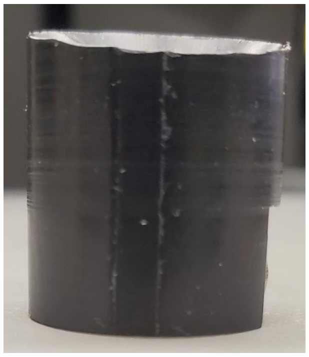
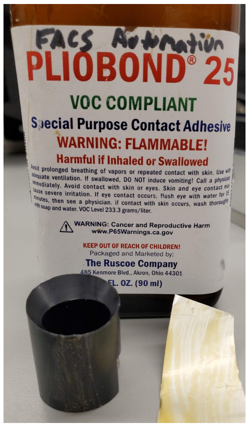

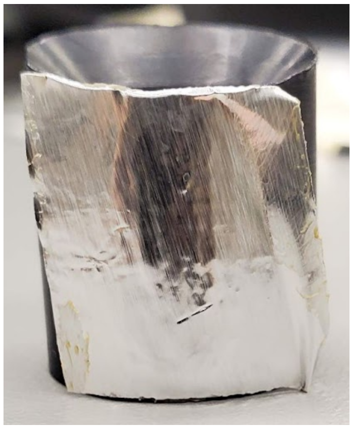
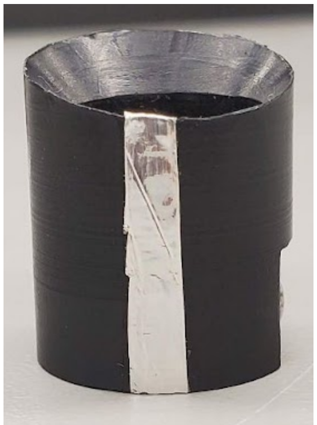
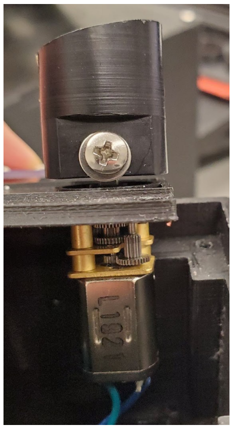

#### Final 2-0056 Tube Housing Assembly

1. Attach the 18 Pololu motors with M1.6x0.35 x 3 mm socket hex screws to the two 3-0281 motor plates.

2. Insert the motor shaft in the 3-0280 tube socket such that the flat face of the motor shaft is aligned with the side hole.
3. Use a M3x6mm pan head screw and M3 washer to fix the motor on the tube socket in place.
4. Place the two assembled motor plates into the 3-0282-18T tube housing and screw in place with #3x1/4” round head Phillips thread-forming screws.

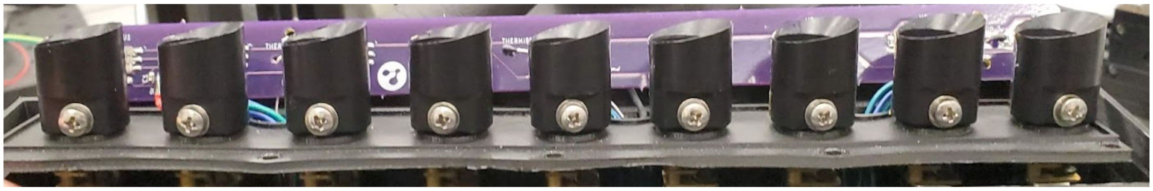

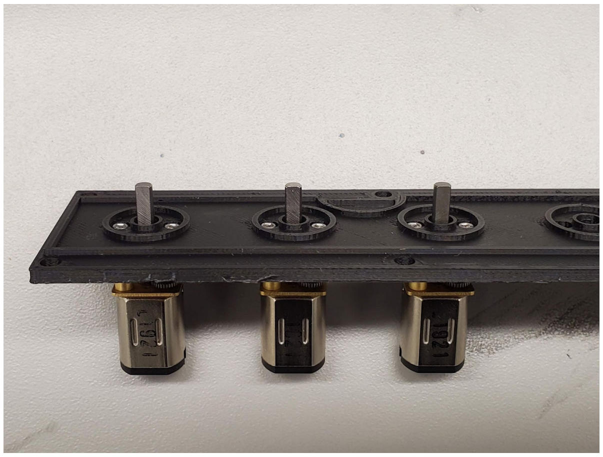

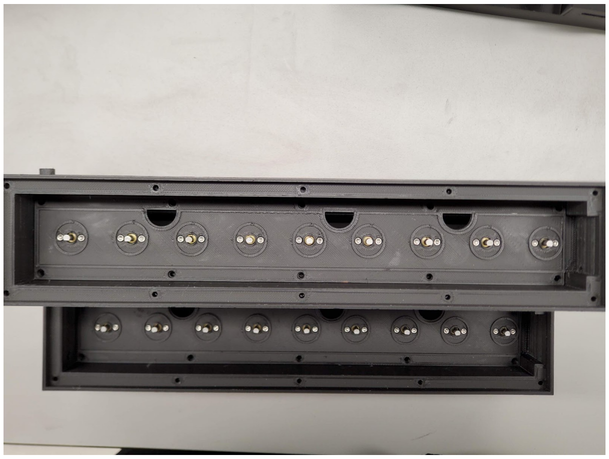

5. Cut 18x 9-inches of 26 AWG blue and green wire, and 18x 12-inches of 26 AWG blue and green wire. Strip off about 0.2” of insulation on both sides of all wires.
6. For the motors closest to the rear panel of the tube housing, use the set of 9-inch wires. With the semicircles of the motor plate facing you, solder the blue wire on the left side of the motor, and the green wire on the right side. Twist the wires together tightly. Label each set of wires with its respective motor number.
7. Repeat step 6 for the other motor plate, using the 12-inch set of wires.
8. Pre-assemble the motor encoders for the lower rail of tubes with 4x #2-56x1” socket hex screw, the 5-0007 FACS Photointerruptor Breakout Board, and 4x 3-0614 motor encoder PCB spacer.
9. Mount in the tube housing locking in place with #2 nuts.
10. Repeat for the upper rail using 4x #2-56x0.625” socket hex screws instead.
11. Route the wiring through the ports in the motor plates and to the back of the tube housing.
12. Route the thermistor leads through the holes under the labels ‘THERMISTOR_#’ on the PCB. It should be such that the epoxied head of the thermistor points towards the motor sockets.

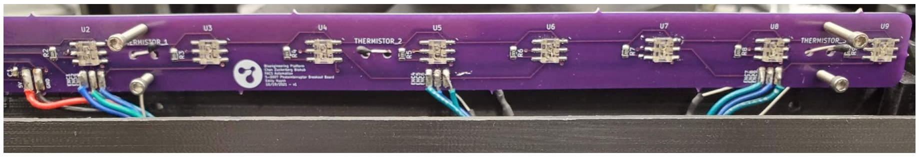

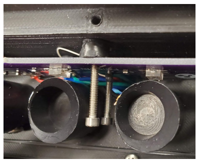

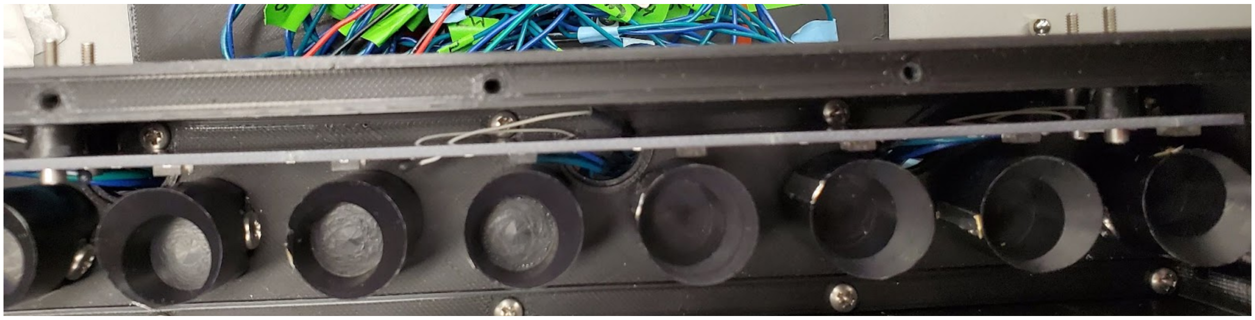

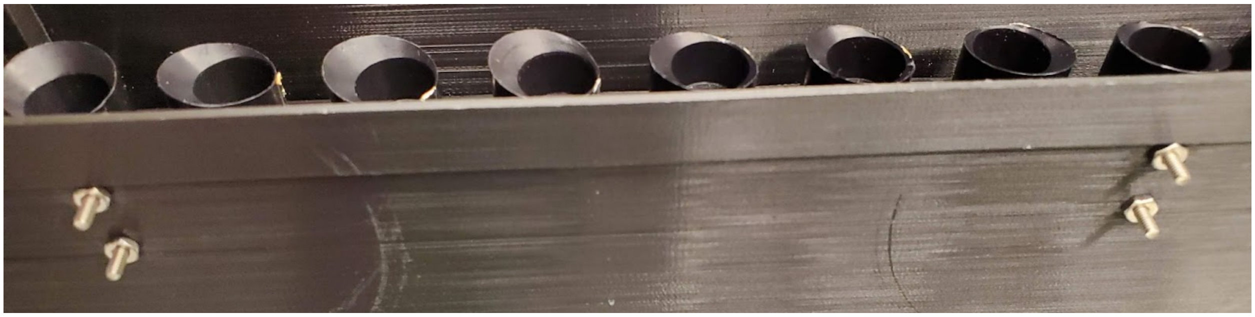

13. Connect the motor, thermistor and photointerrupter wires to the DB25 (2057-DB25-PD-ND, Digikey) and DB15 (AFR15B-ND, Digikey) connectors according to the schematic below. Prior to soldering each wire to the solder cup, add about 0.1” of D1.27mm heat shrink to apply over the solder joint after soldering.

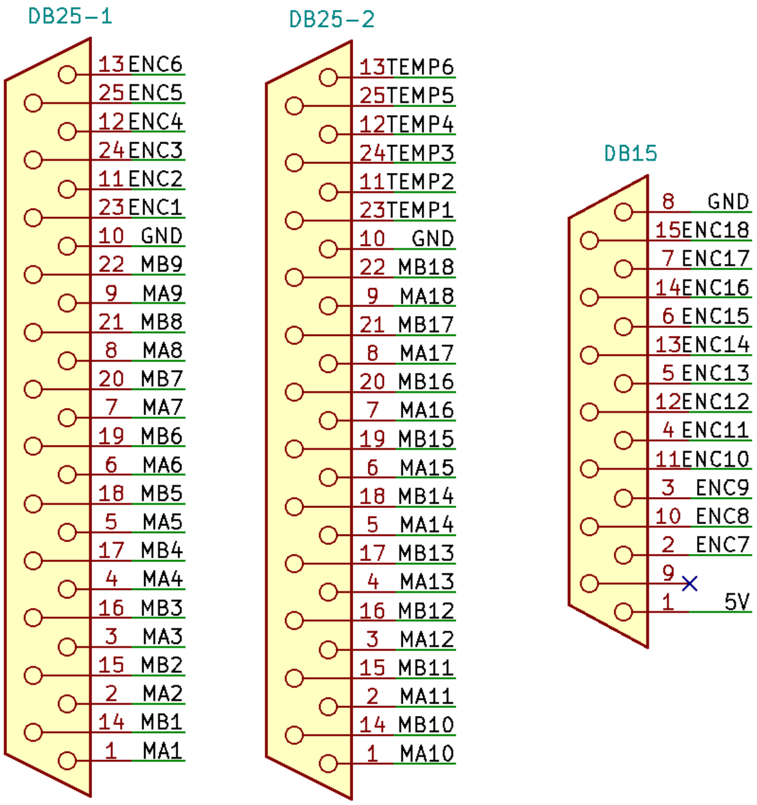

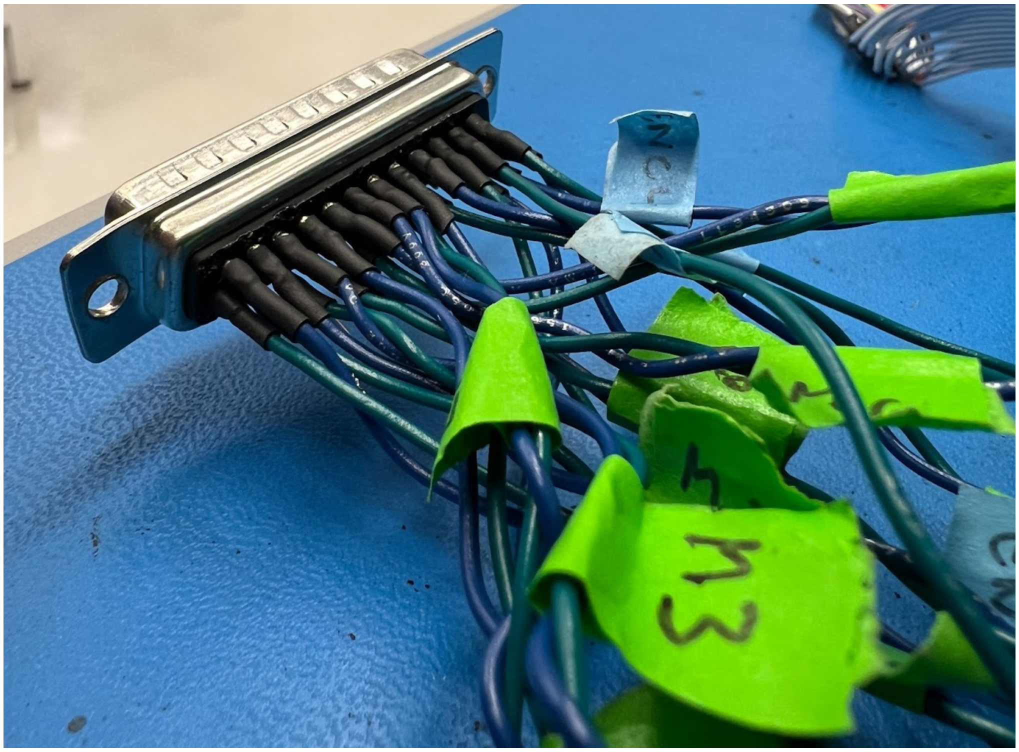


14. Screw the DB25 and DB15 connectors onto the 3-0712 rear access plate with #4x3/8” round head pan head machine screws and #4 nuts.

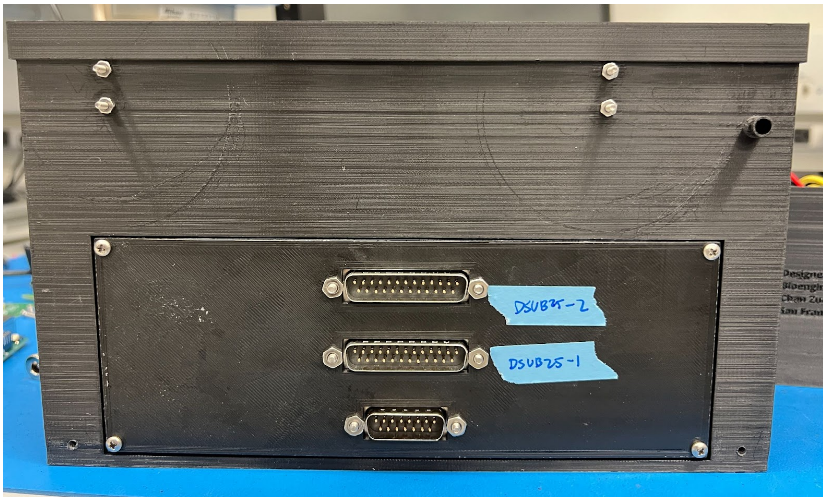


15. Place the 3-0713 rear gasket and 3-0712 rear access plate onto the back of the 3-0282-18T housing and screw in place with #4x1/2” round head Phillips thread-forming screws.
16. Sandwich the 3-0283 test tube guide, 3-0284 rubber insert, and 3-0285 top plate on 3-0282-18T tube housing and screw in place with #3x3/8” round head Phillips thread-forming screws.
17. Insert the 3-0774 tube housing location block into the upper rail left-most position (when facing the CZ Biohub logo).

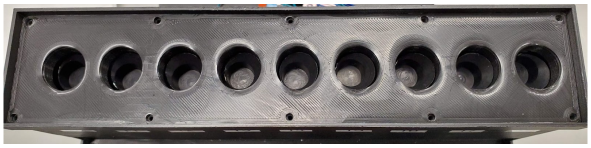
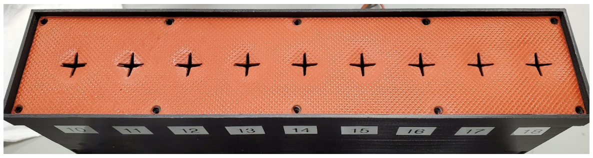
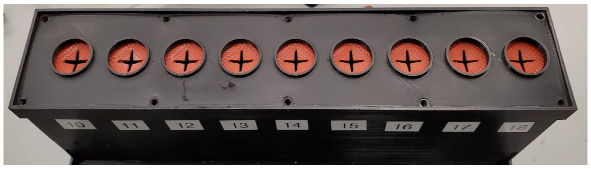
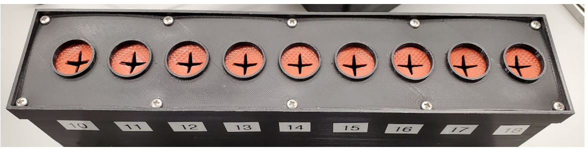
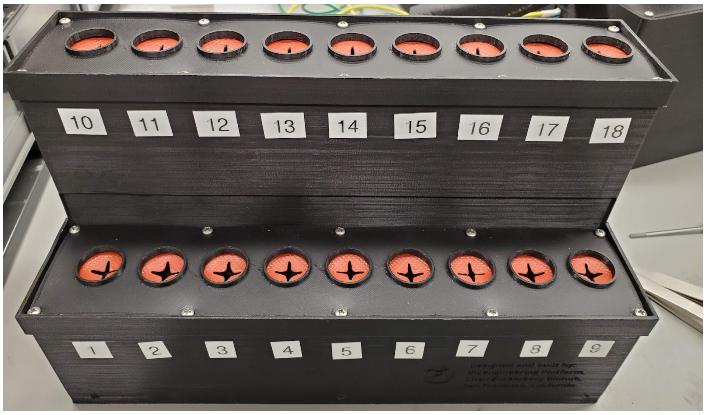

#### 2-0132 FACS Automation Electronics Box

##### 3-1095 FACS Electronics Box Case

1. Print the FACS electronics box case with the bottom on the build plate.
2. Print in Tough PLA with PVA support material with the Ultimaker S5.
3. Wash in the water bath to dissolve the PVA support material.
4. Install the threaded 3x M3 inserts on the inside of the bottom face of the box using the soldering iron to heat set them. Refer to the leftmost picture below for reference.
5. Add 3x M3 M-F standoffs to the inserts.
6. Mount the Arduino Mega on top of the standoffs and attach with 3x M3x0.5x5 hex screws.
7. Align headers of the 5-004 FACS PCB v2 to the Arduino Mega and push into place.

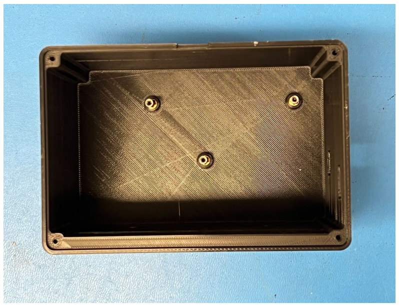
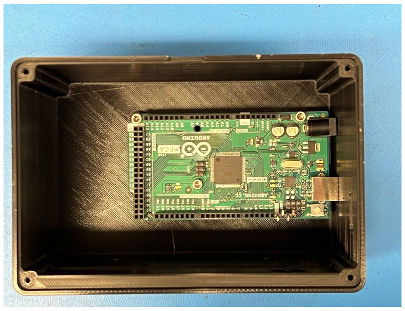
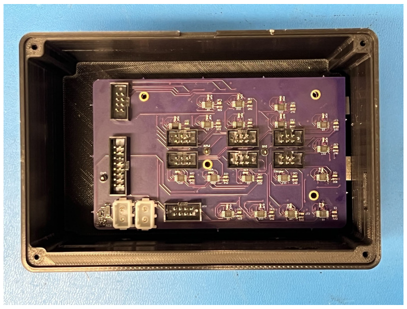

##### 3-1096 FACS Electronics Box Lid

1. Print the FACS electronics box lid with the top on the build plate.
2. Print in Tough PLA without support material with the Ultimaker S5.
3. Attach the lid to 3-1095 FACS Electronics Box Case using 4x #3x3/8” round head Phillips thread forming screws.

##### 3-1209 Electronics Box Adapter Strip

The 3-1209 Electronics Box Adapter Strip is used whenever an adjustment is needed between the location of the electronics box to the L-brackets mounted on the automation alignment plate. This may be needed if the ribbon cables are a little short.

1. Laser cut the strip from 6-6.35 mm thick acrylic with the standard laser cutter settings for acrylic.
2. Tap the holes with the ¼”-20 tap.
3. Loosely attach the adapter strip to the exterior of 3-1095 FACS Electronics Box Case using the two slots and ¼”-20 x 0.5”socket head screws and washers.

#### 3-0674 Solenoid Adapter Plate

1. Print the solenoid adapter plate with the counterbore surface as the top surface.
2. Print in Tough PLA without support material with the Ultimaker S5.
3. Install the threaded inserts using the soldering iron to heat set them.
4. Mount the Asco Red Hat solenoid valve to the adapter plate with the M5x16 mm screws in the counterbore holes.

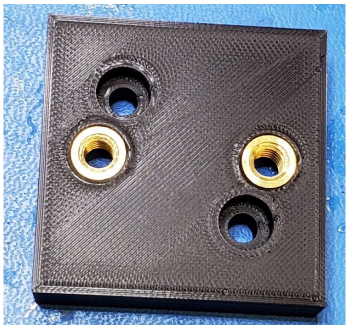
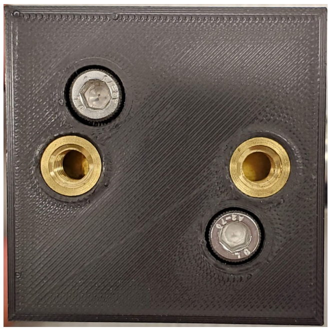
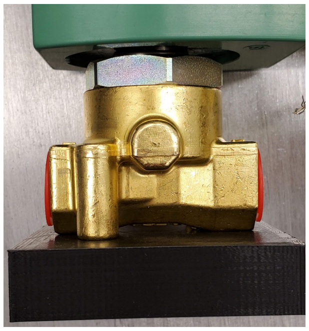

#### 3-0627 Air Filter Holder Base

1. Print the air filter holder base with the bottom on the build plate.
2. Print in Tough PLA without support material with the Ultimaker S5.

#### 3-0722 Air Filter Holder Top

1. Print the air filter holder top in Tough PLA with PVA support material with the Ultimaker S5. The top and bottom were split to decrease the time to print and reduce the amount of PVA support material.
2. Wash in the water bath to dissolve the PVA support material.

3. Screw 3-0627 air filter holder base and 3-0722 air filter holder top together with #4x3/4" round head Phillips thread forming screws.
4. Insert the air filter such that it rests atop the holder.

### 2-0128 FACS Automation Cold Air System

#### 3-0541 Air Gun Adapter

1. Print the air gun adapter in Tough PLA with PVA support material with the Ultimaker S5.
2. Wash in the water bath to dissolve the PVA support material.
3. The piping portion may need some clearing after soaking with a drill bit (No. 76 or smaller) or a small tool to remove the dissolving PVA support material.
4. Return the part to the water bath to fully eliminate all of the PVA support material.

#### 3-0314 Air Gun Adapter Sleeve

1. Print the air gun adapter sleeve in Tough PLA with PVA support material with the Ultimaker S5.
2. Wash in the water bath to dissolve the PVA support material.
3. Install the 4xM3 threaded inserts using the soldering iron to heat set them.

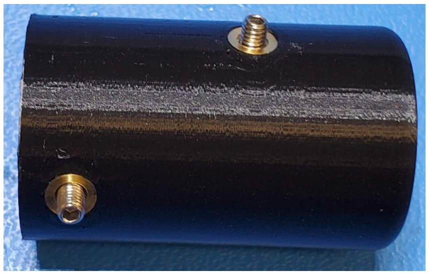

1. Remove the flexible nozzle assembly from the cold air gun to access the cold end fitting.
2. Push the 3-0541 air gun adapter over the cold air fitting.
3. The inner diameter of the 3-0541 air gun adapter may need to be drilled open with a 31/34" drill bit.
4. Slide the 3-0314 air gun adapter sleeve over the 3-0541 air gun adapter.
5. Screw the M3 set screws into place in the threaded inserts in the 3-0541 air gun adapter. Two of the M3 set screws insert into two holes in the side of the air gun exhaust. The other two M3 set screws impinge on the 3-0314 air gun adapter sleeve.

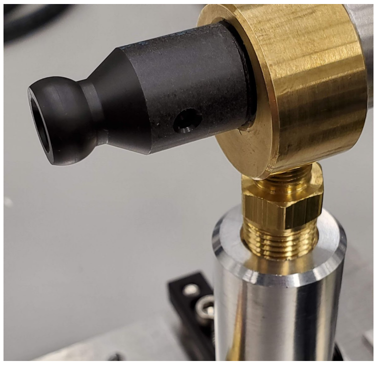
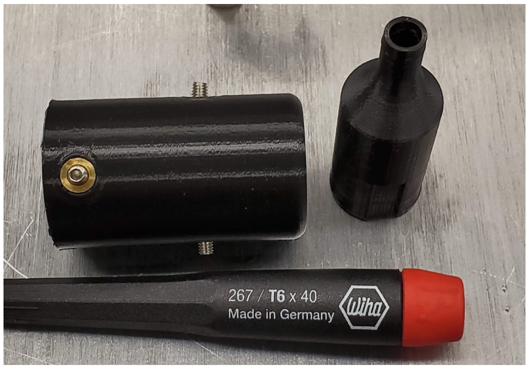

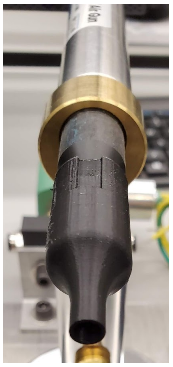
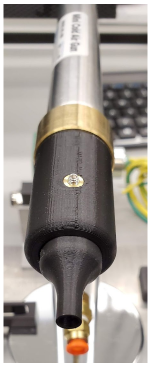

#### Attachment of subassemblies to plate

1. Mount the 2-0059 FACS Zaber stage sub-assembly to the 2-0119 FACS automation alignment plate aligning the Zaber LSQ450X-E01T4A-ENG2710 stage with the 3 alignment dowel pins on the plate.
2. Screw the stage in place with ¼”-20 x0.25” screws.

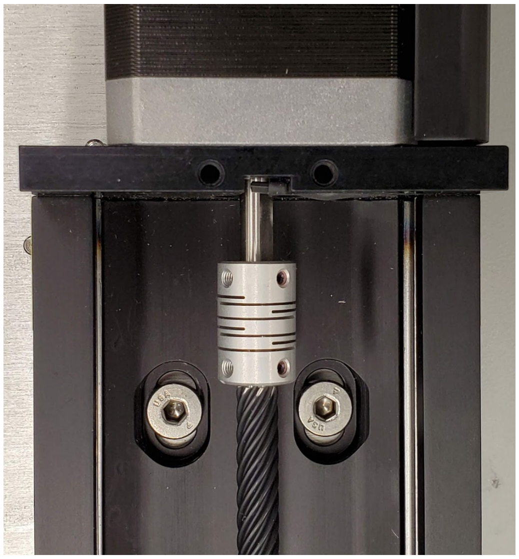
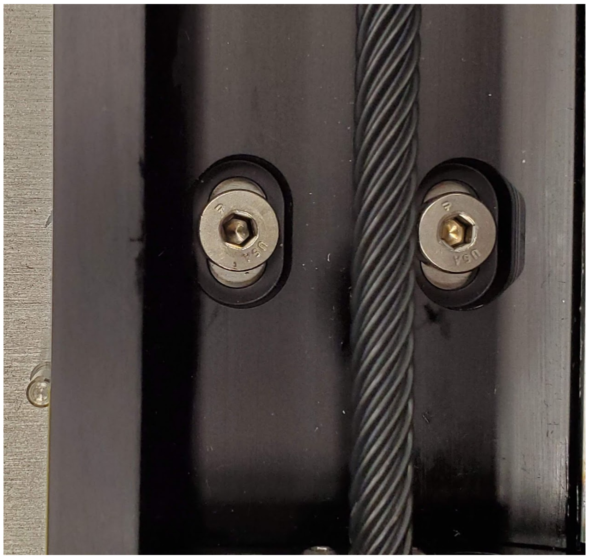

3. Attach the four McMaster-Carr L-brackets to the 3-0312 mounting plate with ¼”-20 x 3/8” screws.
4. Attach the 3-0674 solenoid attachment plate to the rear two L-brackets with ¼”-20 x 0.5” screws.

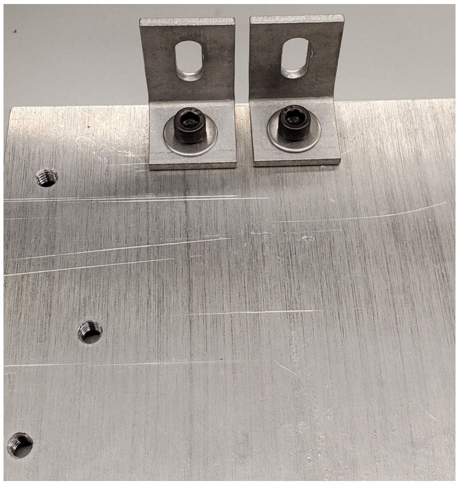
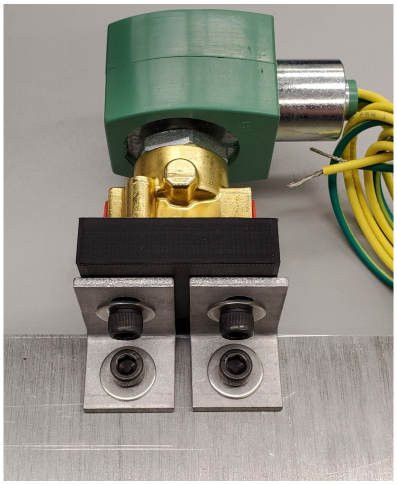

5. Screw the 2-0132 FACS Automation Electronics Box to the two side L-brackets with ¼”-20 x 0.375” screws into the tapped holes of the 3-1209 Electronics Box Adapter Strip.
6. Adjust the screws in the slots of the adapter strip as needed.

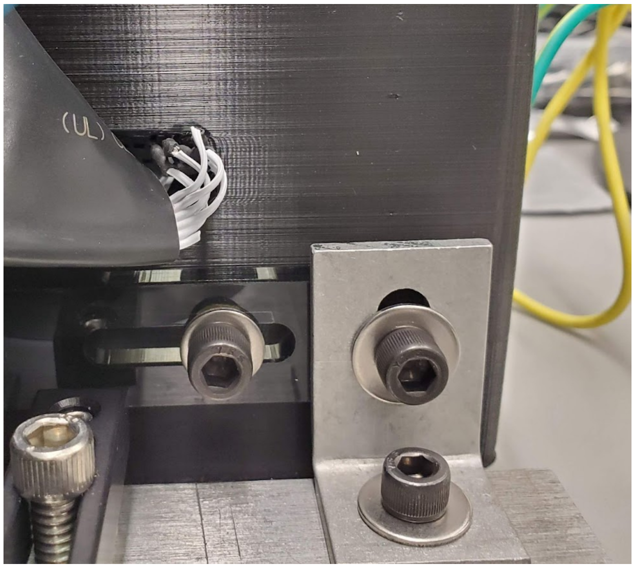

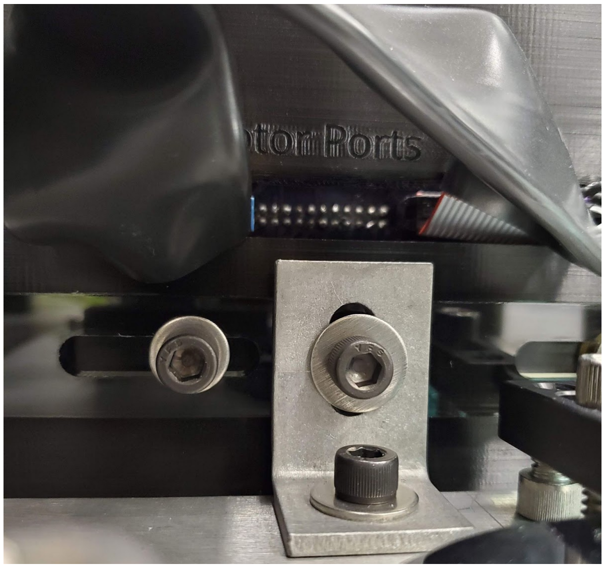

7. Screw the 3-0627 air filter base to the 3-0312 mounting plate with #6-32x0.5” screws.
8. Attach the regulator to the 3-0312 mounting plate with ¼”-20 x5/16” screws.

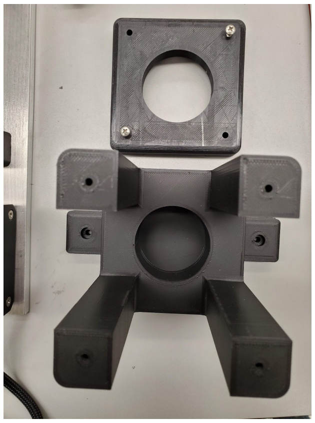
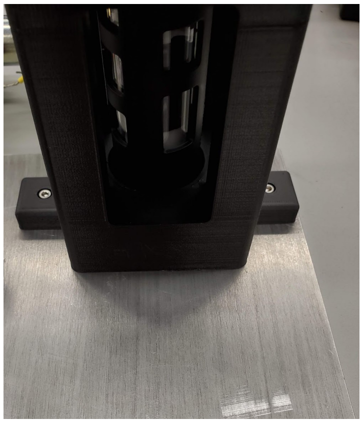

9. Screw the two Thorlabs CL3 clamps to the 3-0312 mounting plate with ¼”-20x1” in the clamp slot and ¼”-20 x 0.5” screws in the clamp threads.
10. Align the base of the 2-0128 cold air system such that the cold air exit faces toward the direction of the tube housing.
11. Lock the base in place with the Thorlabs CL3 clamps.

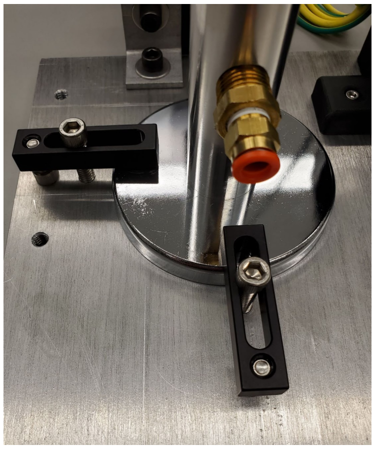
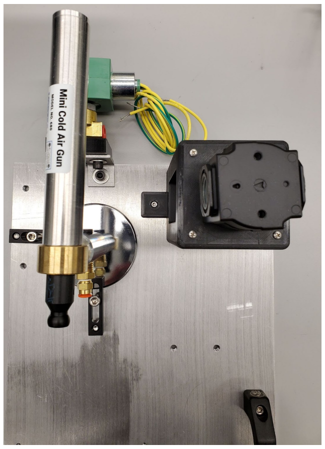

12. Screw in the push-to-connect to NPT adapters into both sides of the solenoid valve, both sides of the filter, and both sides of the regulator.
13. Cut the ¼” tubing into 6” to 1 foot pieces.
14. Insert the tubing to plumb from the solenoid valve into the filter, from the filter into the regulator, and from regulator into the air gun.
15. The regulator should be set to approximately 65 PSI, though it may vary based on the measured temperature inside of the tube housing.

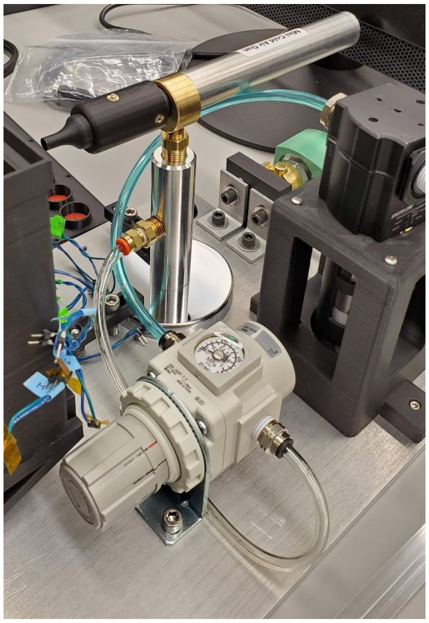
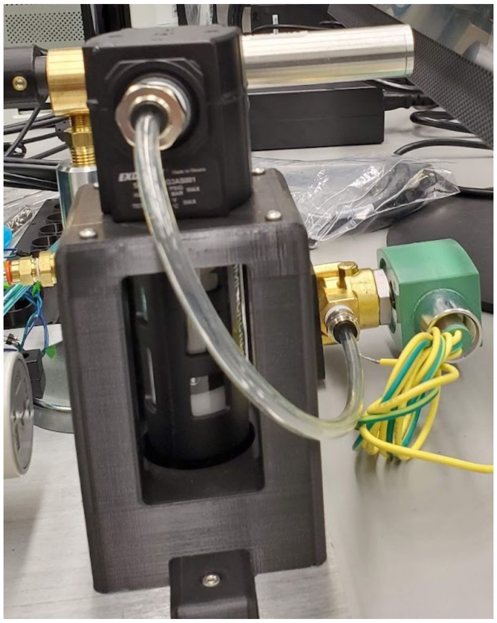

16. Attach the 3-0574 FACS tube holder adapter with #6-32 x0.5” screws. The positions of the adapter part can be adjusted in the slot during final assembly to ensure access to all of the required positions (tube housing and Sony sorter).

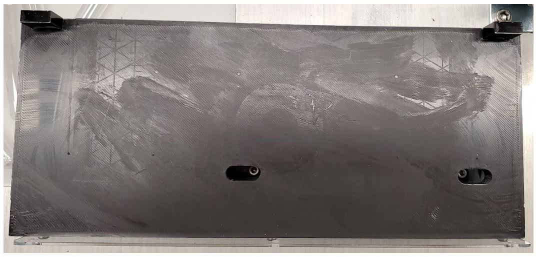
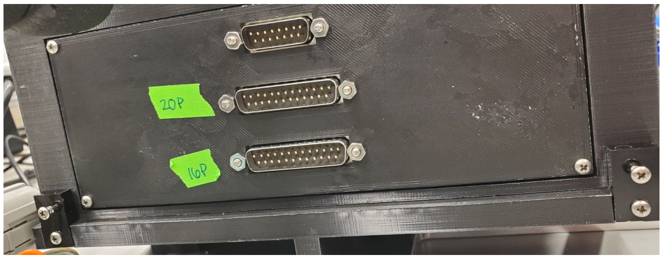


17. Attach the 2x 3-1116 tube housing attachment brackets to the 3-0574 FACS tube holder adapter with #4x1/2” round head Phillips thread forming screws.
18. Attach the 3-1168 tube housing front plate through the 3 bottom screw holes with #4x1/2” round head Phillips thread forming screws into the 3-0574 FACS tube holder adapter.
19. Slide the 2-0056 tube housing assembly onto the 3-0574 FACS tube holder adapter.
20. Screw in the 2x #4x1/2” round head Phillips thread forming screws into the top holes of the two 3-1116 tube housing attachment brackets and into the 2-0056 tube housing assembly.
21. Screw in the #4x1/2” round head Phillips thread forming screws into the top 3 screw holes of 3-1168 tube housing front plate and into the 2-0056 tube housing assembly.

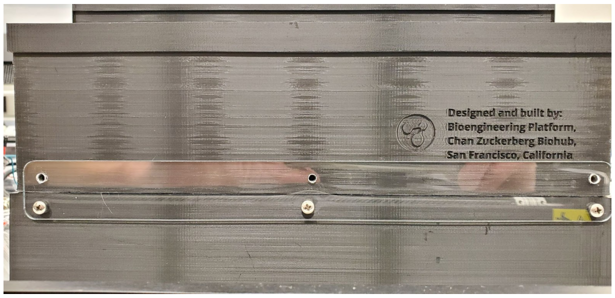

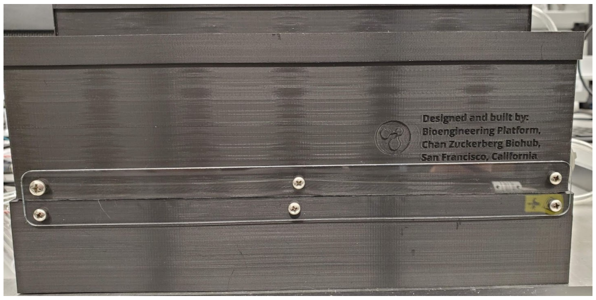


22. Assemble the tubing and insulation from the air gun to the tube housing.
23. Wrap the air gun adapter 3D printed parts in the insulation. This will require slicing the insulation open along one side with a razor blade.
24. Use zip ties to hold the insulation in place.
25. Attach a Tee push-to-connect fitting to the exit of the 3-0541 air gun adapter tube.
26. Connect 6” of ¼” tubing to each side of the T exit.
27. Wrap the tubing with insulation.

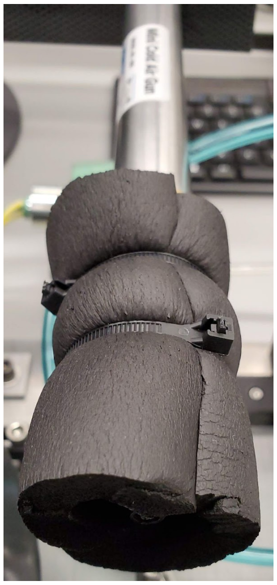
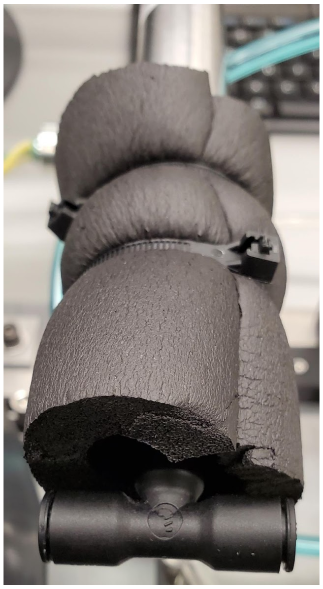
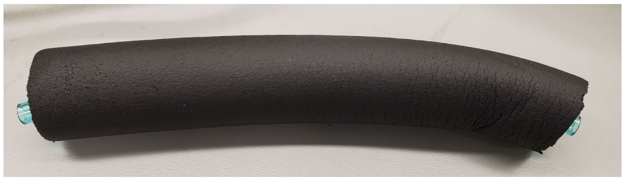

28. Connect the right exit tubing with a Wye push-to-connect fitting, and the left exit tubing with a Tee push-to-connect fitting.
29. Connect 6” of ¼” tubing to all 4 of the exits of the fittings and wrap the tubing with insulation.
30. Connect a 90 degree elbow fitting to all four of the tube exits.
31. Connect the four fittings to the tube housing cold air inlet ports on final installation.

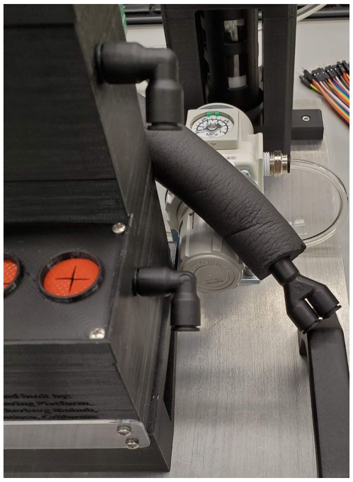
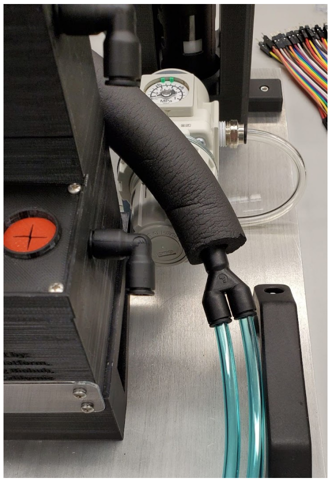
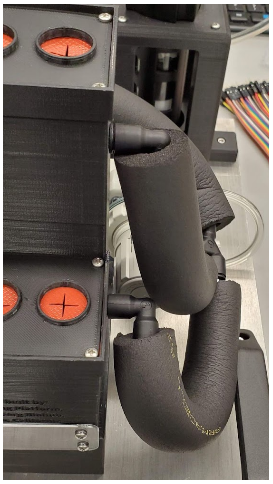
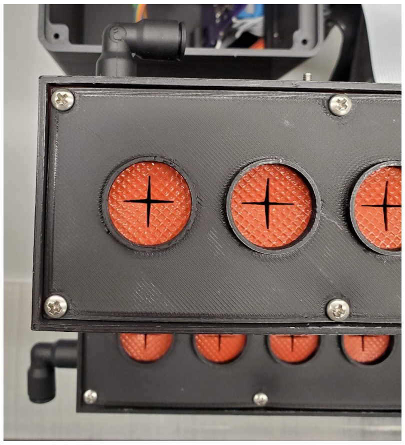
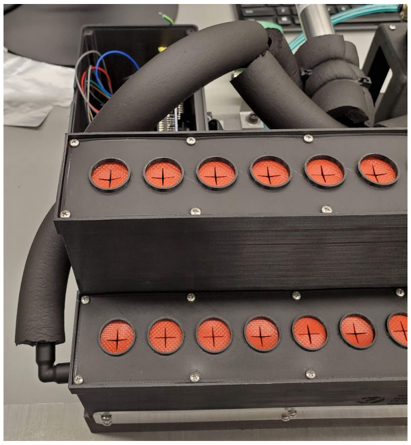

32. Screw the McMaster-Carr handles onto the 3-0312 mounting plate with #10-32 x3/4” screws.

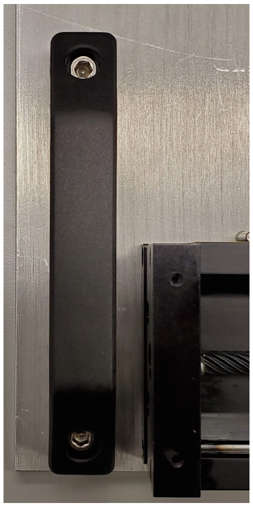
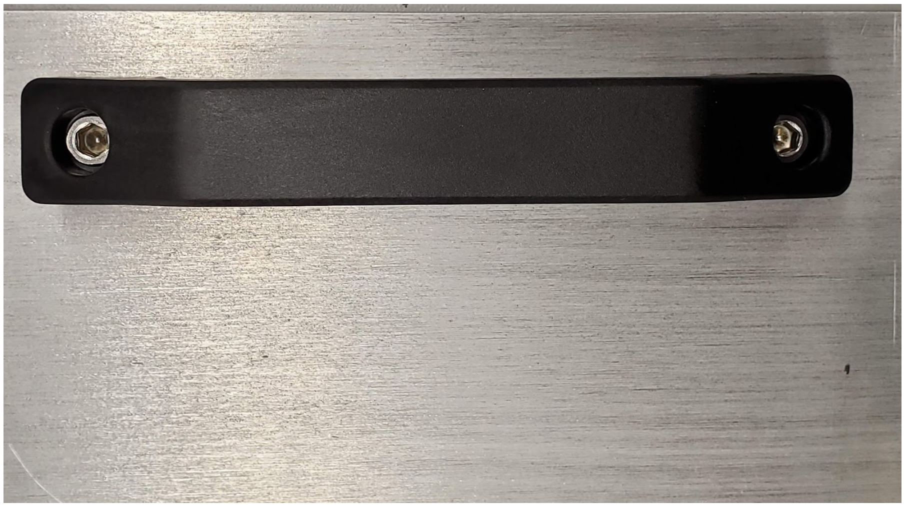

### Sub-assembly 2-0135 FACS Zaber Controller

#### Zaber X-MCC3 Universal Motor Controller

1. Connect each of the three linear axis stages and gripper to the Zaber Universal Motor Controller with the provided cables, and in the following order, as seen in the photo below:
    a. The X stage (LSQ450X-E01T4A-ENG2710) cable is inserted into the ‘Axis 1’ slot.
    b. The Y stage (LSQ075B-T4-ENG2690) cable is inserted into the ‘Axis 2’ slot.
    c. The Z stage (LSQ150B-T4A) cable is inserted into the ‘Axis 3’ slot.
    d. The gripper (X-GLP-E03) cable is inserted into the ‘Next’ slot.

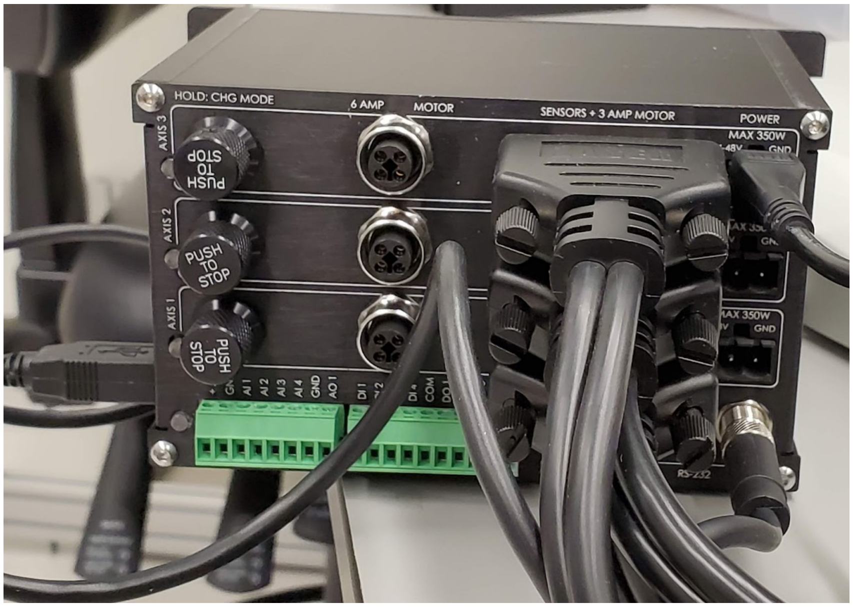


#### 3-0694 Motor Controller Plate

1. Print the motor controller plate in Tough PLA with no support material with the Ultimaker S5.
2. Install the 5x ¼”-20 threaded inserts using the soldering iron to heat set them.
3. Mount the Zaber X-MCC3 controller onto the plate with the ¼”-20 x0.5” screws.
4. Install the Eyebolt into the top threaded insert.
5. A hole must be drilled into the stainless steel side wall on the right hand side. Use a No 32 cobalt steel drill bit. Install the hook with the provided No. 5 sheet metal screw.

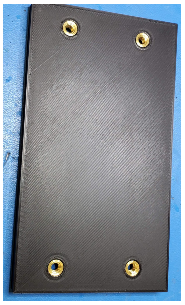 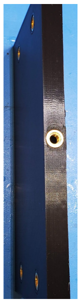

#### Zaber Controller Configuration

1. Download the Zaber Console Software onto a computer with a Windows operating system.
2. Perform the initial set-up using the Zaber Console Software.
3. Test moving the stages with the home button in the Zaber Console Software.
4. Cross-check the names of the Zaber stages with the names in the *zaber_controller.py* class. Successfully connected devices appear like below.

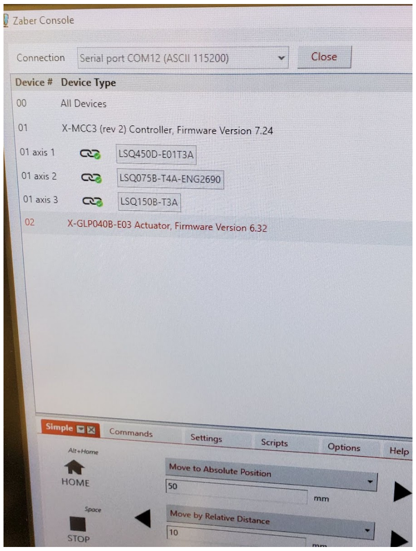


### Final assembly 1-0015 FACS Assembly

1. Pre-assemble the 2-0129 FACS Sony Clamps with the #6-32 x1.5” screws.
   a. The screw on the inner side of the clamp should be fastened slightly.
   b. The screw on the end should be free and inserted into the 3-0310-V1.1 primary clamp.
2. Do this for both copies of the clamps.

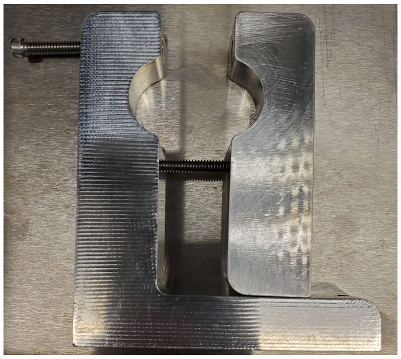

3. Remove the front plate of the Sony biosafety cabinet and roll the base of the cabinet forward.
4. Separate the primary and secondary clamps as far as possible and rotate the secondary clamp in order to envelop the locking nuts on the front foot of the Sony SH800.

5. Do this for both feet.

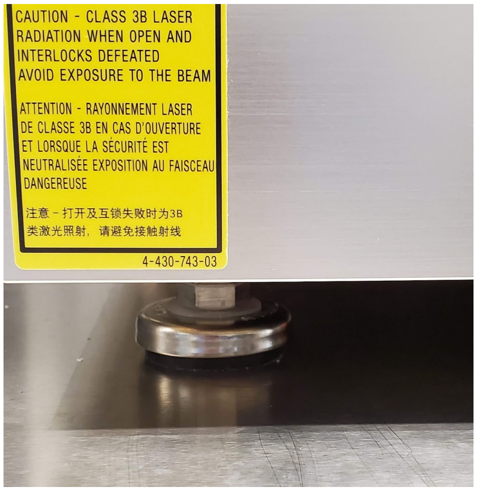

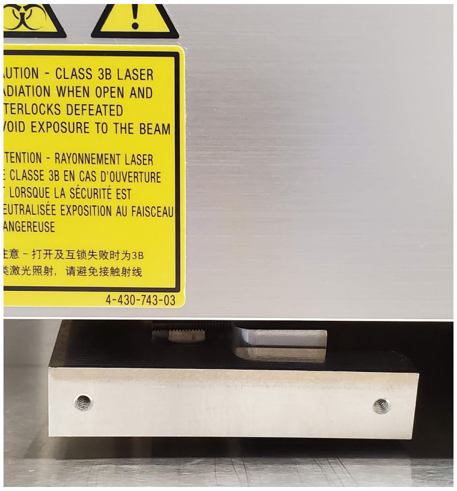

6. Loosely start to screw in both screws alternating between the two screws for both clamps.
7. Tighten the screws so that the primary clamp front face is planar to the Sony SH800.

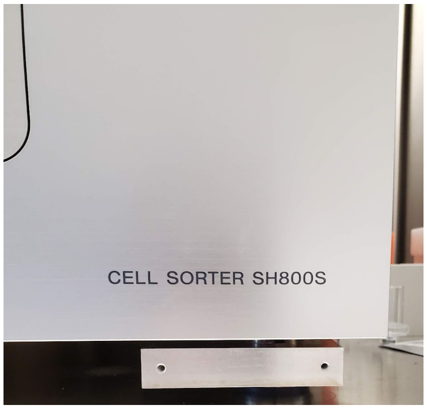

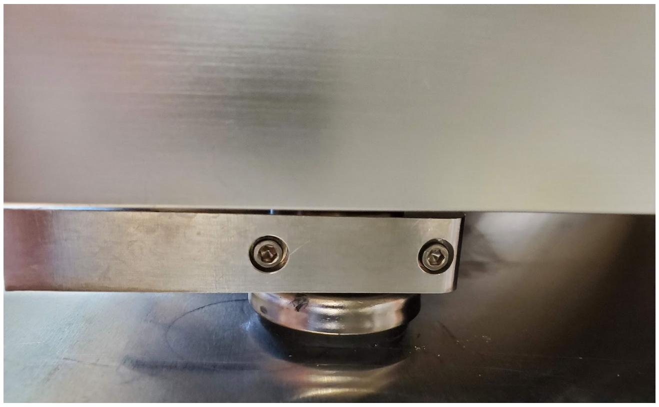
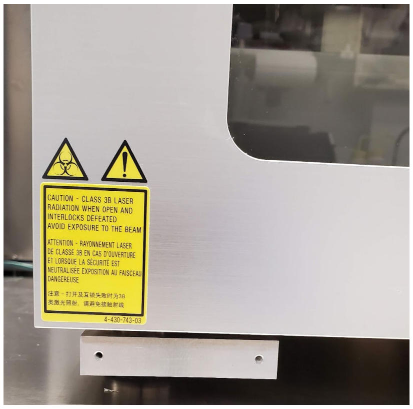
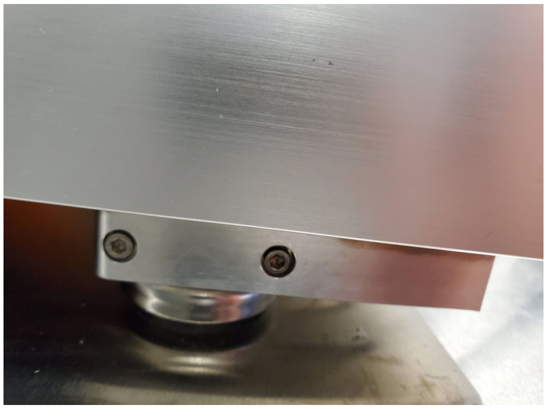

8. Position and screw in place the 3-0313-V1.1 clamp mounting block to the 2-0120 FACS Sony Clamps with #6-32 x0.625" screws.

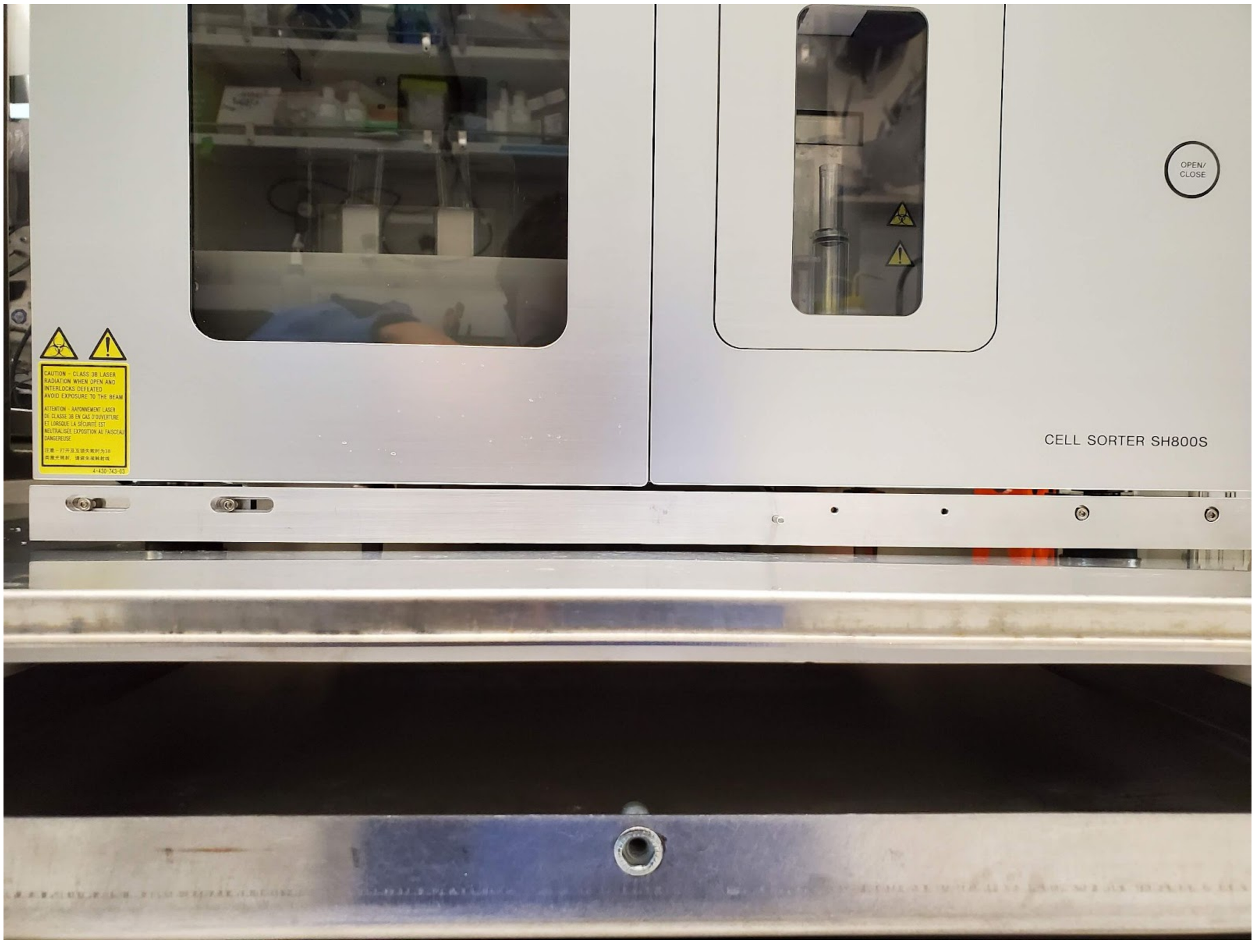
CAUTION - CLASS 3B LASER RADIATION WHEN OPEN AND INTERLOCKS DEFEATED AVOID EXPOSURE TO THE BEAM
ATTENTION - RAYONNEMENT LASER DE CLASSE 3B EN CAS D'OUVERTURE ET LORSQUE LA SÉCURITÉ EST NEUTRALISÉE EXPOSITION AU FAISCEAU DANGEREUSE
4-430-743-03
OPEN/ CLOSE
CELL SORTER SH800S

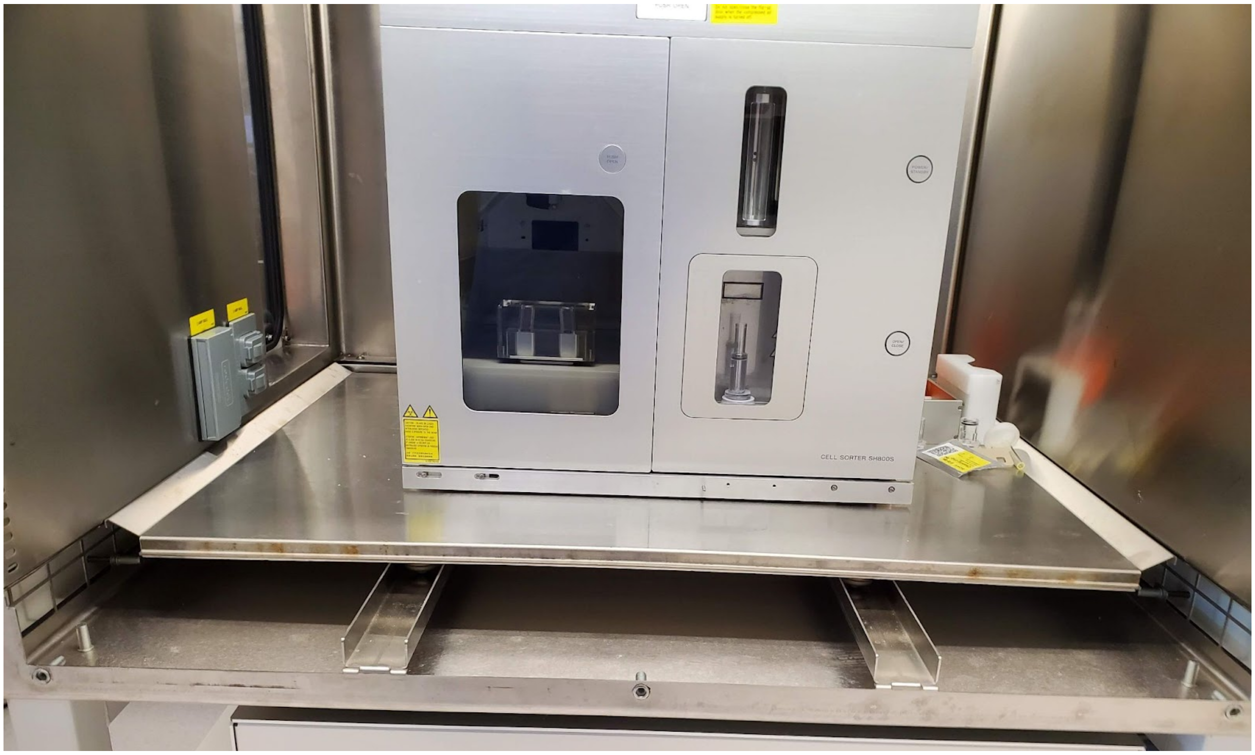
CELL SORTER SH800S

9. Move the 2-0126 Sony cell sorter and adapter as far to the left and as far back in the biosafety cabinet as possible to allow clearance for the 2-0119 FACS automation alignment plate.

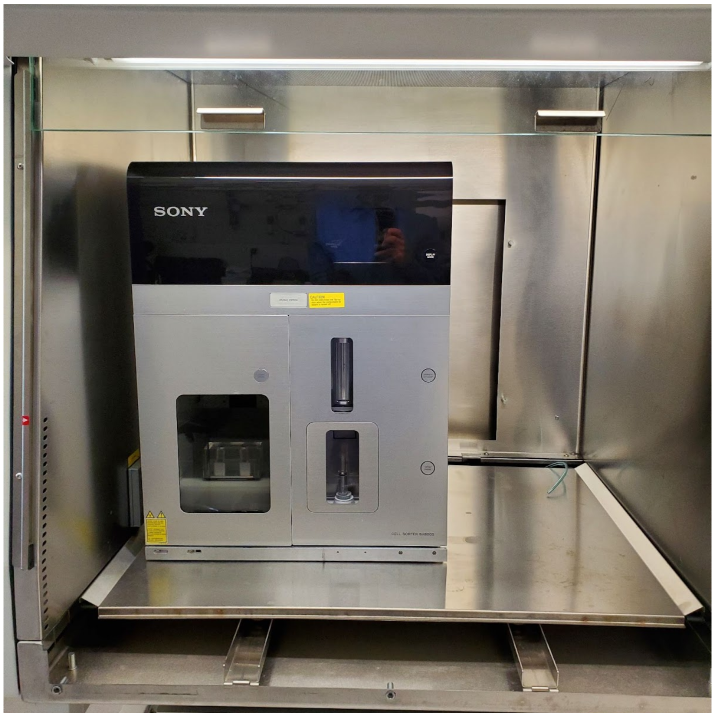


10. Insert the 2-0119 FACS automation alignment plate into the biosafety cabinet.
11. Place the 3-0628 L-bracket clamp onto 2-0119 FACS automation alignment plate.
12. Push against the dowel pin inserted into the 3-0313-V1.1 clamp mounting block and screw to the 2-0119 FACS automation alignment plate with 2x #6-32 x0.375" screws and to the 3-0313-V1.1 clamp mounting block with 2x #6-32x 7/16".
13. Center the 2-0056 tube holder with the through holes in the 3-1116 tube housing attachments brackets.
14. Check for clearance of all of the required positions by the Zaber stage.
15. Reposition 3-0574 FACS tube holder adapter, as needed.
16. Screw in place the 2-0056 tube holder with #4x1/2" round head Phillips thread forming screws.
17. Connect the 4x90 degree elbow ¼"push to connect fittings to the 4 cold air inlet ports on the 2-0056 Tube Housing.
18. Roll the base back into place and replace the front plate of the Sony biosafety cabinet.

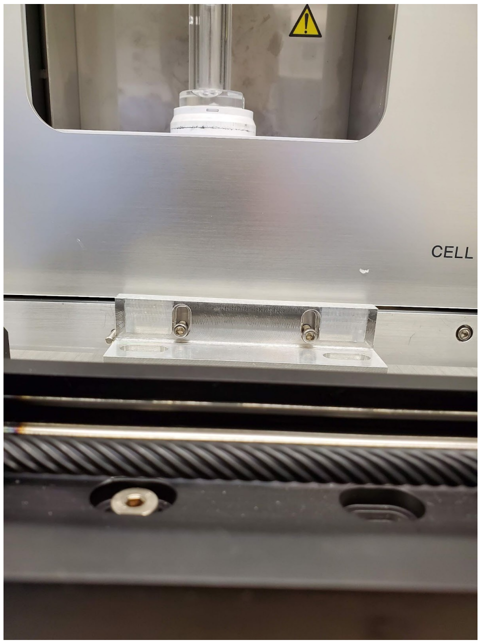

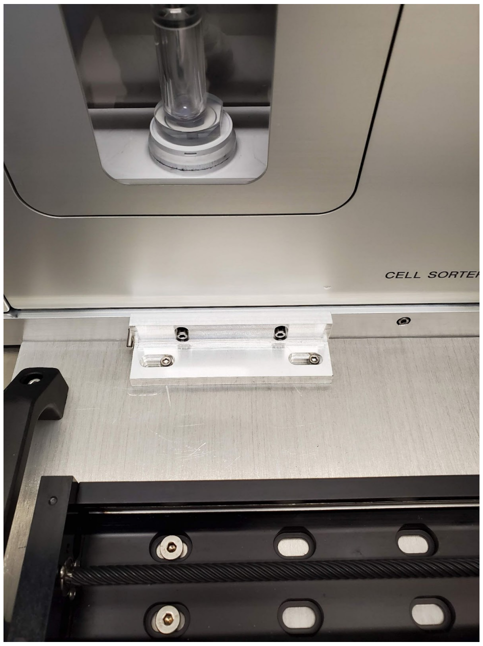

#### Compressed Air Tubing

Compressed air is required for the Vortec cold air gun.

1. From the house compressed air a ¼” tube is run to the biosafety cabinet for the Sony SH800.
2. Attach a push-to-connect on/off valve to the end of the ¼” tubing.
3. Remove the front plate of the Sony biosafety cabinet and roll the base of the cabinet forward.
4. Plumb ~5 feet of ¼” tubing through the left side port of the biosafety cabinet under the base and up into the main cabinet on the right side.
5. Roll the base back into place and replace the front plate of the Sony biosafety cabinet.
6. Attach the end of the tubing at the left side port to the push-to-connect on/off valve.
7. Connect the right end of the tubing to the solenoid valve attached to the 2-0119 FACS Automation Alignment Plate.
8. Connect ¼” tubing from the solenoid valve to the inlet of the air filter. The tubing is 6” to 12” long.
9. Connect ¼” tubing from the exit of the air filter to the inlet of the pressure regulator. The tubing is 6” to 12” long.
10. Connect ¼” tubing from the exit of the pressure regulator to the inlet of the air gun. The tubing is 6” to 12” long.

#### Power / Cable Management

The cabling for the Zaber stage control must not interfere with the free movement of the stage.

1. Use zip ties to bundle the cabling from the LSQ150B, LSQ075B, and X-GLP gripper together. Ensure that there is enough slack in the line for each stage.

2. Thread the 0.037” diameter 304 stainless steel wire (McMaster-Carr, 8908K38) through the grate in the ceiling of the biosafety cabinet.
3. Hang the 2-0135 FACS Zaber Controller from the wire hanger to the grate in the ceiling of the biosafety cabinet.
4. Thread another 304 stainless steel wire through another section of the grate in the ceiling of the biosafety cabinet. Use it to hold the bundle of cabling from the ceiling.

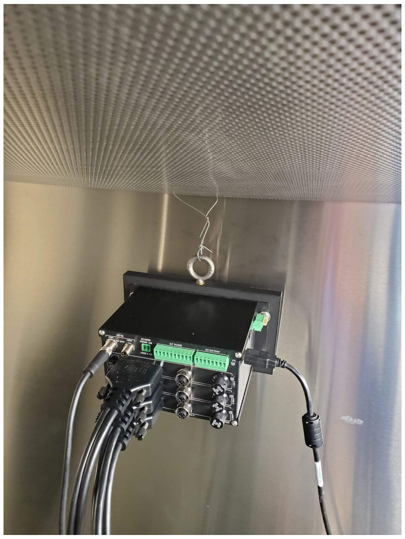
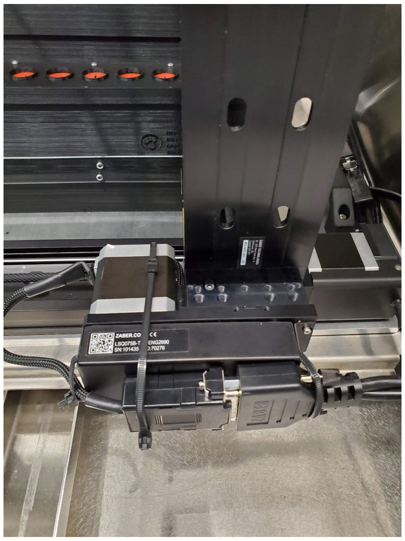
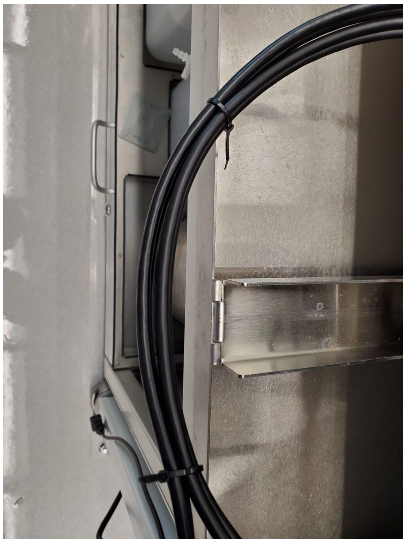
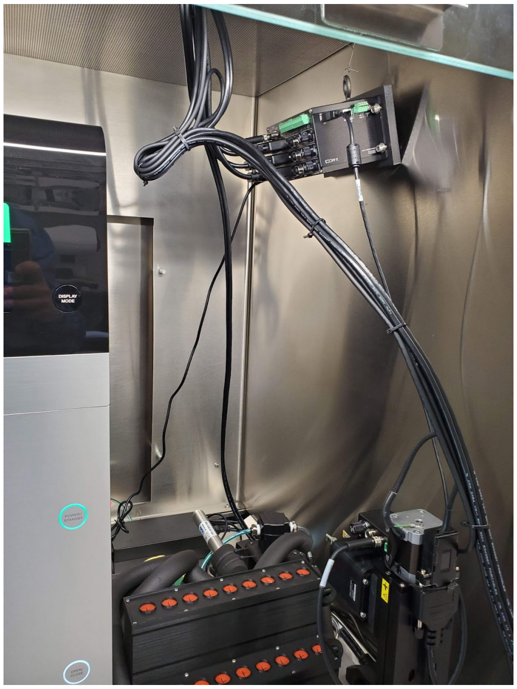
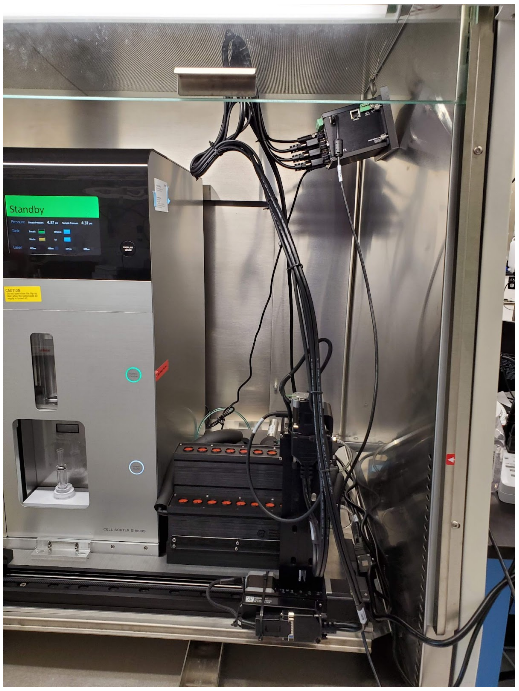


5. Affix the power strip for the Zaber controller and electronics box in a convenient location with Velcro sticky tape.
6. Use a strip of Velcro sticky tape to attach the USB adapter for the Zaber controller and electronics box in a convenient location. The USB adapter connects into the computer running the Sony SH800.

### Bill of Materials (BOM)

See the following documents:
S2 File FACS Hardware BOM
S3 File FACS PCB BOM

### Laser cutter & 3D printers

3D printed parts were printed in-house on an Ultimaker S5 FFF 3D printer, or Formlabs Form3 SLA 3D printer, as specified above and in the Hardware BOM.

All in-house laser cutting was done with a Universal Laser Systems PLS6.15D laser cutter.
