## Supplementary material for "An open-source FACS automation system for high-throughput cell biology": Hardware Bill of Materials

### Hardware BOM - FACS Automation

| Part number | Quantity | Vendor | Name | Description |
|---|---|---|---|---|
| **1-0015-V1.1 FACS Assembly** | | | **FACS Automation project** | **Integration of automated software and hardware to the Sony Sorter SH800 instrument. A 3-axis stage with gripper is used to move sample tubes from a cooling chamber housing to the Sony Sorter instrument.** |
| SH800S | 1 | Sony | Sony SH800S Cell Sorter | Sony SH800 FACS instrument rendering; not the official CAD file. |
| LE-U3M004 SYX | 1 | Sony | Sony LE-U3M004 SYX SH800 Sample Holder (5mL) | Tube adapter for the Sony SH800. It magnetically seats into the instrument, and holds the sample tube. Not official CAD file. |
| 3-0630 | 1 | 3D Printed (SLA) | 3-0630 Sony Adapter Funnel | Funnel used for FACS automation so that the grippers can deliver the tube into the Sony 0.5 mL tube adapter. |
| 352235 | 18 | Falcon | Falcon 352235 Culture Test Tube | 12x75 mm sample tube used with the Sony Sorter FACS instrument |
| 3-0613 | 1 | Laser Cut | 3-0613 Sony Sorter Acrylic Fluidics Maintenance Door | Acrylic door to replace the factory door of the Sony Sorter for the FACS Automation project. This is to allow for the clearance within the biosafety cabinet of the door to access the fluidics inside of the Sony Sorter. |
| 3-0757 | 1 | Outside Machine Shop | 3-0757 Sony Vacuum Plate Replacement | A replacement plate for the HEPA exhaust vent plate to add an L-shaped tube fitting. This allows for the Sony Sorter to be moved inside of the BSC so that the FACS automation hardware fits inside of the BSC. |
| 93475A230 | 12 | McMaster-Carr | Chamfered plain washer normal M4 | Chamfered plain washer normal M4 Stainless Steel |
| 91292A118 | 6 | McMaster-Carr | Hex socket head cap screw M4x0.70 x 16 | Hex socket head cap screw M4x0.70 x 16 Stainless Steel |
| 91828A231 | 12 | McMaster-Carr | Hex nut grade A & B M4x0.7 | Hex nut grade A & B M4x0.7 Stainless Steel |
| LSQ450x E01T4A ENG2710 | 1 | Zaber | Zaber LSQ450x E01T4A ENG2710 Linear Stage | Zaber Technologies 450 mm linear translation stage |
| LSQ075B-T4-ENG2690 | 1 | Zaber | Zaber LSQ075B-T4-ENG2690 | Zaber Technologies 75 mm linear stage |
| LSQ150B-T4A | 1 | Zaber | Zaber LSQ150B-T4A Customer Facing | Zaber Technologies 150 mm linear stage |
| X-GLP-E03 | 1 | Zaber | Zaber X-GLP-E03 | Zaber Technologies grippers. with 20 mm travel |
| 3-0297-TBO | 1 | Outside Machine Shop | 3-0297-TBO Top Claw Blunted Off-axis | Claw affixed to the Zaber gripper to grab and hold FACS sample tubes for the FACS automation project |
| 3-0297-BBO | 1 | Outside Machine Shop | 3-0297-BBO Bottom Claw Blunted Off-axis | Claw affixed to the Zaber gripper to grab and hold FACS sample tubes for the FACS automation project |
| 3-0312-V1.1 | 1 | Outside Machine Shop | 3-0312-V1.1 Mounting plate | The plate for mounting all of the hardware for the FACS Automation project. Version 1.1 |
| 3-0310-V1.1 | 2 | Outside Machine Shop | 3-0310-V1.1 Primary clamp finger | Primary clamp finger to attach and align the automation rig to the Sony sorter foot. V1.1 |
| 3-0311-V1.1 | 2 | Outside Machine Shop | 3-0311-V1.1 Secondary clamp | Secondary clamp, version 1.1 |
| 92196A168 | 4 | McMaster-Carr | Socket head cap screw #6-32 x 1.75 | Socket head cap screw #6-32 x 1.75 Stainless Steel |
| 3-0313-V1.1 | 1 | Outside Machine Shop | 3-0313-V1.1 Clamp mounting block | Block to align and mate between the Sony clamps and the automation plate, version 1.1 |
| 3-0574 | 1 | 3D printed (FFF) | 3-0574 FACS tube holder adapter | Adapter part between mounting plate and tube holder to align the tube holder at the appropriate location for FACS automation |
| 92196A148 | 4 | McMaster-Carr | Socket head cap screw #6-32 x 0.5 | Socket head cap screw #6-32 x 0.5 Stainless Steel |
| 2471N2 | 2 | McMaster-Carr | McMaster 2471N2 Pull Handle | McMaster-Carr pull handle with mounting hole center-to-center width of 3 15/16", fits M5 screws, and load capacity 220 lbs. |
| 92196A272 | 4 | McMaster-Carr | Socket head cap screw #10-32 x 0.75 | Socket head cap screw #10-32 x 0.75 Stainless Steel |
| 680 | 1 | Vortec | Vortec 680 Mini Cold Air Gun and Filter | Vortec Mini Air Gun and Filter |
| 3-0541 | 1 | 3D printed (FFF) | 3-0541 Air Gun Adapter | 3D printed part to interface between the Vortec mini air gun exhaust and 1/4" quick release fittings |
| 3-0314 | 1 | 3D printed (FFF) | 3-0314 Air Gun Adapter Sleeve | The sleeve mates between the air gun and the air gun adapter to make a more rigid connection to them. |
| 94180A331 | 7 | McMaster-Carr | McMaster 94180A331 M3 threaded insert | M3 threaded insert |
| 91217A070 | 4 | McMaster-Carr | McMaster 91217A070 M3 Set Screw | M3x0.5x8 mm set screw |
| CL3 | 2 | Thorlabs | Thorlabs CL3 Compact Variable Height Clamp | Thorlabs CL3 Compact Variable Height Clamp for Optical Tables and 1/4"-20 screws |
| 92196A542 | 2 | McMaster-Carr | Socket head cap screw 1/4-20 x 1 | Socket head cap screw 1/4-20 x 1 Stainless Steel |
| 92196A537 | 10 | McMaster-Carr | Socket head cap screw 1/4-20 x 0.5 | Socket head cap screw 1/4-20 x 0.5 Stainless Steel |
| 3-0628 | 1 | Outside Machine Shop | 3-0628 L-bracket clamp to plate | L-bracket to align and secure the 1/4" mounting plate to the bracket clamped to the Sony Sorter for the FACS Automation project. |
| 92196A146 | 2 | McMaster-Carr | Socket head cap screw #6-32 x 0.375 | Socket head cap screw #6-32 x 0.375 Stainless Steel |
| 3-0627 | 1 | 3D printed (FFF) | 3-0627 Air Filter Holder Base | Base of housing for the air filter suppled with the Vortec cold air gun to keep the filter upright, and provide mounting locations. |
| AR30-N03BG-Z-B | 1 | SMC | SMC AR30-N03BG-Z-B Pressure Regulator | Air regulator |
| 97395A451 | 4 | McMaster-Carr | McMaster 97395A451 0p125in dia. x 0p75in long SS Dowel Pin | Dowel pin, 1/8" dia, 3/4" long, 316 stainless steel |
| 2313N793 | 4 | McMaster-Carr | McMaster 2313N793 CORNER MACHINE BRACKET | L-bracket using 1/4" screws. Used in the FACS automation project to screw the electronics box to the mounting plate off of the side of the plate. |
| 92196A535 | 6 | McMaster-Carr | Socket head cap screw 1/4-20 x 0.375 | Socket head cap screw 1/4-20 x 0.375 Stainless Steel |
| 92196A534 | 2 | McMaster-Carr | Socket head cap screw 1/4-20 x 0.3125 | Socket head cap screw 1/4-20 x 0.3125 Stainless Steel |
| 3-0674 | 1 | 3D printed (FFF) | 3-0674 Solenoid Adapter Plate | Plate to mount the solenoid valve to a bracket that affixes to the mounting plate. The hole spacing for the solenoid is unique such that a standard L-bracket will not suffice without the adapter. |
| 8262H022 | 1 | Asco Red Hat | Asco Red Hat 8262H022 Solenoid Valve | Solenoid valve, normally close, 12 V DC, Cv 0.35. |
| 91292A126 | 2 | McMaster-Carr | Hex socket head cap screw M5x0.80 x 16 | Hex socket head cap screw M5x0.80 x 16 Stainless Steel |
| 93365A160 | 10 | McMaster-Carr | McMaster 93365A160 Heat-Set Inserts For Plastics | 1/4"x20 threaded insert |
| 3-1095 | 1 | 3D printed (FFF) | 3-1095 FACS Electronics Box Case | Case to hold the Arduino Mega and FACS PCB |
| 3-1096 | 1 | 3D printed (FFF) | 3-1096 FACS Electronics Box Lid | Lid to case to hold the Arduino Mega and FACS PCB |
| 99461A825 | 24 | McMaster-Carr | McMaster 99461A825 - SS thread-forming screw, round head, PH drive, #3 - 3/8" long | Stainless steel thread-forming screw for plastics, round head, Phillips drive, #3 - 3/8" long. Drill diameter: 0.081" |
| 93655A220 | 3 | McMaster-Carr | McMaster-Carr 93655A220 Male-Female Threaded Hex Standoff M3 | 18-8 Stainless Steel, 4.500 mm Hex, 11 mm Long, M3 x 0.50 mm Thread |
| 91292A110 | 3 | McMaster-Carr | Hex socket head cap screw M3x0.50 x 5 | Hex socket head cap screw M3x0.50 x 5 Stainless Steel |
| A08000 | 1 | Digikey | Arduino Mega | The 8-bit board with 54 digital pins, 16 analog inputs, and 4 serial ports |
| 3-1209 | 1 | Laser Cut | 3-1209 Electronics Box Adapter Strip | Adapter strip for 1/4"-20 screws to fasten the electronics box to the L-brackets on the FACS automation plate due to the limited length of the ribbon cables |
| 5-2004 | 1 | OSH Park | 5-0004 FACS PCB v2 | Main PCB: H-bridge motor drivers, a transistor for solenoid valve, and circuitry for thermistors |
| 3-0722 | 1 | 3D printed (FFF) | 3-0722 Air Filter Holder Top | Top of housing for the air filter suppled with the Vortec cold air gun to keep the filter upright, and provide mounting locations. |
| 99461A150 | 4 | McMaster-Carr | McMaster 99461A150 - SS thread-forming screw, round head, PH drive, #4 - 3/4" long | Stainless steel thread-forming screw for plastics, round head, Phillips drive, #4 - 3/4" long. Drill diameter: 0.086" |
| 3-1116 | 2 | Laser Cut | 3-1116 Tube housing attachment bracket | Bracket to attach the tube housing to the the tube holder cantilever that attaches to the automation alignment plate |
| 3-0281 | 2 | 3D printed (FFF) | 3-0281 Motor plate | FACS automation plate interface for DC motors to seat within the housing |
| 3-0282-18T | 1 | 3D printed (FFF) | 3-0282-18T Housing | FACS automation housing for the sample tubes, DC motors, coupling parts and cooling. This configuration holds 18 tubes. |
| 3-0283 | 2 | 3D printed (FFF) | 3-0283 Test tube guide | FACS automation part to support test tubes within the tube housing |
| 3-0284 | 2 | Laser Cut | 3-0284 Rubber insert | FACS automation rubber gasket for tube holder to reduce the heat transfer between the cooled tube holder, and ambient conditions |
| 3-0285 | 2 | 3D printed (FFF) | 3-0285 Top plate | FACS automation top plate to close the tube housing and clamp the test tube guide and rubber insert in place. |
| DB25-SF-M1 | 2 | Digikey | Digikey DB25-SF-M1 | DB25 connector from Adam Tech |
| 3-0713 | 1 | Laser Cut | 3-0713 Rear Gasket | A gasket between the rear access plate and the tube housing to limit any cold air from escaping. |
| 3-0712 | 1 | 3D printed (FFF) | 3-0712 Rear Access Plate | A plate to provide access to the back of the tube holder for the FACS automation project |
| 3-0280 | 18 | Outside Machine Shop | 3-0280 Test Tube Socket | Socket interface for the DC motor to rotate test tubes for the FACS automation instrument. |
| 3038 | 18 | Pololu | Pololu Micro Metal Gearmotor HPCB | High-power carbon brush motor |
| 92000A116 | 18 | McMaster-Carr | PH Pan head screw M3x0.50 x 6 mm | PH Pan head screw M3x0.50 x 6 Stainless Steel |
| 98689A112 | 18 | McMaster-Carr | Chamfered plain washer normal M3 | Chamfered plain washer normal M3 Stainless Steel |
| 97259A101 | 18 | McMaster-Carr | McMaster 97259A101 M3 Low-Strength Steel Thin Square Nut | M3 Square Nut Stainless Steel |
| 91292A260 | 36 | McMaster-Carr | Hex socket head cap screw M1.6x0.35 x 3 mm | Hex socket head cap screw M1.6x0.35 x 3 Stainless Steel |
| 99461A815 | 20 | McMaster-Carr | McMaster 99461A815 - SS thread-forming screw, round head, PH drive, #3 - 1/4" long | Stainless steel thread-forming screw for plastics, round head, Phillips drive, #3 - 1/4" long. Drill diameter: 0.081" |
| 92196A108 | 6 | McMaster-Carr | Pan head machine screw #4-40 x 0.375 | Pan head machine screw #4-40 x 0.375 Stainless Steel |
| 99461A130 | 14 | McMaster-Carr | McMaster 99461A130 - SS thread-forming screw, round head, PH drive, #4 - 1/2" long | Stainless steel thread-forming screw for plastics, round head, Phillips drive, #4 - 1/2" long. Drill diameter: 0.086" |
| 5-2007 | 2 | OSH Park | 5-0007 FACS Photointerruptor Breakout Board | The photointerruptor board used in the FACS Automation tube housing to stop the motors at a specific location for ease of tube grabbing. |
| 3-0614 | 8 | Laser Cut | 3-0614 Motor Encoder PCB Spacer | Spacer to ensure the optical reflectors are within the distance of detection of the tube sockets to be used as motor encoders for the FACS Automation project. |
| 92196A140 | 4 | McMaster-Carr | Socket head cap screw #2-56 x 1 | Socket head cap screw #2-56 x 1 Stainless Steel |
| 91841A003 | 8 | McMaster-Carr | Hex machine screw nut #2-56 | Hex machine screw nut #2-56 Stainless Steel |

| | | | | |
|---|---|---|---|---|
| 92196A083 | 4 | McMaster-Carr | Socket head cap screw #2-56 x 0.625 | Socket head cap screw #2-56 x 0.625 Stainless Steel |
| 215ME-ND | 1 | Digikey | Digikey 215ME-ND | 15 Position D-Sub Plug, Male Pins Connector |
| 91841A005 | 6 | McMaster-Carr | Hex machine screw nut #4-40 | Hex machine screw nut #4-40 Stainless Steel |
| 92196A147 | 2 | McMaster-Carr | Socket head cap screw #6-32 x 0.4375 | Socket head cap screw #6-32 x 0.4375 Stainless Steel |
| 92196A150 | 4 | McMaster-Carr | Socket head cap screw #6-32 x 0.625 | Socket head cap screw #6-32 x 0.625 Stainless Steel |
| 3-1168 | 1 | Laser Cut | 3-1168 Tube housing front plate | A front cross plate to keep the tube housing mounted to the tube holder adapter |
| 92196A533 | 4 | McMaster-Carr | Socket head cap screw 1/4-20 x 0.25 | Socket head cap screw 1/4-20 x 0.25 Stainless Steel |
| X-MCC3 | 1 | Zaber | Zaber X-MCC3 Universal Motor Controller | Motor controller for Zaber stages. |
| 3-0694 | 1 | 3D printed (FFF) | 3-0694 Motor controller plate | Plate to mount the Zaber motor controller and hang from the sidewall of the FACS automation BSC. |
| 3013T45 | 1 | McMaster-Carr | McMaster 3013T45 Steel Eyebolt Without Shoulder | Steel eyebolt for lifting up to 500 lbs. |
| 5648K69 | 12 ft | McMaster-Carr | Firm Polyurethane Tubing for Air and Water, 5/32" ID, 1/4" OD | Tubing for compressed air supply to cold air gun |
| 5779K669 | 1 | McMaster-Carr | Push-to-Connect Tube Fitting for Air, Inline Tee Reducer, for 1/4" x 3/8" Tube OD | Reducing fitting for compressed air tubing |
| 5779K34 | 2 | McMaster-Carr | Push-to-Connect Tube Fitting for Air, Tee Connector, for 1/4" Tube OD | T connector fitting for compressed air tubing |
| 5779K44 | 1 | McMaster-Carr | Push-to-Connect Tube Fitting for Air, Wye Connector, for 1/4" Tube OD | W connector fitting for compressed air tubing |
| 5779K24 | 4 | McMaster-Carr | Push-to-Connect Tube Fitting for Air, 90 Degree Elbow Connector, for 1/4" Tube OD | 90 deg connector fitting for compressed air tubing |
| 45975K45 | 1 | McMaster-Carr | Gray PVC On/Off Valve, Straight, Push-to-Connect for 1/4" Tube OD, Full Port | On-Off valve for compressed air tubing |
| 5779K109 | 7 | McMaster-Carr | Push-to-Connect Tube Fitting for Air, Straight Adapter, for 1/4" Tube OD x 1/4 NPT Male | NPT to tubing connector for compressed air tubing |
| IPAPT03838 | 4 ft | Armacell | AP Armaflex Insulation, ⅜" wall thickness, 1/4" Copper | Insulation for cold air tubing |
| 8908K38 | 1 pack | McMaster-Carr | 304 Stainless Steel Wire, 0.037" dia x 12" | 2 wires needed from pack |
| 5777T11 | 1 | McMaster-Carr | Silicone rubber sheet with crisscross texture, 50A durometer, 1/32" thick, 12" x 12" | Silicone sheet for tube housing sample tube gasket |
| 3788T22 | 1 | McMaster-Carr | Silicone rubber sheet with smooth texture, 50A durometer, 1 mm thick, 12" x 12" | Silicone sheet for gaskets |
| 8589K62 | 1 | McMaster-Carr | Clear Scratch- and UV-Resistant Acrylic Sheet, 12" x 24" x 3/16" | Acrylic sheet stock for laser-cut acrylic door |
| RS-F2-DUCL-02 | 1 L | Formlabs | Durable Resin | 3D printing resin |
| M-KVD-VTPW | 0.75 kg | MatterHackers | Ultimaker PVA Filament - 2.85mm | 3D printing material |
| HTP32830-BLK | 3 kg spool | Protopasta | Black Opaque HTPLA Filament | 3D printing material |
| 7468A62 | 8 fl oz | McMaster-Carr | Pliobond 25 | Rubber cement for claws and tube sockets |
| 402026-WHT-12x2 | 2 | Cable Matters | Outlet Surge Protector Power Strip with USB, 12 ft long Extension Cord with Low Profile Plug | Surge protector for computer and automation hardware |
| HB-UM43 | 1 | Sabrent | 4 Port uSB 3.0 Hub with Individual LED Power Switches | USB hub for automation hardware |
| 90086 | 1 | Velcro | Velcro Sticky Back Tape, 5' x ¾" | Velcro strips for cable management / other device management |
| 70215K101 | 1 | McMaster-Carr | 170-Piece UV-Resistant Cable Tie Assortment | ZIp-ties for cable management |
| 6334K41 | 1 | McMaster-Carr | Heat Shrink Tubing Kit | 12-8 AWG (3/8" Expanded), 2 pieces for the aluminum claws |
| 458742928317 | 1 | Reynolds Wrap | Aluminum Foil 200 sq ft. | Foil for tube sockets reflective strips |
