## Supplementary material for "An open-source FACS automation system for high-throughput cell biology": Electronics Bill of Materials

### Electronics BOM

| Item | Quantity | Part number | Vendor | Name | Description |
|---|---|---|---|---|---|
| SMD291SNL | 1 | SMD291SNL-ND | Digikey | SOLDER PASTE NO-CLEAN LF 5CC SYR | High-temperature solder paste |
| SMDLTLFP10T5 | 1 | SMDLTLFP10T5-ND | Digikey | SOLDER PASTE LOW TEMP LF T5 10CC | Low-temperature solder paste |
| **5-0004 FACS Automation PCB** | 1 | **5-0004** | **OSH Park** | **FACS Automation PCB** | PCB for controlling motors, encoders, cooling, and thermistors. |
| 5-0004 FACS Automation PCB Stencil | 1 | -- | OSH Stencil | FACS Automation PCB Stencil | Solder paste stencil for FACS Automation PCB |
| C1, C2, C3, C4, C9, C10, C11, C12, C17, C18, C19, C | 18 | 1276-1869-1-ND | Digikey | CAP CER 10UF 25V X5R 0603 | General 10uF capacitors |
| C5, C6, C7, C8, C13, C14, C15, C16, C21, C22, C23, | 18 | 1276-1935-1-ND | Digikey | CAP CER 0.1UF 50V X7R 0603 | General 0.1uF capacitors |
| D1 | 1 | US1A-FDICT-ND | Digikey | DIODE GEN PURP 50V 1A SMA | Diode |
| J5, J6 | 2 | A1411-ND | Digikey | CONN PLUG HSG 2POS 5.08MM | Power + valve connector |
| #J5, #J6 | 2 | A14367-ND | Digikey | CONN HEADER VERT 2POS 5.08MM | Power + valve header |
| #J5, #J6 | 4 | A25555-ND | Digikey | CONN SOCKET 18-24AWG CRIMP TIN | Power + valve crimps |
| #J6 | 1 | 839-1291-ND | Digikey | CONN PWR JACK 2.1X5.5MM SOLDER | Power plug |
| J7, J8, J10, J11, J13, J14 | 6 | 732-5443-ND | Digikey | CONN RCPT 6POS IDC 28AWG GOLD | 2x03 headers |
| #J7, #J8, #J10, #J11, #J13, #J14 | 6 | 732-5394-ND | Digikey | CONN HEADER VERT 6POS 2.54MM | 2x03 connectors |
| J9, J12 | 2 | 732-5444-ND | Digikey | CONN RCPT 8POS IDC 28AWG GOLD | 2x04 headers |
| #J9, #J12 | 2 | 732-5395-ND | Digikey | CONN HEADER VERT 8POS 2.54MM | 2x04 connectors |
| J15 | 1 | S9171-ND | Digikey | CONN HEADER VERT 16POS 2.54MM | 2x08 headers |
| #J15 | 1 | S9288-ND | Digikey | CONN HEADER R/A 16POS 2.54MM | 2X08 connectors |
| P1 | 1 | SAM15923-ND | Digikey | CONN HEADER SMD 36POS 2.54MM | 2x18 header |
| P2, P3, P4, P6, P7 | 5 | WM14828-ND | Digikey | CONN HEADER SMD 8POS 2.54MM | 1x8 header |
| P5 | 1 | WM8617-ND | Digikey | CONN HEADER SMD 4POS 2.54MM | 1x4 header |
| Q1 | 1 | IRLML2803PBFCT-ND | Digikey | MOSFET N-CH 30V 1.2A SOT23 | Solenoid valve MOSFET |
| TP1, TP2, TP3 | 3 | 36-5006-ND | Digikey | PC TEST POINT COMPACT BLACK | Test points |
| R1 | 1 | RMCF0805FT330RCT-ND | Digikey | RES 330 OHM 1% 1/8W 0805 | 330 Ohm resistor |
| R2, R3, R4, R5, R6, R7, R8, R9, R10, R11, R12, R13, | 20 | RNCP0603FTD10K0CT-ND | Digikey | RES 10K OHM 1% 1/8W 0603 | 10k Ohm resistor |
| R21, R22, R23, R25, R33, R34, R35, R36, R45, R46, F | 18 | YAG1235CT-ND | Digikey | RES SMD 100K OHM 0.1% 1/10W 0603 | 100k Ohm resistor |
| R29, R30, R31, R32, R41, R42, R43, R44, R53, R54, R55, R56, R63, R64, R65, R72, R73, R74 | 18 | CSR0603FKR500CT-ND | Digikey | RES 0.5 OHM 1% 1/8W 0603 | 500mOhm resistor |
| U1, U2, U3, U4, U5, U6, U7, U8, U9, U10, U11, U12, U | 18 | MAX14870ETC+CT-ND | Digikey | IC MOTOR DRIVER 4.5V-36V 12TDFN | Motor drivers |
| DB25-1, DB25-2 | 2 | 2057-DB25-SF-M1-ND | Digikey | CONN D-SUB RCPT 25POS IDC | DSUB25 connectors |
| #DB25-1, #DB25-2 | 2 | 2057-DB25-PD-ND | Digikey | 2057-DB25-PD-ND | DSUB25 headers |
| DB15 | 1 | 215ME-ND | Digikey | CONN D-SUB PLUG 15POS PNL MNT | DSUB15 connectors |
| #DB15 | 1 | AFR15B-ND | Digikey | CONN D-SUB RCPT 15POS IDC | DSUB15 headers |
| -- | 3 | 3M157833-1-ND | Digikey | CBL RIBN 26COND 0.05 GRAY 1' | Shielded ribbon cables |
| #TH1, #TH2, #TH3, #TH4, #TH5, #TH6 | 6 | 615-1146-ND | Digikey | THERMISTOR NTC 10KOHM 3575K BEAD | Thermistors |
| -- | 1 | 839-1291-ND | Digikey | CONN PWR JACK 2.1X5.5MM SOLDER | Power barrel jack connector |
| **5-0007 FACS Photointerruptor Breakout Board** | 1 | **5-0007** | **OSH Park** | **FACS Photointerruptor Breakout Board** | PCB for socket encoders |
| 5-0007 FACS Photointerruptor Breakout Board | 1 | -- | OSH Stencil | FACS Photointerruptor Breakout Board Ster | Solder paste stencil for photointerruptor breakout board |
| U1, U2, U3, U4, U5, U6, U7, U8, U9 | 18 | 516-2467-1-ND | Digikey | SENSOR OPT REFLECTIVE 2MM 6SMD | Optical encoders |
