## Supplementary material for "An open-source FACS automation system for high-throughput cell biology": Example Summary Report of Sorted Samples

**Summary PDF reports of five- 96 well plate sorts**

A summary PDF report is produced after each 96 well plate sort. Below is the summary report from five- 96 well plates. This representative array of 480 samples sorted with the automated system shows the diversity of fluorescence profiles between different gene targets and the ability of the gating algorithm to tailor-make gates for each population. The summary report is generated by bespoke software that exports the sorting data from the sorter GUI. The report facilitates examination of the run performance.

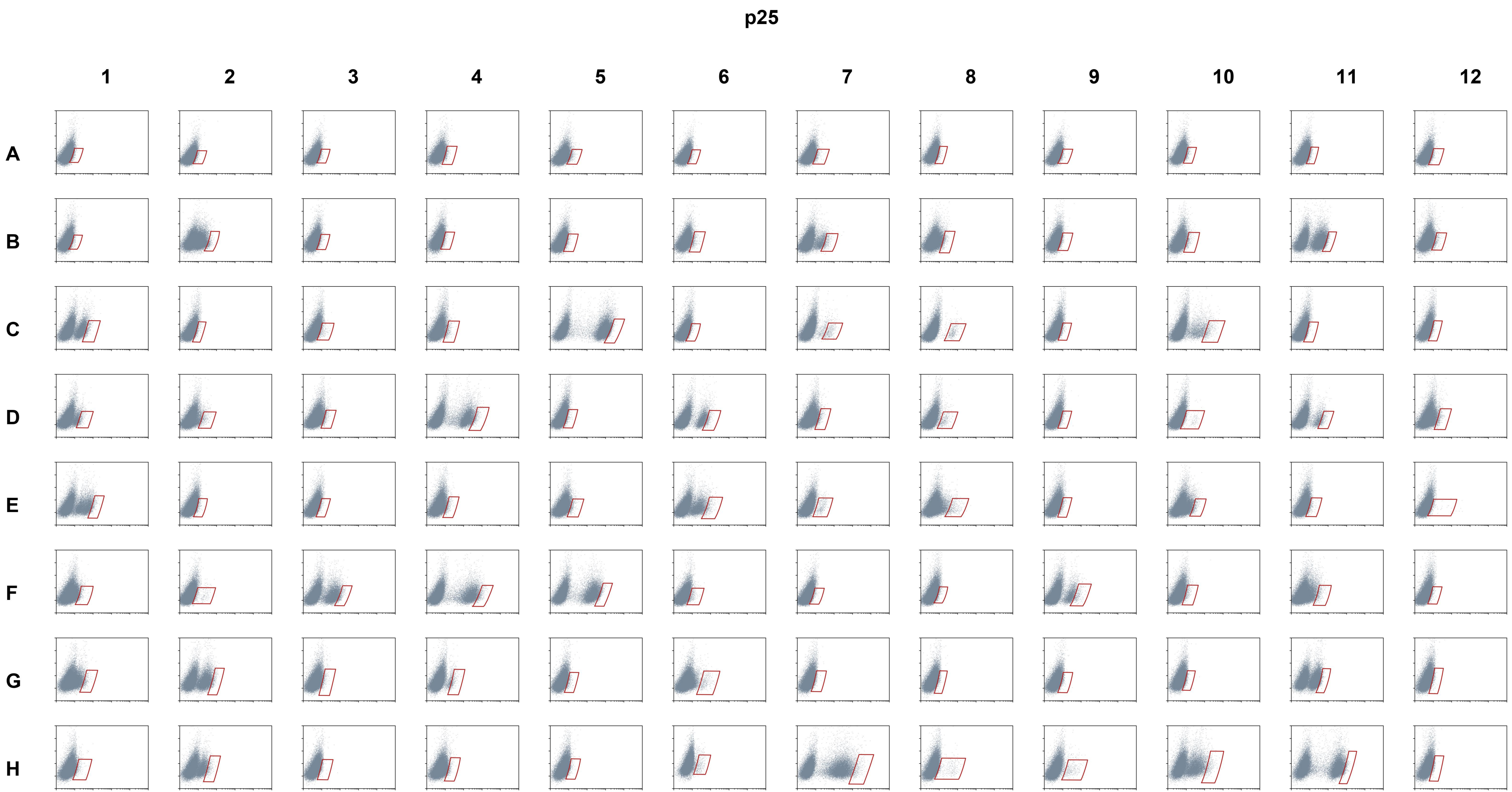
p25
1
2
3
4
5
6
7
8
9
10
11
12
A
B
C
D
E
F
G
H

# p27

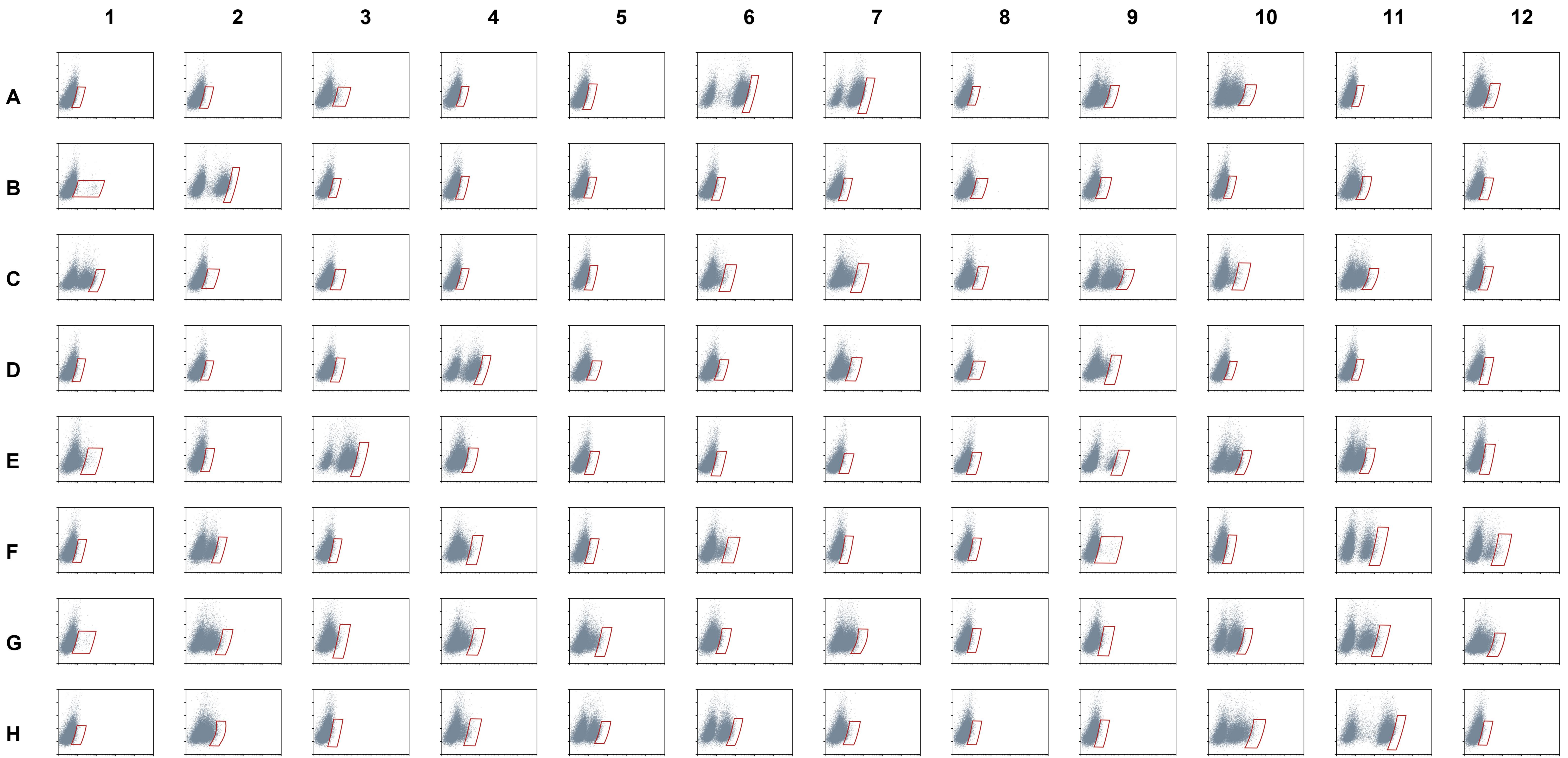

p28

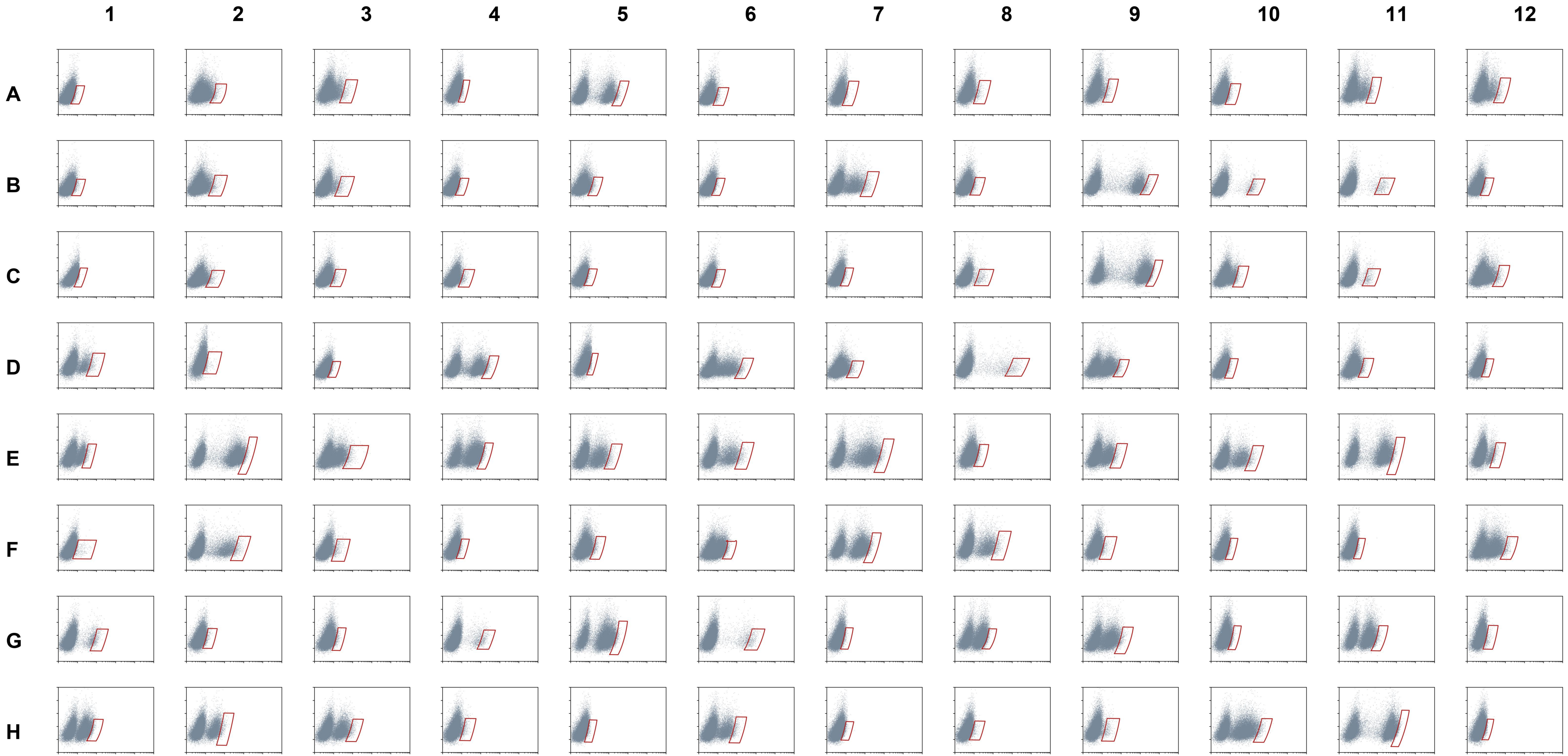

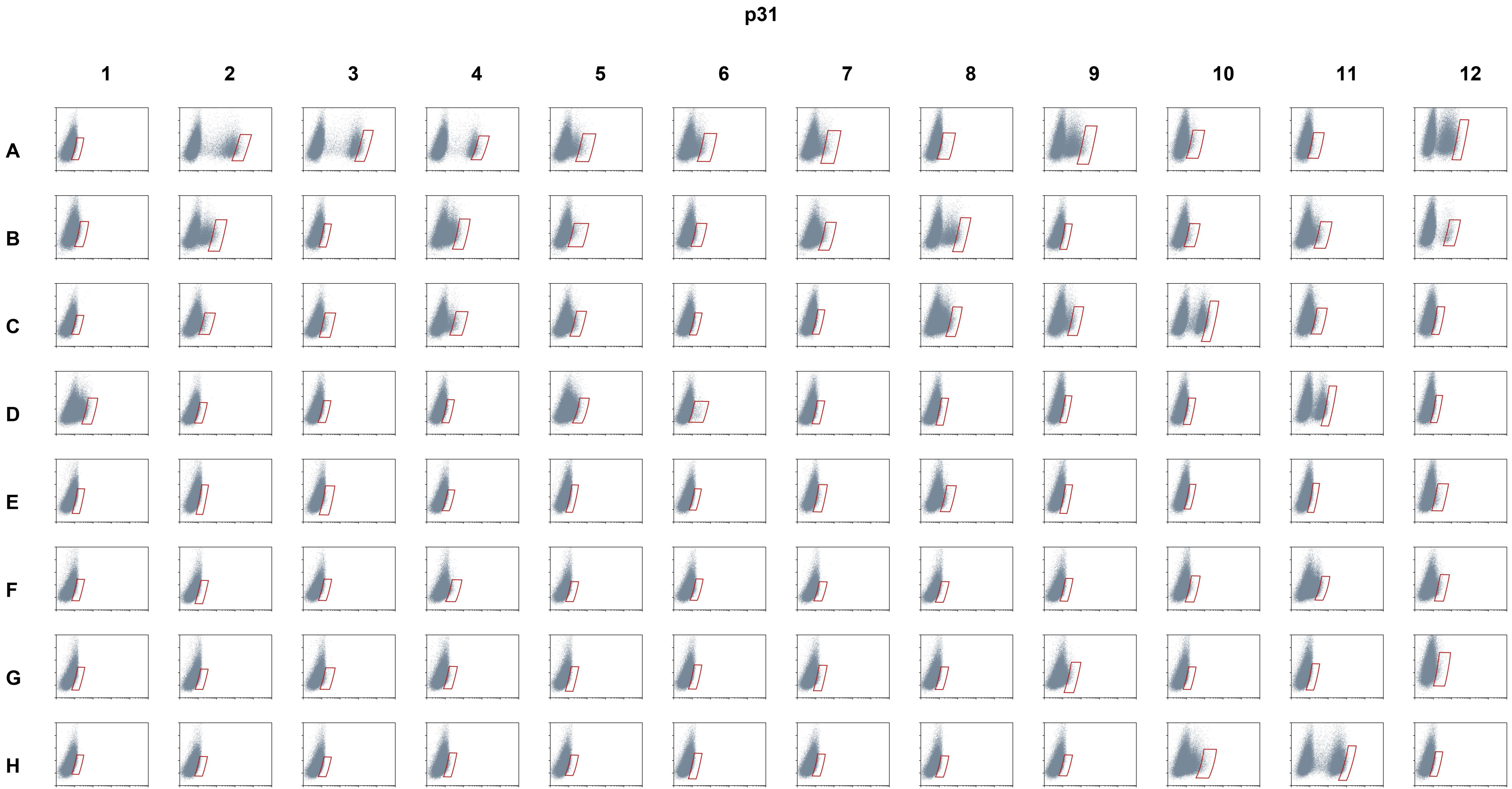
p31
1
2
3
4
5
6
7
8
9
10
11
12
A
B
C
D
E
F
G
H

p32

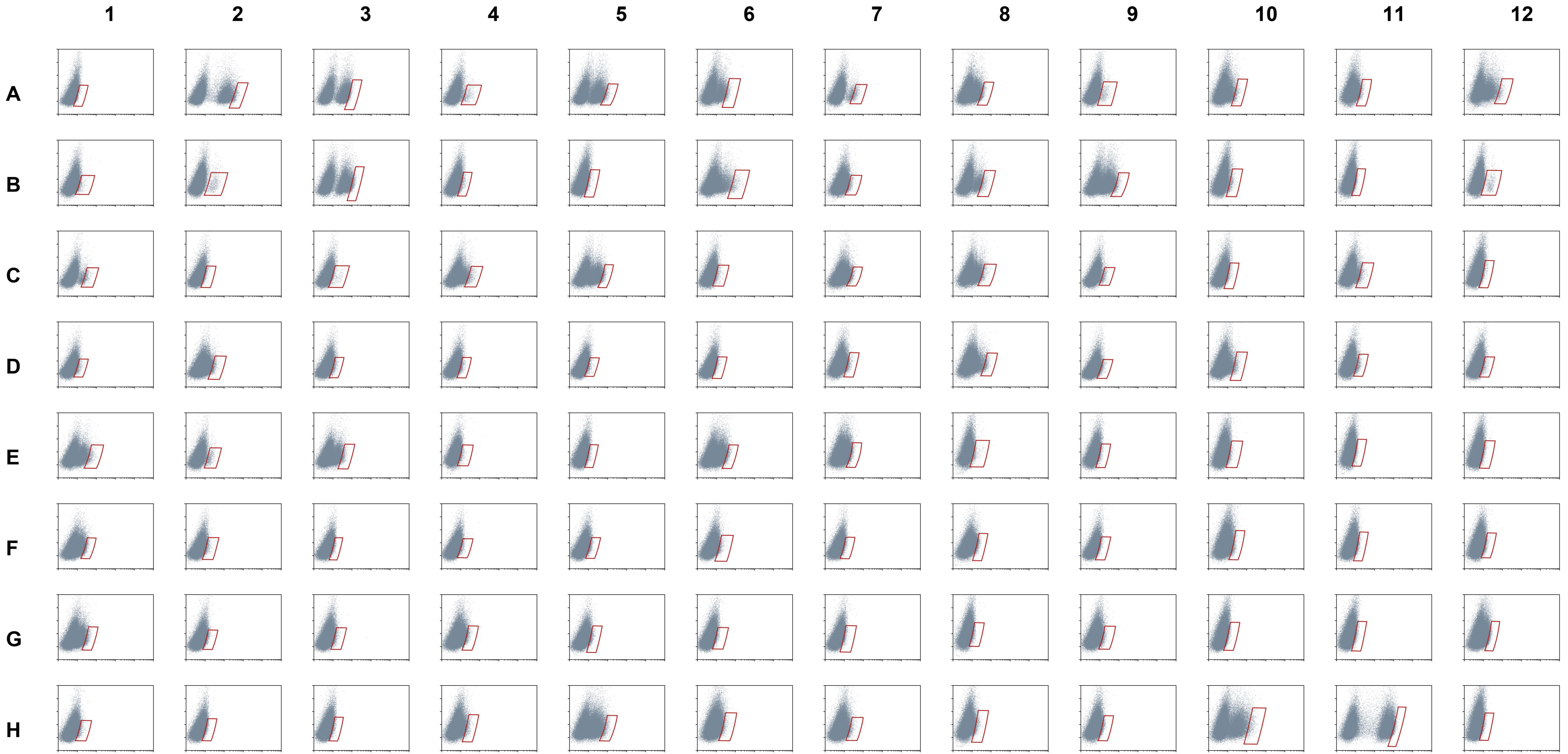
