## Supplementary material for "An open-source FACS automation system for high-throughput cell biology": Automation Standard Operation Procedure

### FACS Automation SOP

**Diane Wiener and Emily Huynh**
**Last updated: 2023-02-24**

#### Introduction

This SOP explains, in as much detail as possible, the procedure for using the FACS automation system for auto-loading samples into the Sony Cell Sorter SH800S and automatic gating for up to 18 samples at a time.

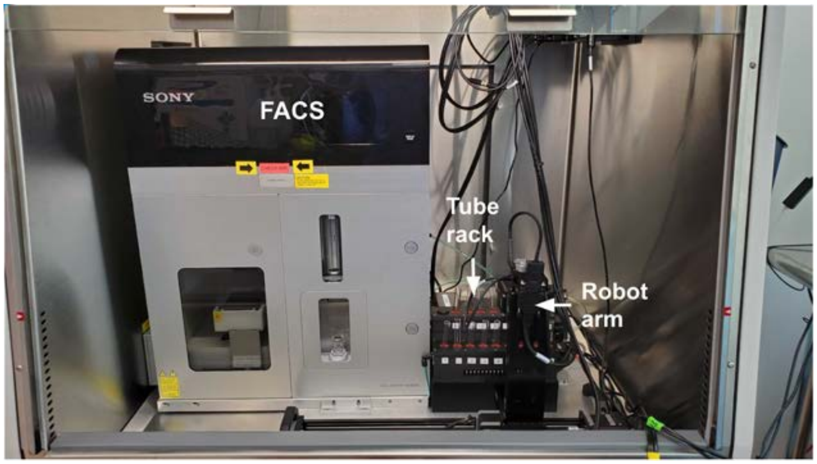


**Figure 1: The FACS automation system. Labeled are the Sony SH800 (FACS), sample tube housing (Tube rack), and Zaber stage (Robot arm).**

#### Part 1: Sample & system prep

**Step 1: Turn on the BSC fan.**
Turn on the BSC fan to keep the inside of the hood cool.

**Step 2: Restart the computer.**
Only keep open applications that are necessary (Sony Cell Sorter Software and GitBash).

**Step 3: Power on the FACS Automation system.**
Turn on the power supply affixed to the side of the Sony biosafety, Figure 2. This provides power to the Zaber stage (robot to move the samples) and the Arduino, which controls custom electronics for the automation.

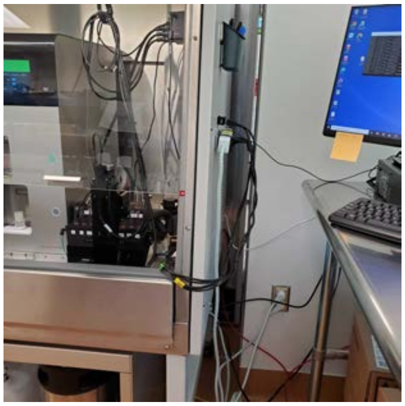

**Figure 2: The automation power supply is affixed to the side of the biosafety cabinet.**

**Step 4: Start cooling the tube housing.**
Go to the Desktop and click the program titled "`1 - CHILL HOUSE`."
The sample tube housing needs a minimum of 10 minutes to equilibrate. The sample tube housing (Figure 1, Tube rack) feels cool to the touch after reaching equilibrium.

Two audible cues indicate that the solenoid valve has opened:

1. a loud click of the valve opening
2. the sound of air flowing

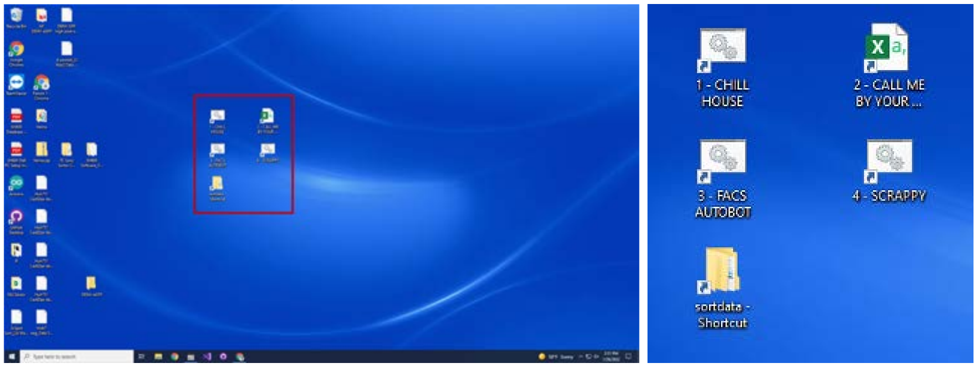


**Figure 3: Desktop image of the CHILL HOUSE.**

**Step 5: Prepare the FACS Automation Sample CSV script configurations.**
Go to the Desktop and click the program titled "`2 - CALL ME BY YOUR NAME`."

**Do not use underscores _ in your sample names! These will result in a fatal error. Each sample name within a Sony sorter experiment must be unique.**

Each row indicates a single sample to be sorted. It includes the following information:

1. Column 1: Tube Number
    a. This is the location of the sample tube in the tube housing, 1-18.
2. Column 2: Name

    a. This is the name of the sample that is used by the Sony software for both profiling and sorting.
    b. Do not use underscores “_” in sample names.
    c. The sample name must be unique within a Sony sorter experiment.
3. Column 3: Well
    a. This is the Well ID location for where the sample is to be sorted into in the destination plate.
    b. The naming convention is Row then Column.

| | A | B | C |
|---|---|---|---|
| 1 | Tube | Name | Well |
| 2 | 1 | facsML0010-p24-D1 | B1 |
| 3 | 2 | facsML0010-p24-D2 | B2 |
| 4 | 3 | facsML0010-p24-D3 | B3 |
| 5 | 4 | facsML0010-p24-D4 | B4 |
| 6 | 5 | facsML0010-p24-D5 | B5 |
| 7 | 6 | facsML0010-p24-D6 | B6 |
| 8 | 7 | facsML0010-p24-D7 | B7 |
| 9 | 8 | facsML0010-p24-D8 | B8 |
| 10 | 9 | facsML0010-p24-D9 | B9 |
| 11 | 10 | facsML0010-p24-D10 | B10 |
| 12 | 11 | facsML0010-p24-D11 | B11 |
| 13 | 12 | facsML0010-p24-D12 | B12 |

**Figure 4: Example CSV. Remember to SAVE IT.**

**Step 6: Prepare the Sony FACS Cell Sorter SH800S.**
Turn on the compressed air supply for the Sony Sorter. Open the sash of the biosafety cabinet. Turn on the fan. Power on the Sony FACS. See Figure 5.

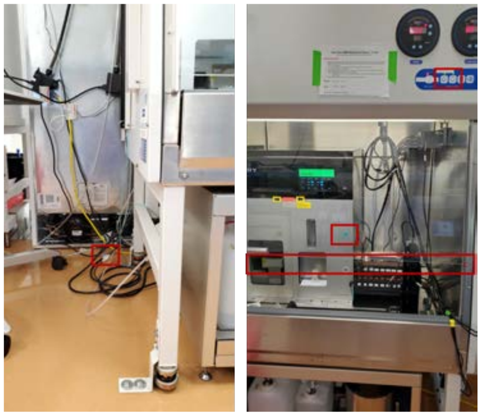

**Figure 5: The biosafety cabinet with the location of the compressed air supply (left), and the sash height, fan button, and power button (right).**

Open the Cell Sorter Software, Figure 6.

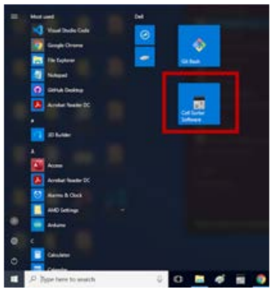

**Figure 6: The Cell Sorter Software icon and location on the TC computer.**

- Below the biosafety cabinet is the fluidics cart with three containers, Figure 7. Unlock the cart and pull it open to access it.

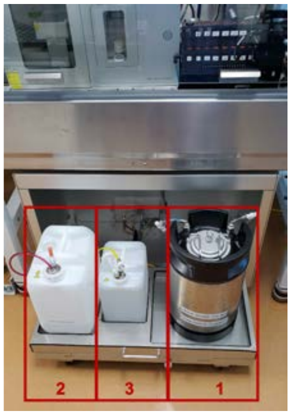


**Figure 7: The Fluidics Cart. Box 1 is for the sheath fluid, box 2 for the non-hazardous waste, and box 3 for the ethanol.**

1. Close the sheath vent at the far right. If the sheath fluid is low, fill it with the sheath fluid, Thermo NERL Diluent 2.
2. Empty the non-hazardous waste at the far left and fill the bottom with about 0.5-1 inch bleach.
3. Fill the ethanol tank with ethanol, if it is low.

- There is a rear left side panel to access the left side of the Sony Sorter. Disconnect the DI water tank, and fill it with MilliQ water, Figure 8.

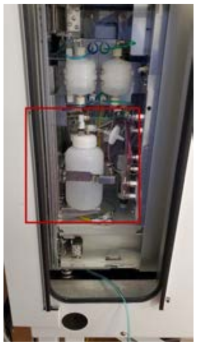

**Figure 8: The rear left side panel with the DI-water tank boxed in red.**

- Follow the Cell Sorter Software prompts to insert a new microfluidic sorting chip and complete droplet calibration. Prepare 12 mL of fresh DI-water and bleach in 15 mL conical tubes.

**These steps are important to do prior to a run, as fluidic errors can potentially cause errors in the Sony Software (e.g., requiring droplet re-calibration).**

**Step 7: Prepare the cells for sorting, and a 96 well plate for the sorted cells.**
Follow standard procedures for preparing cells in FACS buffer for sorting. Please use Falcon test tubes (blue cap test tubes) for your samples.

**Note that the expected sample volume is 300 µL.**

- The OpenCell sort protocol is found here:
    - https://www.protocols.io/edit/opencell-sort-days-bqh2mt8e

**Step 8: Replace Sony FACS tube holder with the 5 mL sample holder with automation funnel.**
Replace the tube holder in the Sony FACS with the holder that contains the tube funnel (Figure 9). This is necessary for proper placement of the sample tubes by the automation robot.

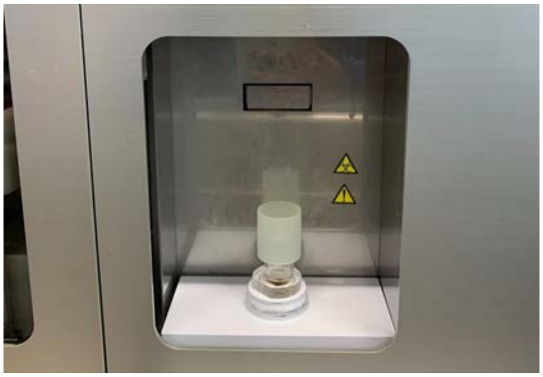

**Figure 9: The modified 5 mL sample tube holder with automation funnel.**

**Step 9: Place samples into tube housing.**
Vortex the sample tubes before placing them into the tube housing. The samples must be placed into the tube housing according to the locations in the CSV file created in Part 1, Step 4. Ensure that the sample tubes are uncapped.

If running a control sample, insert it into the sample tube holder to set up the gates.

**Step 10: Place the 96 well plate with media into the collection door.**
Place the 96 well plate into place in the collection door. Well A1 is positioned in the left corner closest to the door.

#### Part 2: FACS Sorting

**Step 1: Complete the pre-automation checklist.**
If not already done in Part 1, Step 9, vortex the first tube and place it into the sample holder in the FACS. Typically, this sample is the negative control by which to set all of the gates.

Start a new experiment in the Cell Sorter Software:
- **Do not use underscores in your experiment name as this will result in a fatal error of the automation program!**
- **Uniquely name the experiment with a timestamp, e.g. yyyymmdd Experiment Name. This is for the final gate to be properly adjusted for each sample in the automated workflow.**

Before starting automation, complete your pre-automation checklist:
**Checklist:**
- [ ] Tanks are full: DI water, ethanol, sheath fluid.
- [ ] The BSC fan is on and the hood is slightly open.
- [ ] Non-hazardous waste is emptied and filled with about 0.5-1 inch of bleach.
- [ ] The experiment name does not contain underscores and uses a date stamp to make it unique.
- [ ] All sample names are unique.
- [ ] Any previous tabs on the Cell Sorter Software are closed.
- [ ] The Cell Sorter Software window is maximized.
- [ ] There is a plate filled with media for the sorted cells.
- [ ] Your first sample is placed in the Sony sample holder. If you plan to sort the first sample you will need to leave the first tube location empty to place it back in the housing after setting your gates up.

**Step 2: Start the automation program.**
Start the automation program by going to the Desktop and clicking on the "`3 - FACS_AUTOMATION`" icon, which will open a new GitBash terminal. After the terminal opens, please lay it out such that the Cell Sorter Software is fully open in the background and the Terminal is in the bottom right, as in Figure 10.

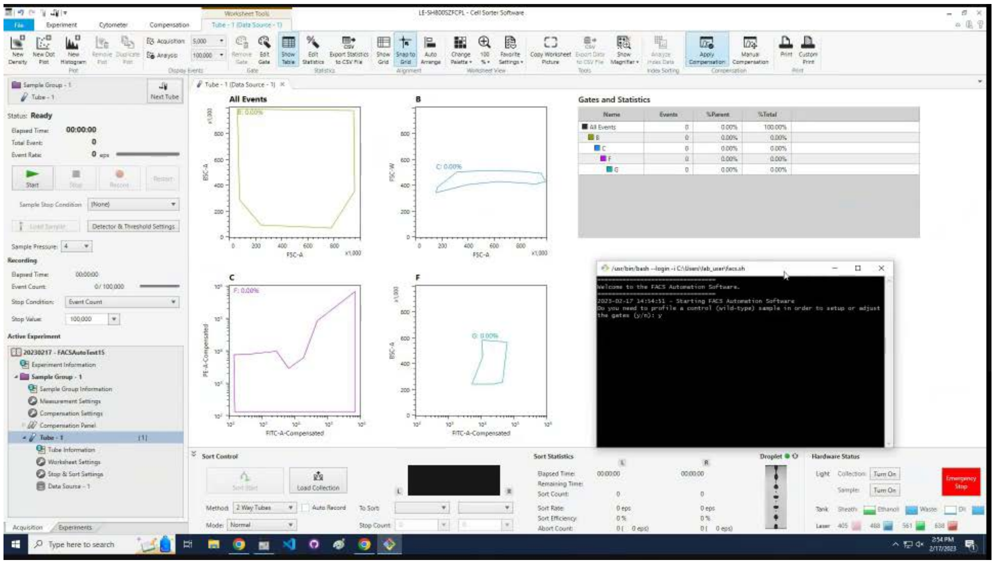


**Figure 10: FACS Automation icon on the desktop, and the terminal in the proper position.**

**Once you open this program, please do not touch the computer unless specifically prompted by the program!**

Opening the terminal will prompt you for several things, which is also summarized in the flow chart at the end of this SOP:

1. It will ask you if you need to set up the first gates and profile a sample.
2. It will ask you for your name so that you will receive Slack messages regarding the sort.
3. Next, it will attempt to find the experiment name by itself. Don't be startled and move the mouse! It will return to the terminal once found to ask for you to confirm if it is the correct experiment name, in which you will type 'y' or 'n'.
4. The automation software prompts for the first tube number (#) to start the automation. This is the number of the first tube to start sorting on the tube housing, e.g. the number in the `Tube` column of `sort_samples.csv` to start. For example, in Fig. 11, if you want to start with `facsML0010-p24-D5` in Row 6, you would type in `5` for the `Tube #`. If you wanted to start at the first sample (`facsML0010-p24-D1`), you would type in `1`.

| | A | B | C |
|---|---|---|---|
| 1 | Tube | Name | Well |
| 2 | 1 | facsML0010-p24-D1 | B1 |
| 3 | 2 | facsML0010-p24-D2 | B2 |
| 4 | 3 | facsML0010-p24-D3 | B3 |
| 5 | 4 | facsML0010-p24-D4 | B4 |
| 6 | 5 | facsML0010-p24-D5 | B5 |
| 7 | 6 | facsML0010-p24-D6 | B6 |
| 8 | 7 | facsML0010-p24-D7 | B7 |
| 9 | 8 | facsML0010-p24-D8 | B8 |
| 10 | 9 | facsML0010-p24-D9 | B9 |
| 11 | 10 | facsML0010-p24-D10 | B10 |
| 12 | 11 | facsML0010-p24-D11 | B11 |
| 13 | 12 | facsML0010-p24-D12 | B12 |

**Figure 11. Example CSV in spreadsheet view.**

**Step 3: Set up your gates after the automated profiling.**
After the automated profiling is complete, the terminal will prompt you to set up your gates.

Use the Polygon tool to draw all of the gates. Double click inside of the gate to create the next gate.

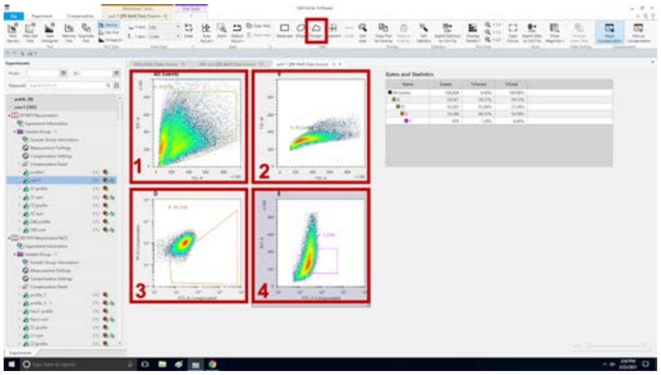


**Figure 12: The four gates that need to be drawn for sorting.**

1. First gate, box 1 in Figure 10, and below:
   - x-axis: FSC-A
   - y-axis: BSC-A

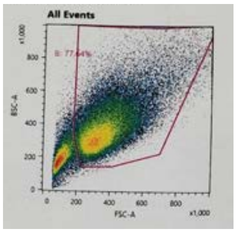


2. Second gate, box 2 in Figure 10, and below:
   - x-axis: FSC-A
   - y-axis: FSC-W

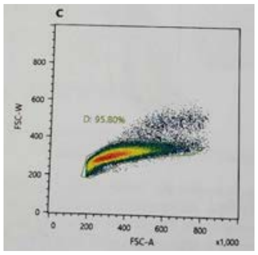


3. Third gate, box 3 in Figure 10, and below:
   - x-axis: FITC-A Compensated
   - y-axis: PE-A Compensated

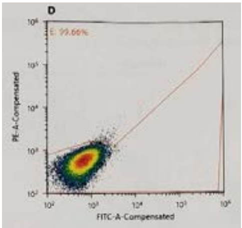


4. Fourth gate, box 4 in Figure 10, and the plot below:
    - X-axis: FITC-A Compensated
    - y-axis: BSC-A

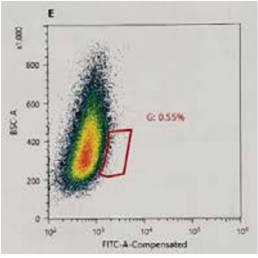


**Step 4: Start the full automation process.**
The automation program will begin once you press enter to confirm that your gates are fully set up. Before doing so, you will want to either discard or place the first sample you used to profile back in the tube housing. **Ensure the Sony tube holder is empty before pressing enter**!

When you are ready, press enter and the first sample you want to sort will begin vortexing.

After 30 seconds of vortexing the sample tube, the robot stage will shuttle the sample tube to the Sony sorter. The automation software will then control all keyboard and mouse inputs. You will not be prompted until the program completes or if errors occur.

**Periodically check on the profiling and sorting of the samples.** You will be messaged on the Slack channel `#autofacs4022` or `#autofacs4028` with any errors.

If any errors or problems occur during the automated sorting, use the following hotkeys if needed:

| Key | Function |
|---|---|
| F5 | Pause |
| F6 | Resume |
| F7 | Stop |

For emergency stop, shut off the power strip closest to you (Figure 2).

#### Part 3: Cleanup

**Step 1: Label and place the 96 well plate in the appropriate incubator.**

**Step 2: Close the solenoid valve to stop the air supply.**
When the automated sorting is complete, the robot stage moves to its home position (all the way to the right of the biosafety cabinet). The automation software will prompt whether to close the solenoid valve. At the end of all of the samples to be sorted for the day, close the solenoid valve by entering ‘y’ for yes. The automation software disconnects from the hardware devices and closes.

If there are multiple runs of the automation software in a day, then enter ‘n’ for closing the solenoid valve. This ensures that the sample tube housing will remain at the proper temperature. Restarting the program will reset the Arduino, which will shut off the solenoid valve. If you are continuing to use the automation platform, type in ‘y’ to turn on the valve.

**Step 3: Power off the hardware devices.**
Turn off the power strip that powers the Zaber stage and the Arduino-controlled electronics (see Part 1, Step 1 image).

**Step 4: Follow the cleanup procedure in the SOP for the Sony Sorter.**

- Add bleach to all of the sample tubes. After 30 minutes, discard as non-hazardous liquid waste.
- Perform a bleach cleaning and DI water rinse for the Sony Sorter.
    - In the Sony Softer, under the Cytometer tab, perform a bleach clean and DI rinse for the probe before hardware and software shutdown.
- Perform an ethanol cleaning weekly.
    - Follow the Sony software prompts for ethanol cleaning.

#### Part 4: Data scraping
This section describes how to scrape your data after you are done sorting.

***The Data Scraping runs the smoothest when there is still an Active Experiment. It is best to run when running the bleach and water cleaning at the end of all of the sorting for the day.***

**Step 1: Set up your experiment to be in view for the scraper to find.**
A. If you are scraping an active experiment:
Scroll the experiment bar all the way to the top, where the experiment icon is, as seen in Fig. 13.

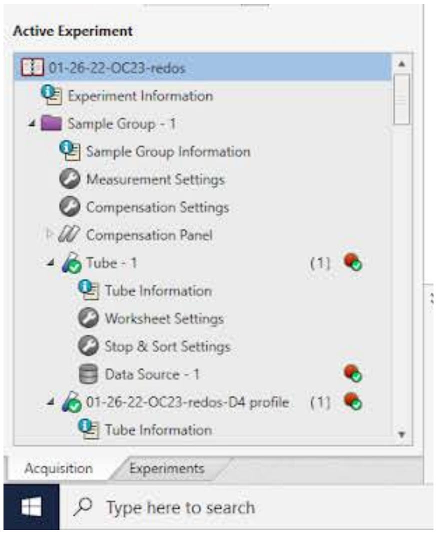


**Fig. 13**. Experiment bar scrolled all the way to the top.

B. If you are scraping a previous (inactive / analysis mode) experiment:
Position the experiment icon such that:

(1) it is the first one at the top,

(2) the icon is in full view, (See Fig. 14 for a correct example, and Fig. 15 for an incorrect example.)

(3) all other experiments are minimized by clicking the side arrow.

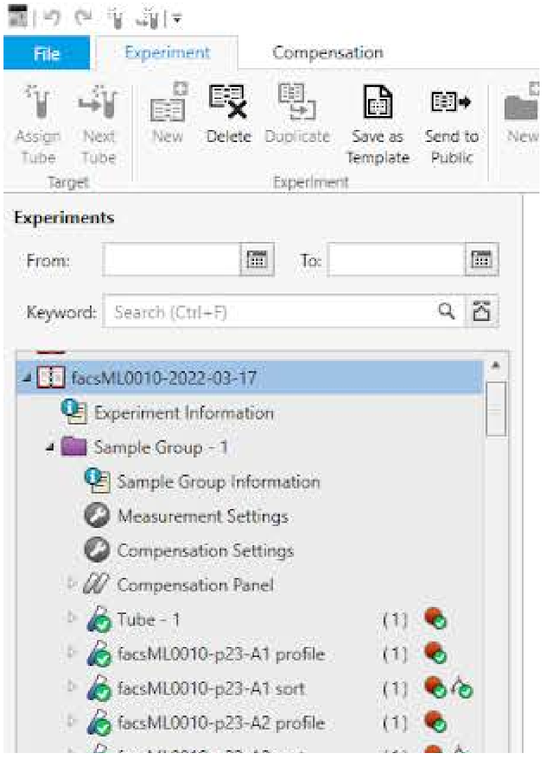

Fig 14. Correct example: The icon for the intended experiment is in full view at the top.

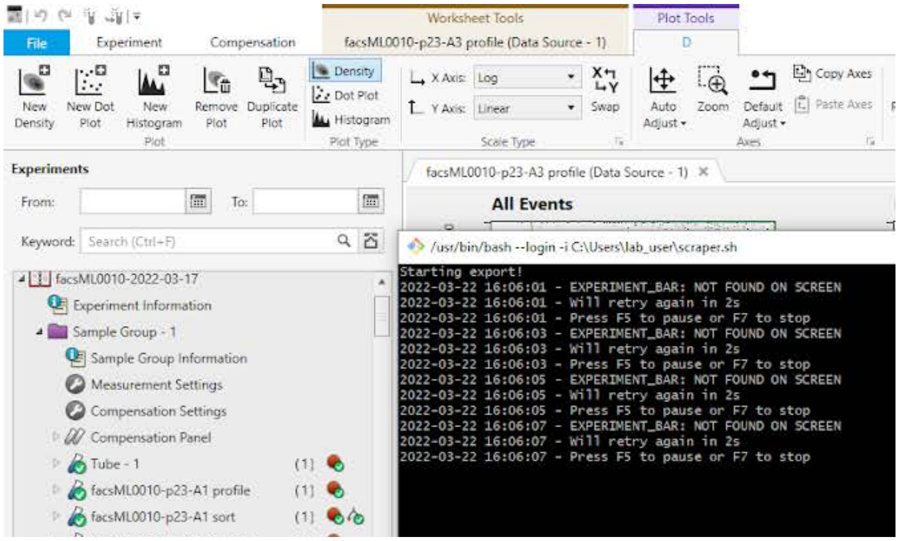


**Fig 15.** Bad example: The icon for the intended experiment is in occluded view. The terminal shows what error message shows up.

Once you properly position the experiment icon, open the experiment so all the tubes that were sorted can be seen, as seen in the figures above.

**Step 2: Start the scraper.**
Go back to the Desktop and click on '`SCRAPPY`' (see Fig. 1 for reference). It will ask if you need to export the samples, or if you want to skip to plotting. Return to the FACS software, and leave the mouse alone. If the scraper is successful it will click on the experiment icon, and continue scraping your data. If it is unsuccessful it will be idle, you will likely need to reposition the experiment icon and restart the scraper.

Leave the scraper alone until it has reached the bottom of the experiment bar. For inactive experiments, this will take awhile, as it needs to scroll through all the past experiments to reach the bottom of the experiment bar.

**Step 3: Answer scraper prompts once it is completed.**
Once the scraper is complete, it will return to the terminal and prompt you for generating plots and experiment summaries. See Fig. 16 below for the prompts.

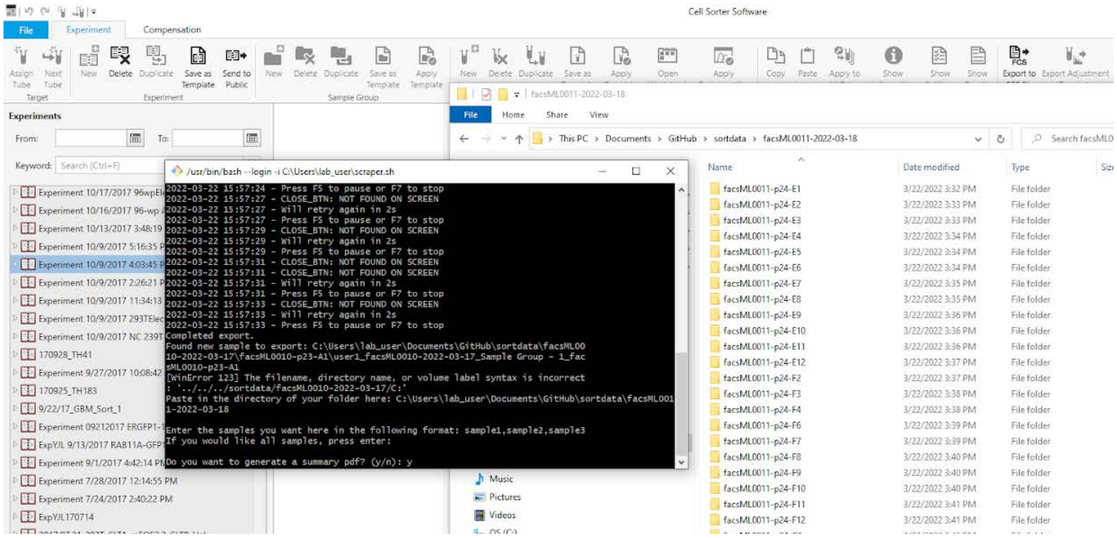


**Fig 16.** Scraper prompts once it is complete.

The first prompt is: '`Paste in the directory of your folder here:`'

Return to the Desktop and click on '`sortdata - Shortcut`' (see Fig. 1).

Click your folder experiment, as seen in Fig. 16.

Copy the file path at the top, and then return to the terminal and `Right Click + Paste + Enter`. **(CTRL+C CTRL+V will not work!!! It is assigned to a different key shortcut on GitBash.)**

The next prompt is: '`Enter the samples you want here in the following format: sample1,sample2,sample3. If you would like all samples, press enter:`'

Answer the final prompt: '`Do you want to generate a summary pdf: (y/n):`'

When it is complete, it will exit out.

#### Part 5: Safety

**Safety when the automation hardware is operating**

The robot arm moves with a high speed and with automated motions that are executed until the next process in the automation software, or if the power supply shown in Figure 2 is turned off. As such, if anything interferes with the intended motions, crashes or potential bodily harm can occur.

1. **Do not place any items on the automation platform as these will interfere with the motion of the robot arm.**

2. **Do not touch or insert your arms / hands inside the biosafety cabinet when the automation software is running. This can cause injuries where the robot pins you between it and another object.**

3. **Do not change the hardware configuration file.**

If there are problems with the automation workflow:
1. switch off the power to the robot arm at the power supply shown in Figure 2
2. press the ‘F5’ or ‘F7’ to pause or end the automation
3. contact Emily or Diane.

**Crashes**

It is very unlikely that the robot arm will crash with FACS automation system shown in Figure 1. The system has been extensively tested, and the locations of the objects shown in Figure 1 are specified in the automation software.

However, crashes can happen, especially if there are foreign objects impeding the path of the robot arm. (As highlighted in the safety when operating section, do not place any items on the automation platform.) The robot arm does have slip / stall detection, which will stop the robot from moving if it has stalled. This can happen if the stage crashes into an object.

When encountering a crash of the robot arm:
1. switch off the power to the robot arm at the power supply shown in Figure 2
2. press the ‘F5’ or ‘F7’ to pause or end the automation, if not automatic
3. contact Emily or Diane.

**Spills**

Spills can occur if a sample tube is not grabbed by the robot arm or inserted into the sample tube housing properly. Small spills of cells should be cleaned immediately per standard safety recommendations. If a sample tube breaks, remove it with tongs, and discard it into a biohazard sharps container. Cover the spill with absorbent material. Pour 1:10 bleach solution onto affected areas, let sit for at least 30 minutes. Then, use paper towels to absorb the spill, and wipe down with clean paper towels soaked with bleach. Discard all waste into a biohazard bag. Keep the biosafety cabinet fan on for at least 15 minutes after cleanup.

If spills occur inside of the sample tube housing, contact Emily or Diane for assistance in appropriate disassembly, cleaning following the above steps, and reassembly of the sample tube housing. Briefly, remove all sample tubes from the sample tube housing. A Phillips screwdriver is located inside of the biosafety cabinet. Use it to remove all of the screws from the top plate of the sample tube housing. Keep the screws in a safe place for reassembly. Remove the top plate, gasket, and test tube guide. Clean as detailed above. Inspect the tube sockets and motor plate inside of the tube housing for any liquid. If there is a spill on the motor plate or tube sockets that requires any disassembly, contact Emily or Diane for assistance. Contact Emily or Diane for reassembly and recalibration of the robot arm positions.

**Automation platform removal**

Automation platform removal should only be performed under the supervision of Emily or Diane. A table or lab cart must be available to hold the automation platform. Turn off the air supply for the sample tube housing. Power on the automation system and click the Chill House executable to open the solenoid valve. This is to purge the remaining pressurized air in the system. Power off the automation system. Disconnect the ¼" blue tubing at the quick disconnect at the rear of the Sony SH800 Sorter. Remove the two #6-32 connected to the bar attached to the Sony Sorter. See Figure 17. For temporary removal, grab the automation platform by the handles and place it on the cart. For long-term removal, disconnect the power cables, and the robot arm controller cables. Grab the automation platform by the handles, place it onto the table for storage.

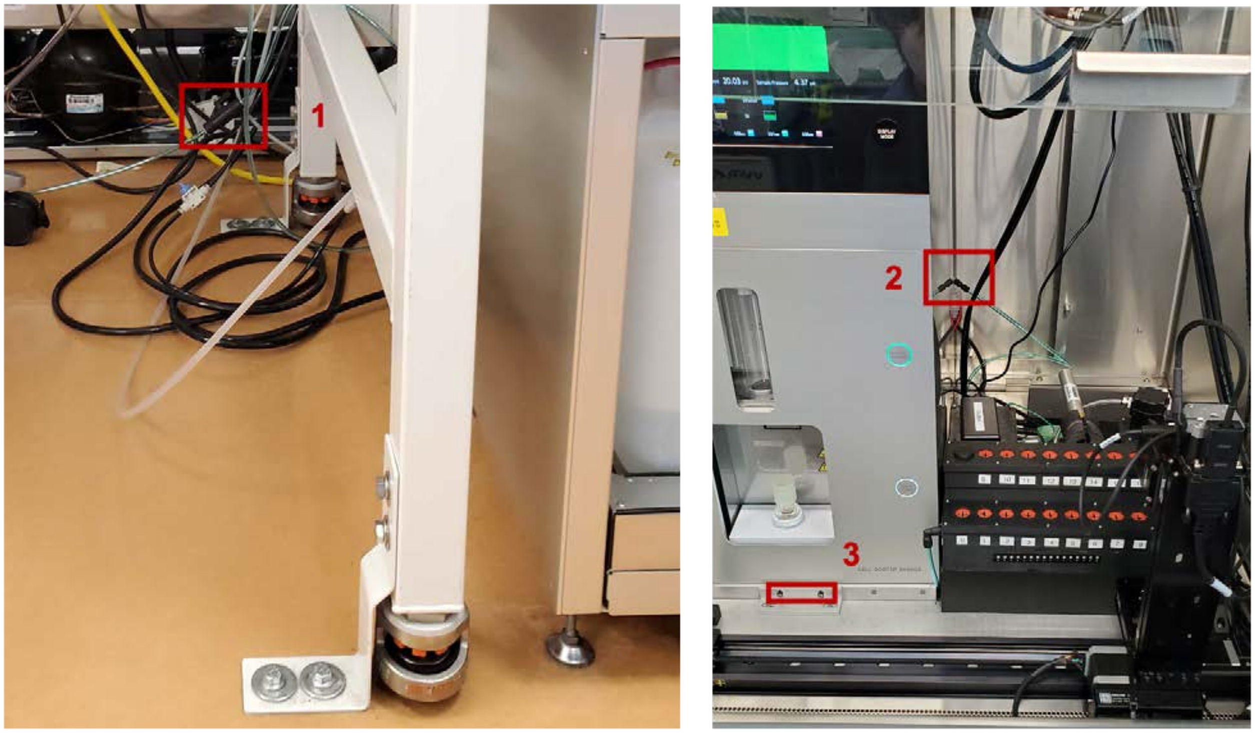


**Figure 17: Locations for the steps necessary for removal of the automation platform.**

#### Part 6: Potential Errors & Fixes

If an error is not listed below, please contact Diane/Emily.

**FACS automation program not accepting username that was before**

If the automation program was not quit properly, sometimes there are background instances of the application running that cause restart of the application to fail. This usually is observed by the `username` not being accepted.

1. Exit out of any open terminal windows.
2. Press `CTRL + ALT + DEL` and select the `Task Manager` option.
3. Navigate through the list searching for active `bash.exe` and `python` instances.
    a. Right click and kill all of them.

4. Restart the automation program, which should now function normally.

**<u>Catching or Restarting the automation program mid-run</u>**

For any error that requires restarting mid-run (Cell Sorter Software errors listed below, automation errors, or any unlisted errors), follow the steps below.

5. Press ‘F5’ to pause the system.
6. Navigate to the log file, which is located in “`C:\Users\lab_user\Documents\GitHub\logs`”, to determine where the automation software failed -- most likely, it failed finding the correct image. The error message when the software cannot find an image is: `[ICON NAME]NOT FOUND ON SCREEN: Will retry again in 2s`. If you cannot find this image, look at the last line on the log file to find out the last action taken by the automation software. Please contact Diane or Emily for any assistance.
7. Backtrack and complete all the steps up to the failure point, which might include duplicating and activating a tube, exporting CSV data, etc. Backtracking may not be possible if a gate was supposed to be set, and it was not.
8. If backtracking is not possible, press ‘F7’ to stop and exit the automation software. Unplug and replug USB cables to make sure that the hardware is no longer connected to the failed run. Decide whether to complete the remainder of the workflow for the current tube to finish the sort manually, or to restart with profiling of that sample. In either case, duplicate and activate the last tube, and close any open worksheet tabs, other than the newly activated tube. Restart the FACS automation script as before, following prompts for the experiment name. Start the process from the tube number you want the automation process to start from, (the number on the tube housing, and in the tube number column). Continue as normal.

**Real Example: “System is frozen.”**

1. Open the log from the FACS Automation system “`C:\Users\lab_user\Documents\GitHub\logs`” and scroll to the bottom of the file to find the current status / problem:

a. 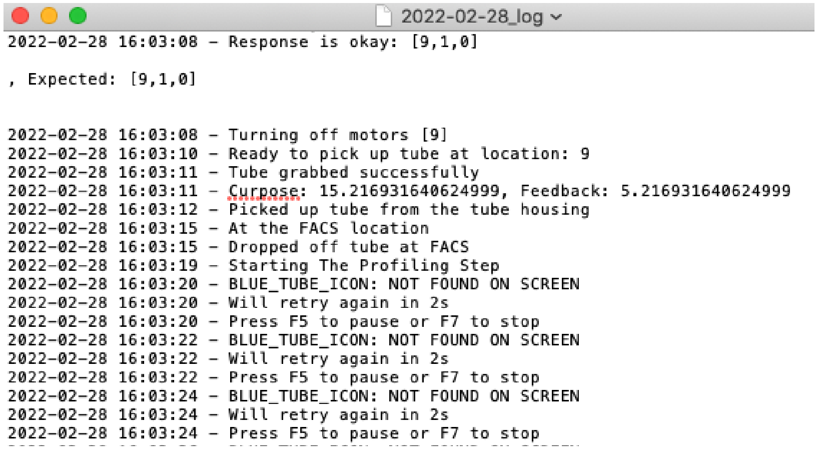
2022-02-28_log

```
2022-02-28 16:03:08 - Response is okay: [9,1,0]

, Expected: [9,1,0]


2022-02-28 16:03:08 - Turning off motors [9]
2022-02-28 16:03:10 - Ready to pick up tube at location: 9
2022-02-28 16:03:11 - Tube grabbed successfully
2022-02-28 16:03:11 - Curpose: 15.216931640624999, Feedback: 5.216931640624999
2022-02-28 16:03:12 - Picked up tube from the tube housing
2022-02-28 16:03:15 - At the FACS location
2022-02-28 16:03:15 - Dropped off tube at FACS
2022-02-28 16:03:19 - Starting The Profiling Step
2022-02-28 16:03:20 - BLUE_TUBE_ICON: NOT FOUND ON SCREEN
2022-02-28 16:03:20 - Will retry again in 2s
2022-02-28 16:03:20 - Press F5 to pause or F7 to stop
2022-02-28 16:03:22 - BLUE_TUBE_ICON: NOT FOUND ON SCREEN
2022-02-28 16:03:22 - Will retry again in 2s
2022-02-28 16:03:22 - Press F5 to pause or F7 to stop
2022-02-28 16:03:24 - BLUE_TUBE_ICON: NOT FOUND ON SCREEN
2022-02-28 16:03:24 - Will retry again in 2s
2022-02-28 16:03:24 - Press F5 to pause or F7 to stop
```

**Figure 18: A log file from the automation software with a BLUE_TUBE_ICON problem.**

b. In this case, a sample finished sorting and a new sample is starting.

        i. The last successful step was `[2022-02-28 16:03:19 Starting The Profiling Step]`
    c. The Blue Tube Icon is not found.
        i. The new sample tube was not duplicated and assigned. (The Sony GUI may have refreshed too slowly and the automated mouse click was not registered.)
    d. Click on the previous sample.
        i. Duplicate the previous sample.
        ii. Close the previous sample worksheet.
        iii. Assign the new sample.
        iv. Ensure that the new sample tube is visible in the sidebar.
    e. Press F6 to resume the automation.
    f. Confirm that the system resumes as expected.
        i. The new, active tube should be renamed at this point.

**<u>Icon Not Found Lookup Table</u>**

The automation log ("`C:\Users\lab_user\Documents\GitHub\logs`") details the progress and any potential errors encountered with the system. When the automation pauses it is often because an icon in the Sony GUI was not found. The error message when the software cannot find an image is: `[ICON NAME]NOT FOUND ON SCREEN: Will retry again in 2s.`

The [Lookup Table] here relates the icon name in the log to the icon to find and what step in the [sequence] the automation and sorting is in.

**<u>Scraper Software Errors</u>**

1. When the software is in analysis mode, it might not find the experiment icon. If it is unsuccessful it will be idle.
    a. Reposition the experiment icon to match Figure 14.
    b. Exit from GitBash.
    c. Restart, by clicking on '`SCRAPPY`'.
2. An error occurs during / after the Scraper prompts:
    a. Check that the file folder contains all of the samples and their data contents.
        i. If the data does not exist, exit out of GitBash and restart by clicking on '`SCRAPPY`'.
    b. Check that there are no additional samples from other experiments.
        i. Make sure only one experiment is open when scraping.
        ii. Delete any unwanted experiment folders that do not contain the proper datasets.
        iii. Follow the prompts on the screen to restart plotting the results.

**<u>Cell Sorter Software Errors</u>**

Sorter software errors will need to be addressed by the user, and the automation software will need a “push” to continue wherever it left off. Instructions to address common errors for the Sony FACS are below. Please follow instructions under “**Catching or Restarting the automation program mid-run”** after addressing Cell Sorter Software errors.

If not already paused, pause the automation system by pressing ‘F5’.

*Tank fluid error*

If tank fluid errors occur in the middle of the automated sorting, address the error by placing the sorter in Standby mode [Cytometer → Settings → Advanced Settings → Standby]. Empty or refill the tank as necessary. Return the sorter into Ready mode.

*Droplet calibration error*

Droplet calibration errors require recalibration of the microfluidic sorting chip. Pause the program (press ‘F5’) and follow the Sony software prompts to recalibrate the droplet.

*Exporting has failed error*

This error (shown in Figure 19) occurs because a new experiment was not created properly. Stop the automation software (Press ‘F7’). Start a new experiment by clicking [File → New], giving the experiment a unique name. Restart the automation. It is possible to restart the automation from the failed sample.

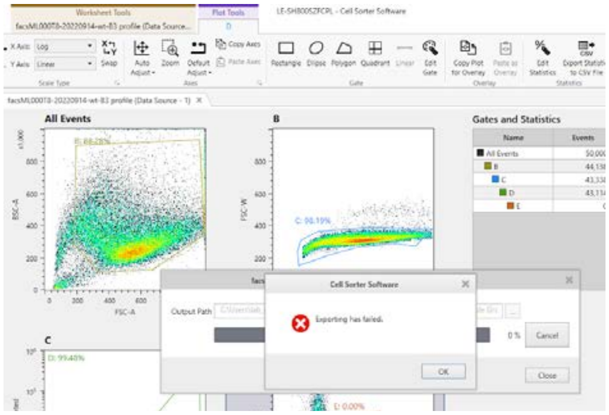


**Figure 19: Exporting failed popup due to a new experiment not created properly.**

*No recorded tube error*

If you were running an experiment in the Sony software and ran into the popup in Figure 20, then the automation software will fail. In order to recover from the error:

a. Press F7 to stop the automation.
b. Finish the sample or stop the Sony sorter sample.

c. Delete any incomplete (blue) tubes.
d. Start a new experiment with a unique name. Ensure the popup does not show up again.
e. It is possible to restart the automation program after the last sample that was fully sorted.

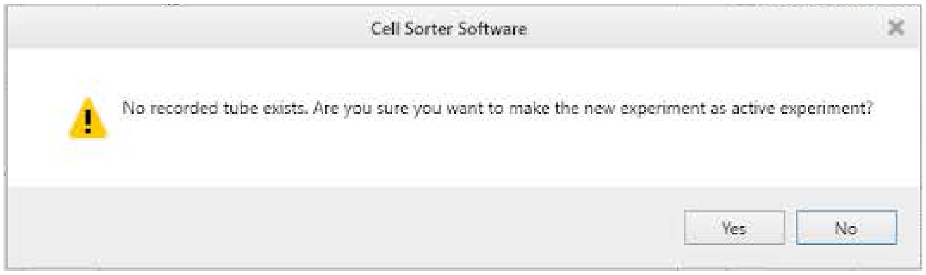


**Figure 20: New experiment popup that will cause the automation to fail.**

#### Naming Conventions

*Unique names are required for the experiment name and sample name*

The experiment name and sample names must be unique for the Gate Vertex Tool (the software written by Sony to access the database of gate vertices). This is because the Gate Vertex Tool will not find database entry for the sample.

The easiest way to ensure unique names is to have a unique experiment name that includes the date.

If a sample was not completed in a particular automation run, then do one of two things, either append the name of the failed profile / sort tubes with something to identify it as failing, or append the name in the CSV file to include some way to identify it as the correct sample when it is run again.

*Using underscores in sample names for the CSV file, or experiment name*

Using underscores in the experiment name is a fatal error for using the Gate Vertex Tool (the software written by Sony to access the database of gate vertices). This is because the data CSV file will not be properly named. Create a new experiment and name the experiment without underscores. Restart the automation program (Ctrl+C, exit out of GitBash if it does not exit out of the program, and unplug and plug the USB cables).

Using underscores in the sample names will result in an error, but the underscores will be replaced with dashes in the Sony Software for the names of the samples. However, the samples will still be sorted.

#### Gating Issues

*Error message: "Tube/Sample [name] cannot be found"*

In the log file, find a line that follows: `Setting file name as: ['user1', '20210416FA', 'Sample Group - 1', 'FA404-2 profile']`. This follows the format [user, experiment name, sample group name, profile name]. If the name is what is expected and the error still occurs, the error may be because of errors when manually drawing and creating all of the gates, which may have caused errors in the exported data. Stop the automation software (Press 'F7'), unplug and replug the USB cables for the Zaber stage and agitation motors, and power cycle the power strip shown in the image for Part 1, Step 1. Restart the Cell Sorter Software. Repeat the fluidics checks. But skip the droplet calibration process which is only required daily. (If the droplet looks bad, you will have to re-calibrate). If this does not fix the issue, contact Diane/Emily.

*The terminal says 'Success.' but no new gate was drawn.*

This is likely because previous worksheet tabs were not closed when the automation software began, and the new gate was drawn on a previous worksheet. Exit the automation software (Press 'F7'), finish the full sort manually for the sample. Then, duplicate and assign (Ctrl+L, F12) the current sample. Close all other worksheet tabs other than the newest sample tube. Scroll to the new tube in the Active Experiment Panel. Restart the automation software as described in "**Restarting the automation program mid-run", Step 4**.

**Hardware Issues**

*Tube agitation is slow / not spinning*

If a tube is not spinning or the agitation is slow, exit the automation software (Press 'F7'). Remove the tube from the sample tube holder. Open the Arduino software, and navigate to the Serial Monitor (Tools > Serial Monitor). If the Arduino software says "`ERROR: Port busy`", unplug and replug the USB cables and try again.

The Serial Monitor should read 'Ready'. In the Serial Monitor, type '`S,[motor_num]`', replacing `motor_num` with the sample tube housing number of the tube that is not spinning minus 1. For example, '`S,[3]`' spins the tube in the position in the sample tube housing that is labeled 4. Place a tube in the sample tube housing to observe whether an empty sample tube spins. If it does not spin contact Diane and/or Emily.

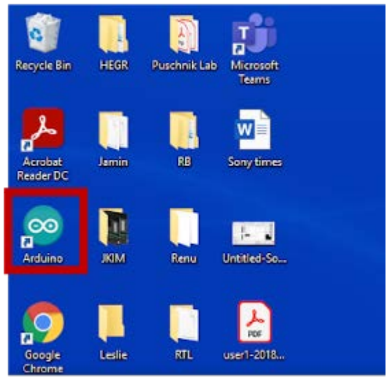


Skip that location for all subsequent sorting experiments (until the issue can be addressed), place the sample tube into a different location in the sample tube housing, and update the sample CSV file with the new position.

Exit the Serial Monitor and the Arduino software. Restart the automation software.

*Temperature reading errors*

If the temperature readings raise errors (by being outside the expected temperature range) for two consecutive readings (20 minutes) after reaching equilibrium, find a temperature probe / thermometer and measure the temperature within the sample tube housing at several locations. There is usually a thermometer in the TC room next to the sink. Note that variation of about 3-5 °C is normal within different areas of the sample tube housing.

If the temperature is accurate between the thermometer and the automation readings, please contact Diane/Emily before continuing to adjust the cold air flow. If the temperature readings do not match, and the temperature probe / thermometer is within the desired temperature range (2-8 °C), then there is a problem with the automation software. The temperature error can be disregarded during the experiment. Inform Diane/Emily once the experiments are complete to repair the system.

#### Part 7: Cleaning & Maintenance

To date, the automation system requires little cleaning and maintenance, though this may change with more frequent use. Below outlines the known for cleaning and maintenance needs.

##### Tube Housing Gasket Cleaning

At times a white powdery substance appears on the gasket around the slots where the tubes are inserted in the two rows of the tube housing. This is caused by wear of the sample tubes by the tube sockets. (It is much less of an issue with new teflon tube sockets.) It can lead to the sample tubes wiggling out of the tube sockets while being shaken.

Clean the gasket using a shammy cloth (located inside the top drawer storage unit next to the computer) moistened with DI-water.

**Gripper Claw: Reapplication of Rubber Cement**

The inner gripping surface of the claws are coated with Pliobond 25 rubber cement to provide some adhesive friction to grab the sample tubes. When there is noticeable slipping of the sample tubes when the gripper attempts to grab the tubes, the rubber cement needs to be reapplied. Apply several coats of Pliobond 25 rubber cement to the inner gripping surfaces of the claws ensuring that none is on the arms of the claws. It might be necessary to reapply the shrink tubing to the claws. Contact Diane/Emily for assistance.

### FACS Automation Setup Flowchart

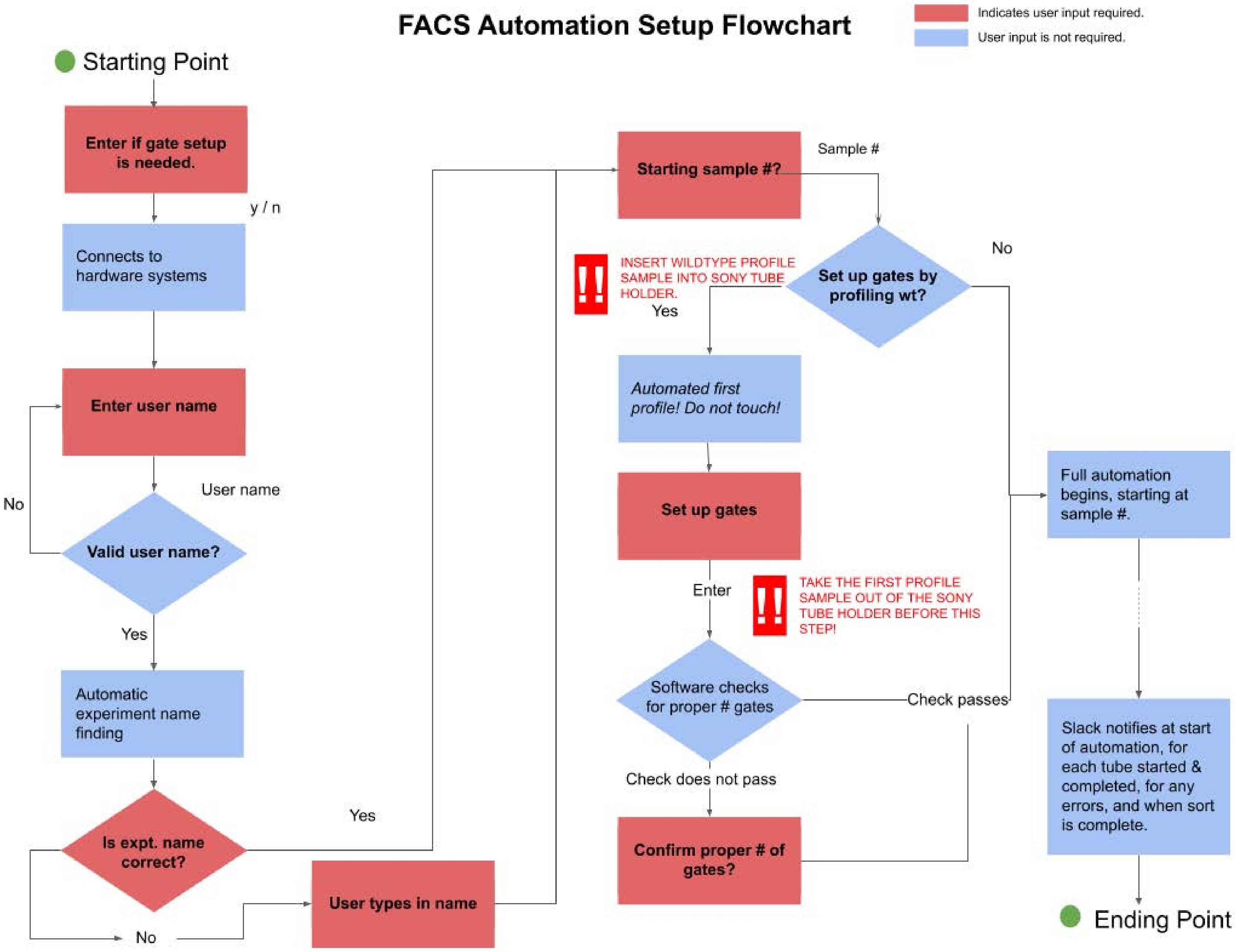
