## Supplementary material for "An open-source FACS automation system for high-throughput cell biology": Automation Controller Software Instructions

### FACS Automation Controller Software

Click through control of Sony Cell Sorter software

Last Updated: 2023-02-23

### Startup sequence

General

### Prepare the Cell Sorter Software GUI.

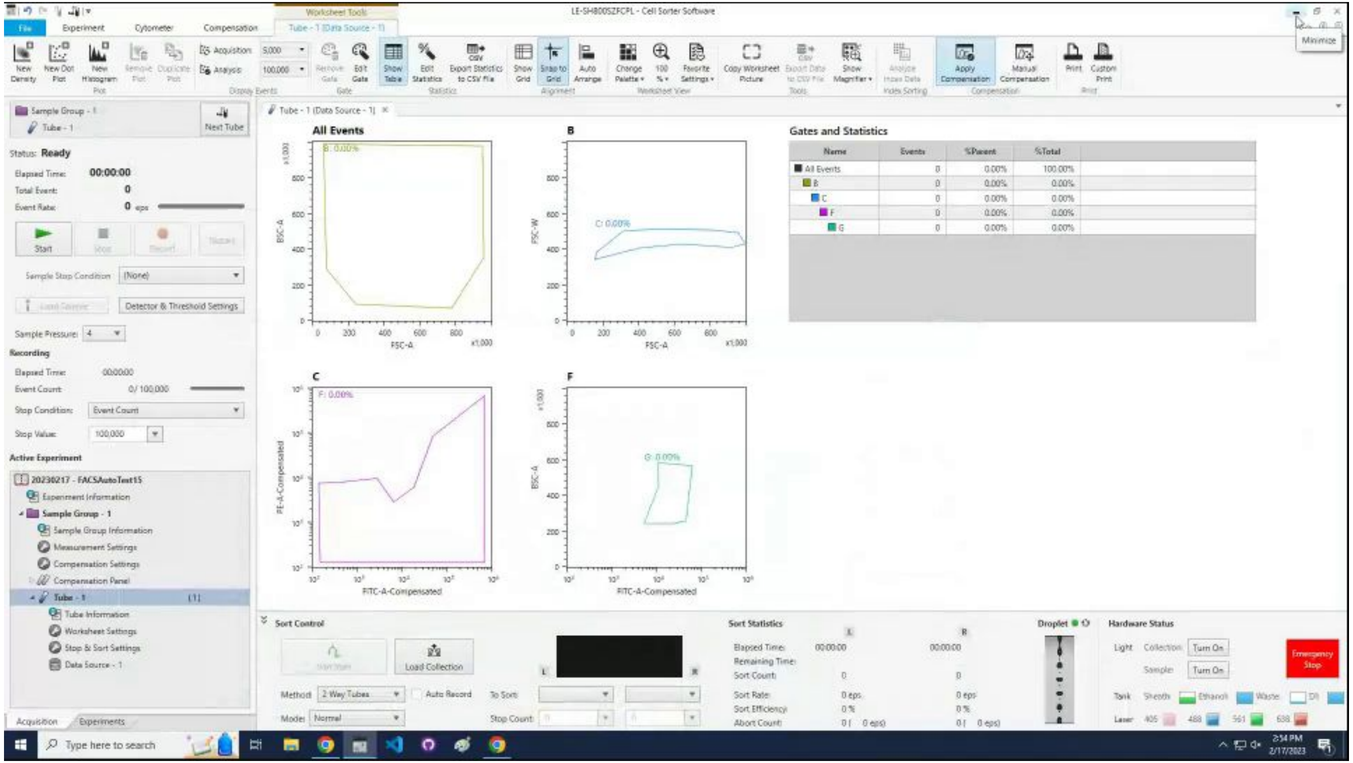

### Open the solenoid valve by clicking `cool`.

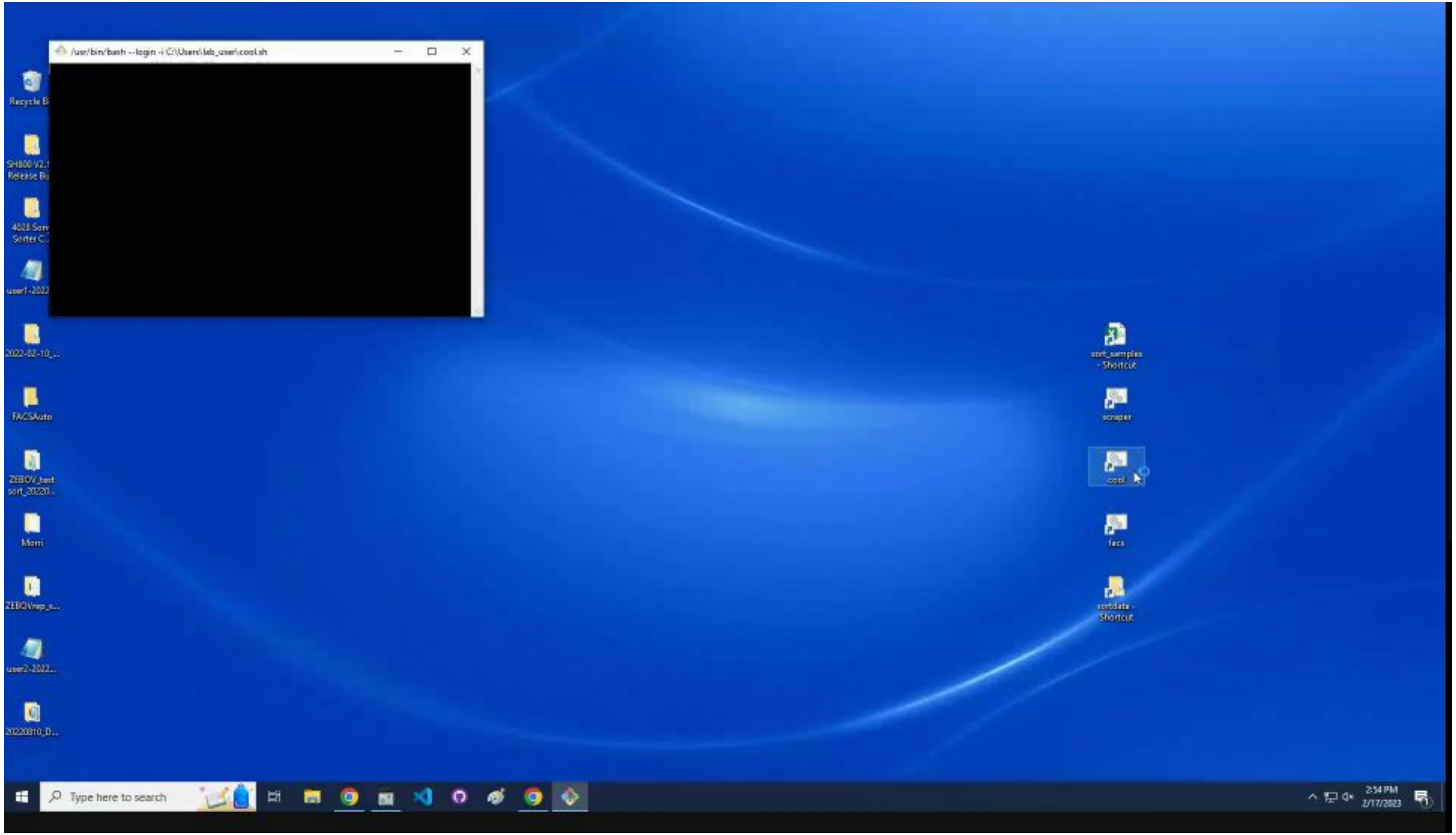

### Open the worklist `sort_samples.csv` file.

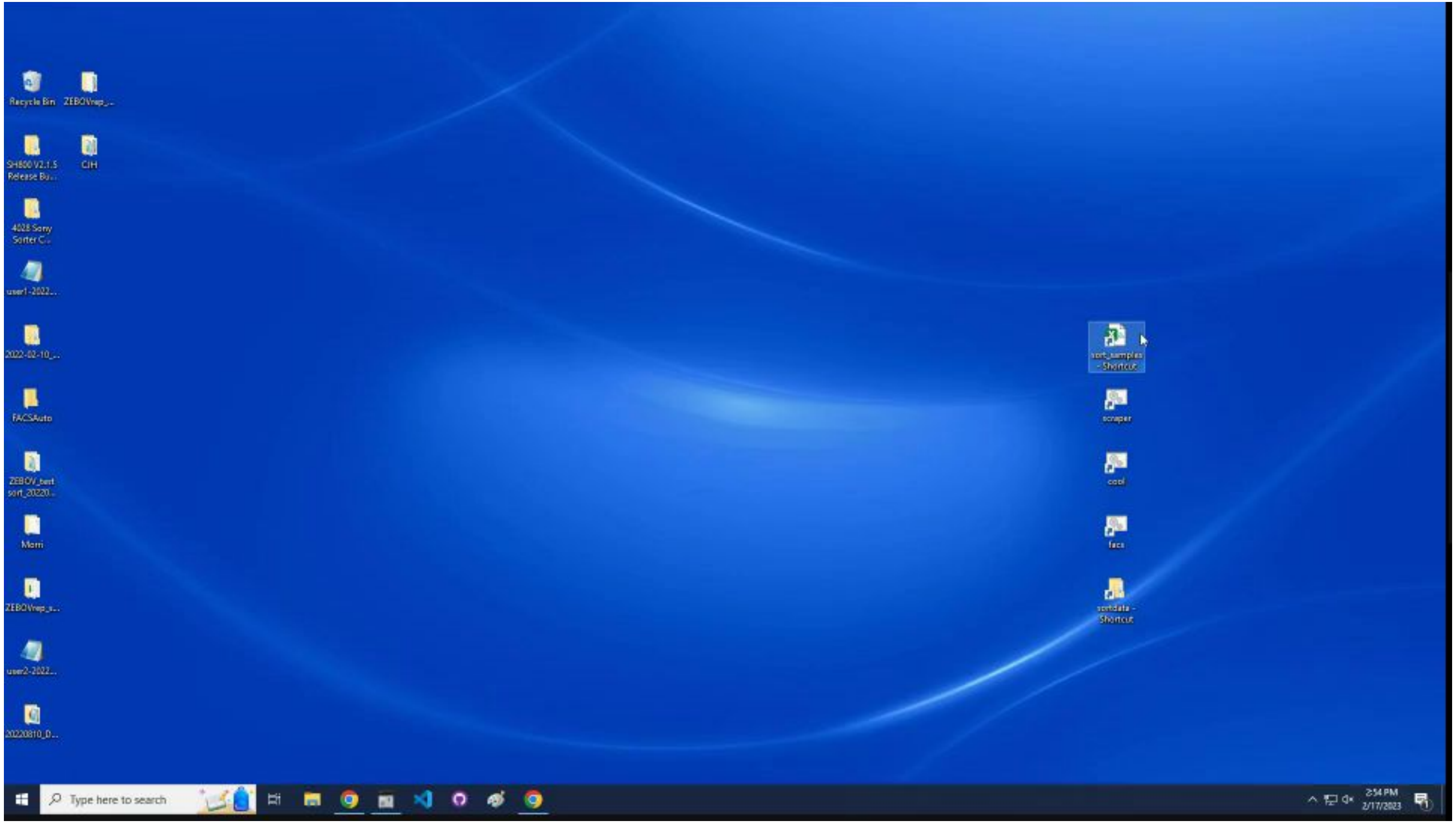

### Fill in tube location, sample name, and well ID, and save.

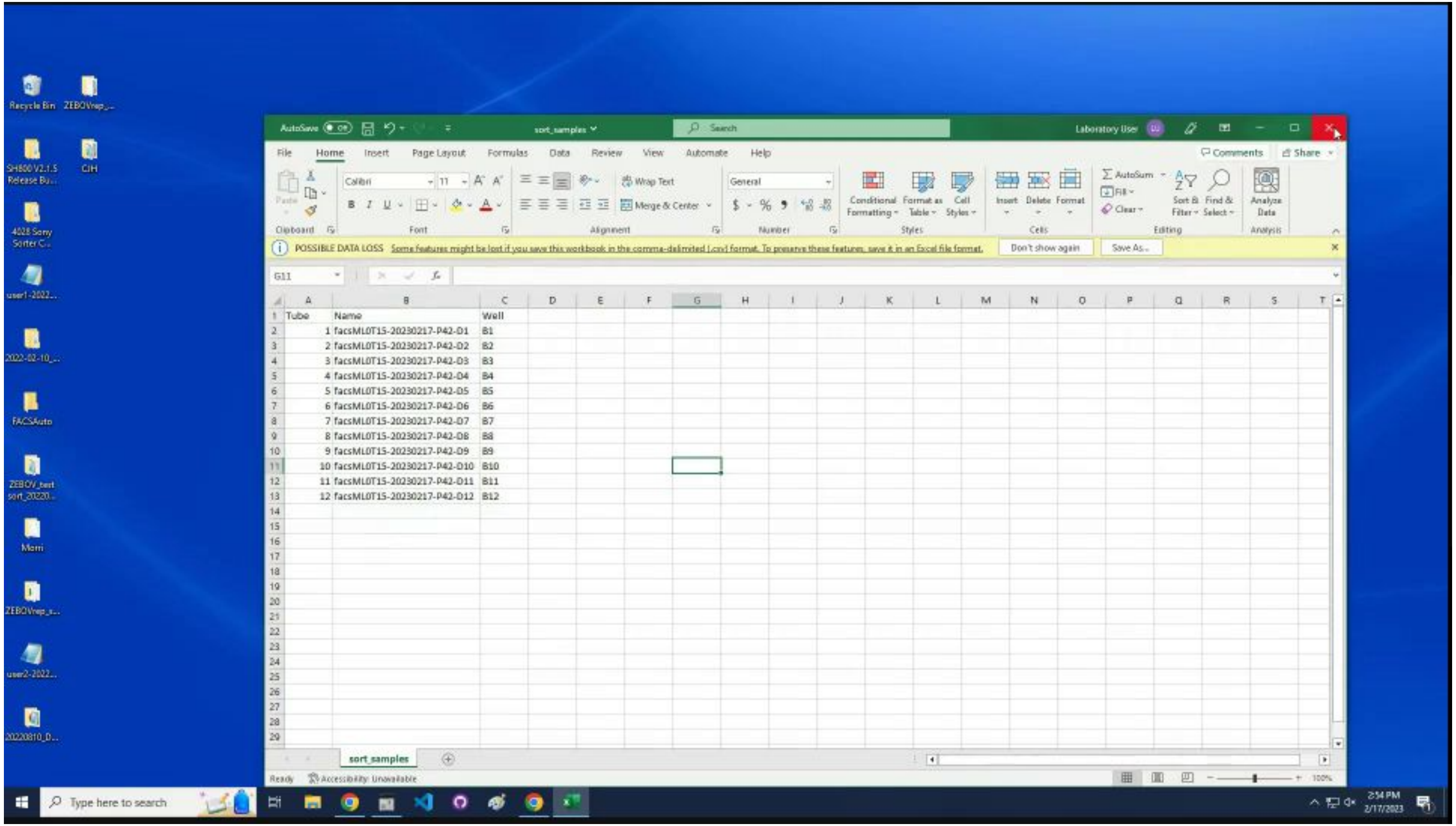

Start the automated sorting by clicking `facs`.

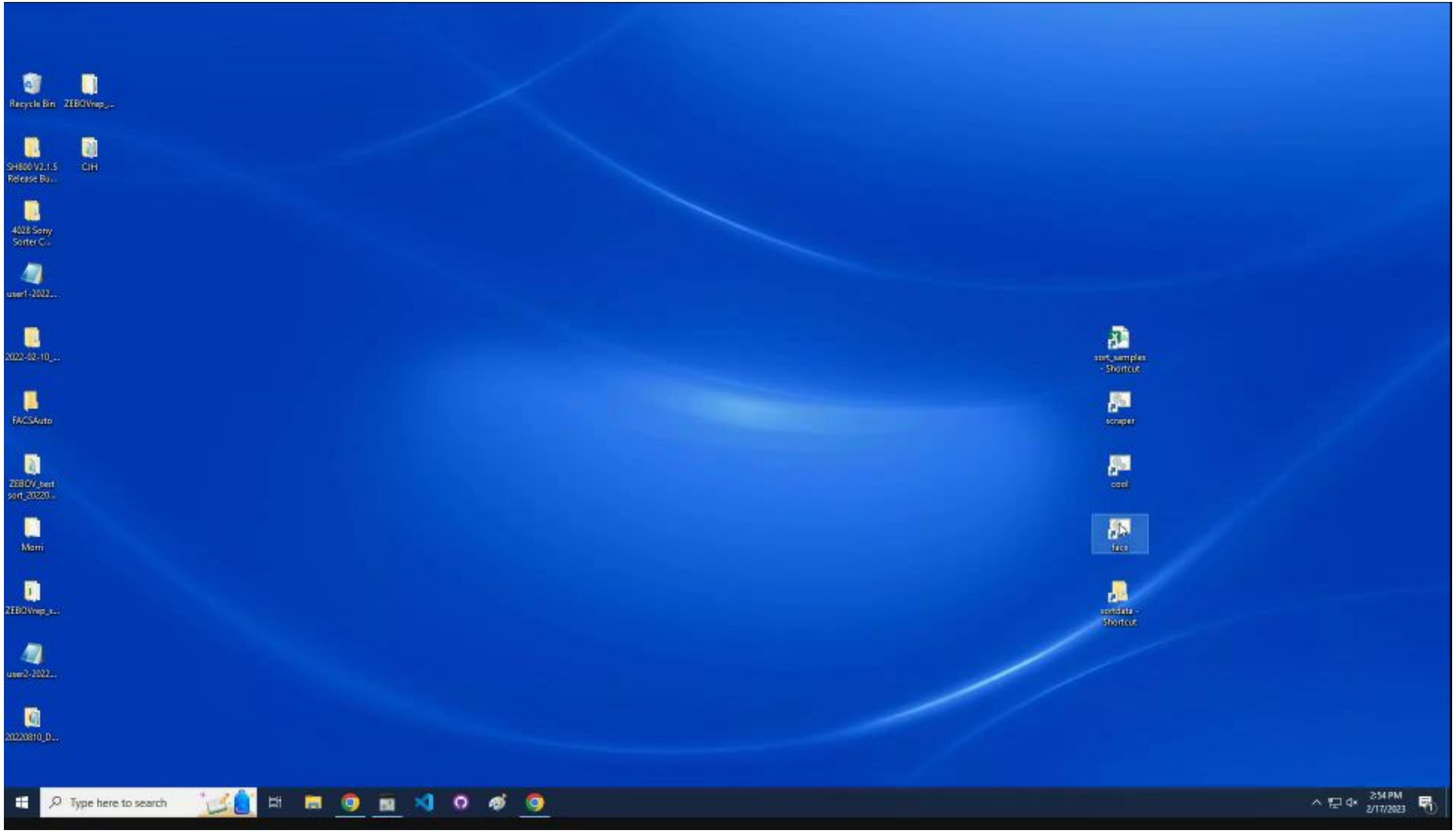

### Startup sequence

Need to set up gates

### Enter `y` to profile a sample to set up the gates.

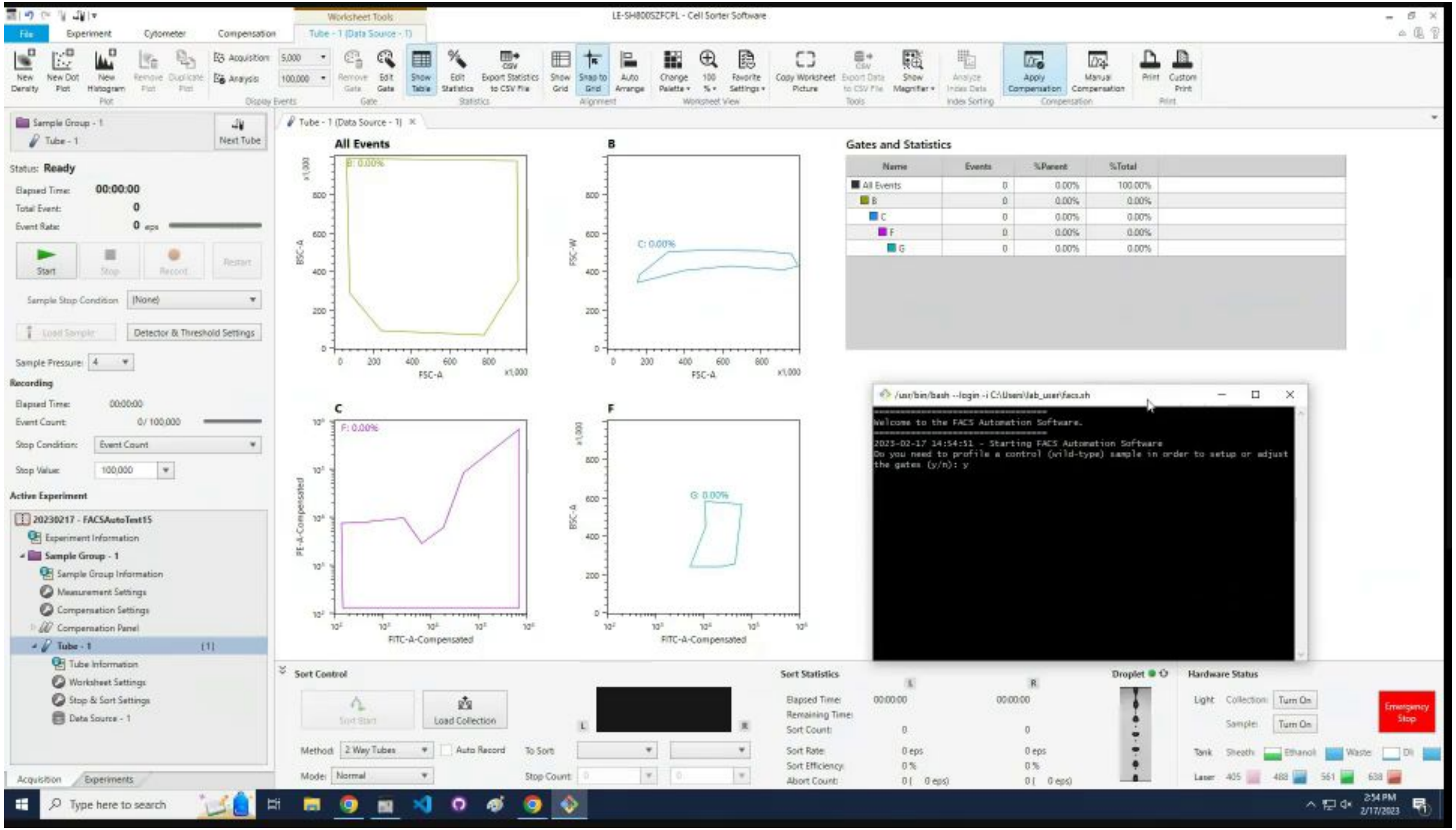

### Enter `user name.`

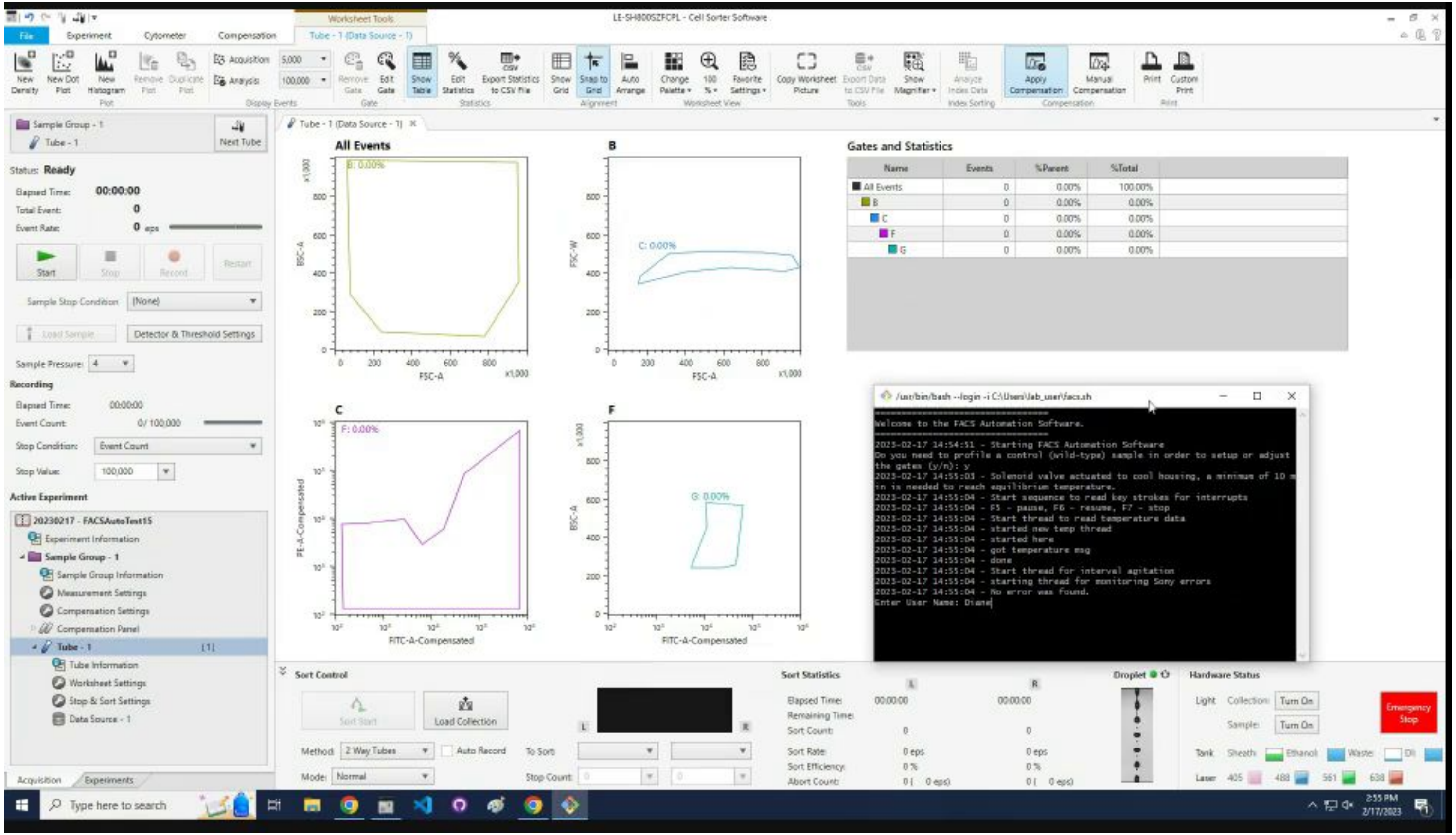

### Find experiment name to enter into program.

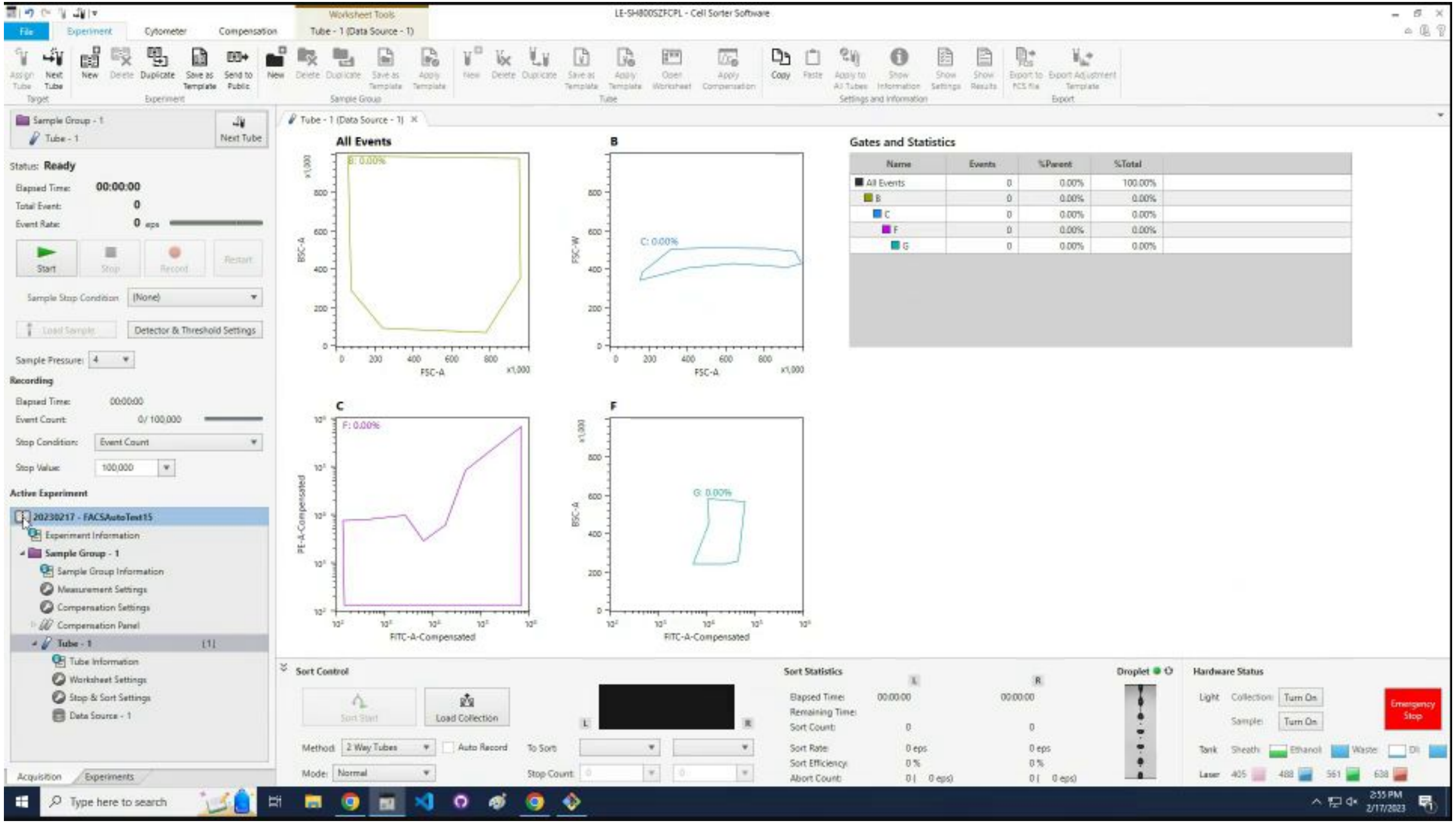

### Confirm that the experiment name is correct.

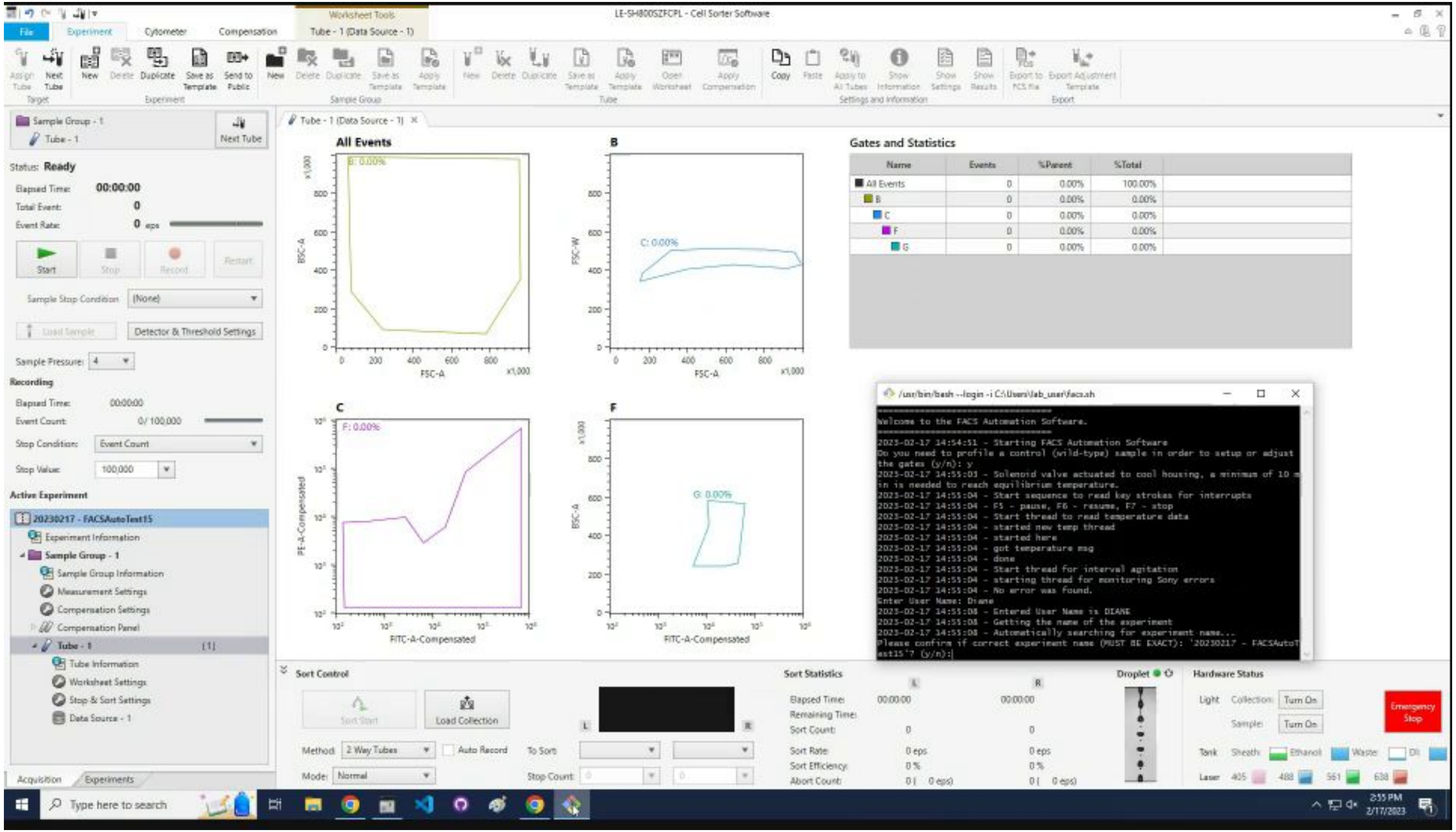

### Enter the first tube number to start the sorting process.

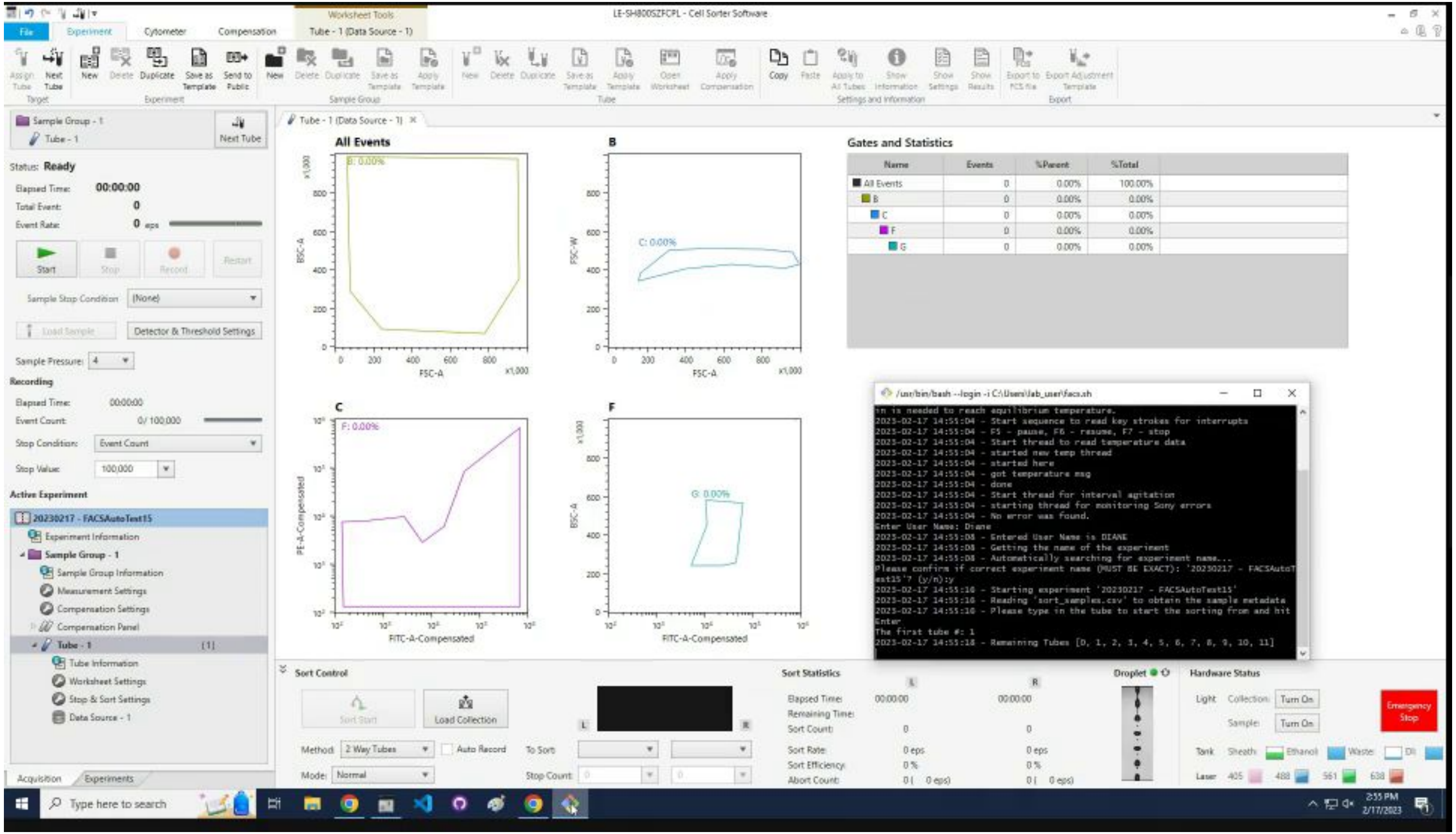

### Automated sorting starts to set up the gates.

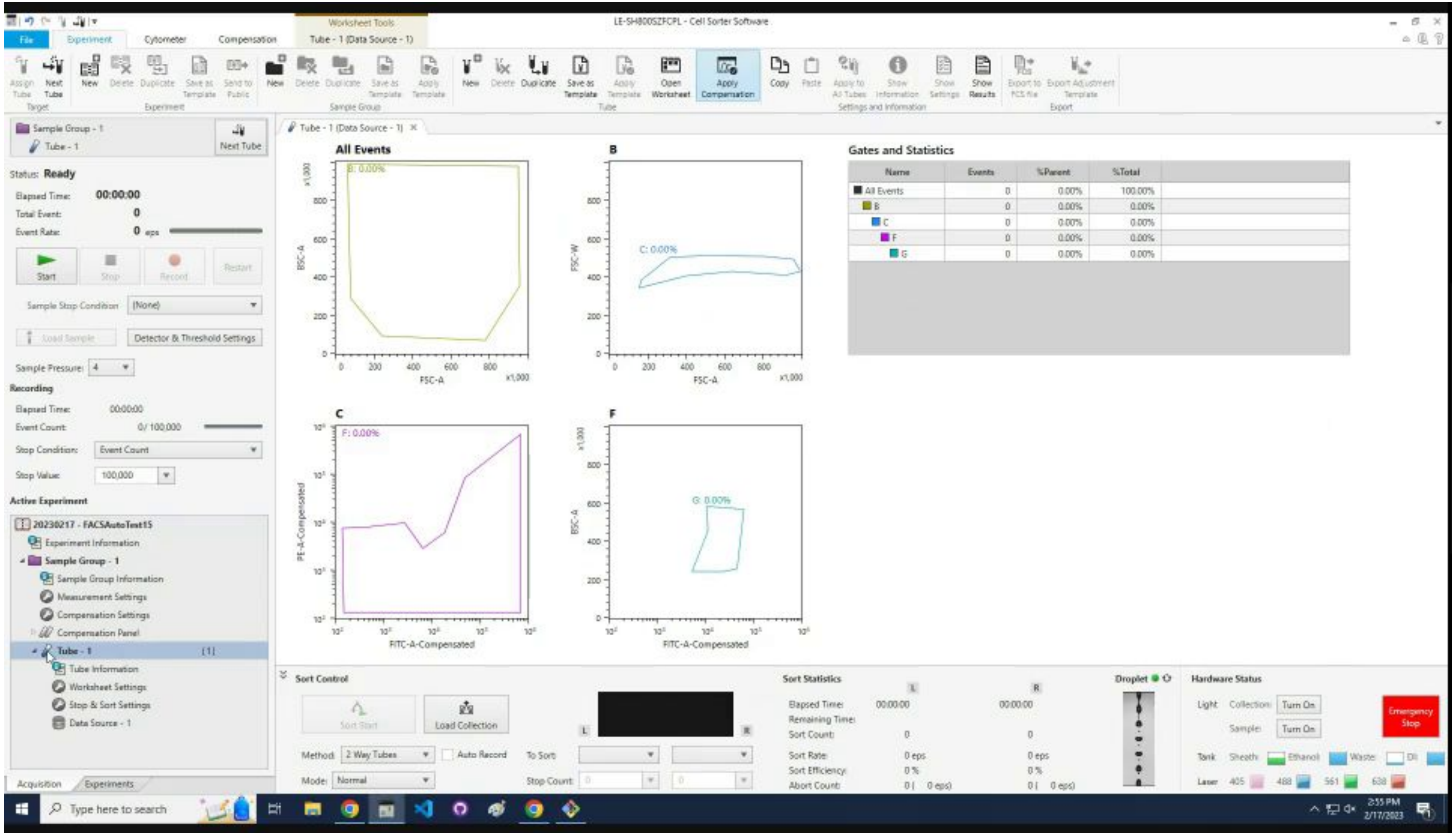

### Change the `Stop Value` to `50,000`.

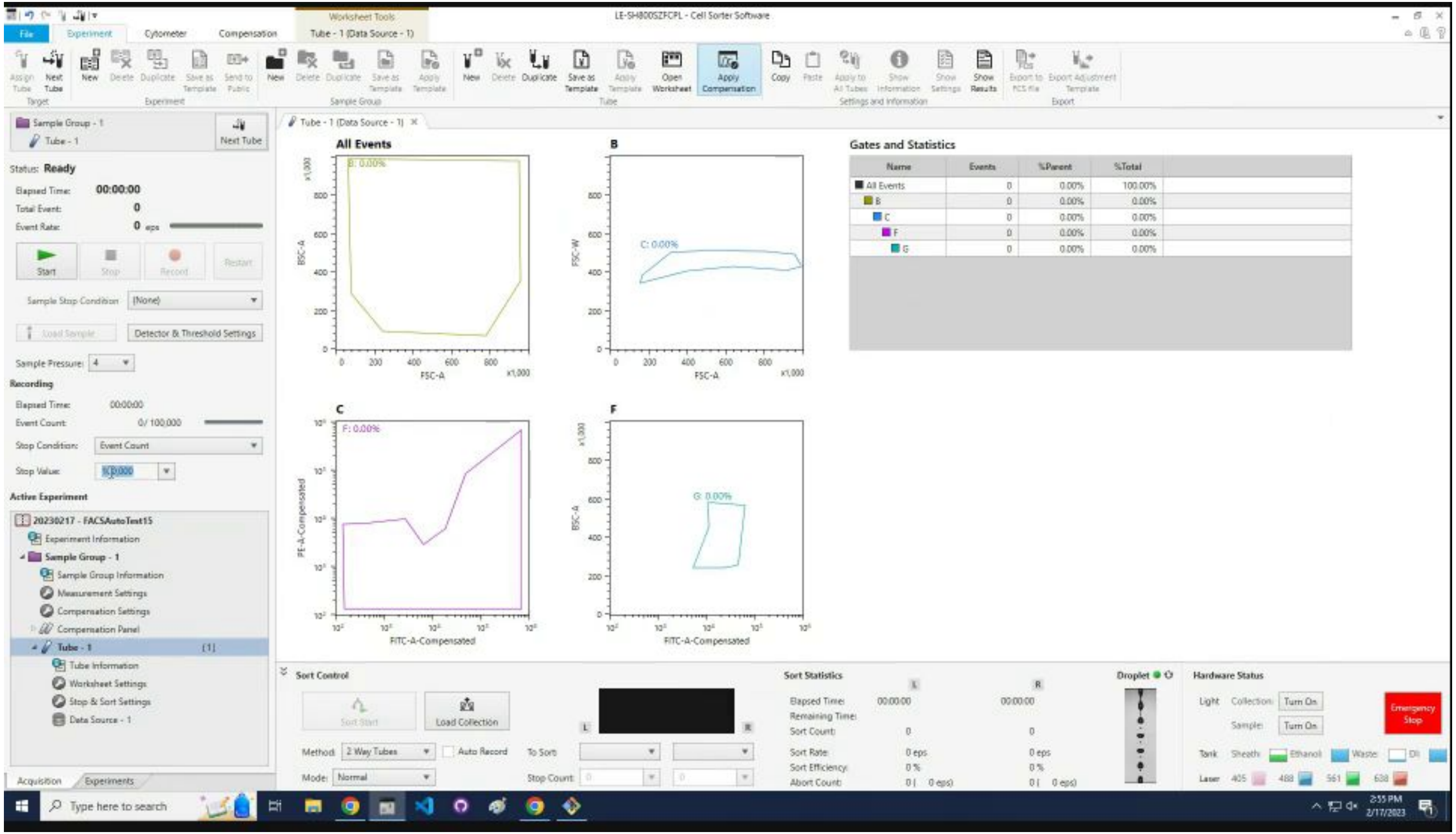

### Change the sorting `Method` to `96 Well Plate`.

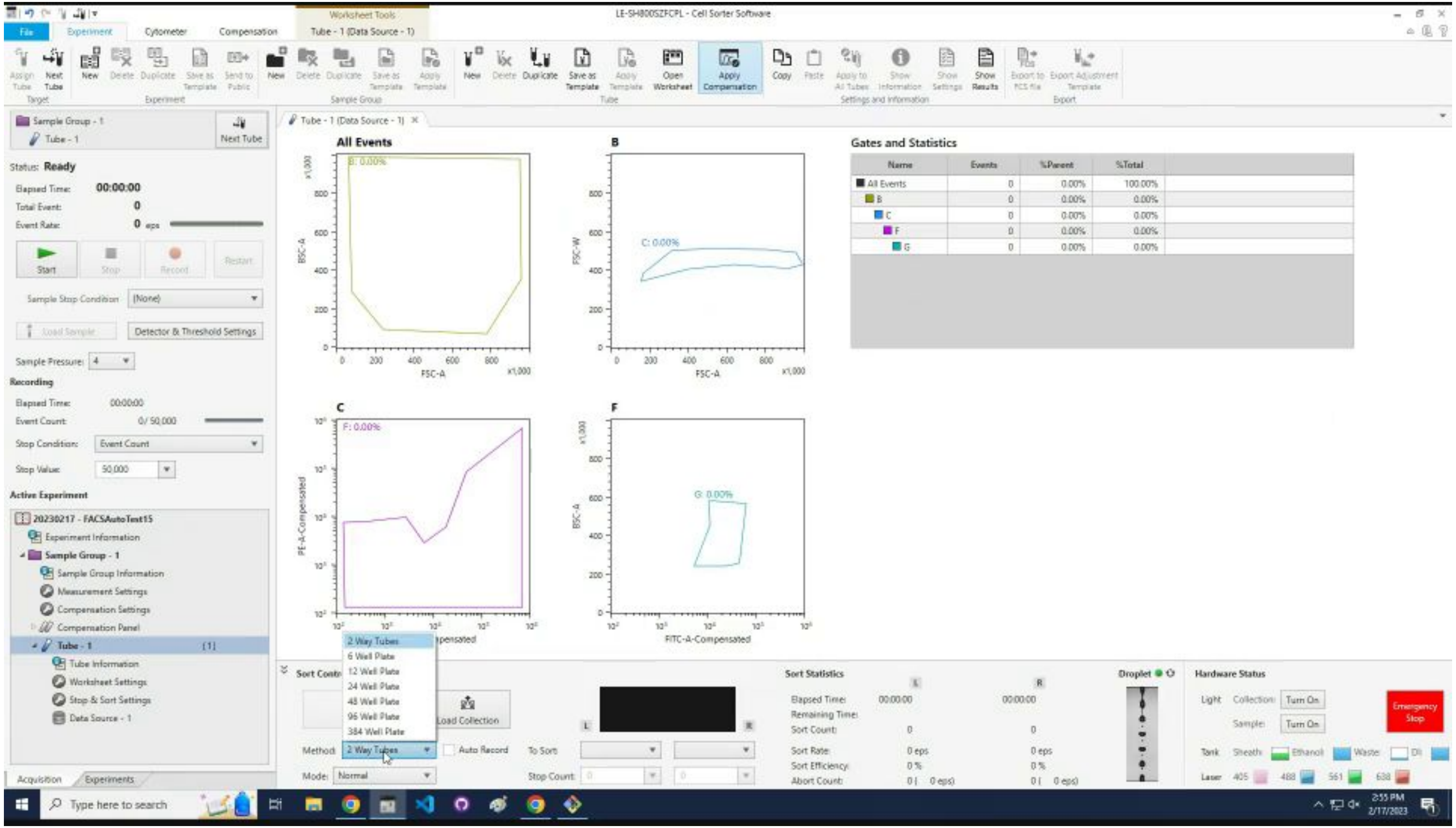

### Change the `Stop Condition` to `Event Count`.

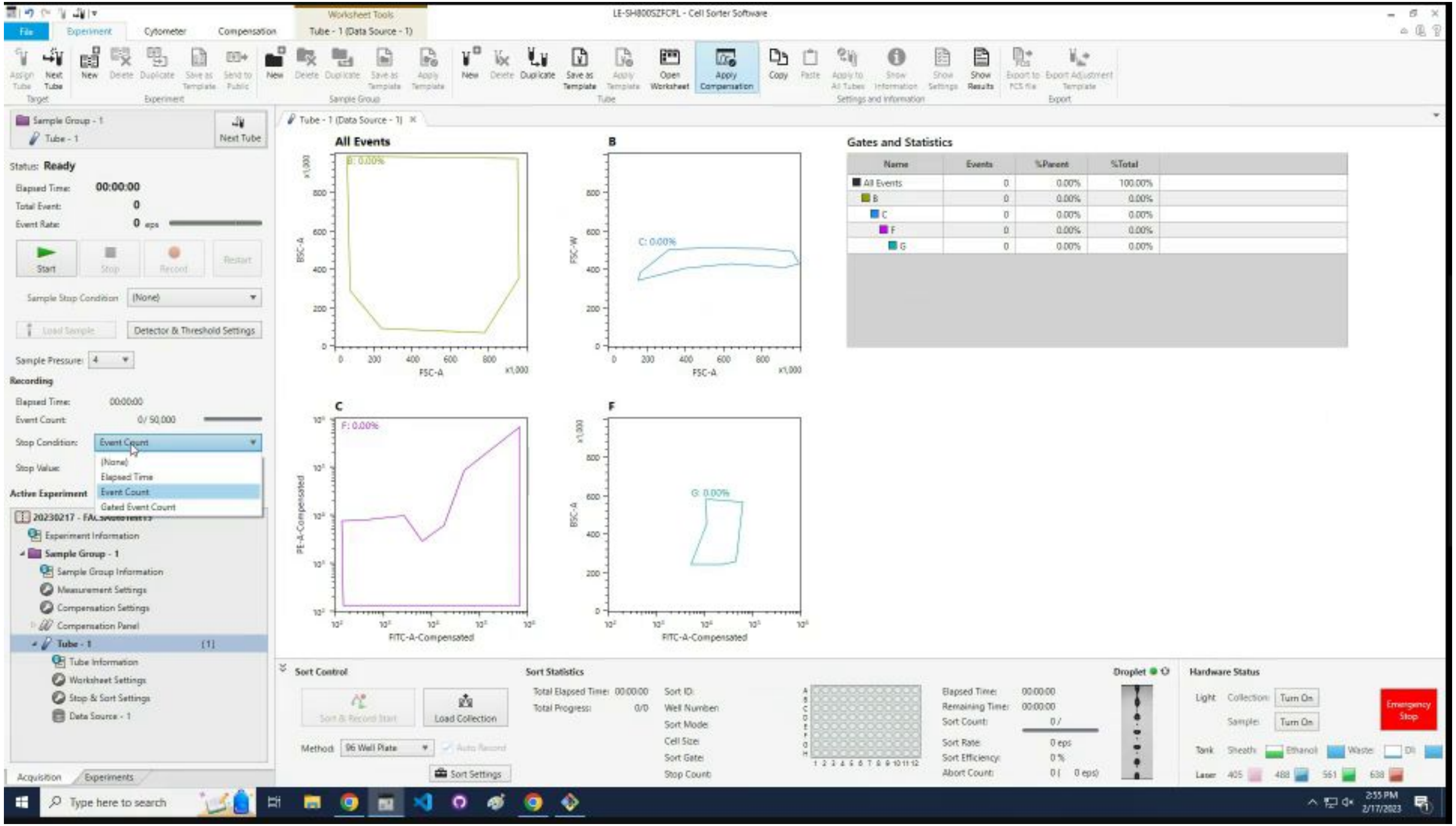

### Change the `Sample Pressure` to 5.

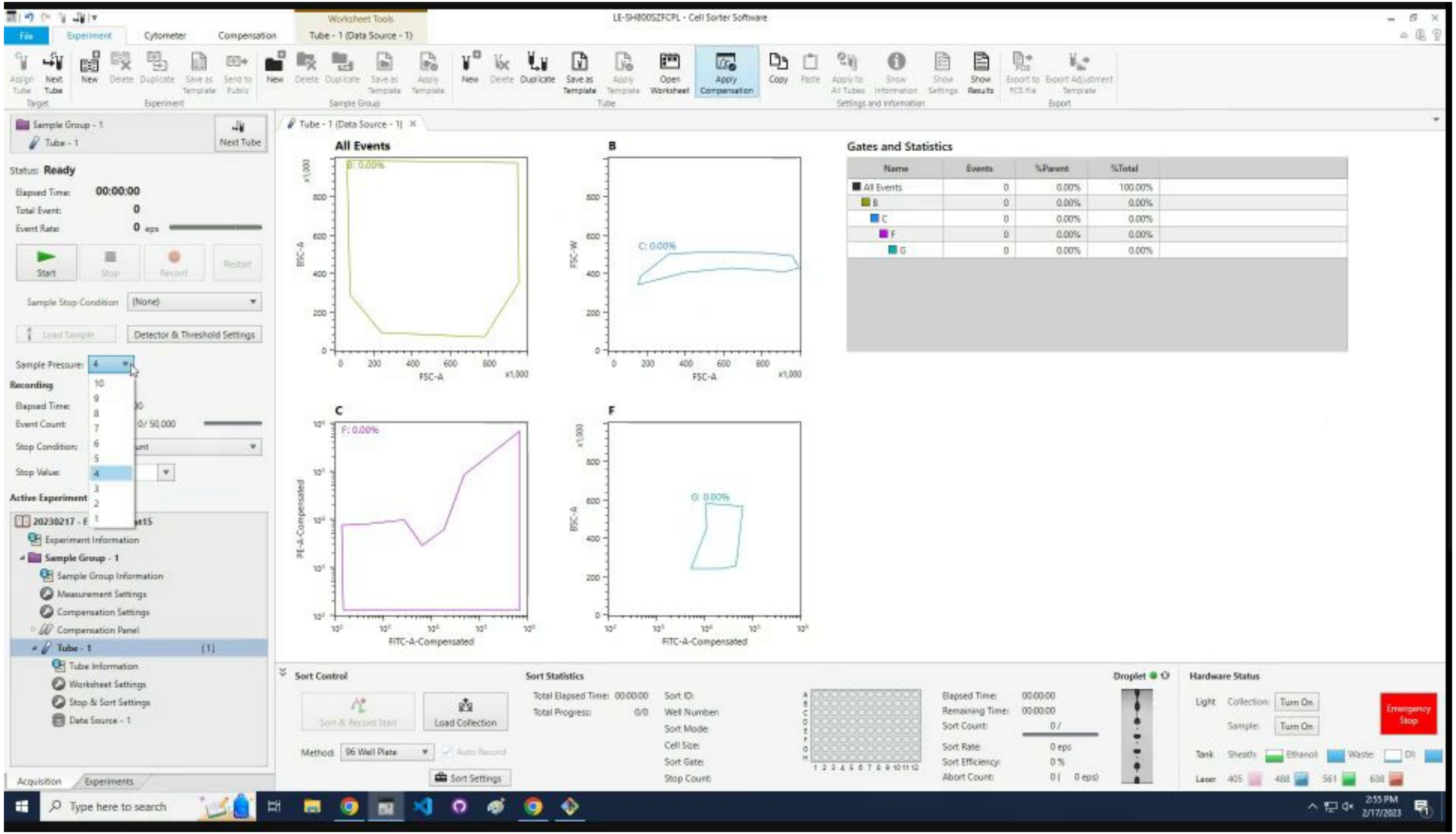

### Click `Start` to begin the profile.

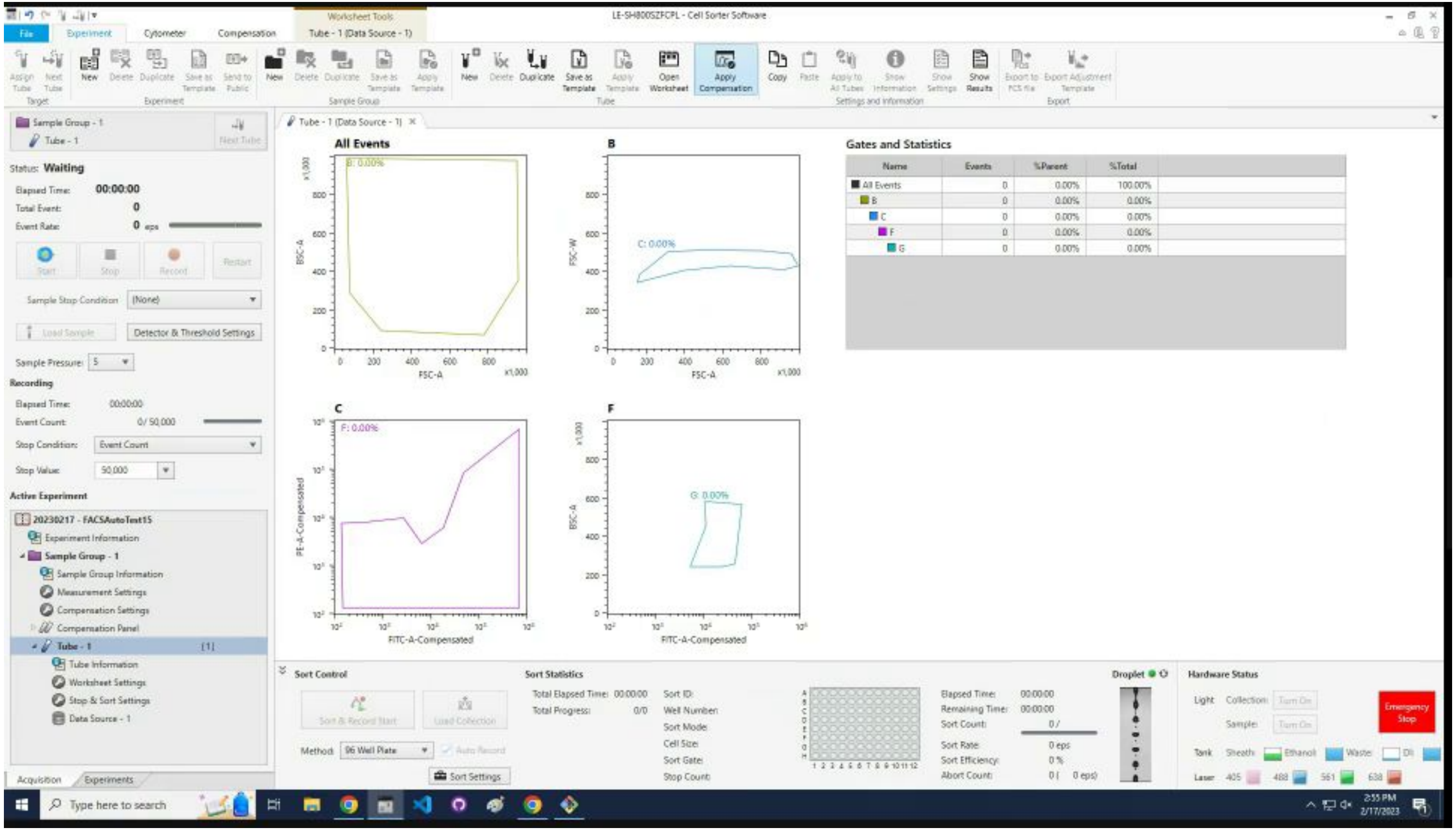

### Wait ~20 seconds for cells to appear. Click `Record`.

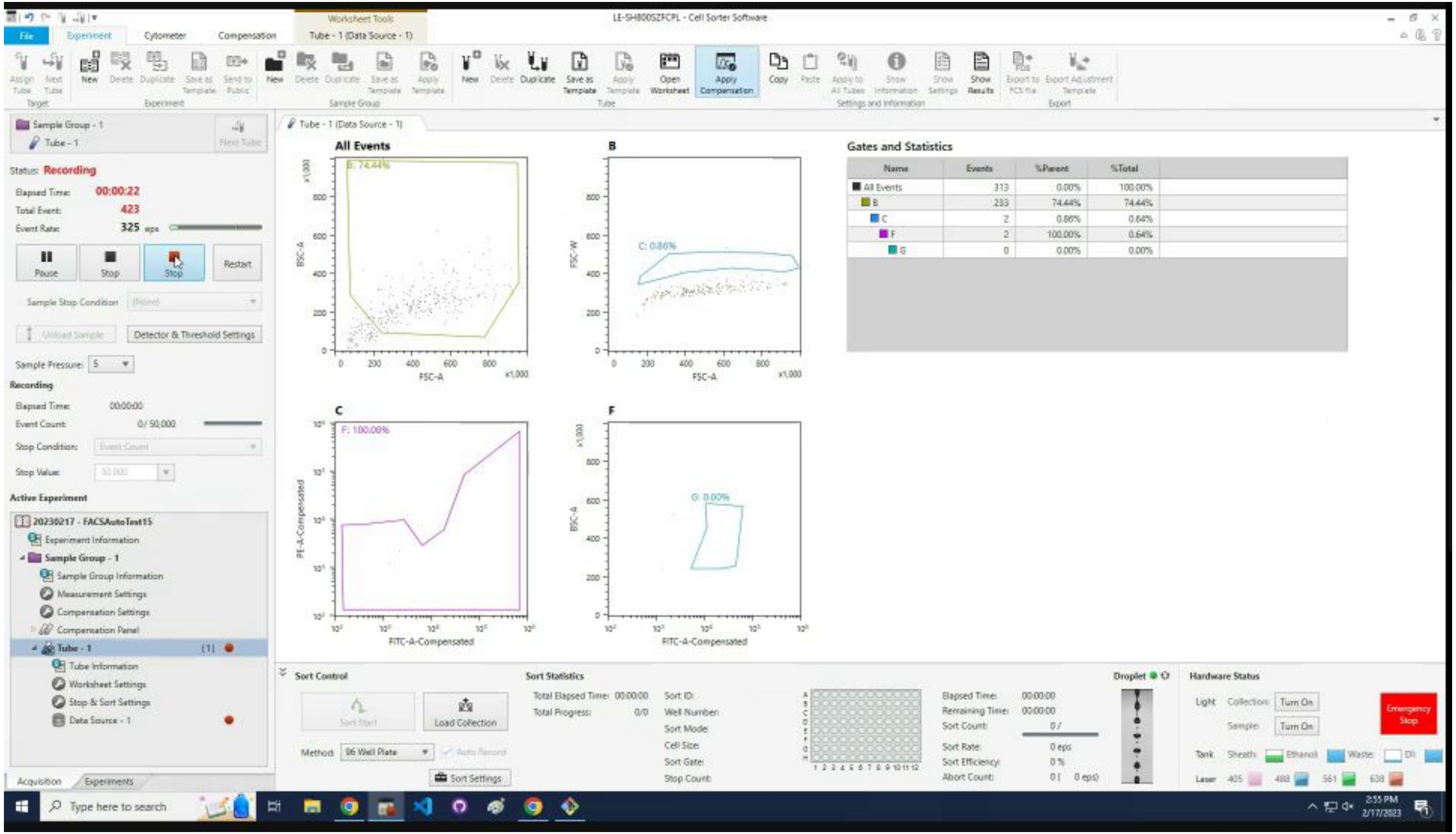

### Wait for profiling to finish then click `OK`.

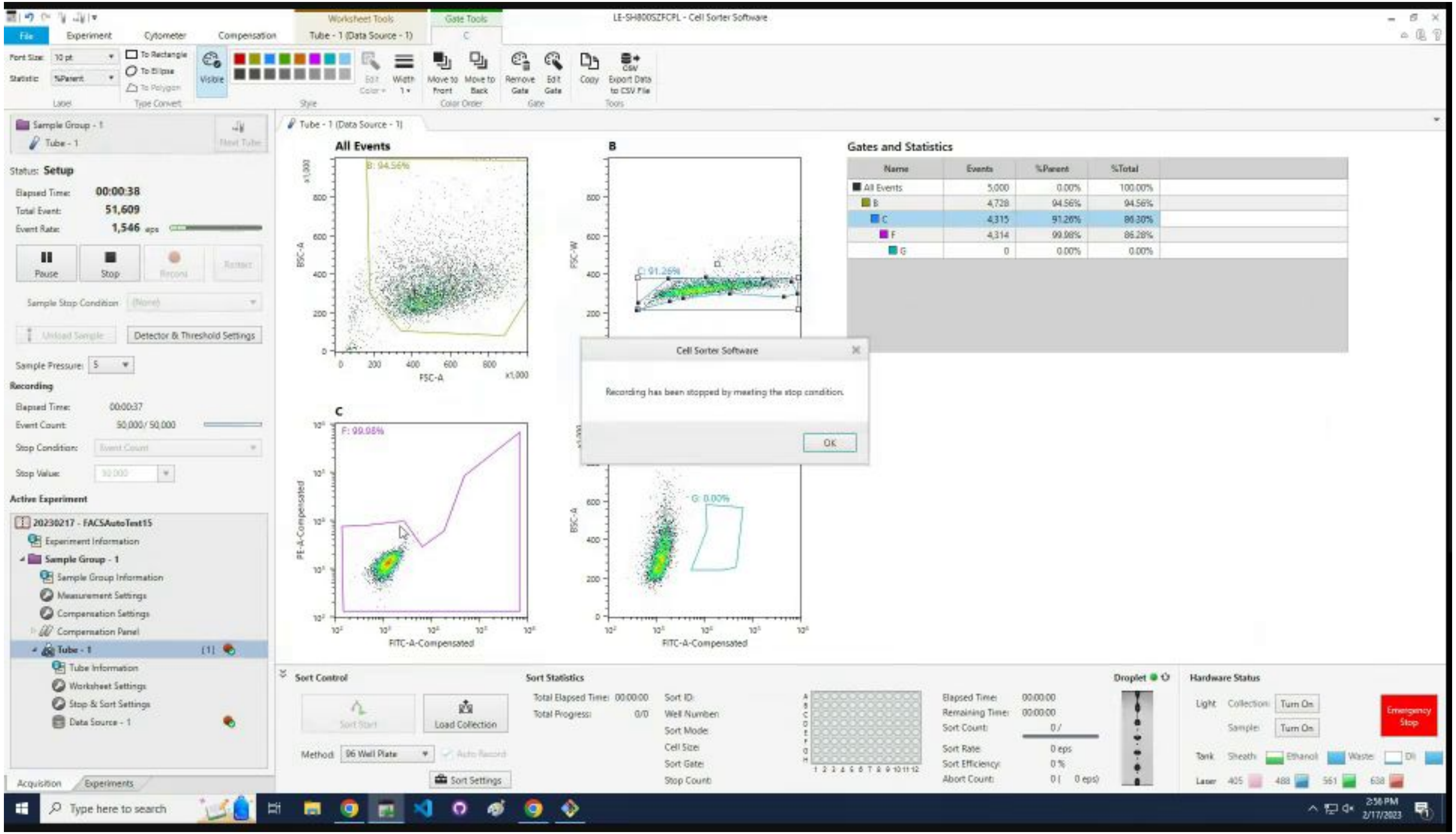

### Click `Stop`.

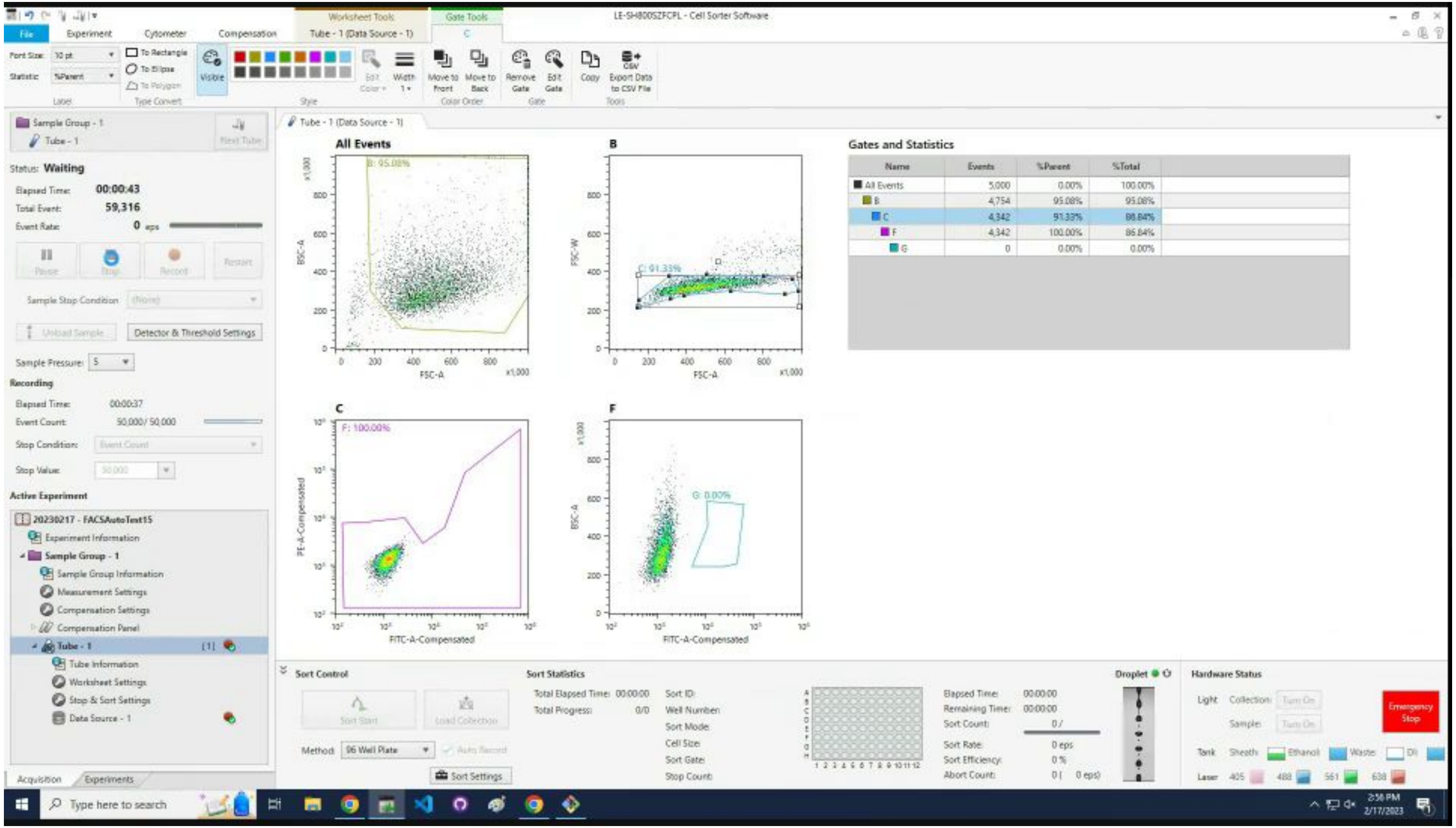

LE-SH800SZFCPL - Cell Sorter Software
All Events
B
C
F
Gates and Statistics
Status: Waiting
Elapsed Time: 00:00:43
Total Event: 59,316
Sort Control
Sort Statistics
Hardware Status
Emergency Stop

### Adjust the first three gates.

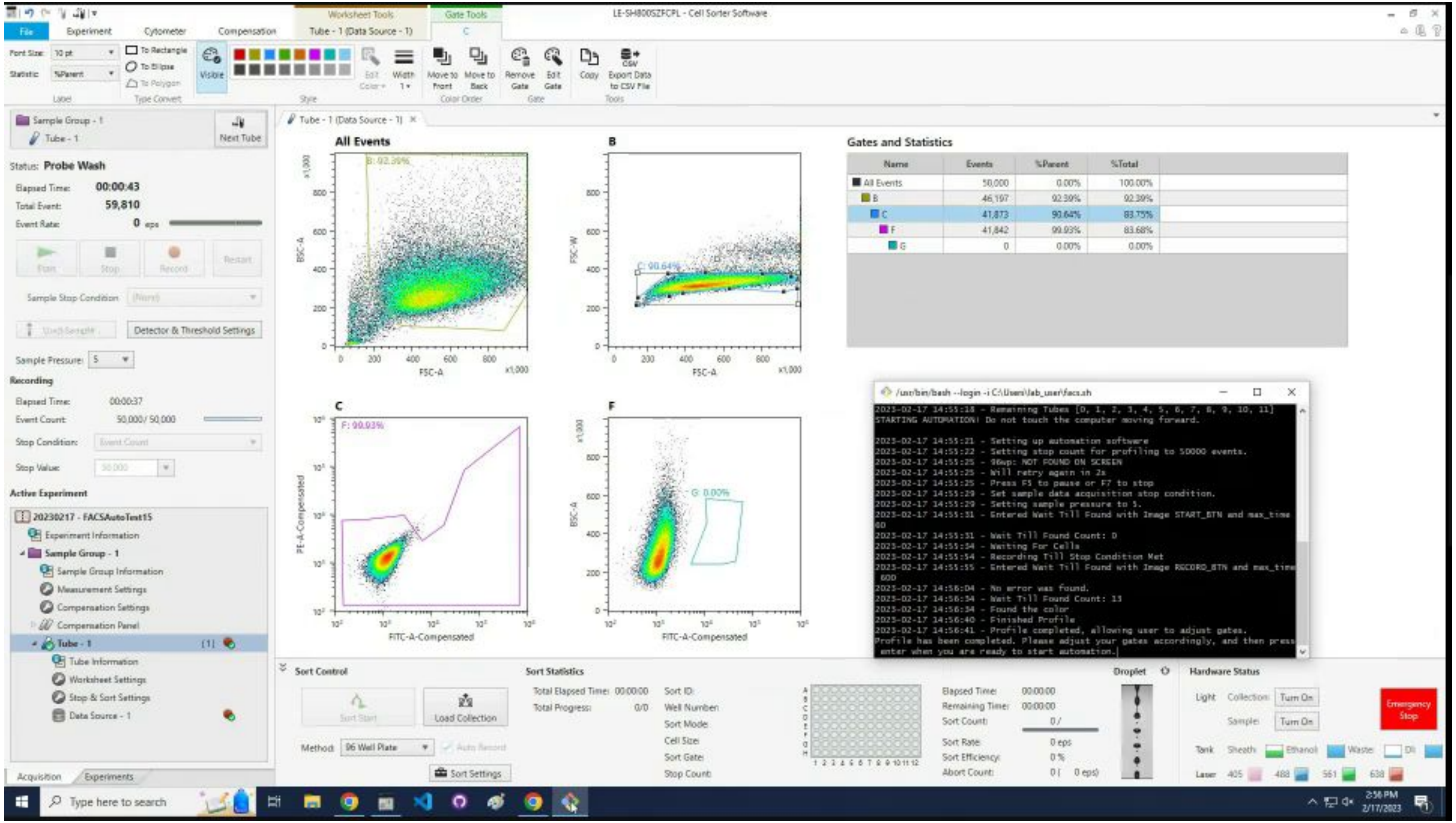

### Click the X to close the tube tab.

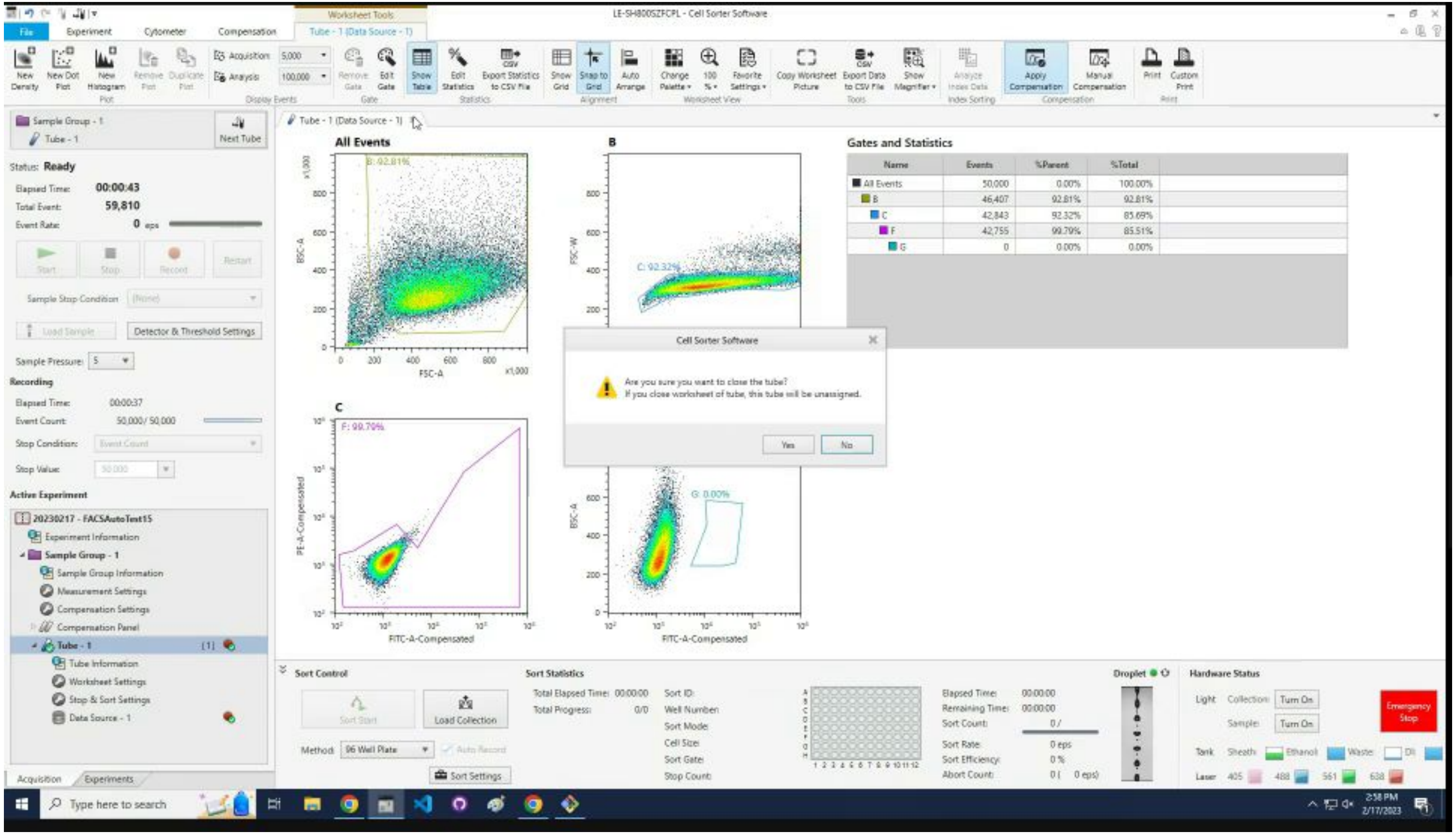

### Click `Yes` to confirm closing the tube tab.

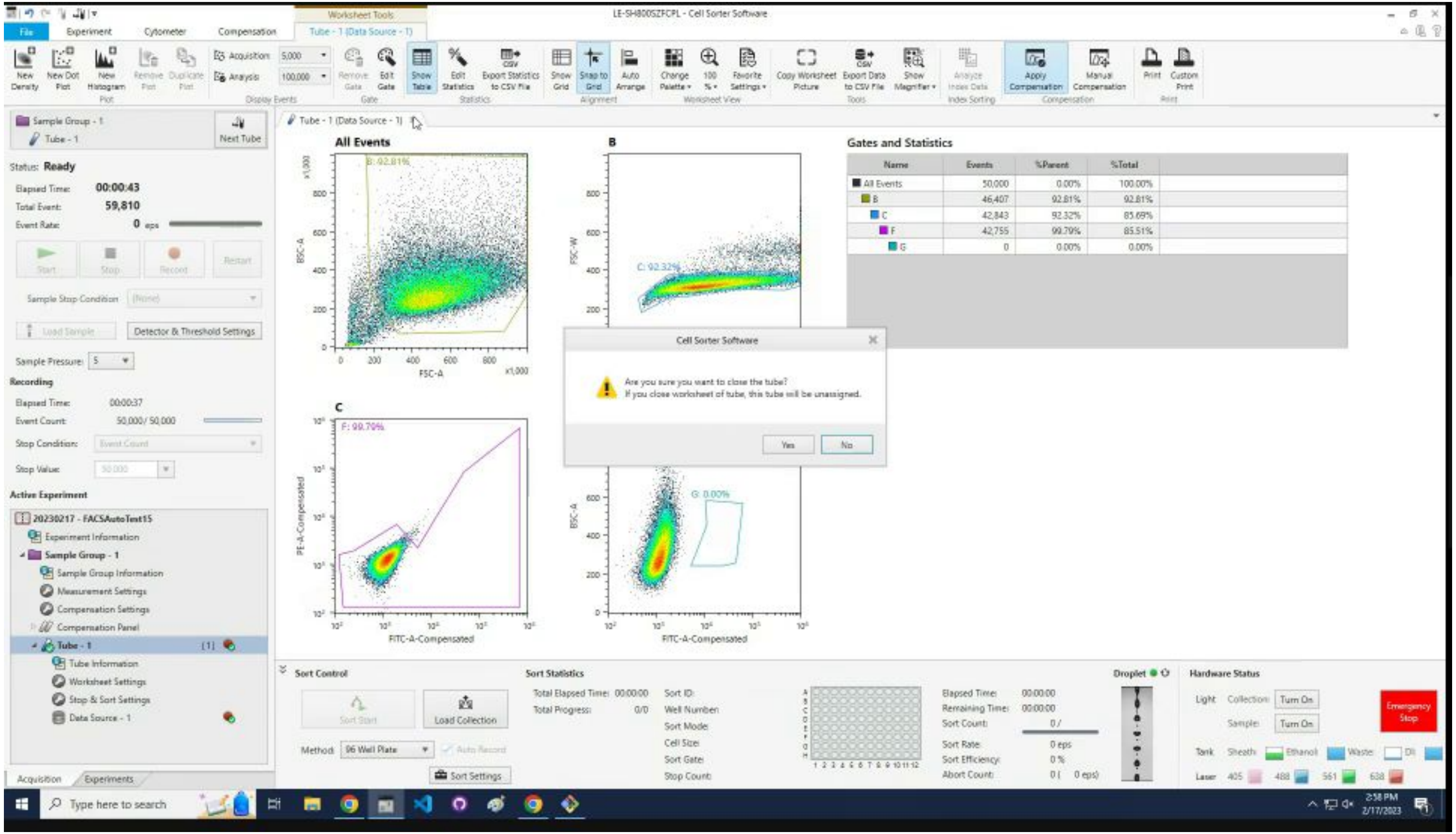

### Click on the profile sample in the list.

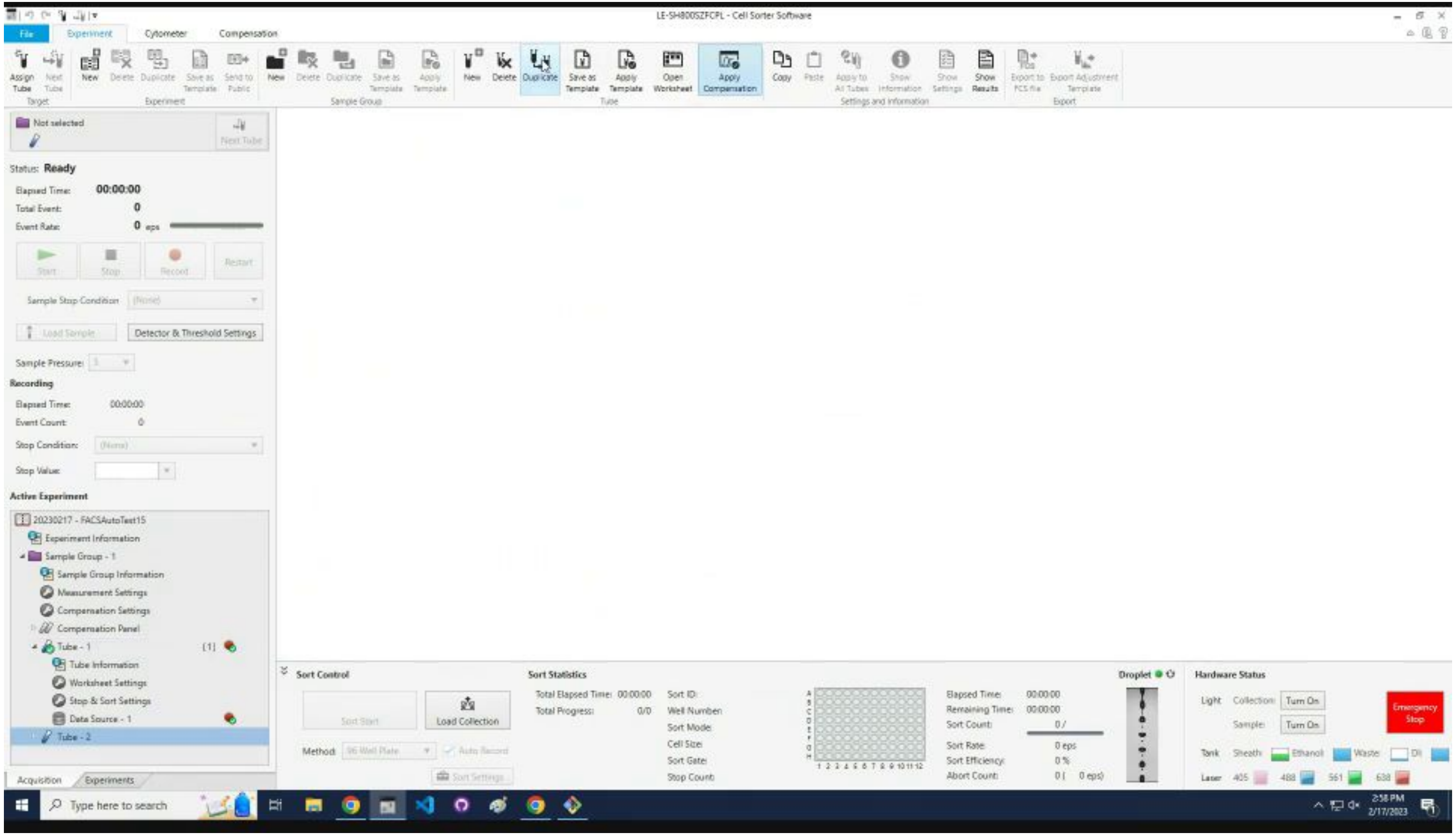

### Click the `Duplicate` icon.

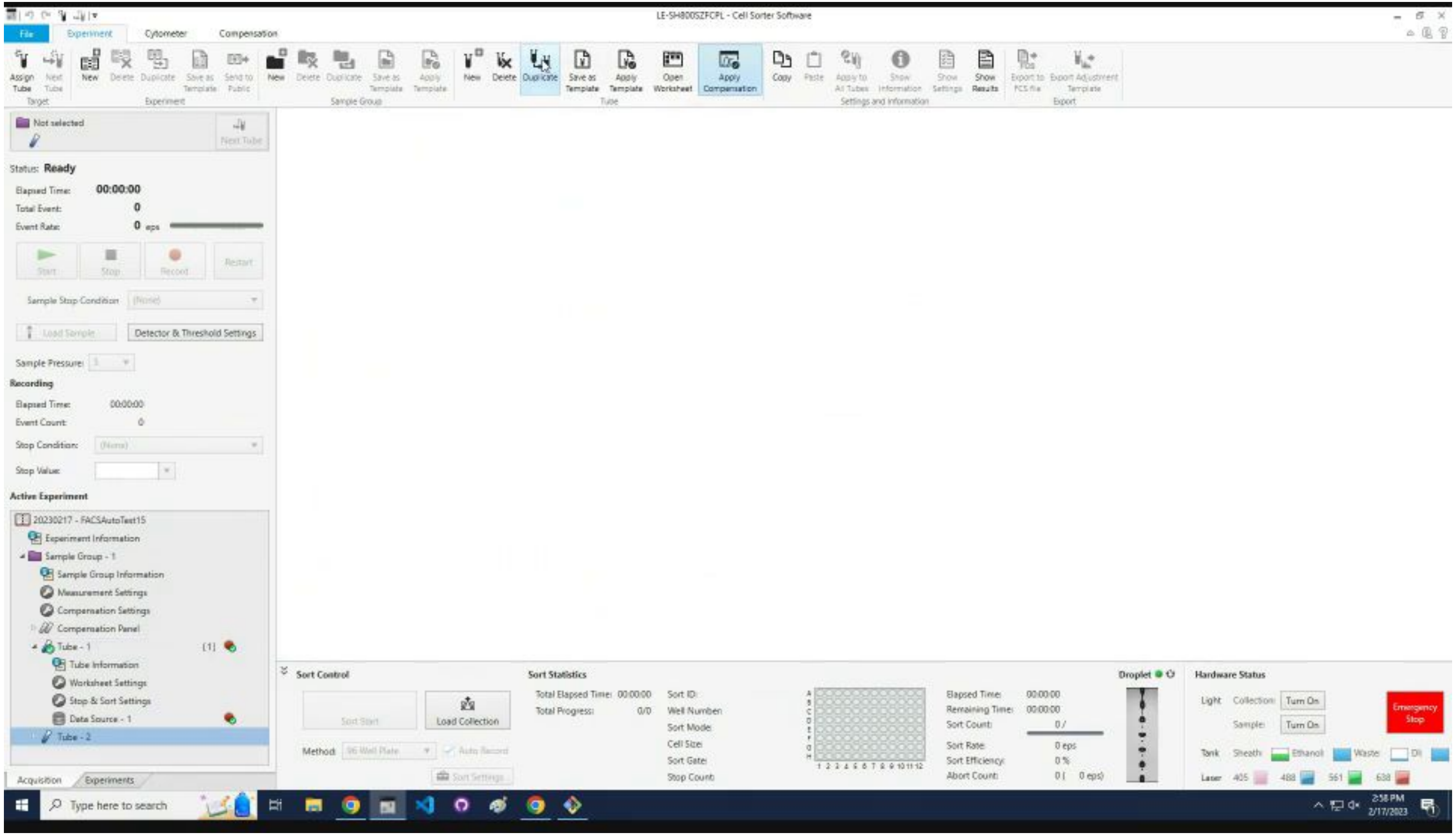

### Click the `Assign Tube` icon.

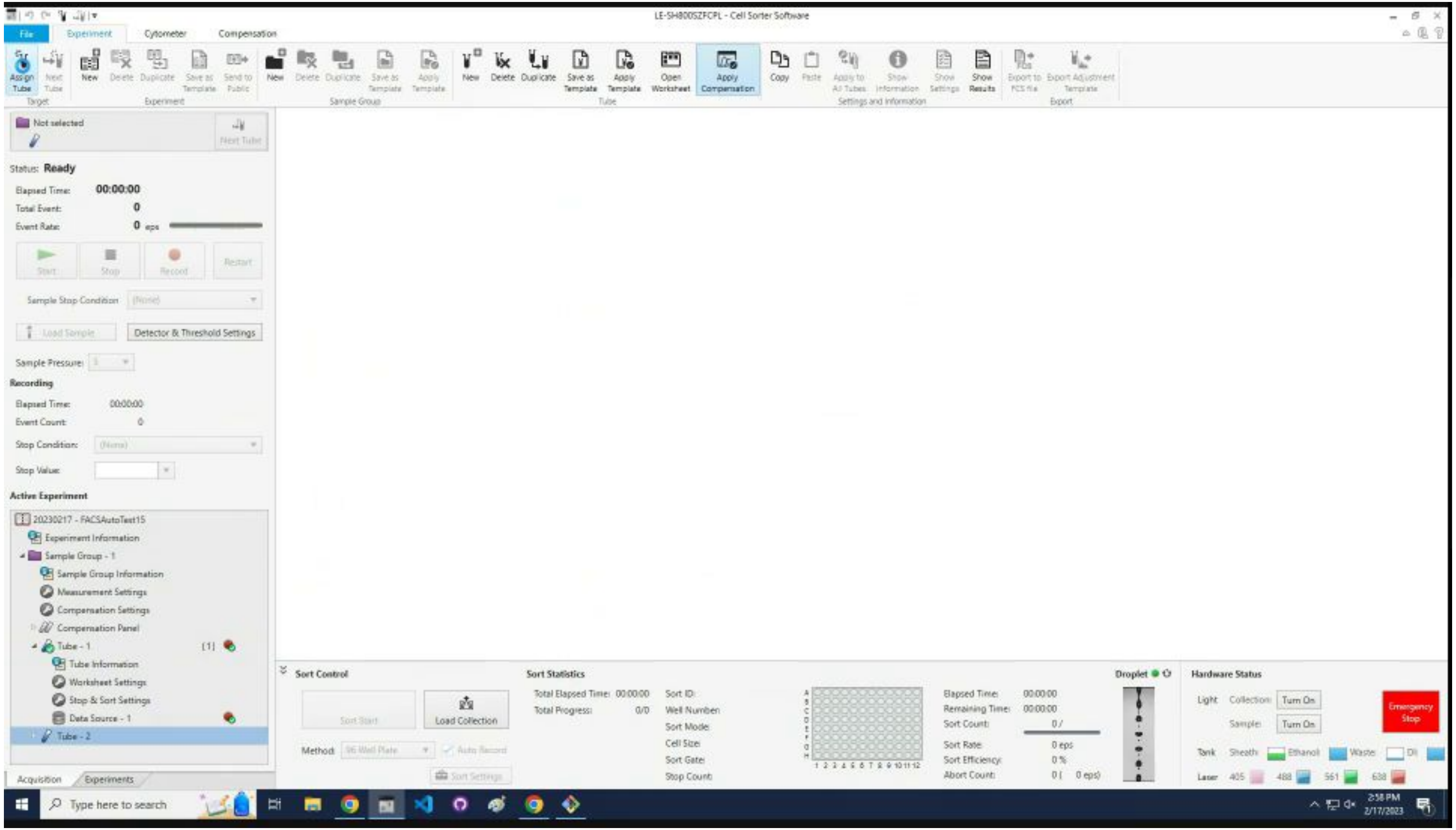

Goto [Iterate through samples](#) section.

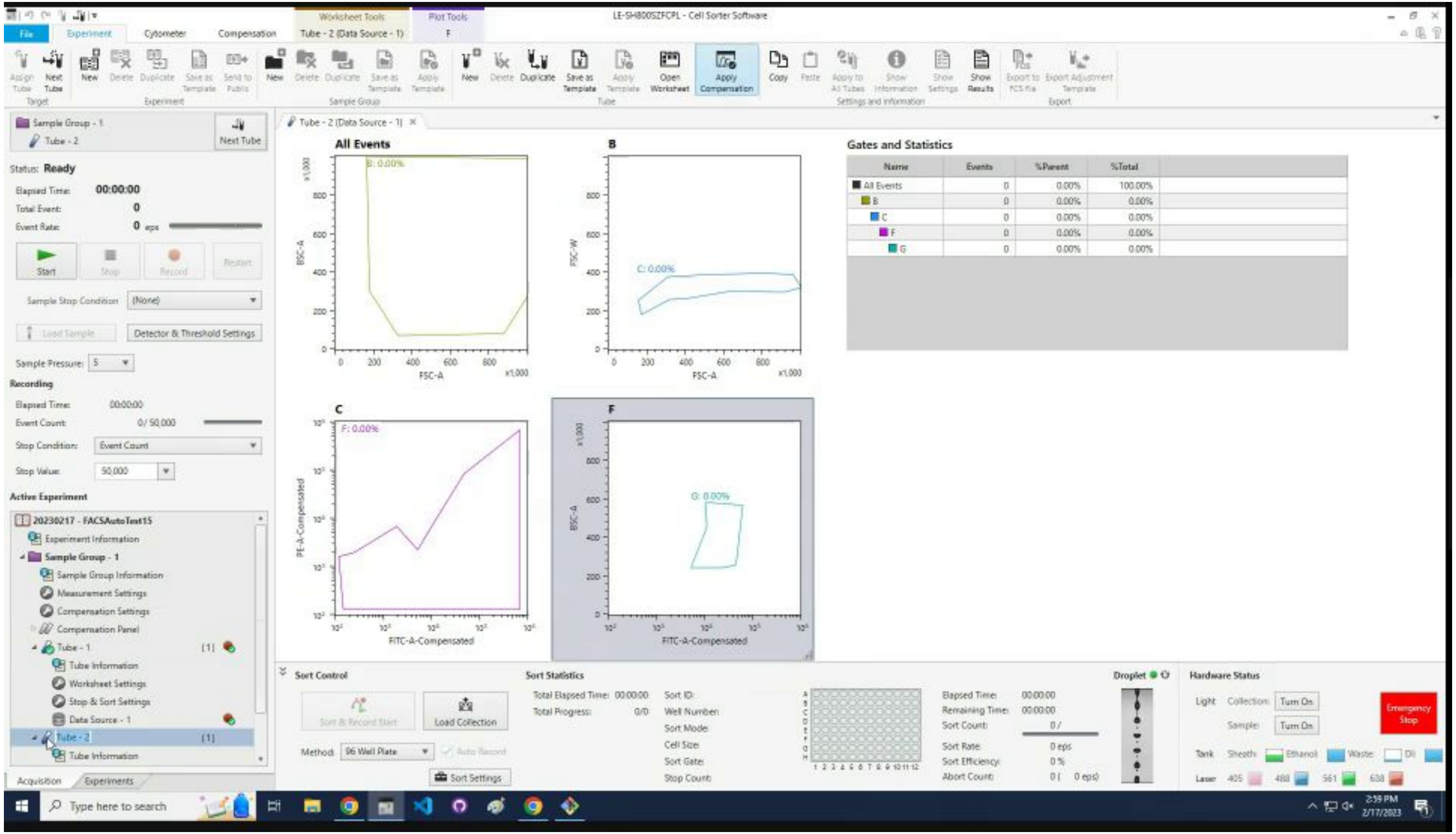

### Startup sequence

Continuation from previous set of samples

### Enter `n` to not set up the gates.

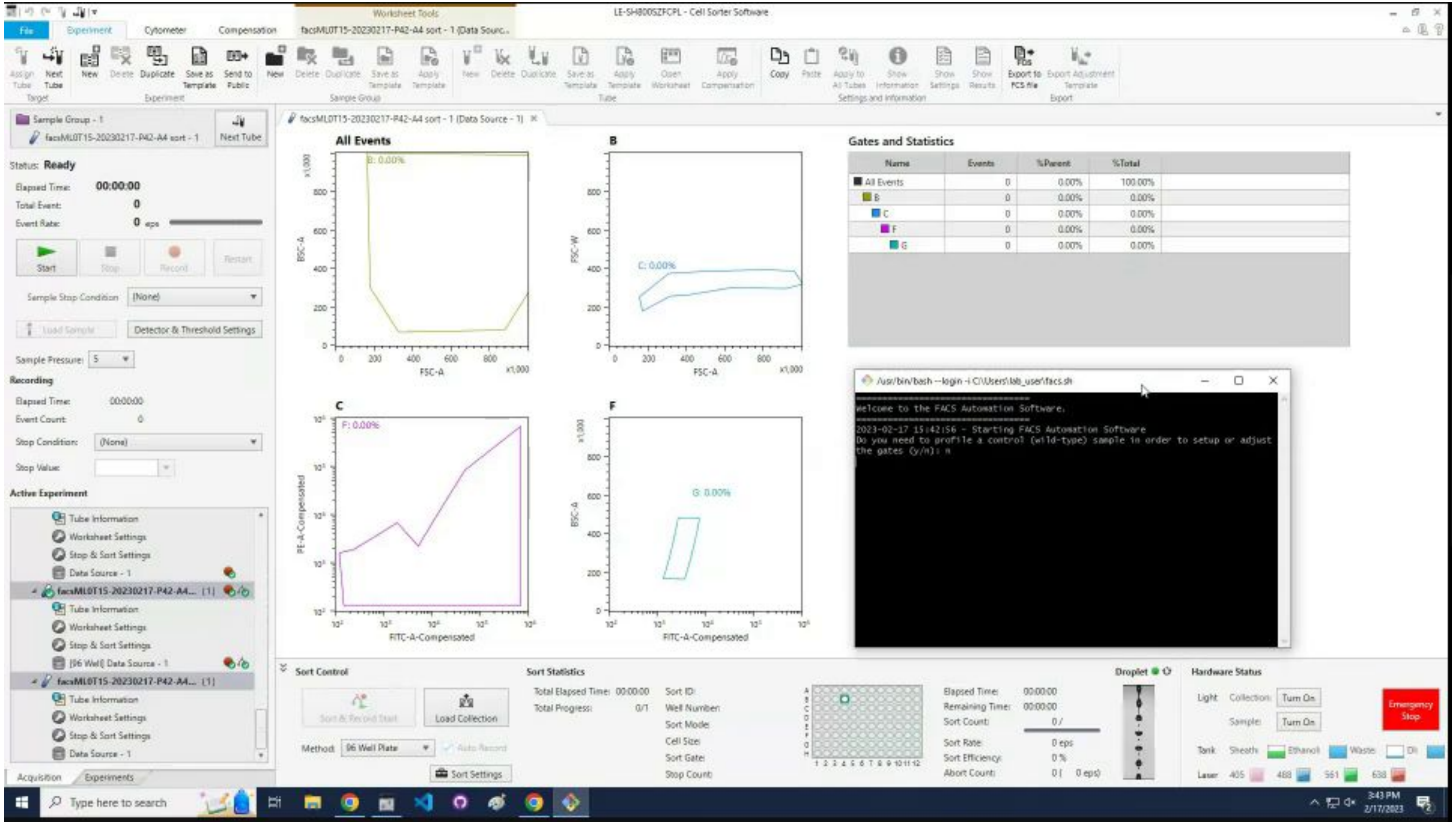

### Enter `user name.`

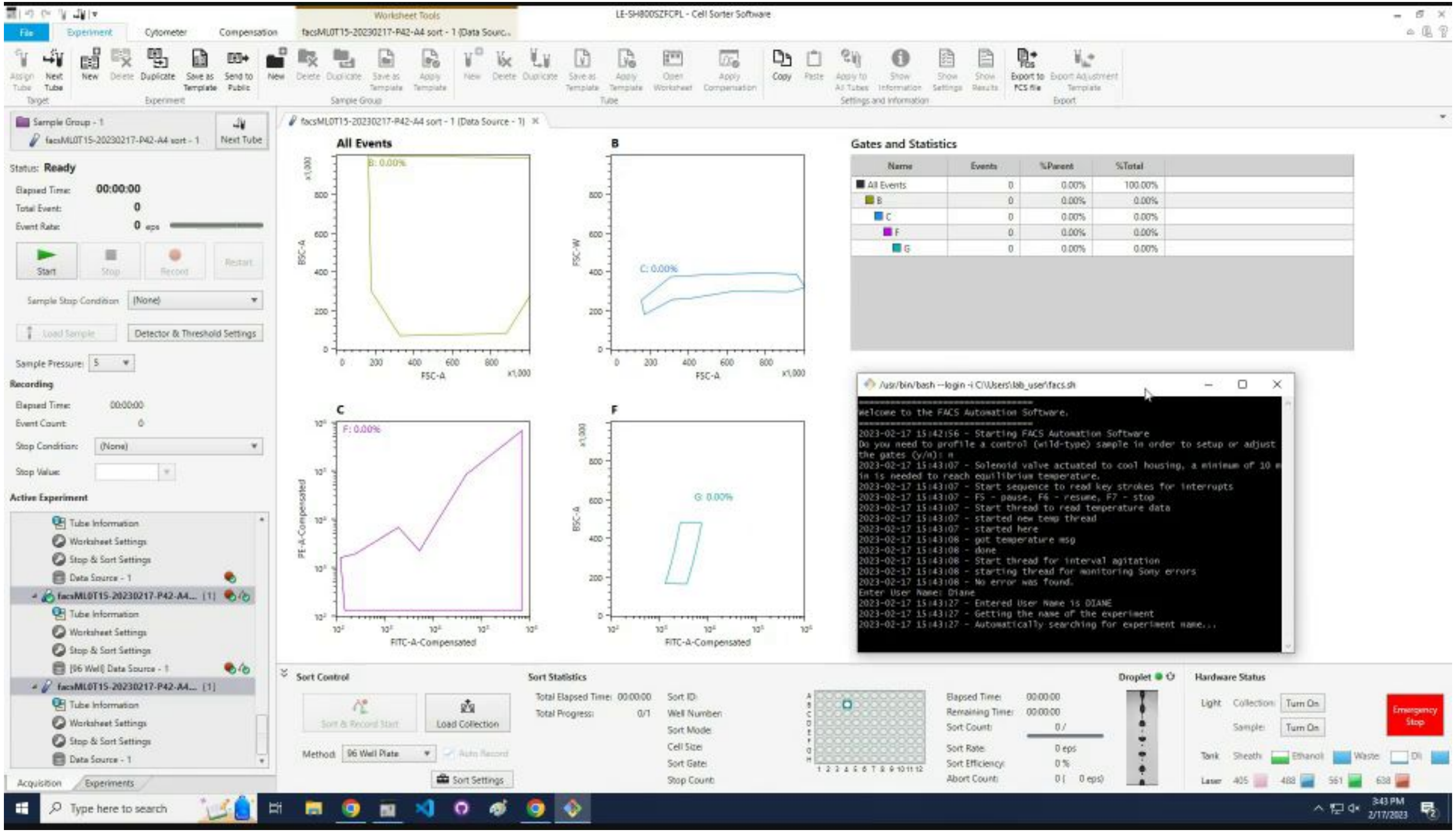

### Find experiment name to enter into program.

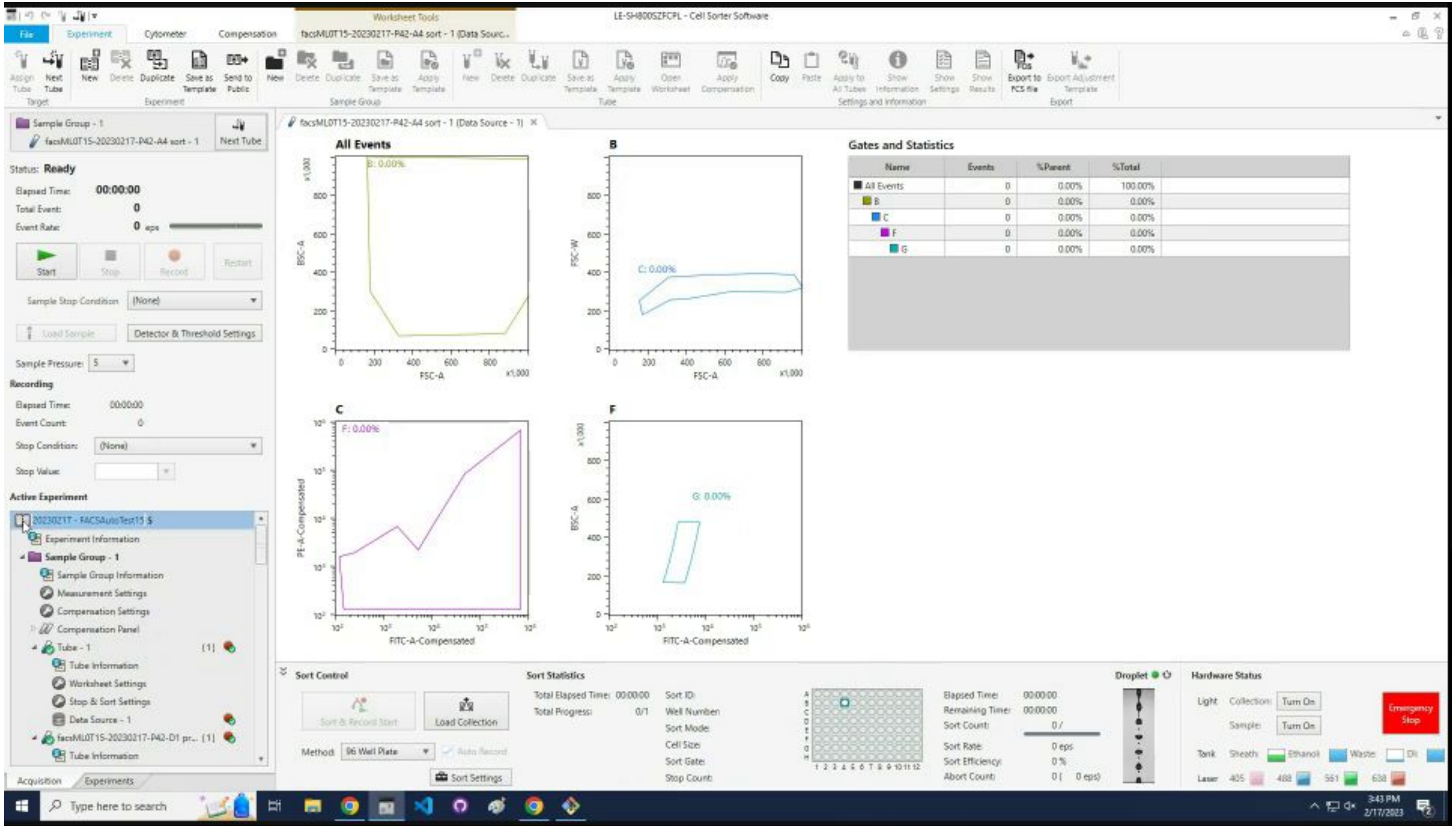

### Confirm that the experiment name is correct.

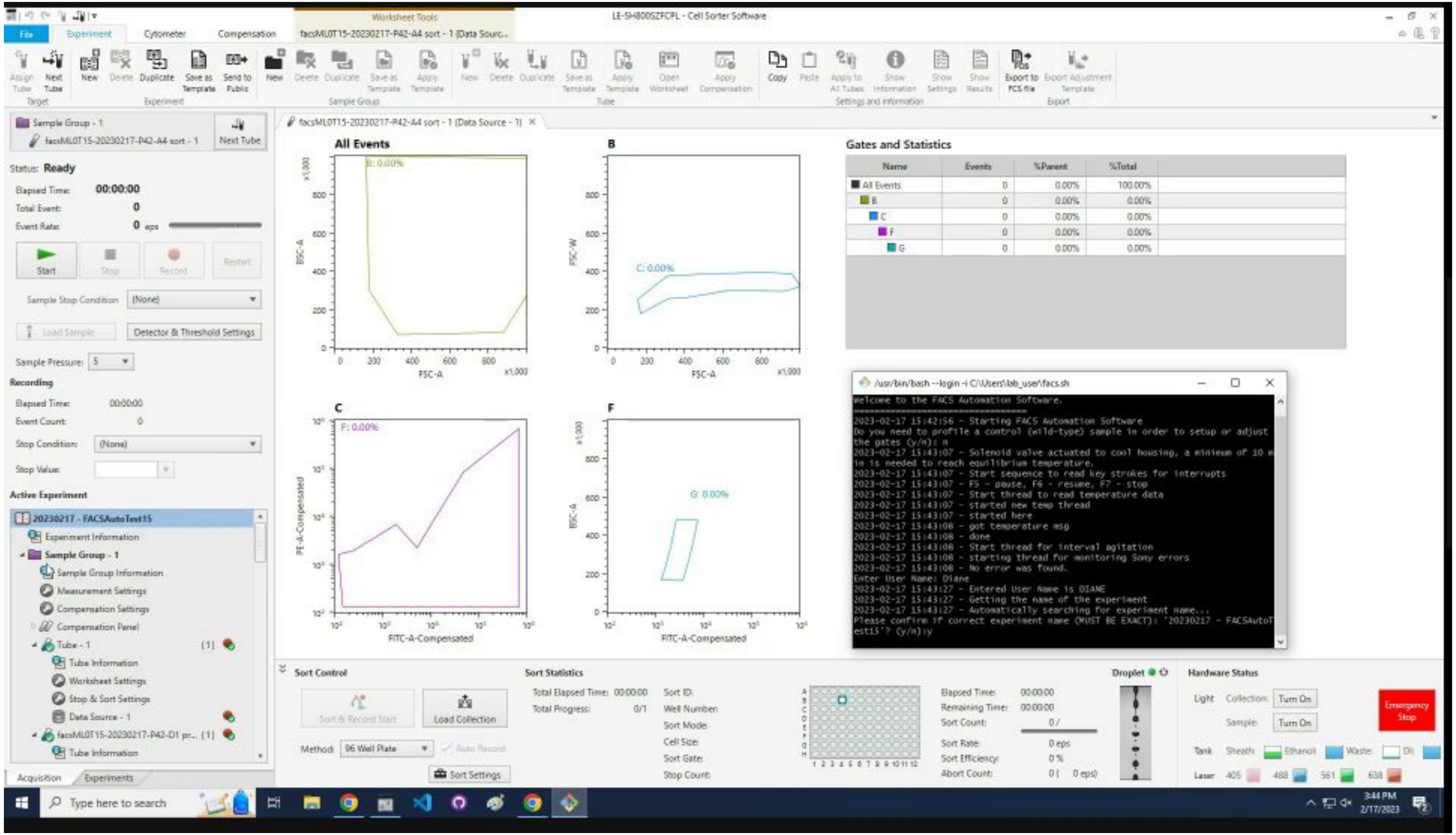

### Enter the first tube number to start the sorting process.

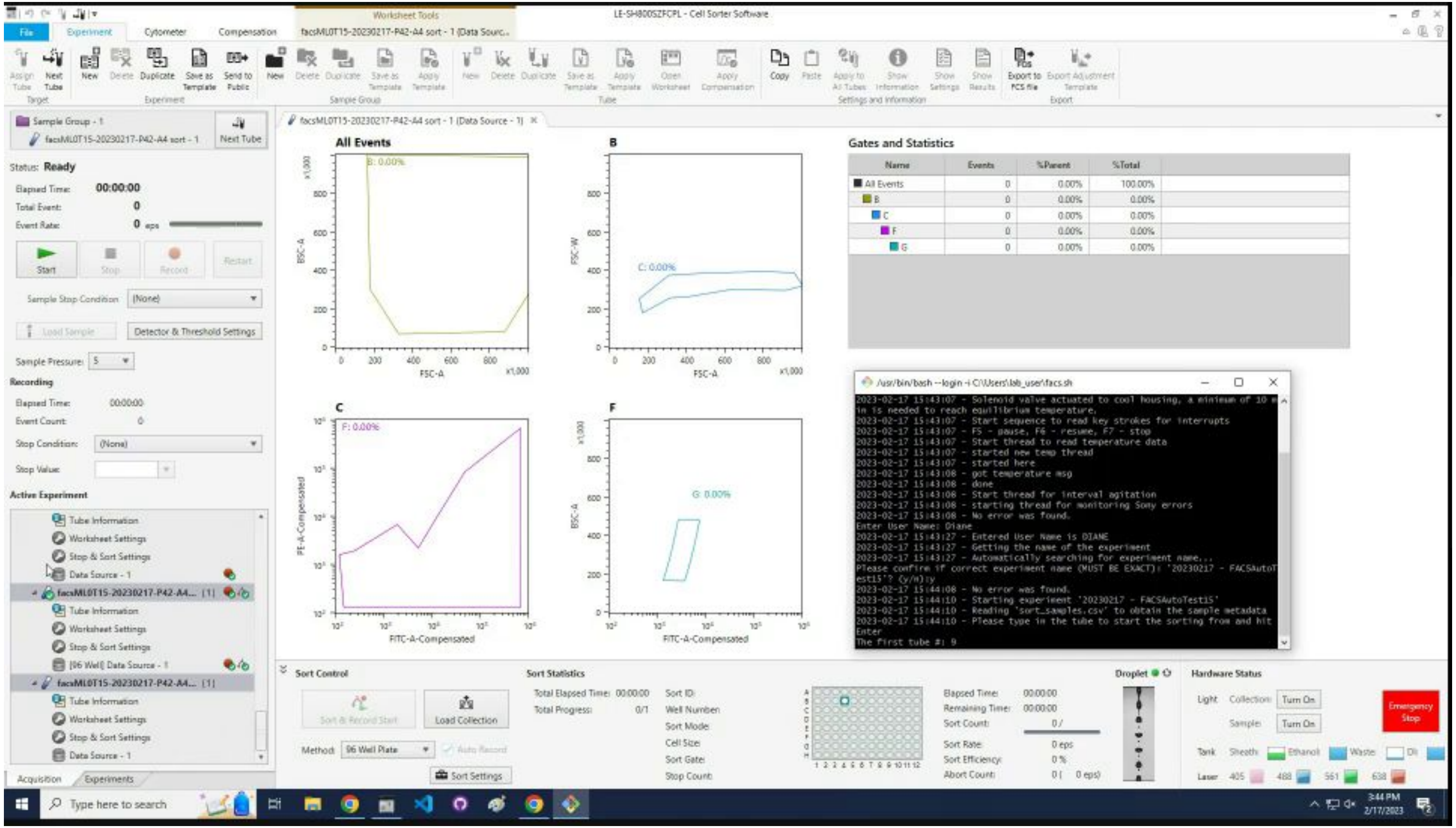

### Click `Sort Settings.`

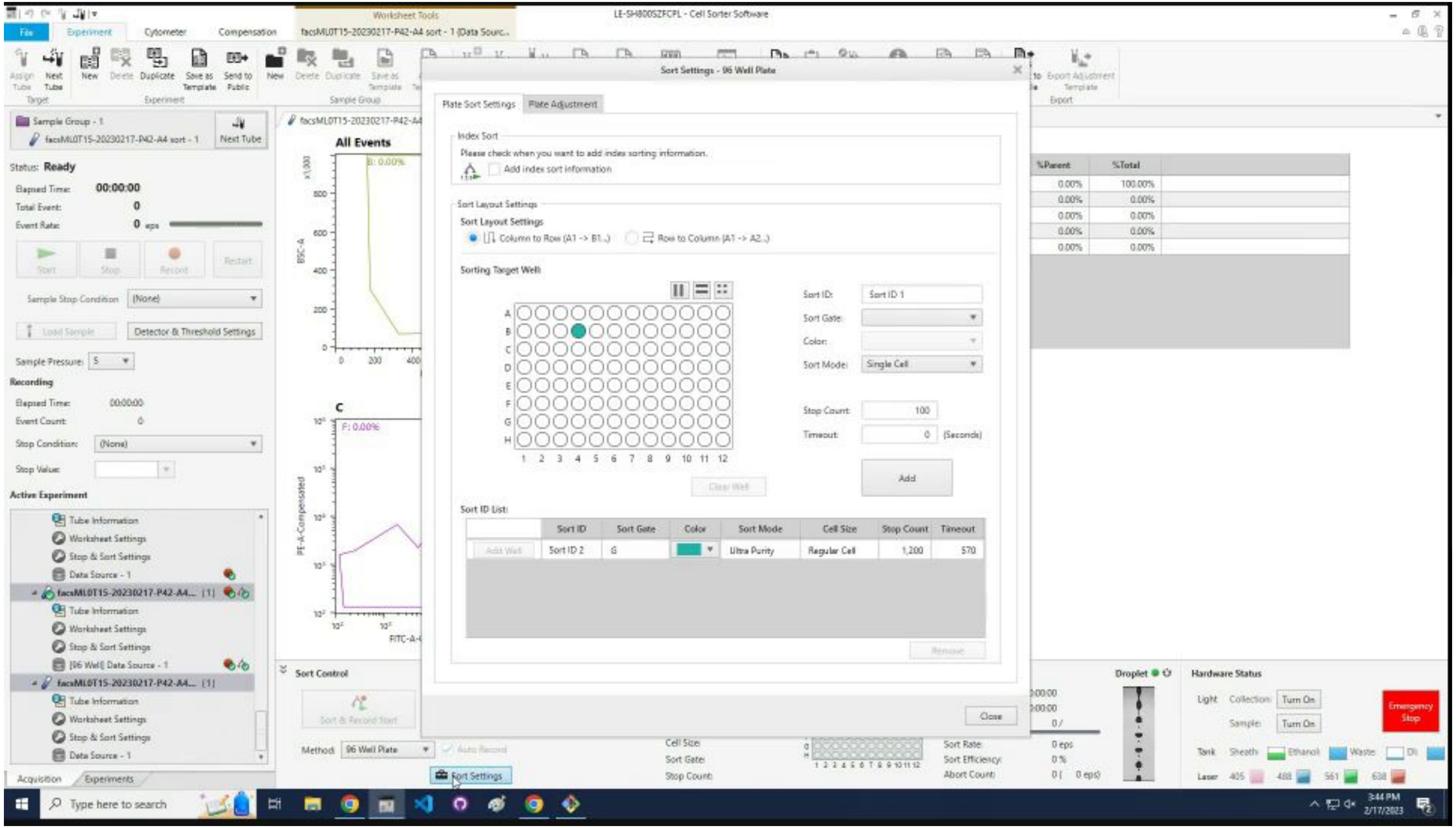

### Select the first line and click `Remove`.

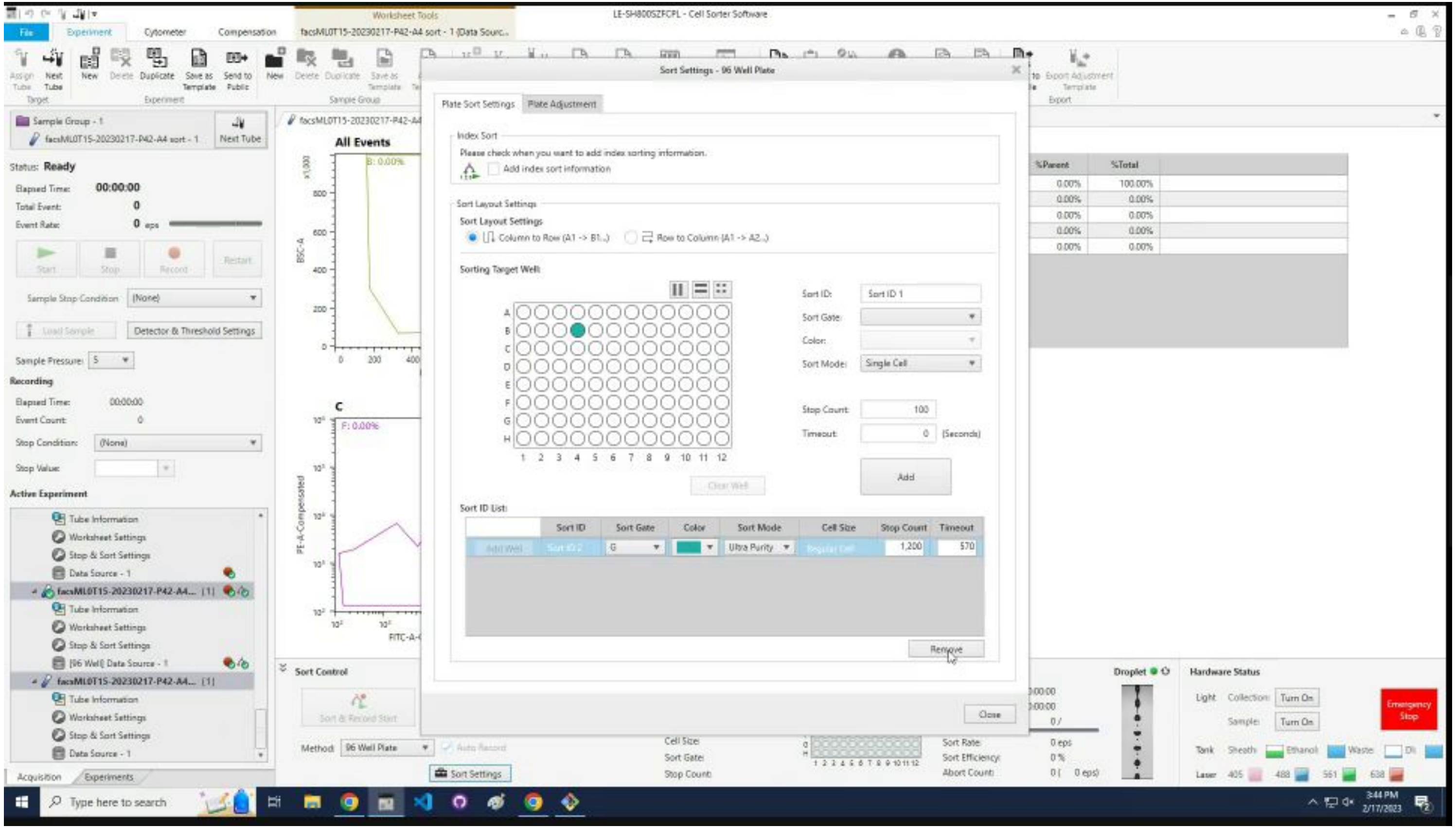

### Click `Close`.

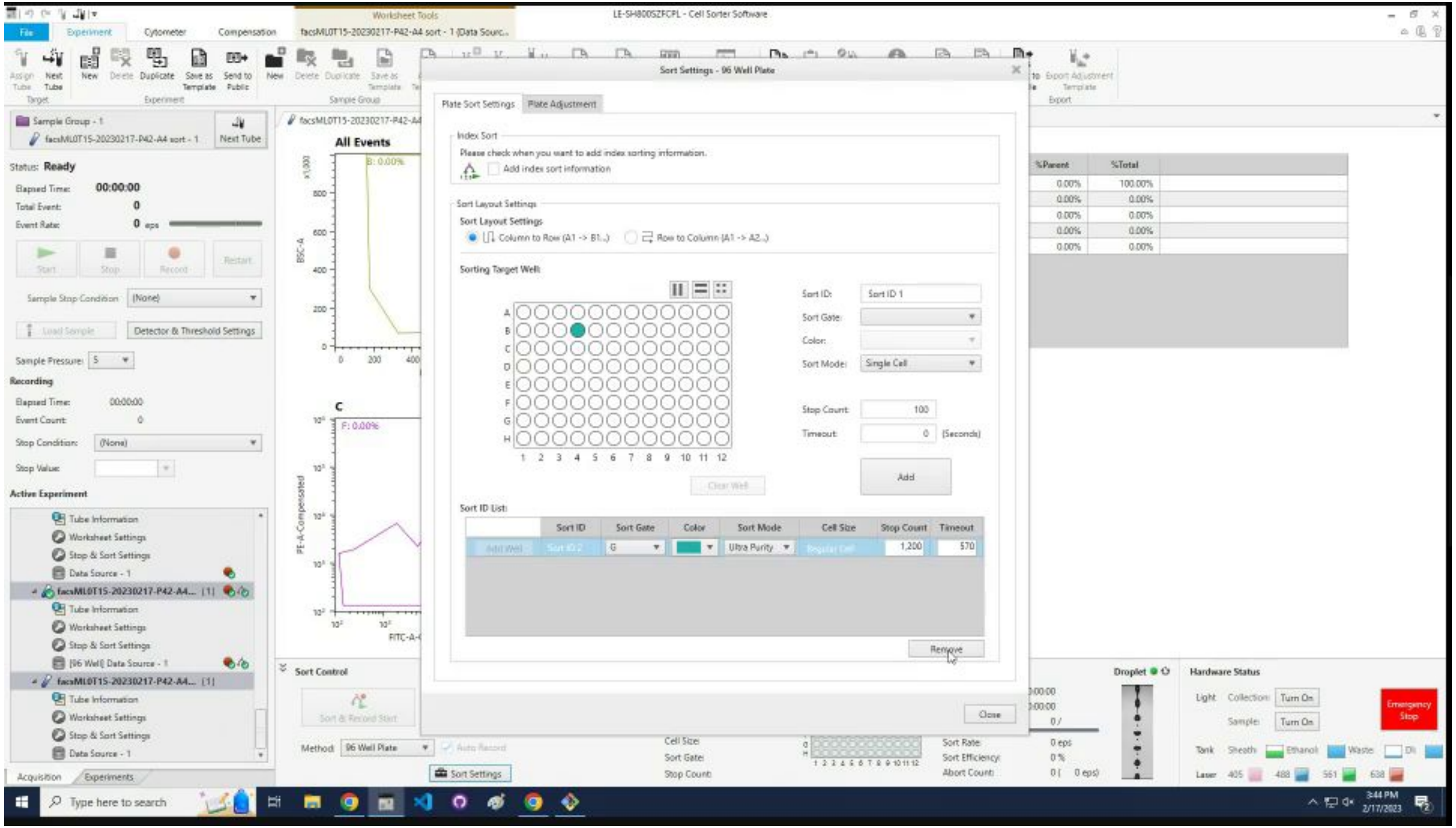
LE-SH800SZFCPL - Cell Sorter Software
Sort Settings - 96 Well Plate
Plate Sort Settings
Plate Adjustment
Index Sort
Please check when you want to add index sorting information.
Add index sort information
Sort Layout Settings
Column to Row (A1 -> B1..)
Row to Column (A1 -> A2..)
Sorting Target Well:
Sort ID:
Sort ID 1
Sort Gate:
Color:
Sort Mode:
Single Cell
Stop Count:
100
Timeout:
(Seconds)
Add
Clear Well
Sort ID List:
Sort ID
Sort Gate
Color
Sort Mode
Cell Size
Stop Count
Timeout
Ultra Purity
1,200
570
Remove
Close
All Events
Sample Group - 1
Next Tube
Status: Ready
Sort Control
Sort Settings
Hardware Status
Emergency Stop

### Click on the assigned tube in the list.

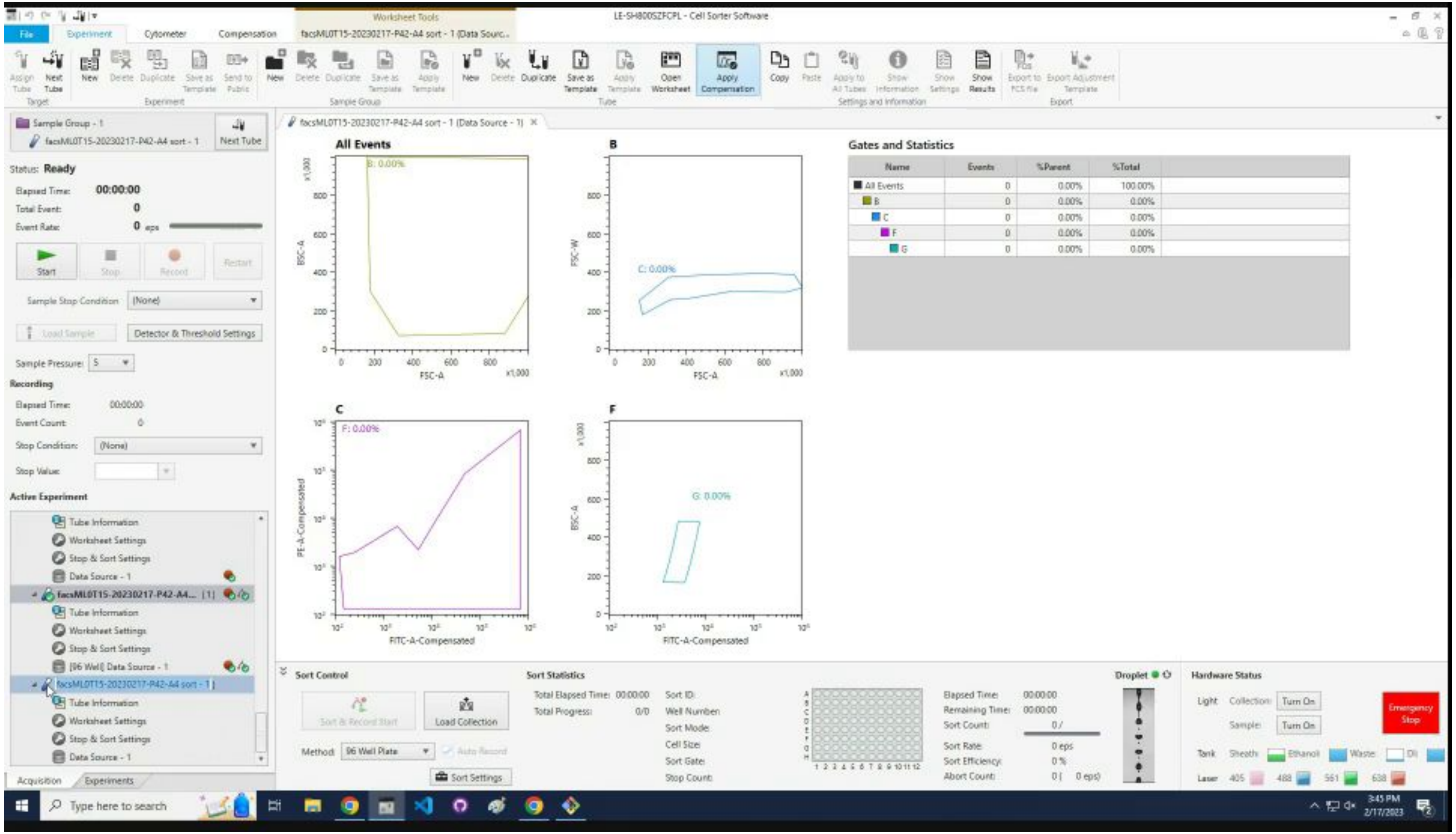

Goto `Iterate through samples` section.

### Iterate through samples

### Press `F2` to rename the active tube.

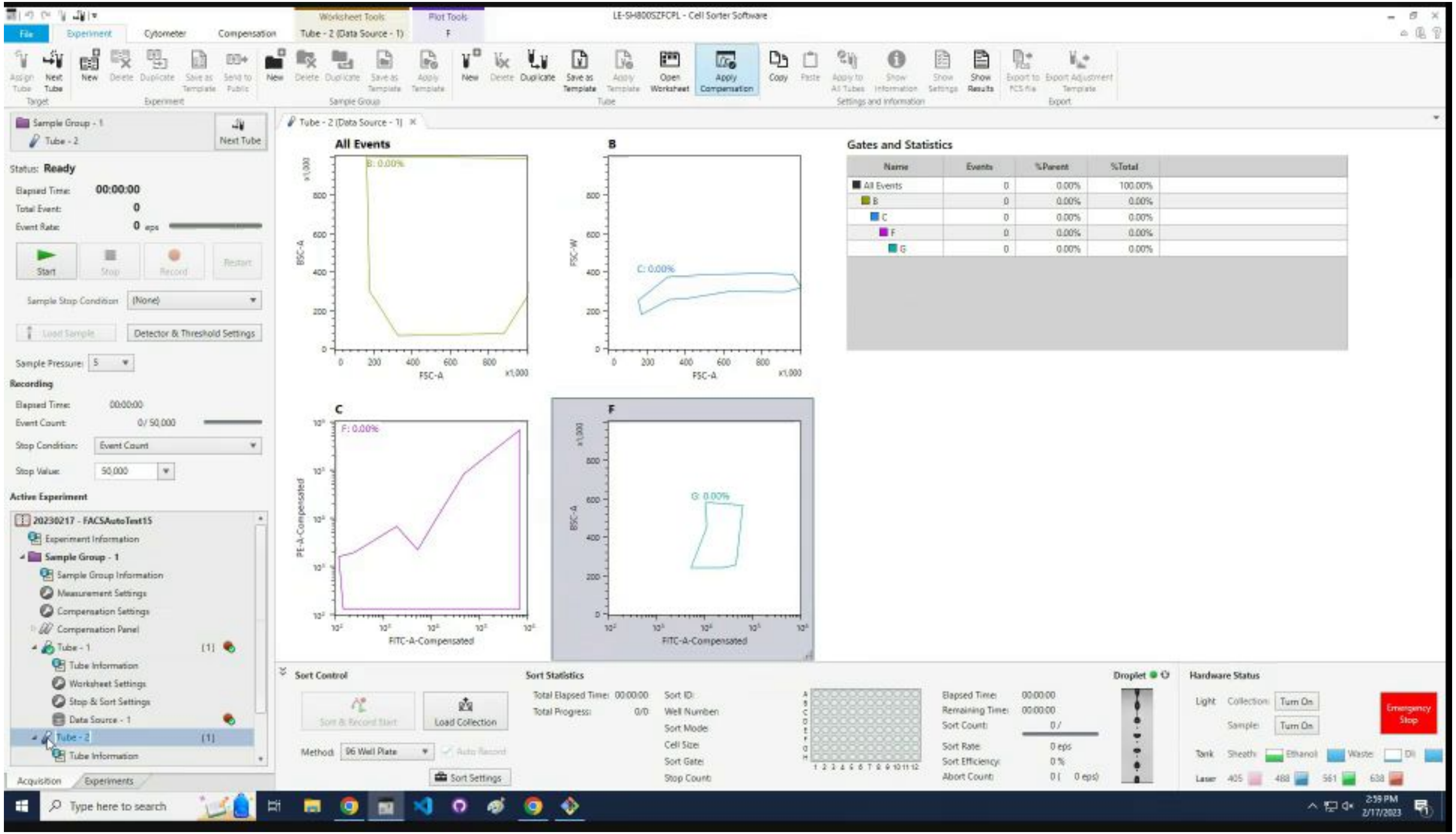

### Rename to the sample name appended with profile.

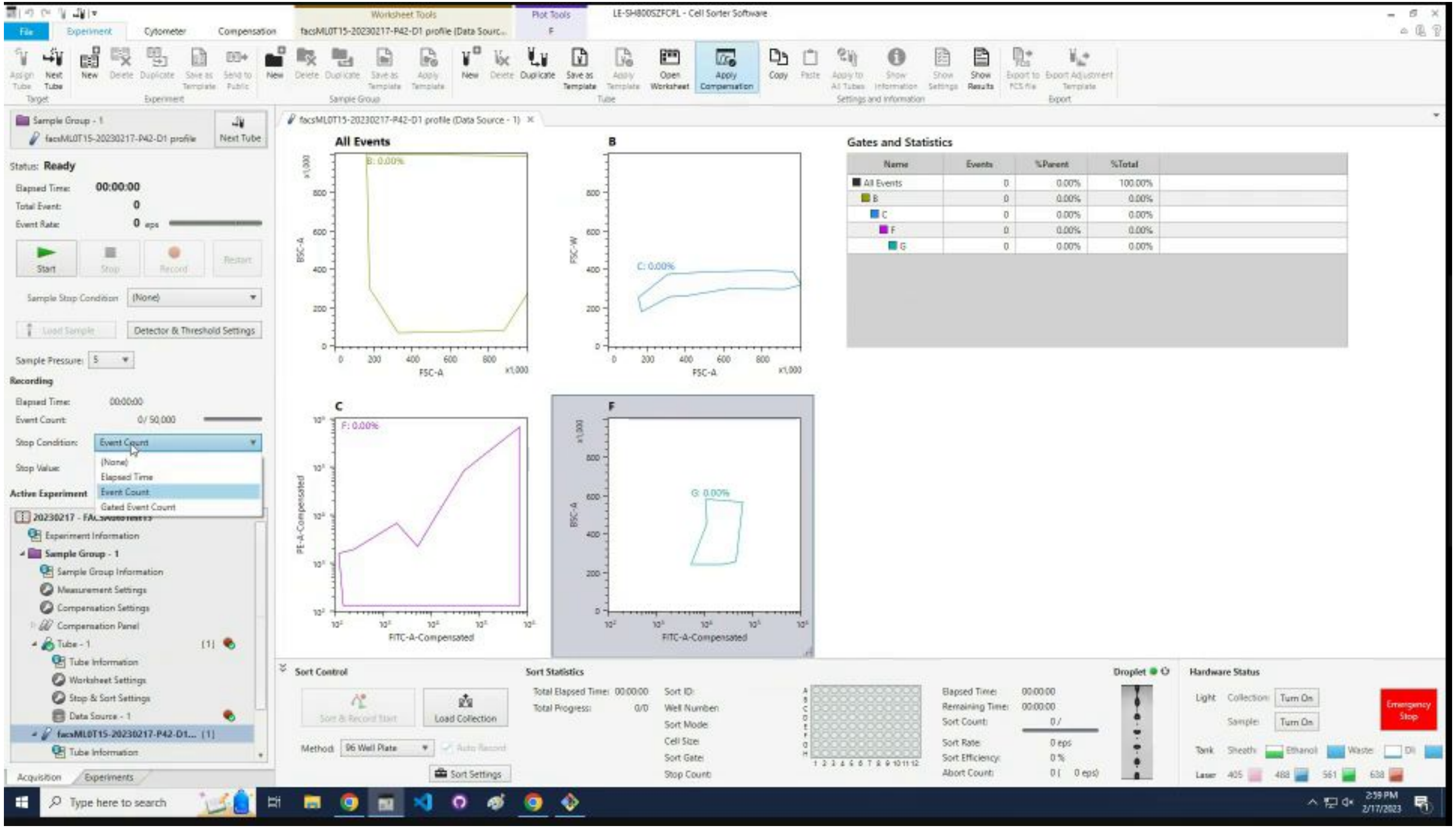

### Change the `Stop Condition` to `Event Count.`

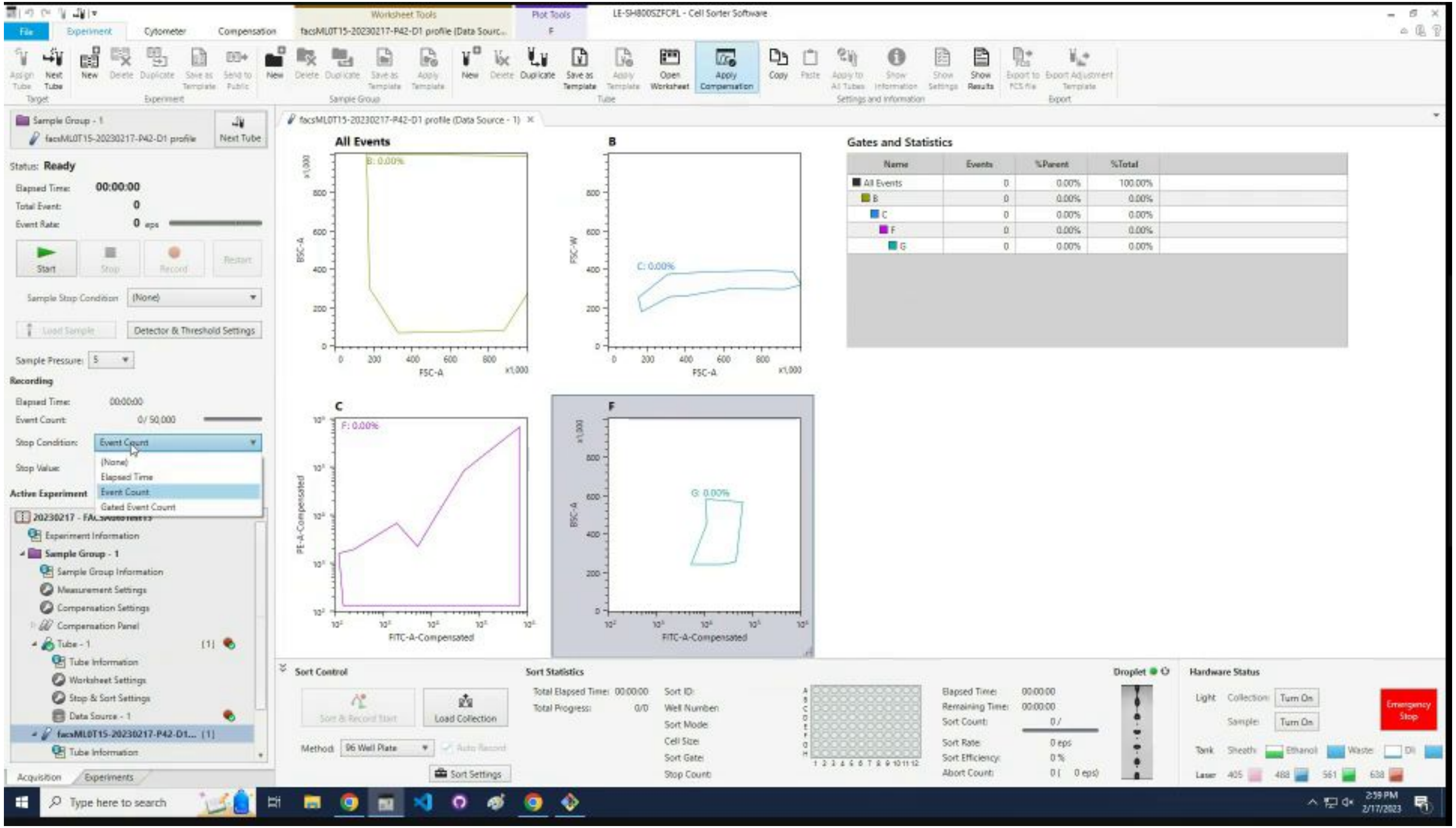

### Change the `Sample Pressure` to 5.

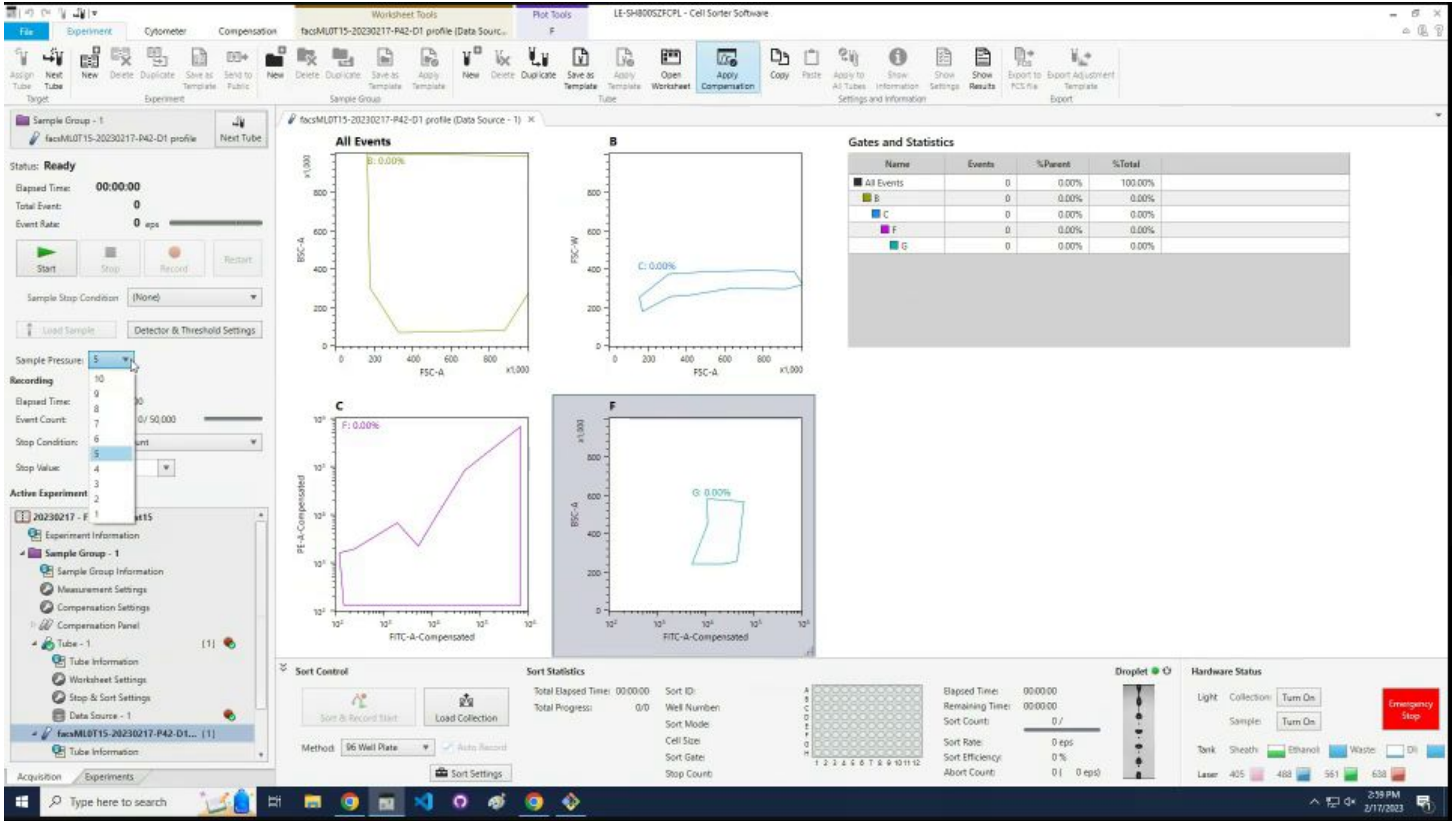

### Press `Start.`

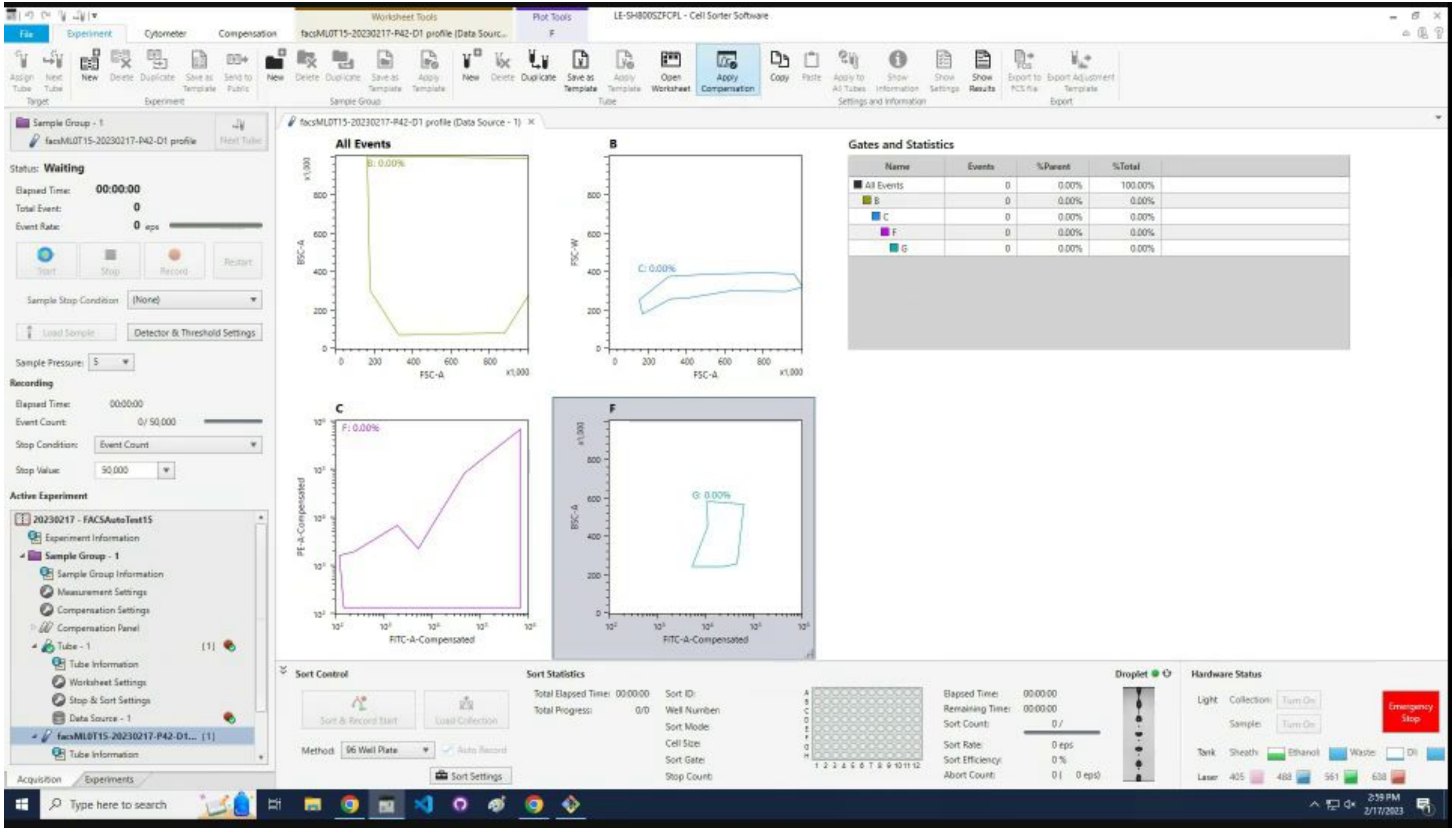
LE-SH800SZFCPL - Cell Sorter Software
File
Experiment
Cytometer
Compensation
Sample Group - 1
Status: Waiting
Elapsed Time: 00:00:00
Total Event: 0
Event Rate: 0 eps
Start
Stop
Record
Restart
Sample Stop Condition (None)
Load Sample
Detector & Threshold Settings
Sample Pressure: 5
Recording
Elapsed Time: 00:00:00
Event Count: 0/ 50,000
Stop Condition: Event Count
Stop Value: 50,000
Active Experiment
20230217 - FACSAutoTest15
Experiment Information
Sample Group - 1
Sample Group Information
Measurement Settings
Compensation Settings
Compensation Panel
Tube - 1
Tube Information
Worksheet Settings
Stop & Sort Settings
Data Source - 1
All Events
B
C
F
FSC-A
BSC-A
FSC-W
PE-A-Compensated
FITC-A-Compensated
Gates and Statistics
Name
Events
%Parent
%Total
All Events 0 0.00% 100.00%
B 0 0.00% 0.00%
C 0 0.00% 0.00%
F 0 0.00% 0.00%
G 0 0.00% 0.00%
Sort Control
Sort & Record Start
Load Collection
Method: 96 Well Plate
Auto Record
Sort Settings
Sort Statistics
Total Elapsed Time: 00:00:00
Total Progress: 0/0
Sort ID:
Well Number:
Sort Mode:
Cell Size:
Sort Gate:
Stop Count:
Elapsed Time: 00:00:00
Remaining Time: 00:00:00
Sort Count: 0 /
Sort Rate: 0 eps
Sort Efficiency: 0 %
Abort Count: 0 ( 0 eps)
Droplet
Hardware Status
Light
Collection:
Sample:
Turn On
Tank
Sheath
Ethanol
Waste
DI
Laser
405
488
561
638
Emergency Stop
Acquisition
Experiments
Type here to search
2:59 PM
2/17/2023

### Wait ~20 seconds for cells to appear. Click `Record`.

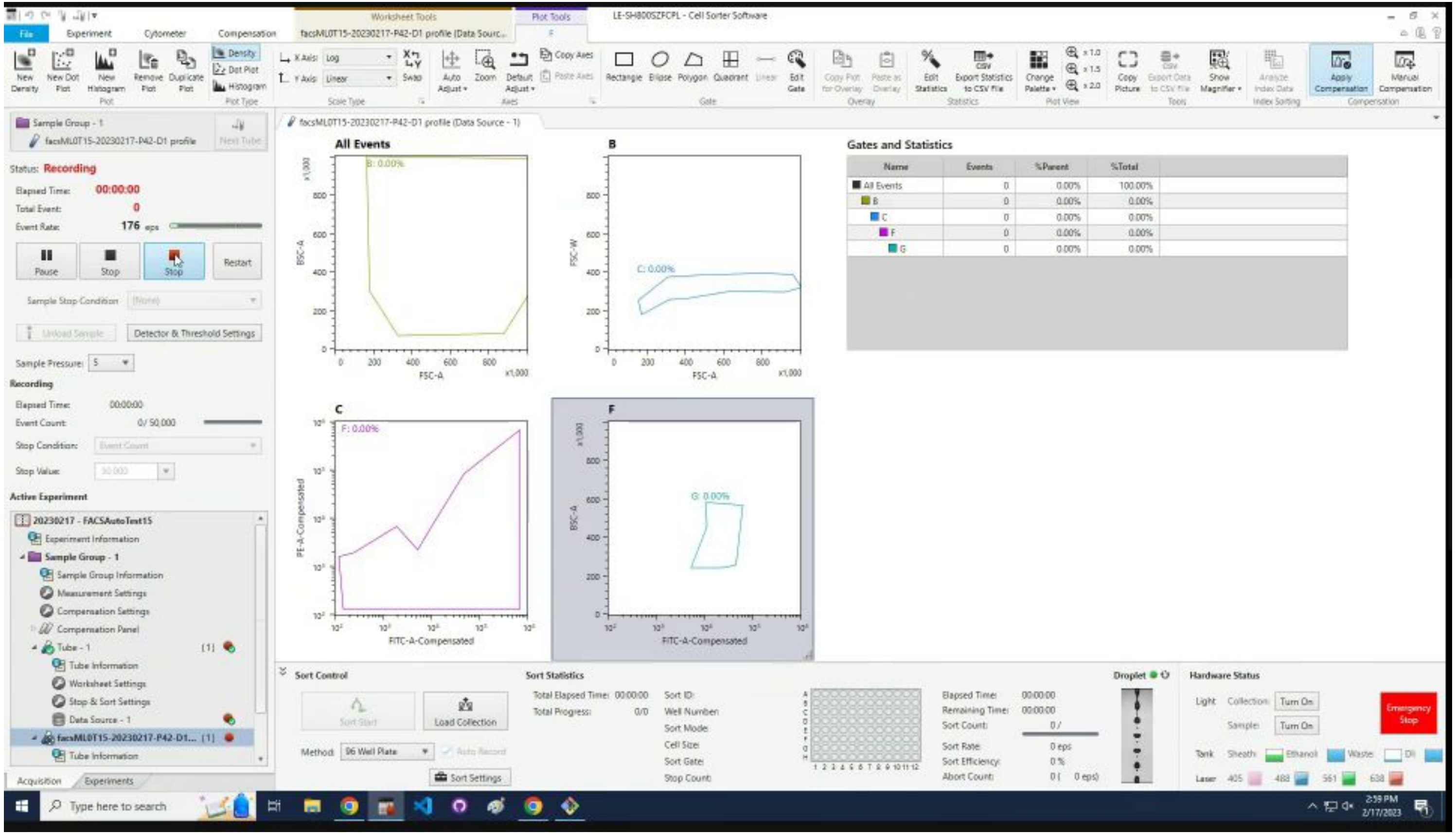

### Wait for profiling to finish then click `OK`.

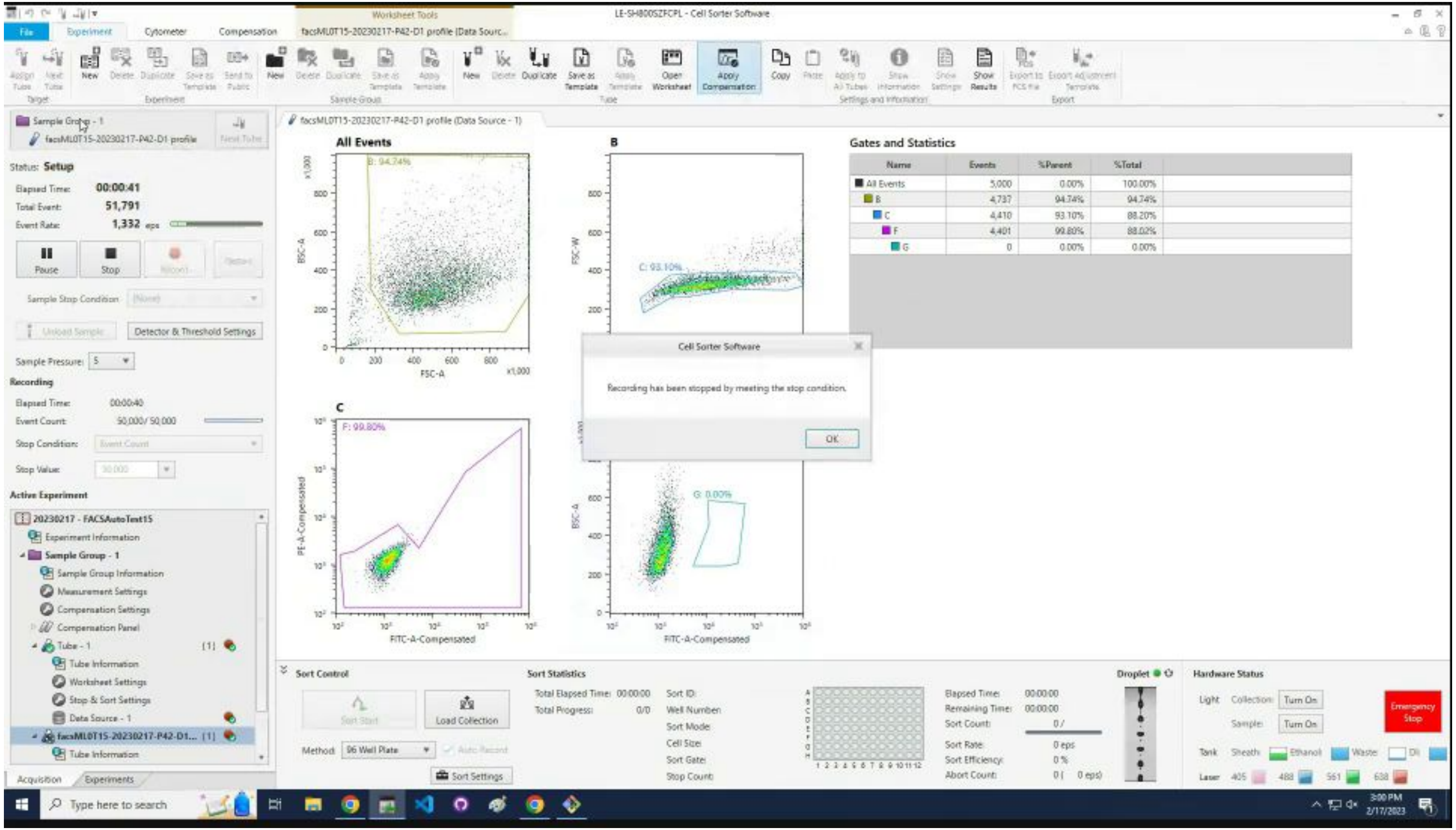

### Click `Stop`.

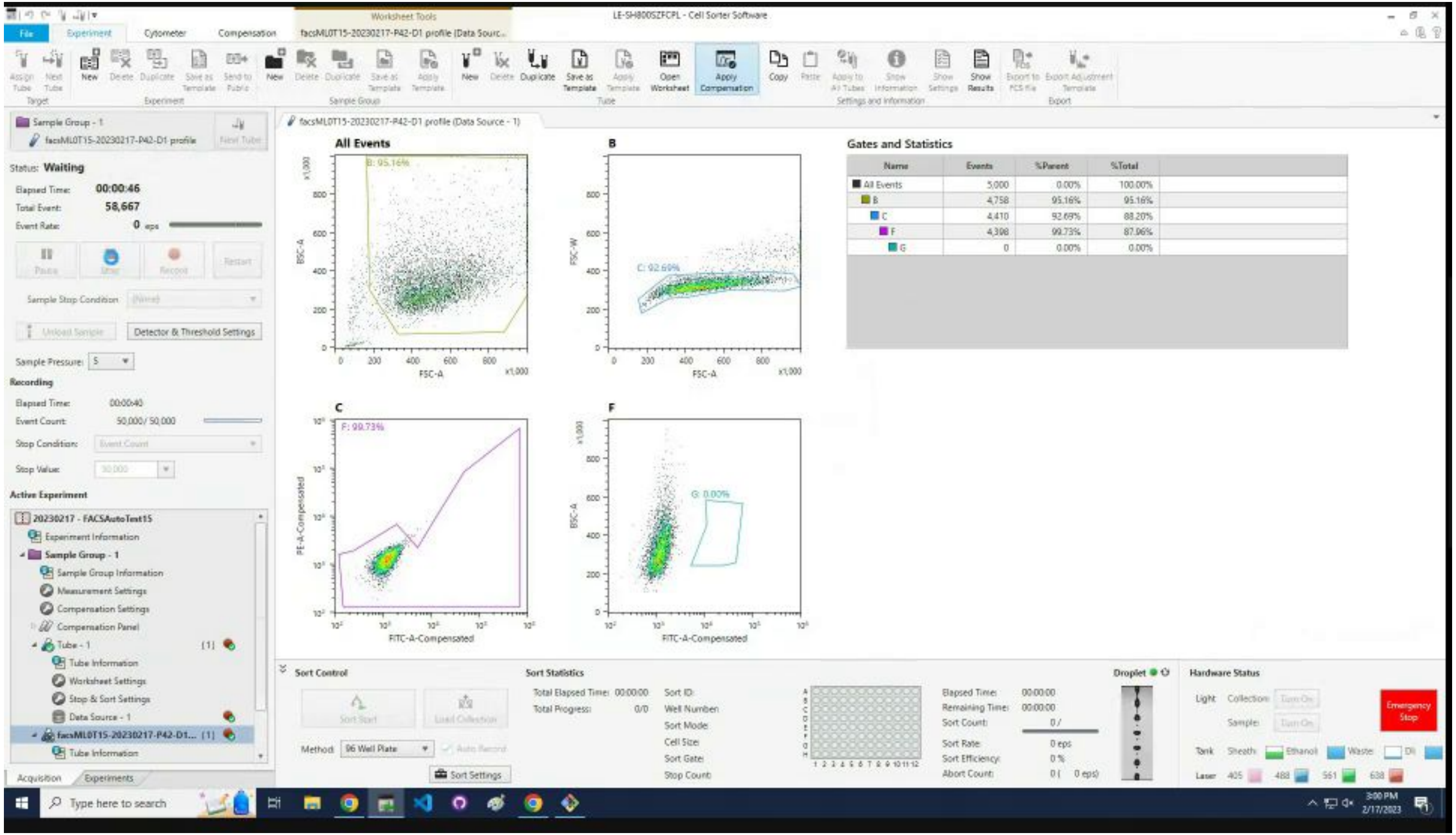

LE-SH800SZFCPL - Cell Sorter Software
Status: Waiting
Elapsed Time: 00:00:46
Total Event: 58,667
Event Rate: 0 eps
Pause
Stop
Record
Restart
Sample Stop Condition
Detector & Threshold Settings
Sample Pressure: 5
Recording
Elapsed Time: 00:00:40
Event Count: 50,000 / 50,000
Stop Condition:
Stop Value:
Active Experiment
20230217 - FACSAutoTest15
All Events
B: 95.16%
C: 92.69%
F: 99.73%
G: 0.00%
FSC-A
FITC-A-Compensated
Gates and Statistics
Name | Events | %Parent | %Total
All Events | 5,000 | 0.00% | 100.00%
B | 4,758 | 95.16% | 95.16%
C | 4,410 | 92.69% | 88.20%
F | 4,398 | 99.73% | 87.96%
G | 0 | 0.00% | 0.00%
Sort Control
Sort Statistics
Hardware Status
Emergency Stop

### Click on the fourth plot.

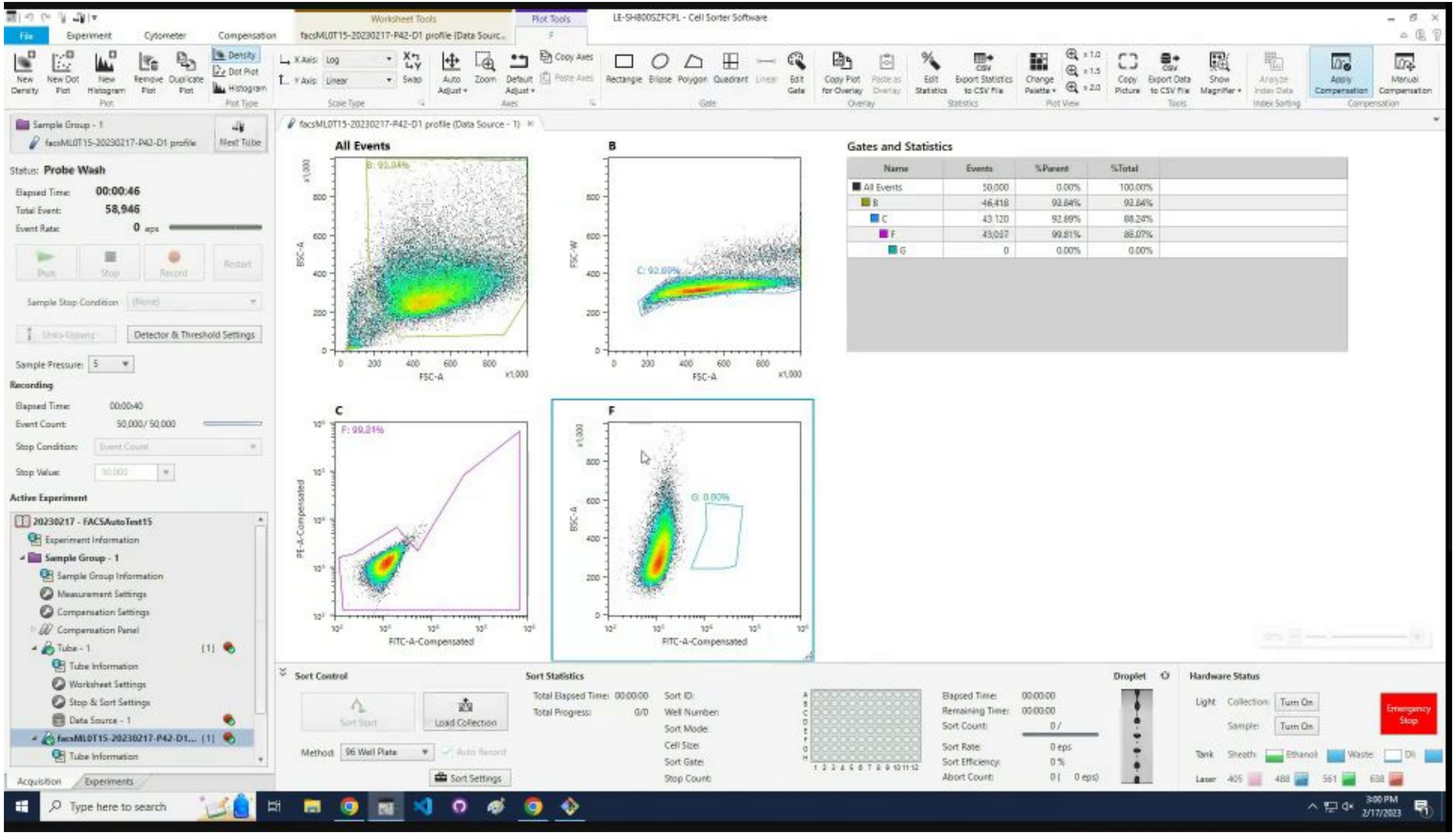

Click `Export Data to CSV File` icon.

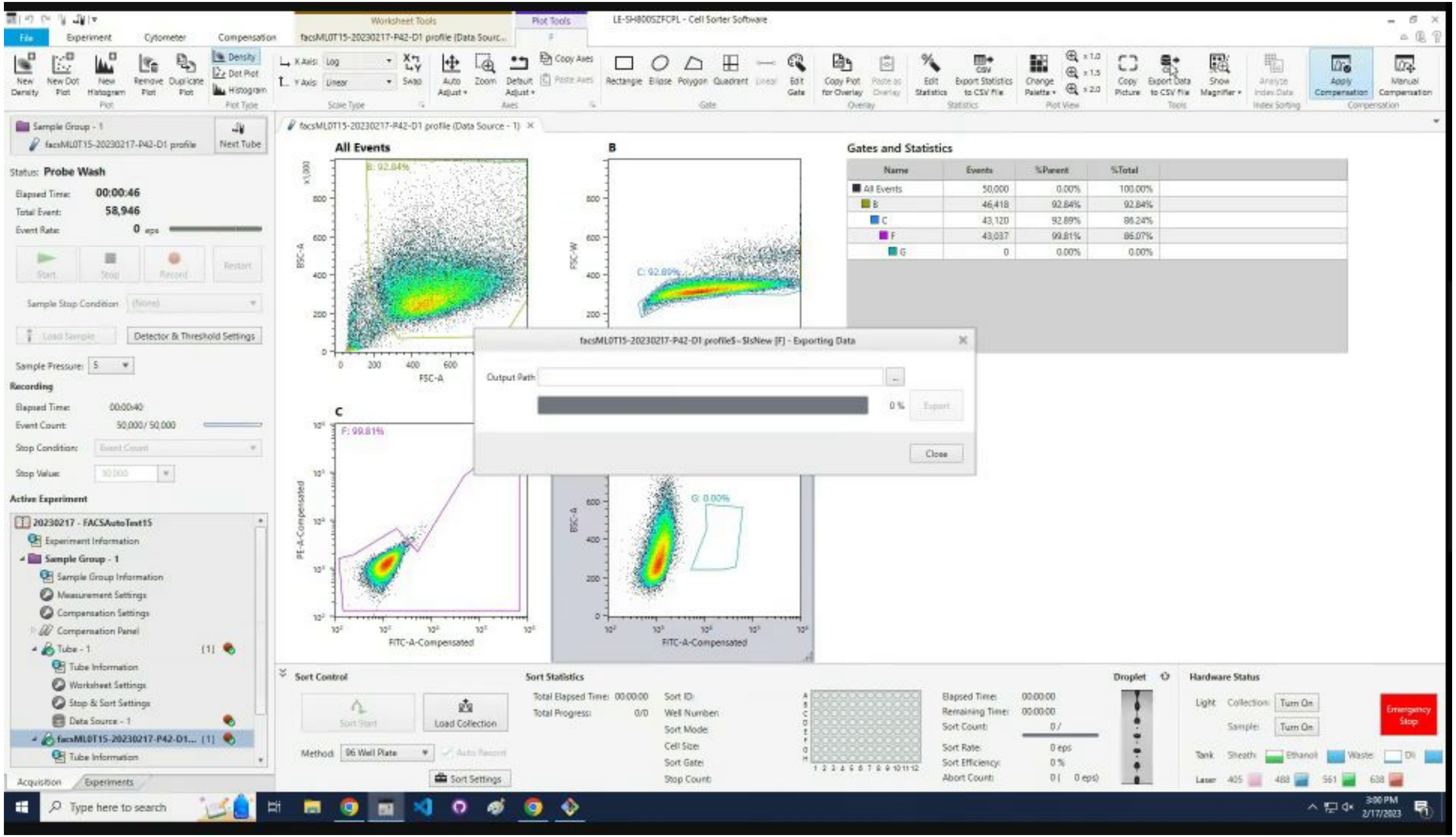

### Click the … button in the popup.

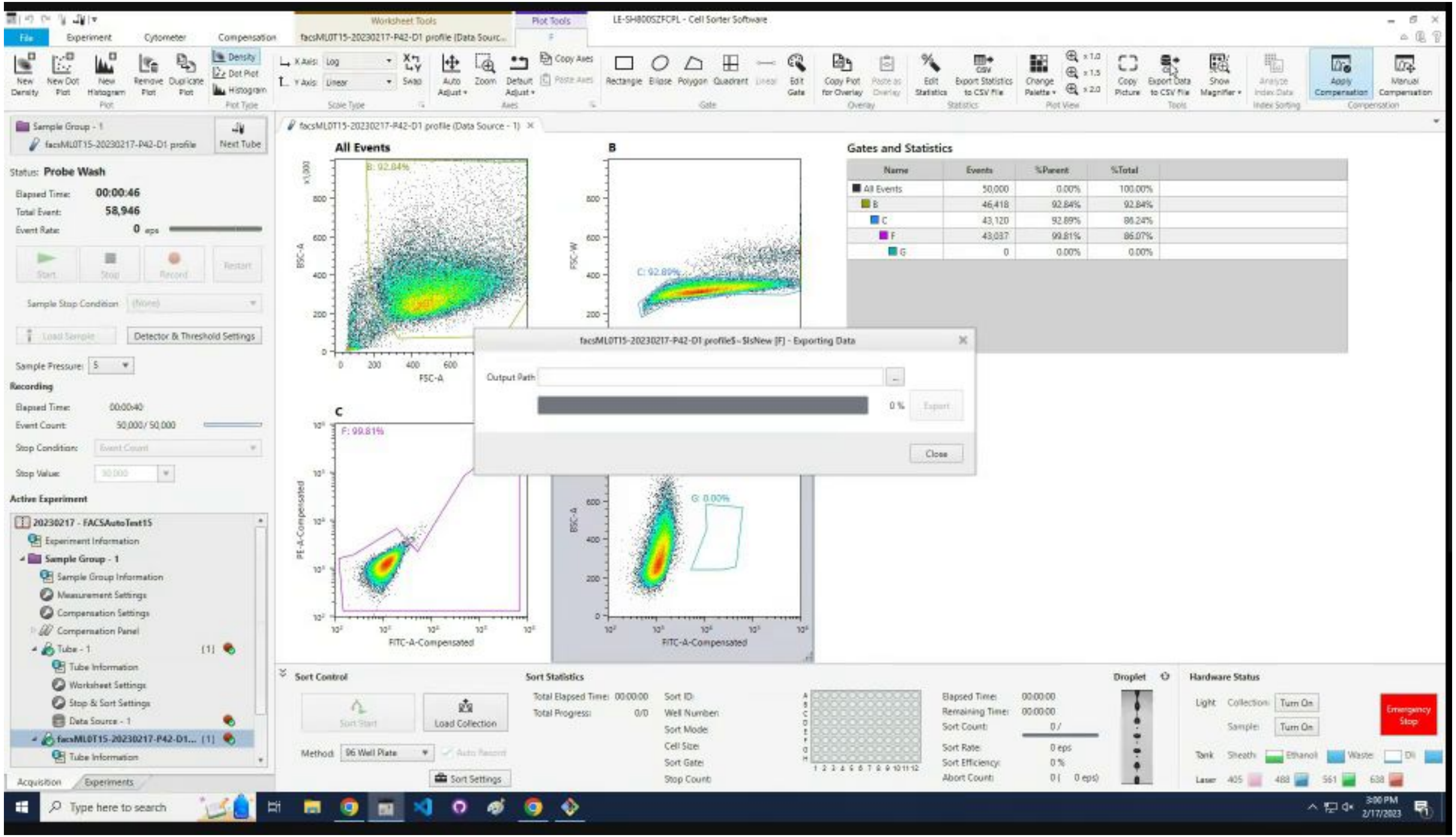

### Select all of the file path and file name.

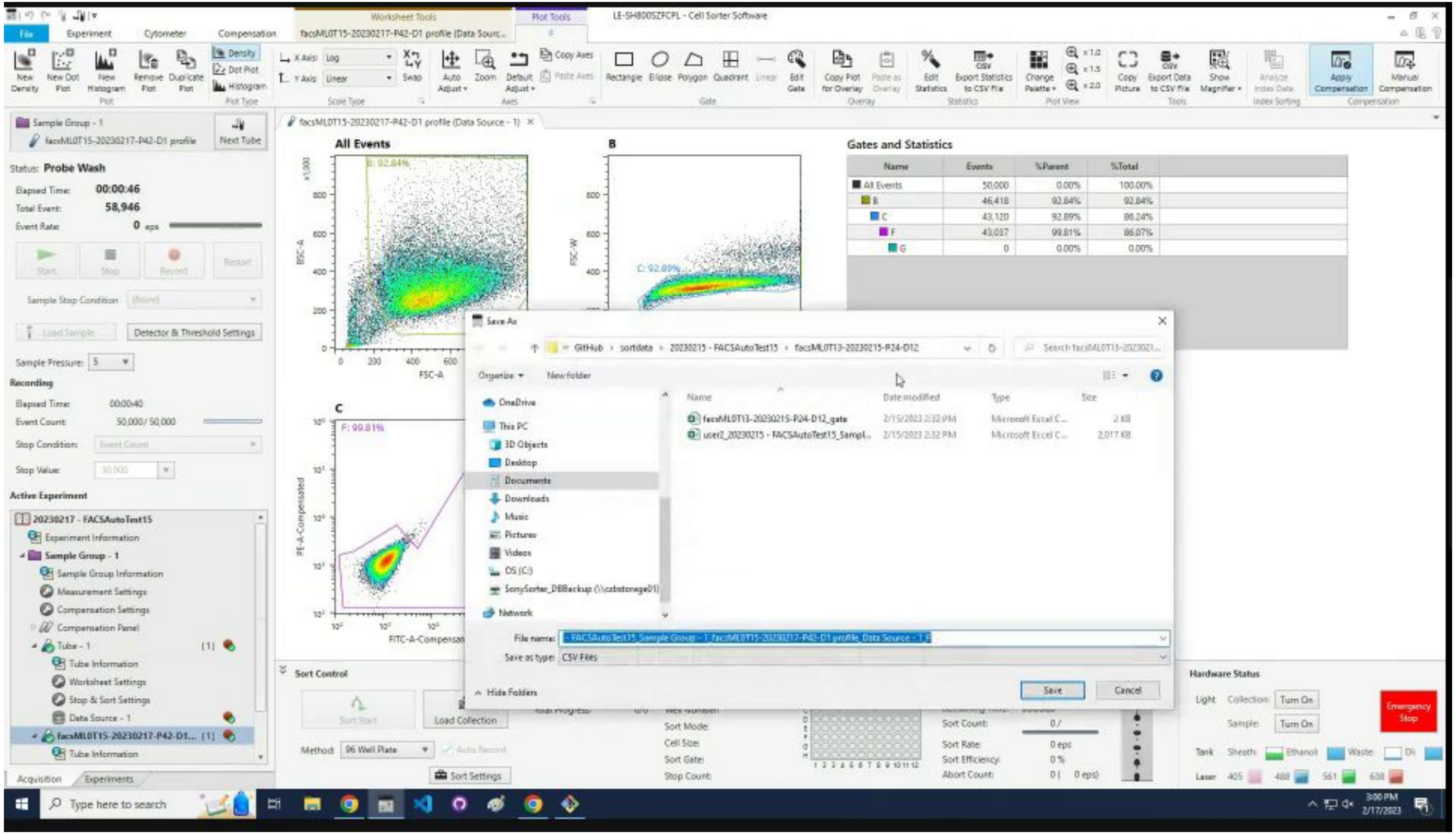

### Rename the file as

`C:\Users\lab_user\Desktop\FACSAuto\{User}_{Experiment}_{SampleGroup}_{Sample}`

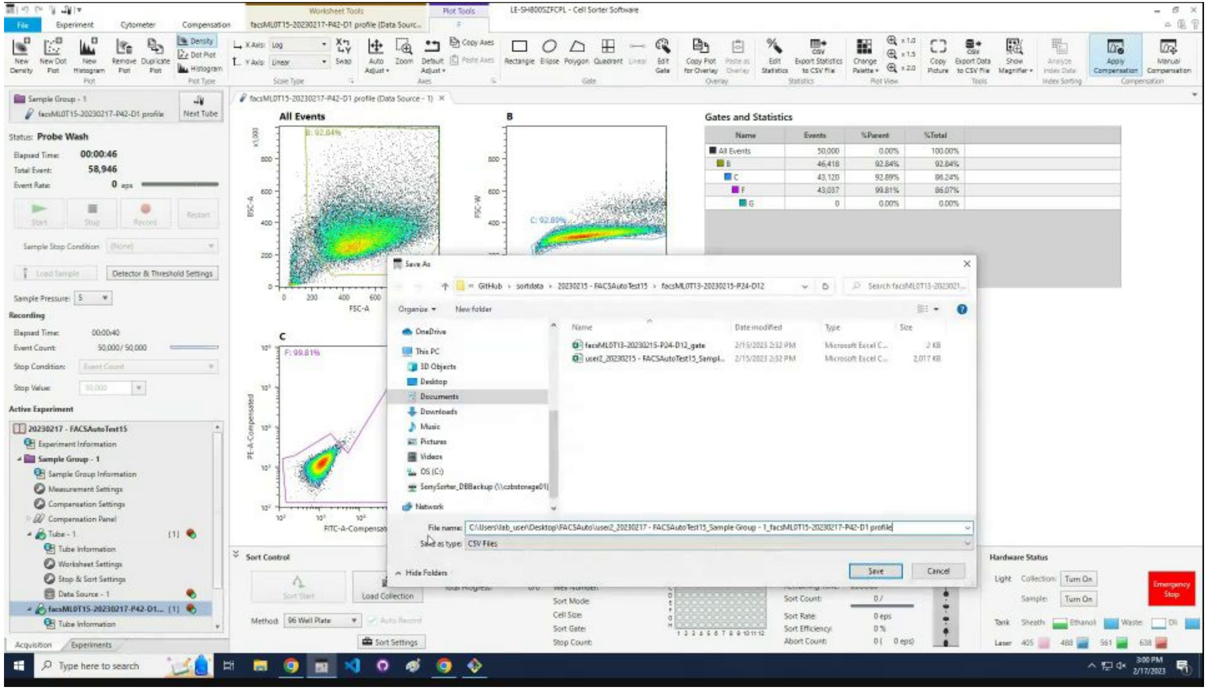

### Click `Save`.

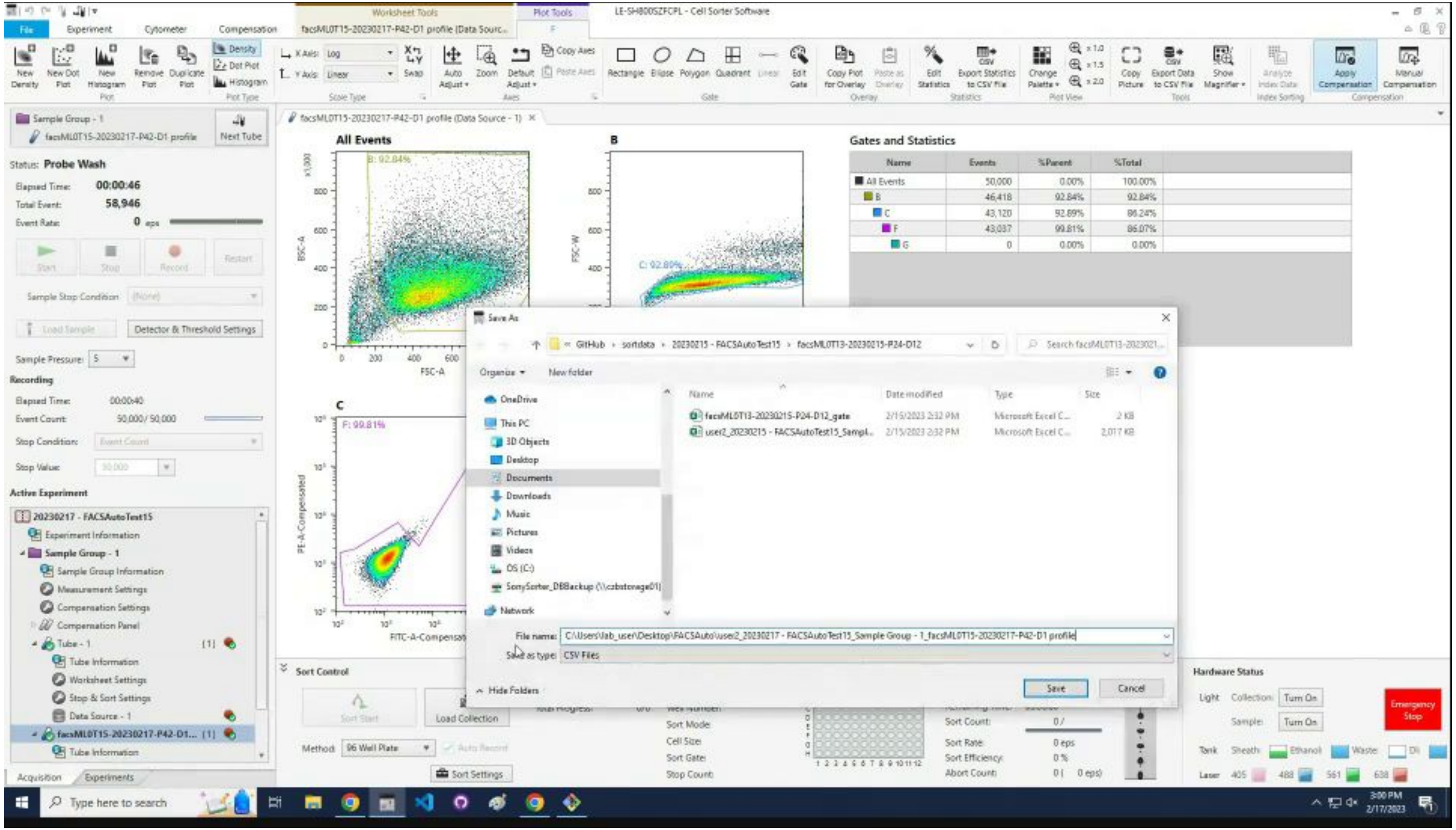

LE-SH800SZFCPL - Cell Sorter Software
File
Experiment
Cytometer
Compensation
Worksheet Tools
Plot Tools
Sample Group - 1
Next Tube
Status: Probe Wash
Elapsed Time: 00:00:46
Total Event: 58,946
All Events
Gates and Statistics
Name
Events
%Parent
%Total
All Events
50,000
0.00%
100.00%
B
46,418
92.84%
92.84%
C
43,120
92.89%
86.24%
F
43,037
99.81%
86.07%
G
0
0.00%
0.00%
Save As
New folder
OneDrive
This PC
3D Objects
Desktop
Documents
Downloads
Music
Pictures
Videos
Network
File name:
Save as type: CSV Files
Hide Folders
Save
Cancel
Sort Control
Hardware Status
Emergency Stop
Active Experiment
20230217 - FACSAutoTest15
Experiment Information
Sample Group Information
Measurement Settings
Compensation Settings
Compensation Panel
Tube Information
Worksheet Settings
Stop & Sort Settings
Data Source - 1
Acquisition
Experiments
Type here to search

### Click `Export`.

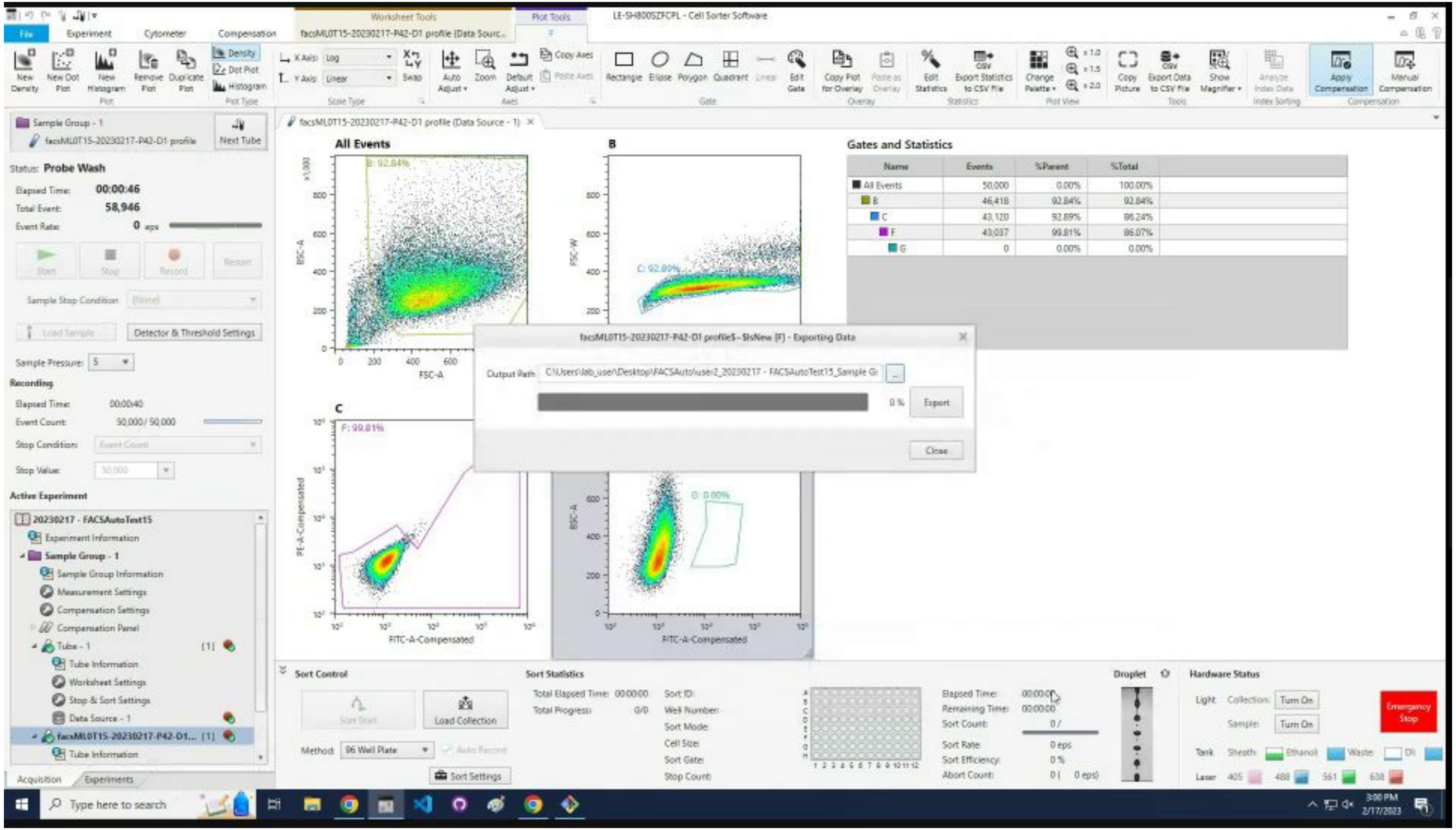

### Wait for the export to finish.

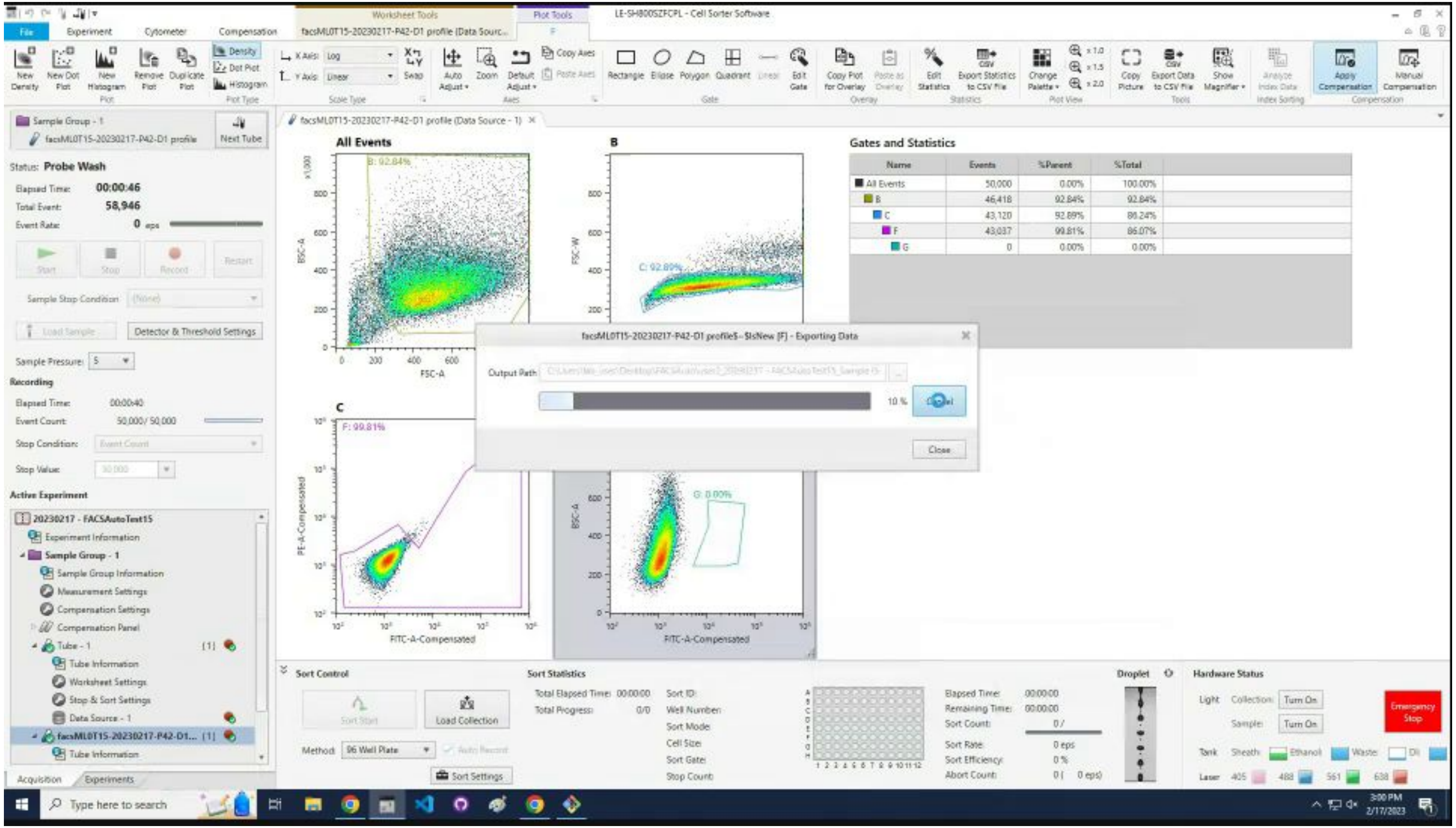

### Click OK.

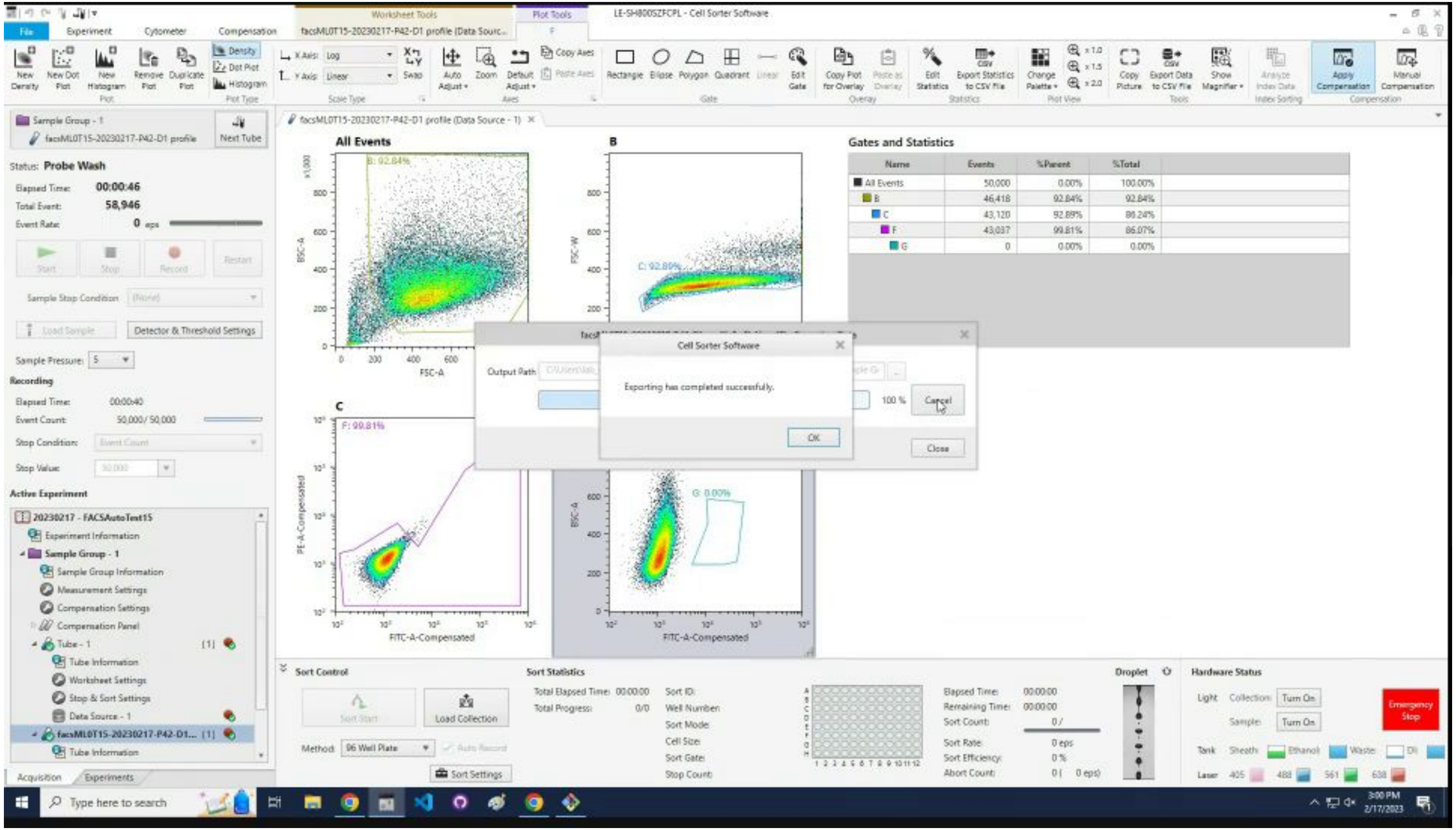
Cell Sorter Software
Exporting has completed successfully.
OK
Cancel
Close
Gates and Statistics
All Events

### Click `Close`.

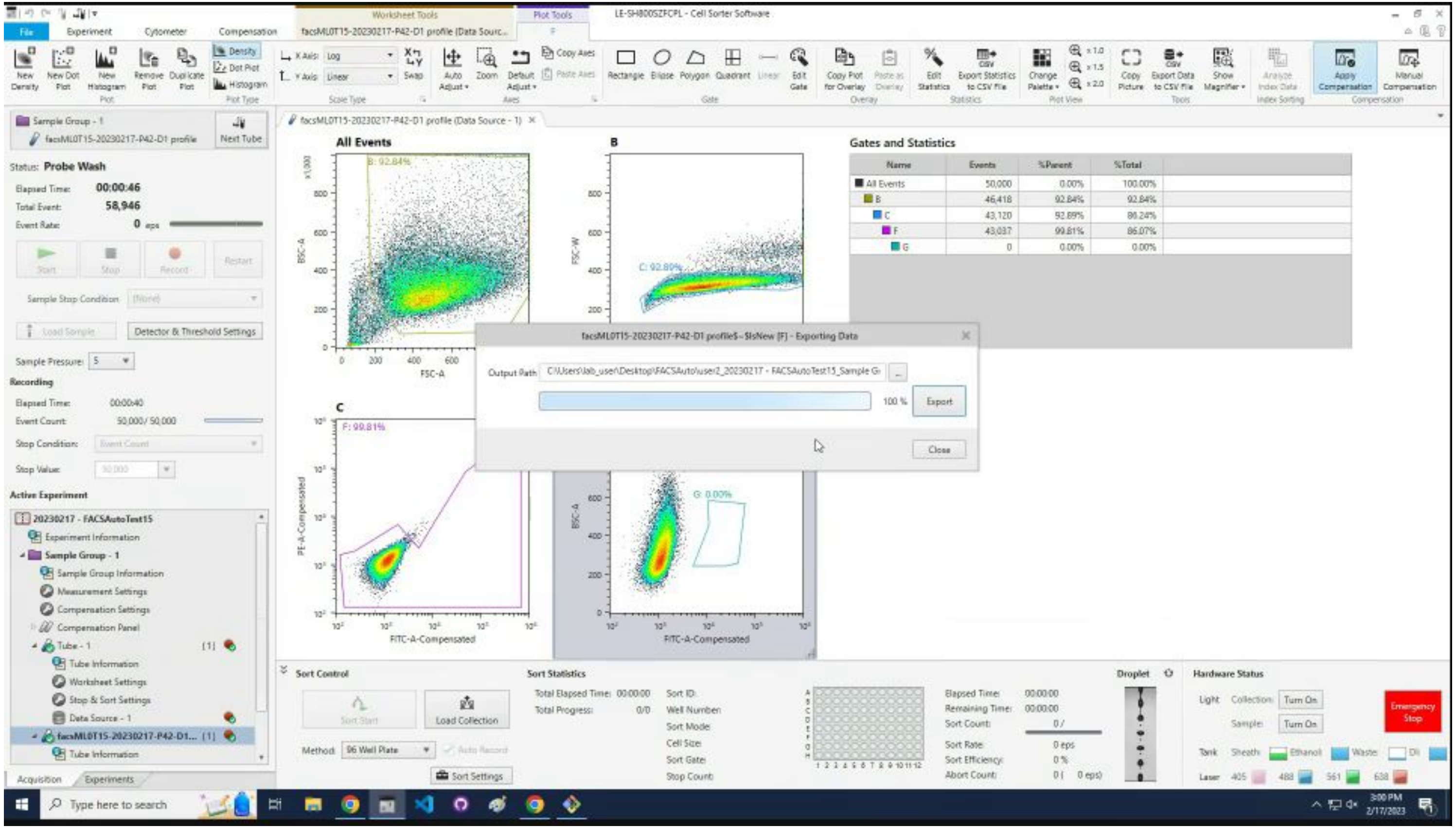
LE-SH800SZFCPL - Cell Sorter Software
facsMLOT15-20230217-P42-D1 profile$-$lsNew (F) - Exporting Data
Output Path
100 %
Export
Close
All Events
Gates and Statistics
Name
Events
%Parent
%Total
Sort Control
Sort Statistics
Hardware Status
Status: Probe Wash
Active Experiment
Type here to search

### Click the X to close the tube tab.

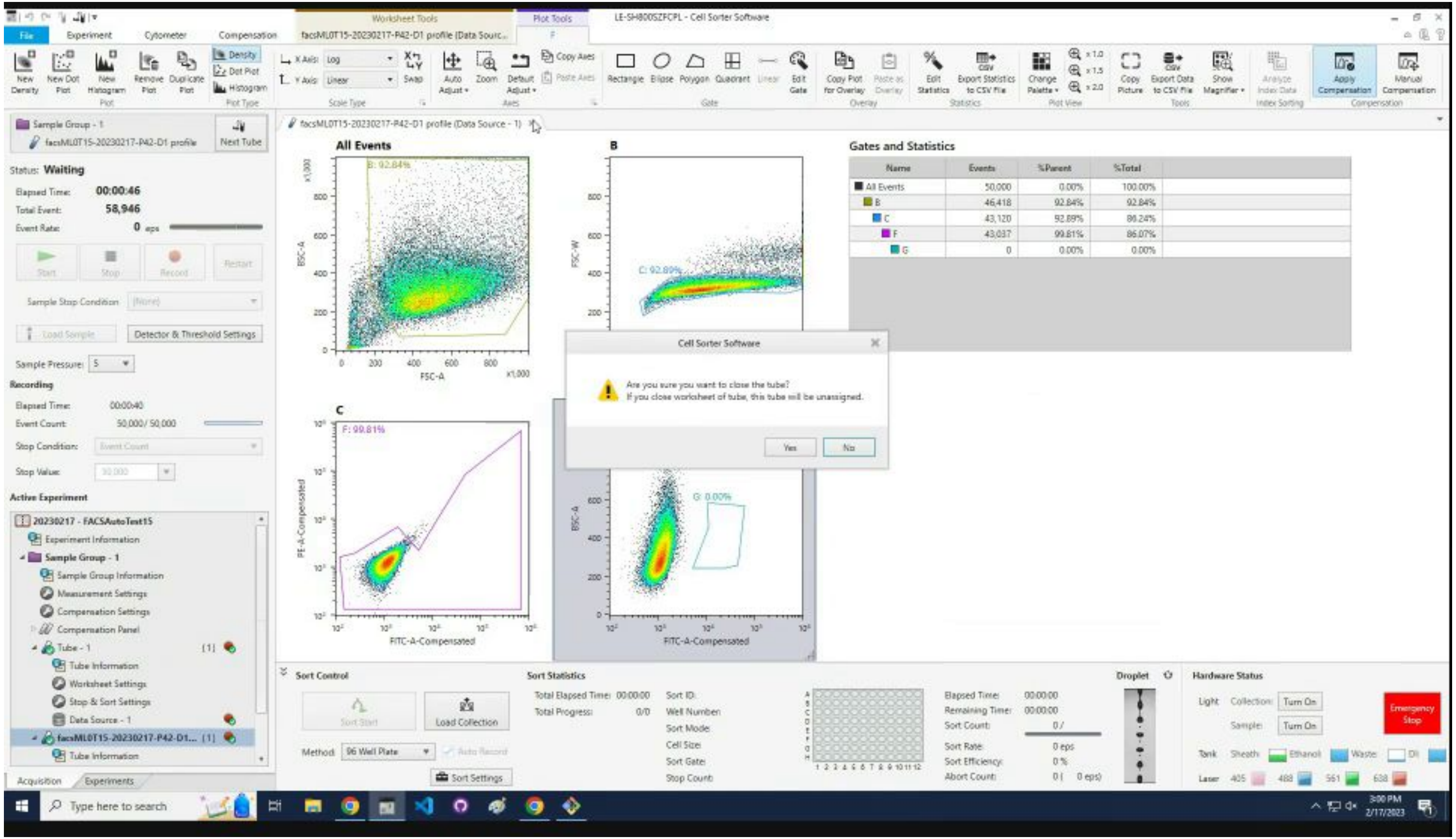

### Click `Yes` to confirm closing the tube tab.

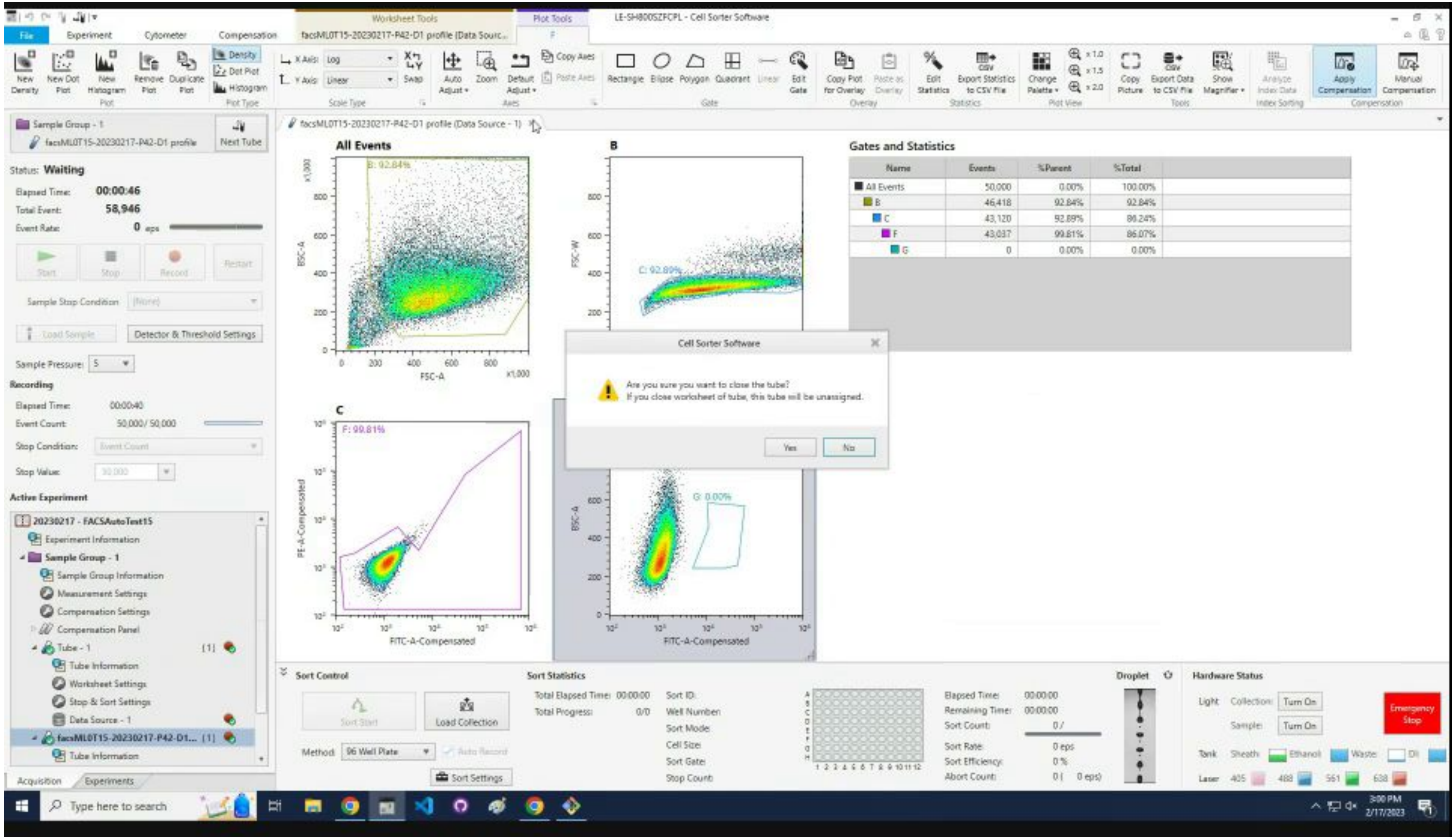

### Wait for the automation to create the gate.

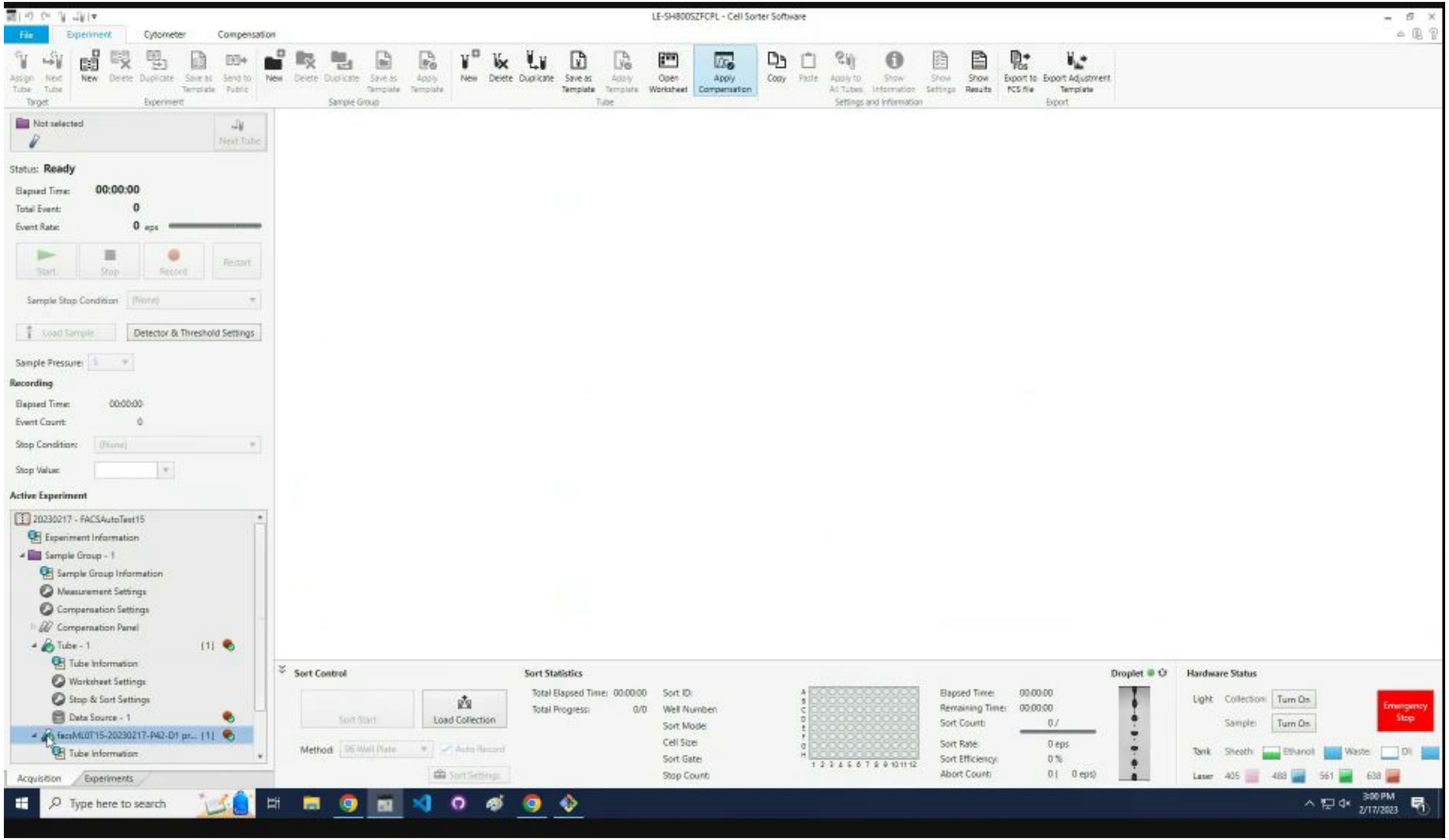

### Click on the profile sample in the list.

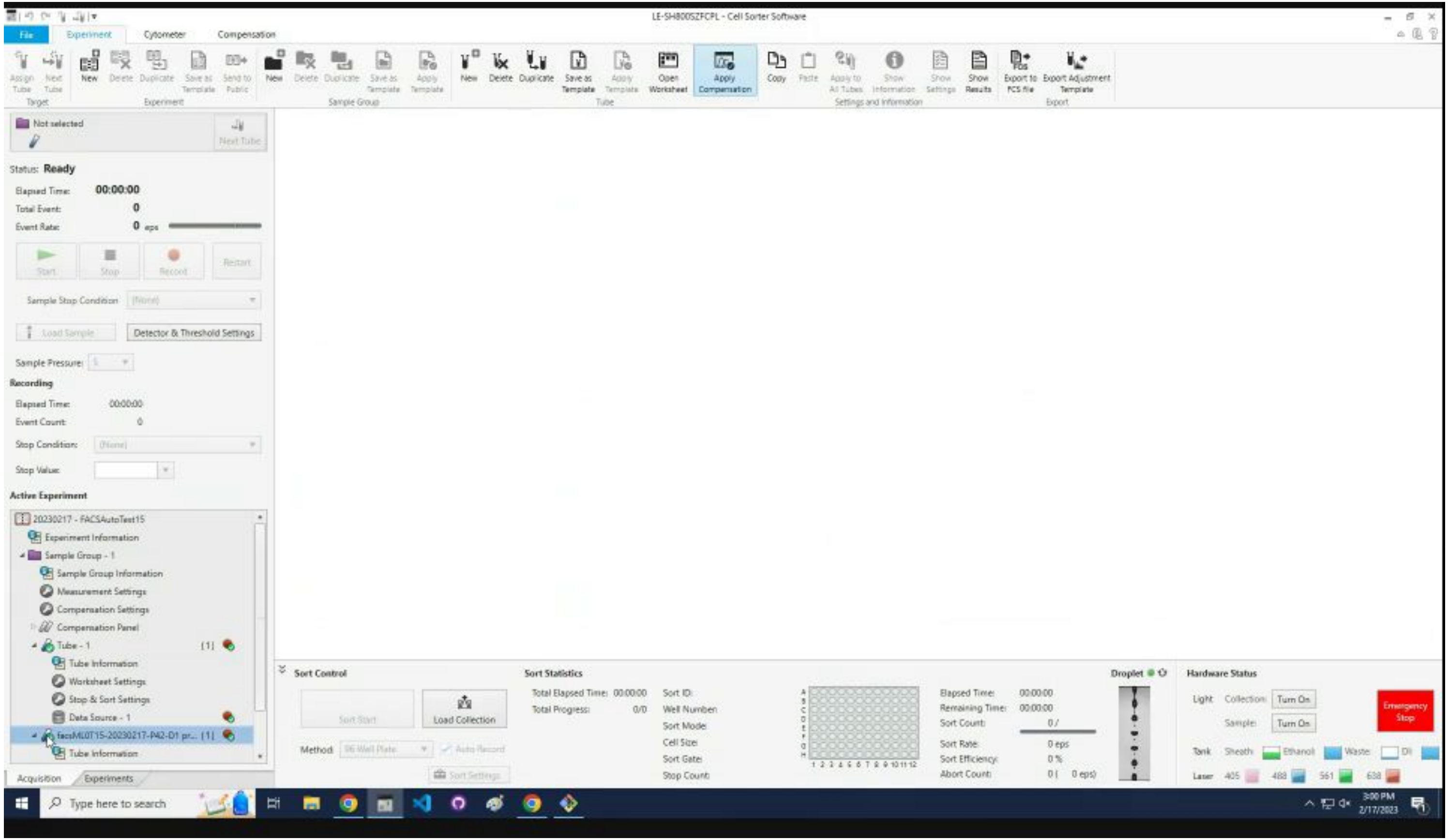

### Click the `Duplicate` icon.

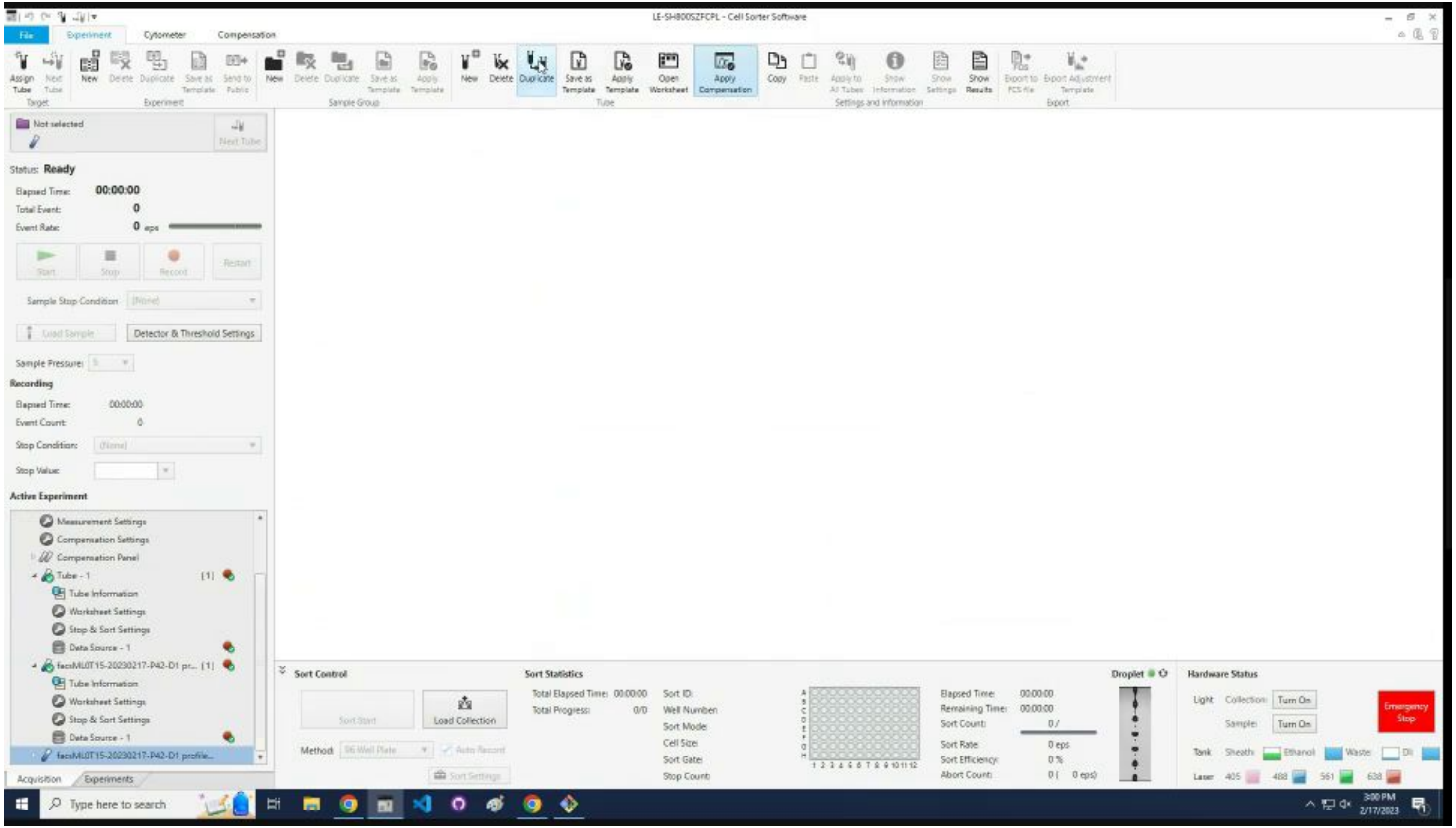

### Click the `Assign Tube` icon.

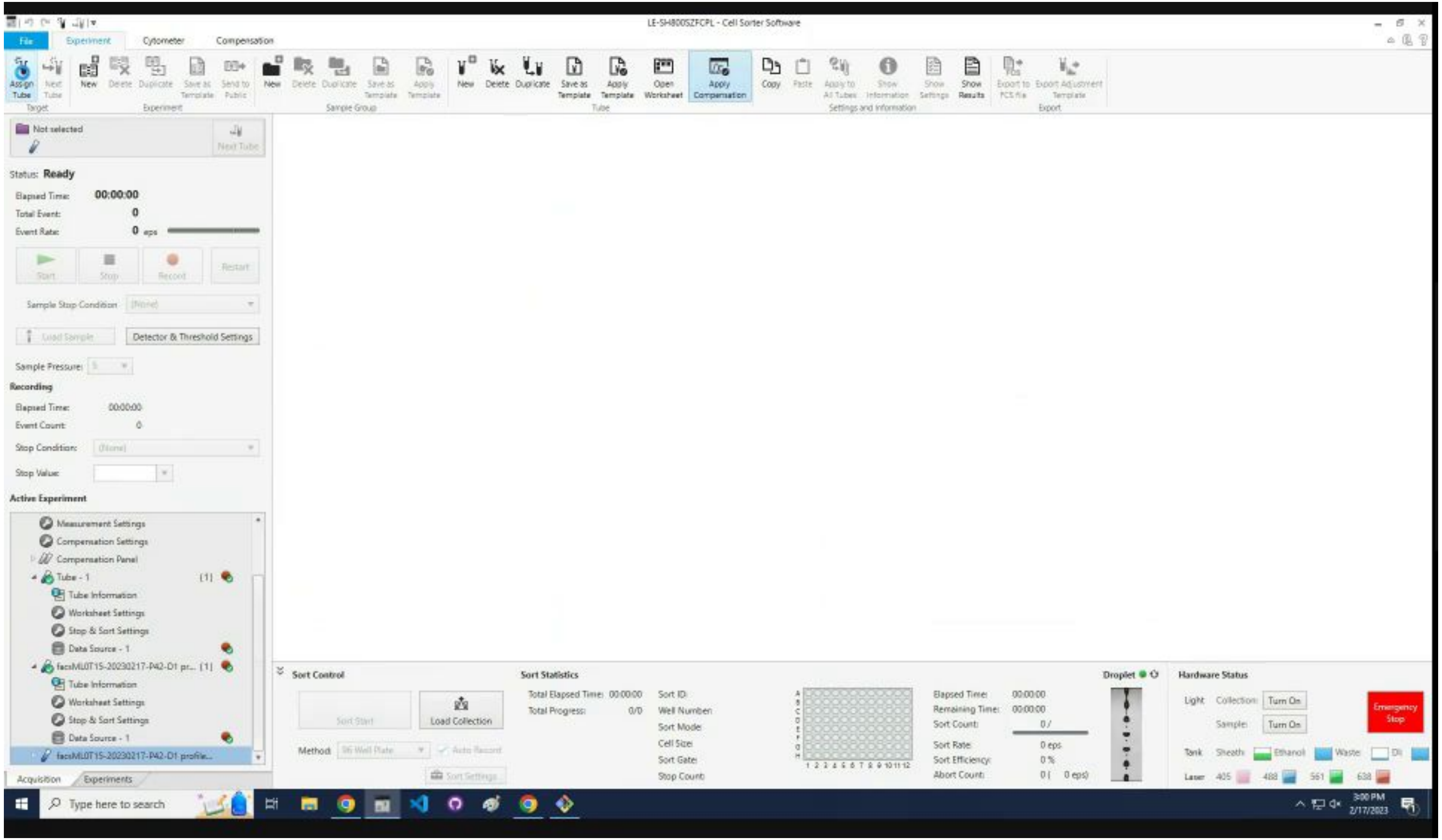

### Check that the gate was properly drawn.

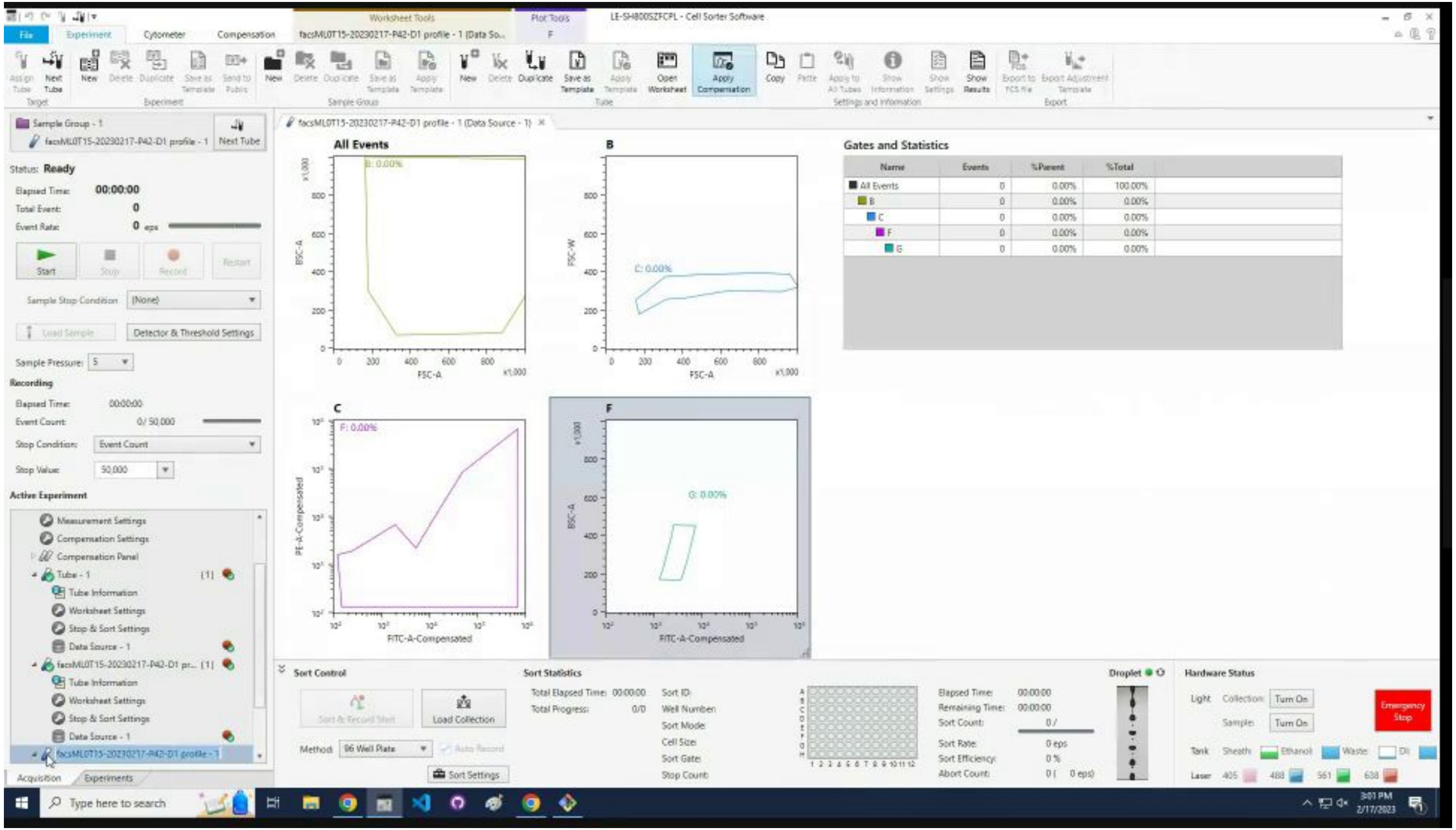

### Press `F2` to rename the tube.

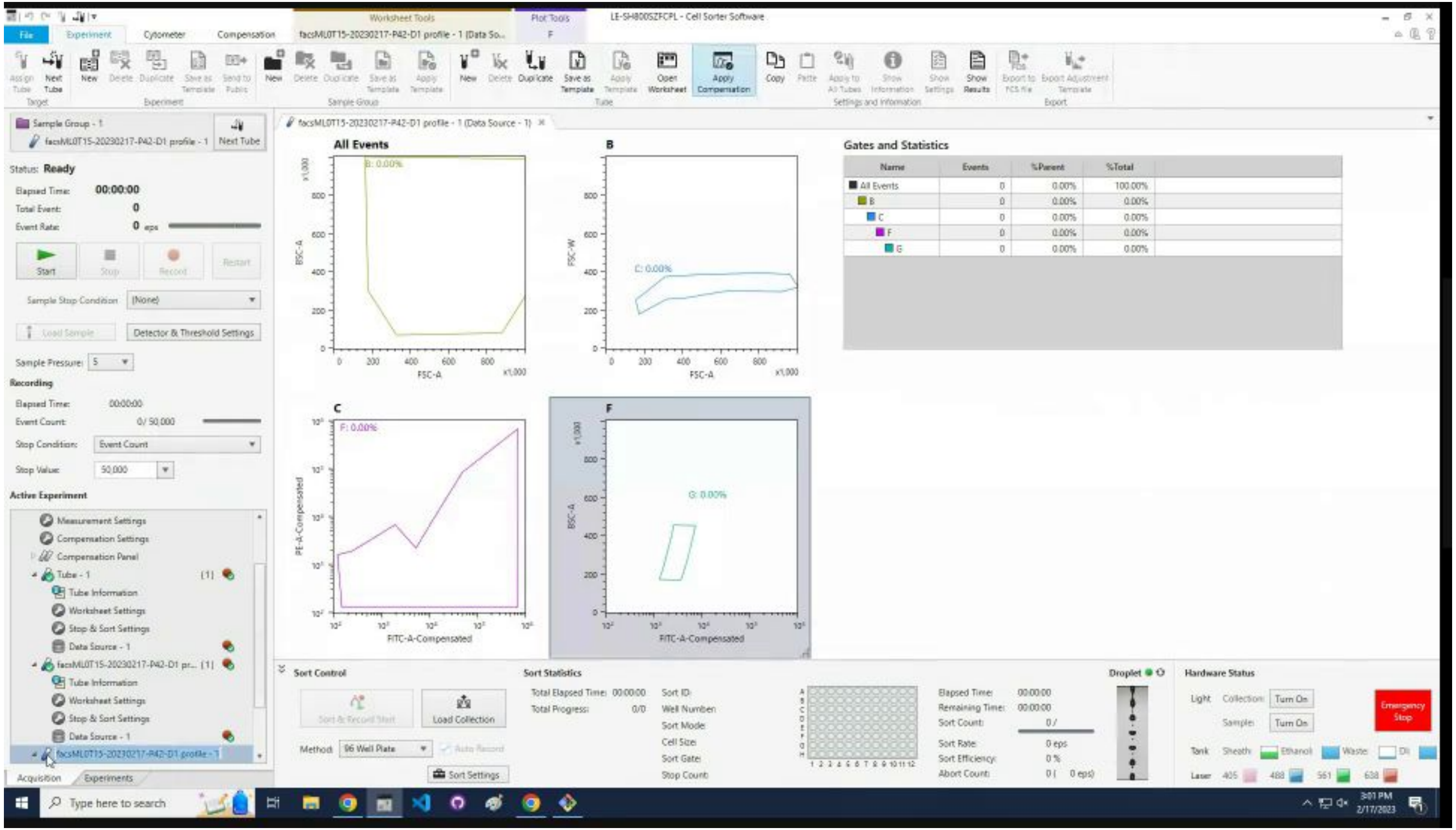

### Rename to the sample name appended with sort.

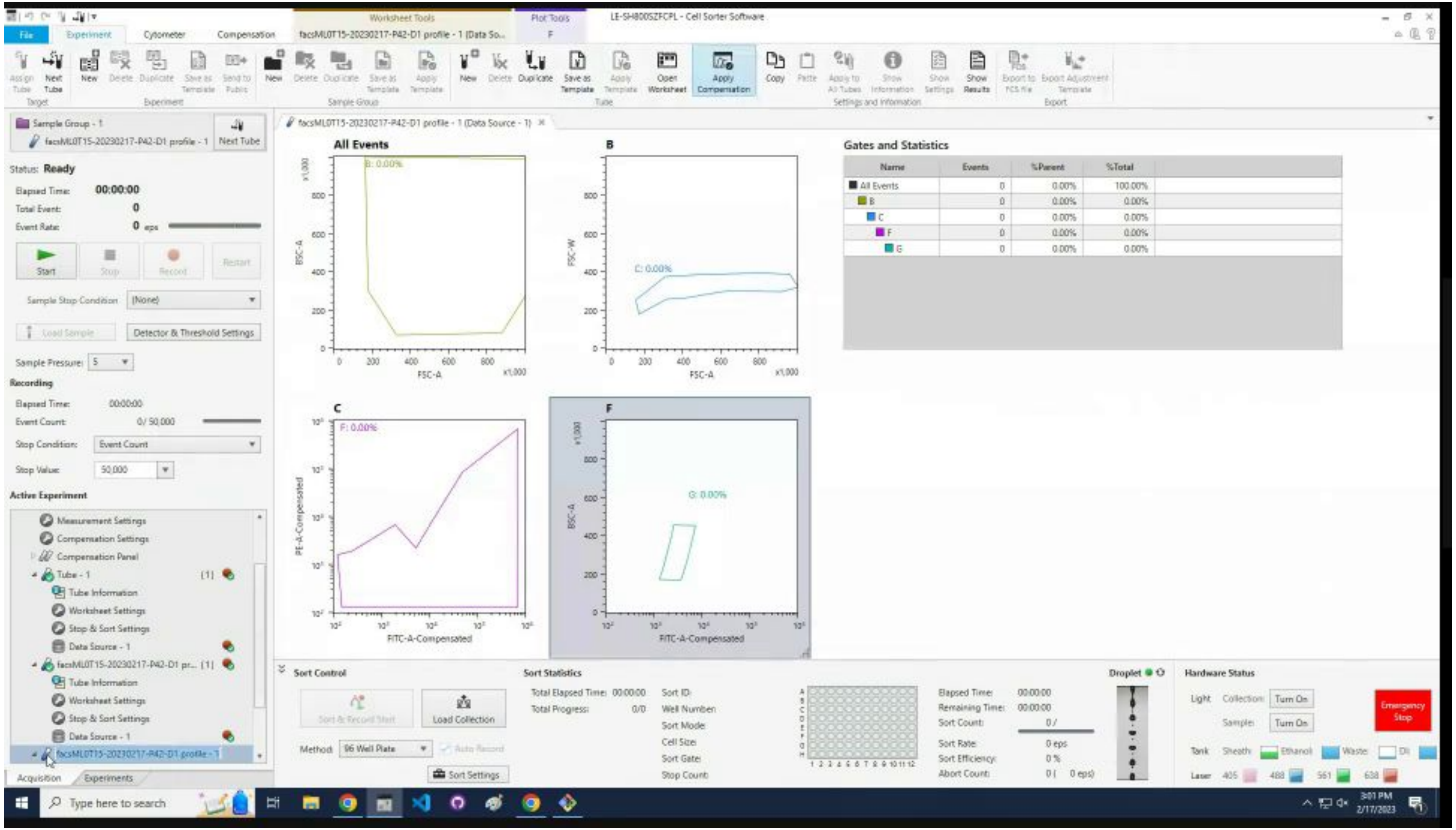

### Set the `Sample Pressure` likely to 5.

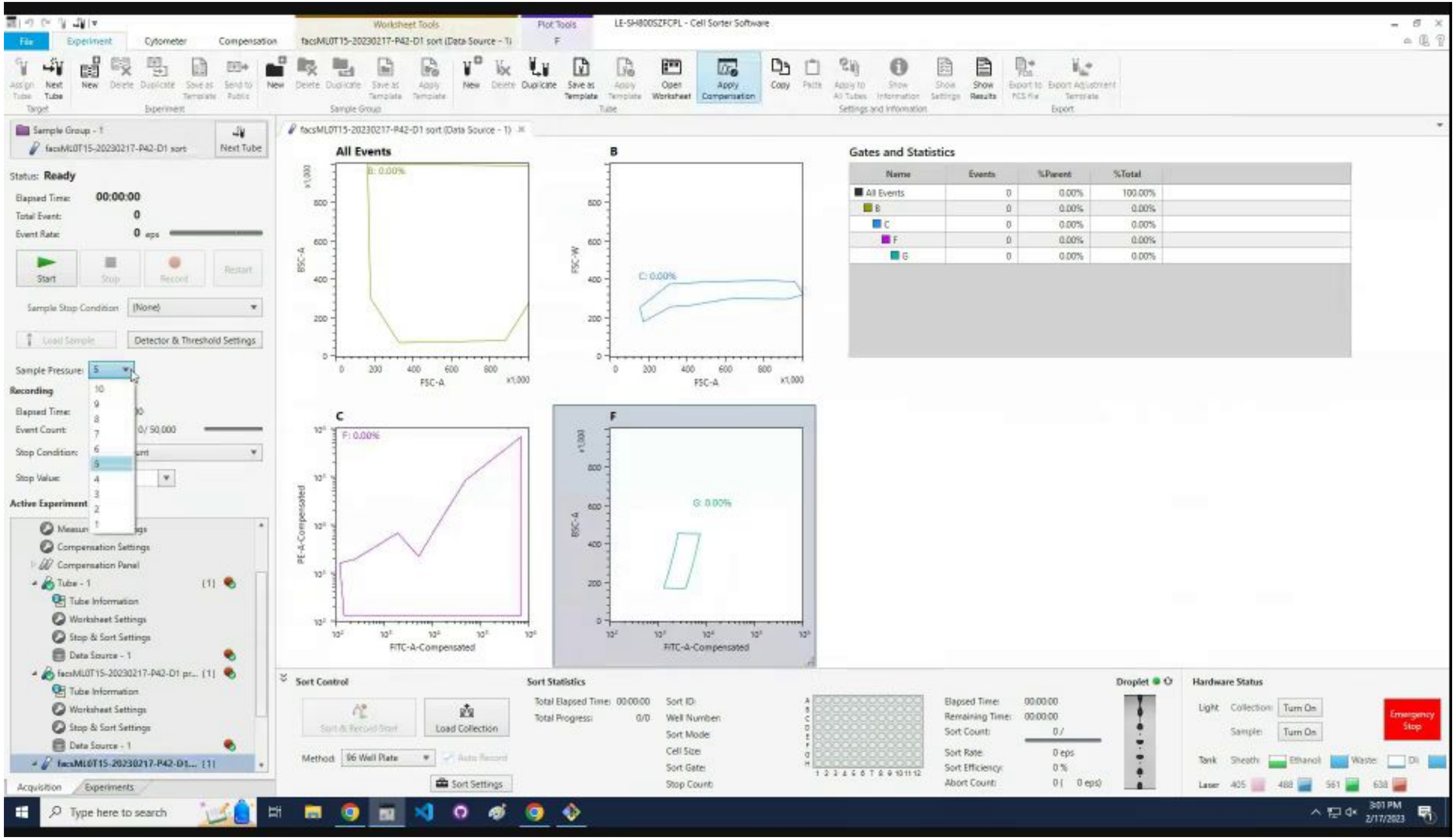

### Change the `Stop Condition` to `None`.

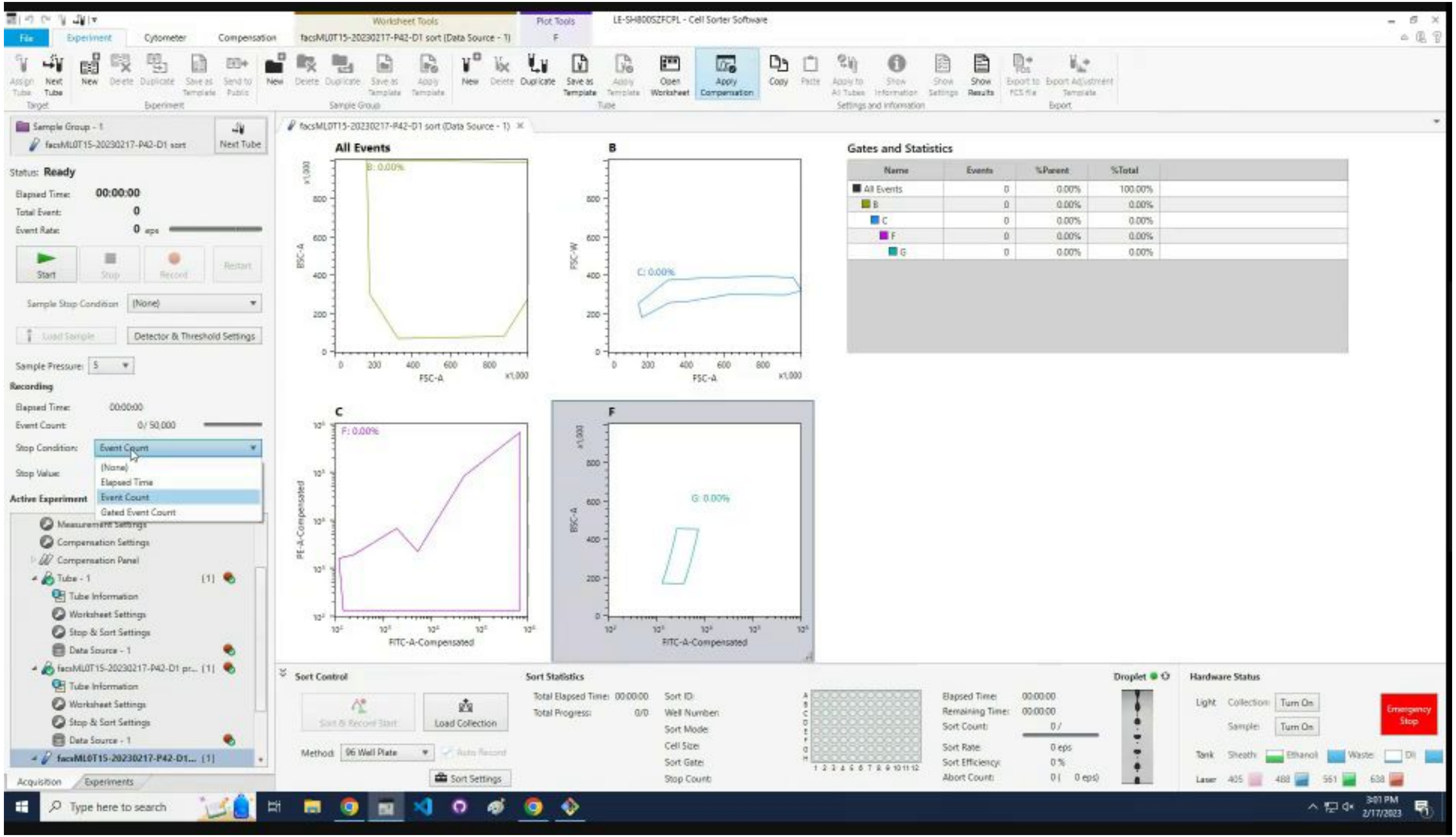

### Click `Sort Settings.`

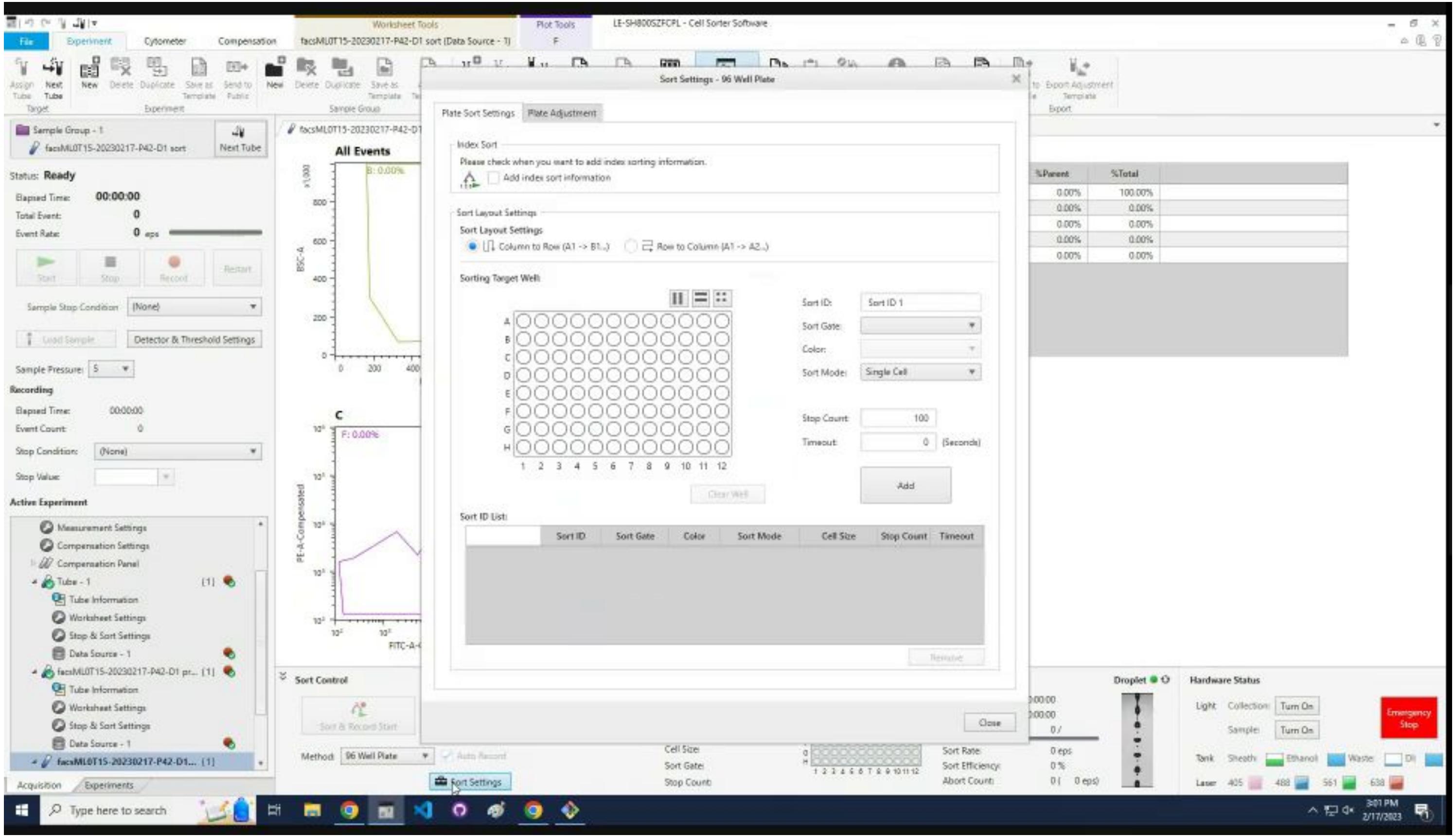

### If it is the first sort, it loads without any sort settings.

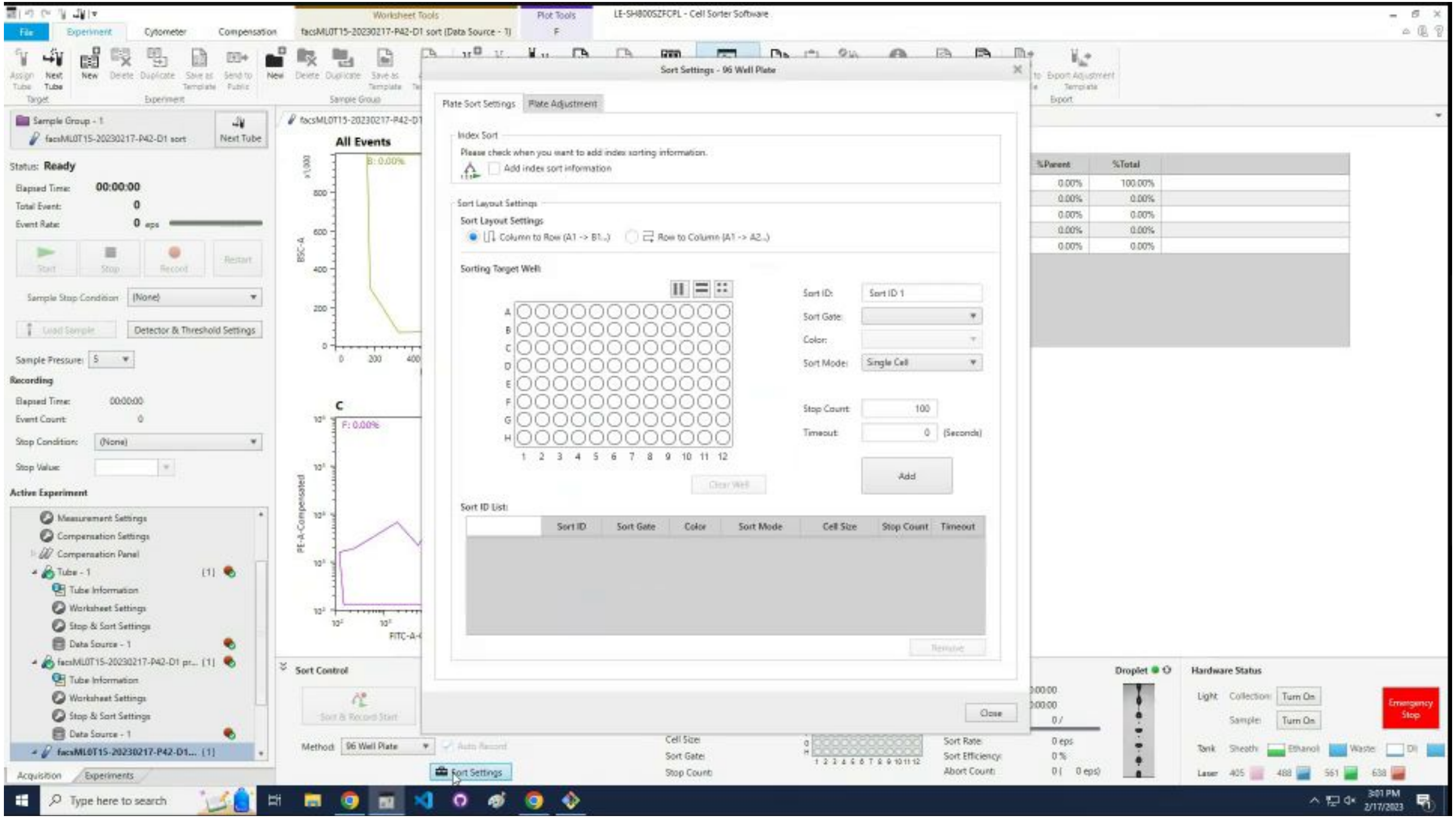

### If is not the first sort, it loads with the previous settings.

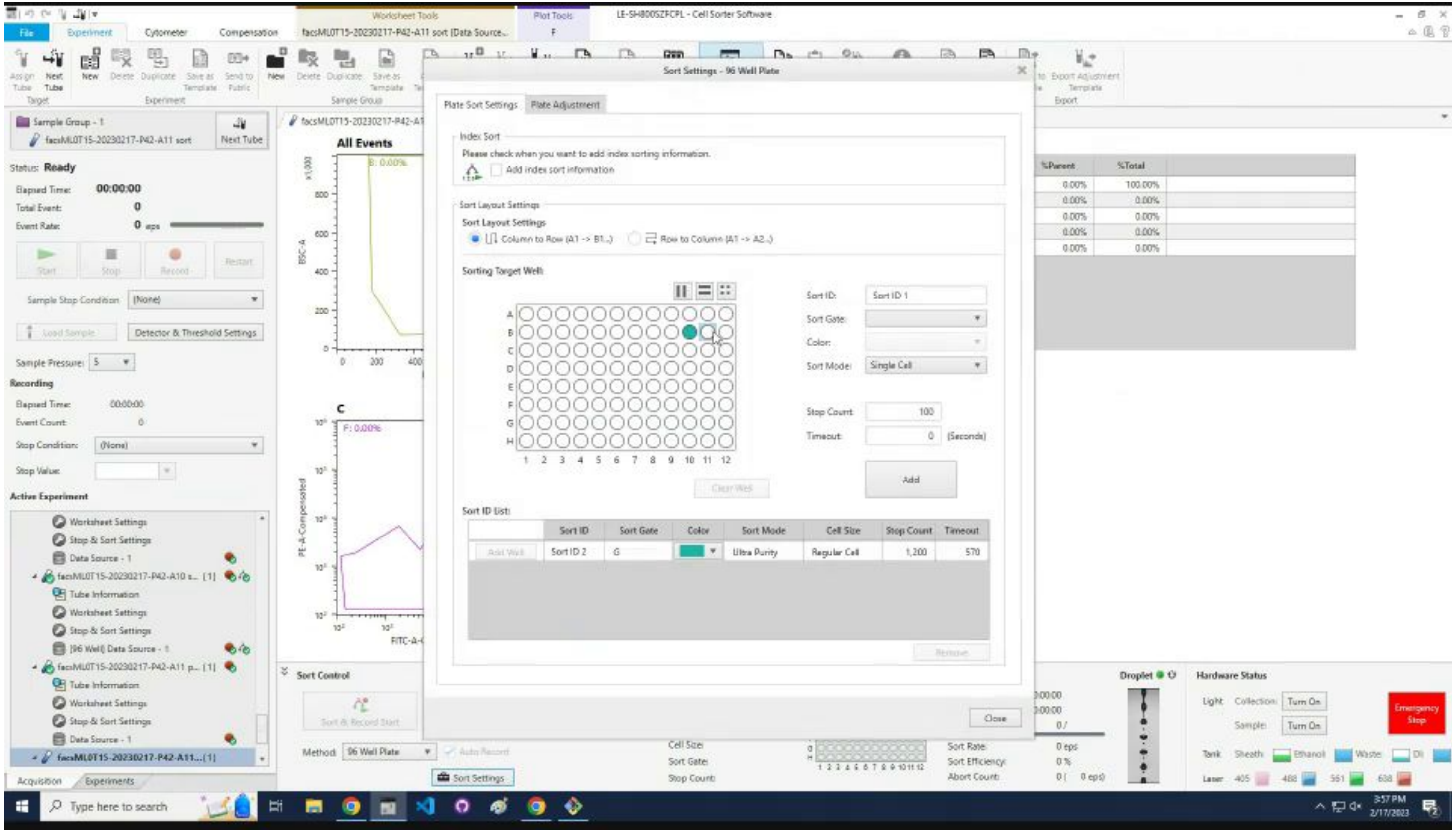

### Select the well into which the desired cells should go.

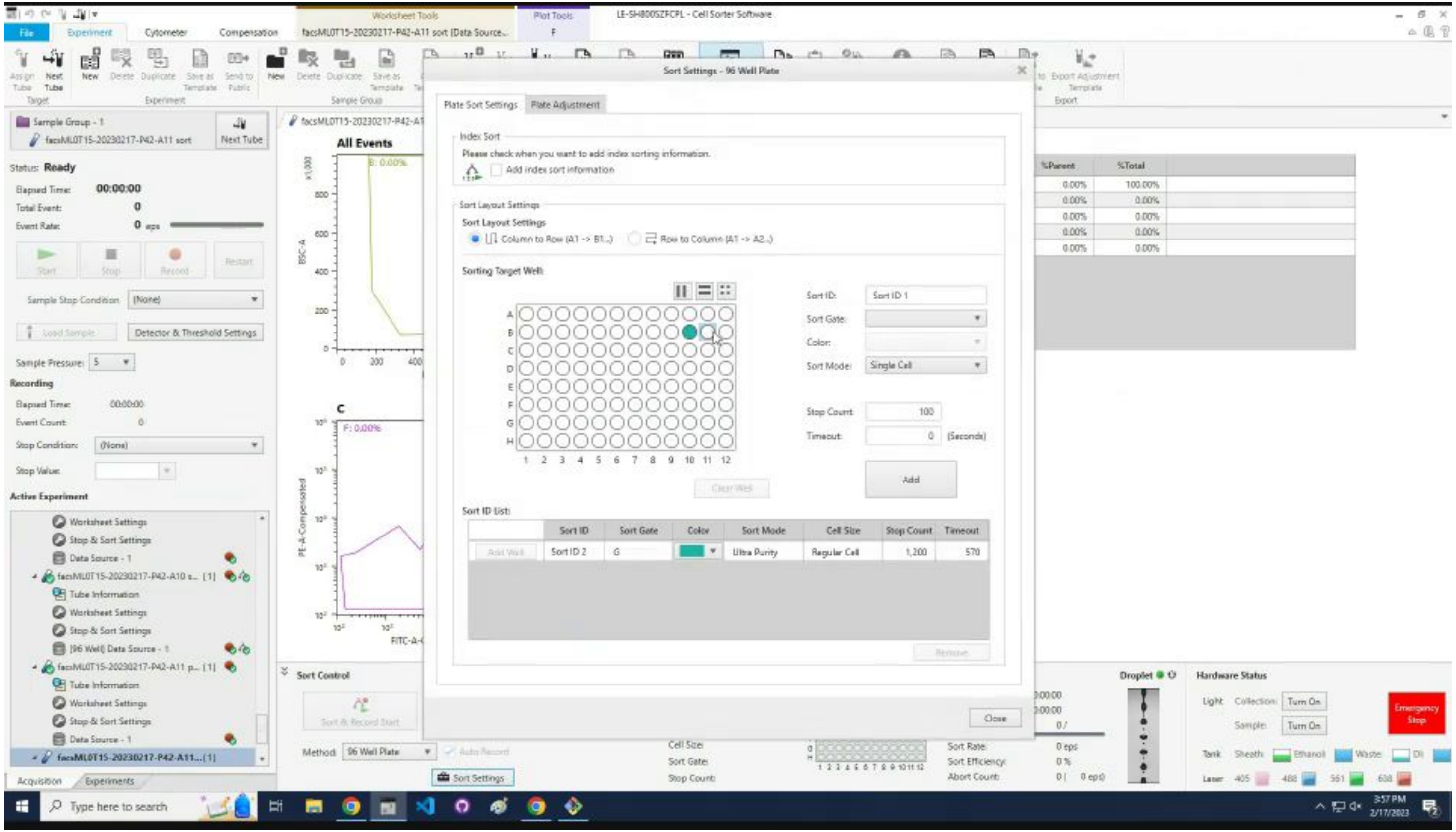

### Click the `Sort Gate` drop down.

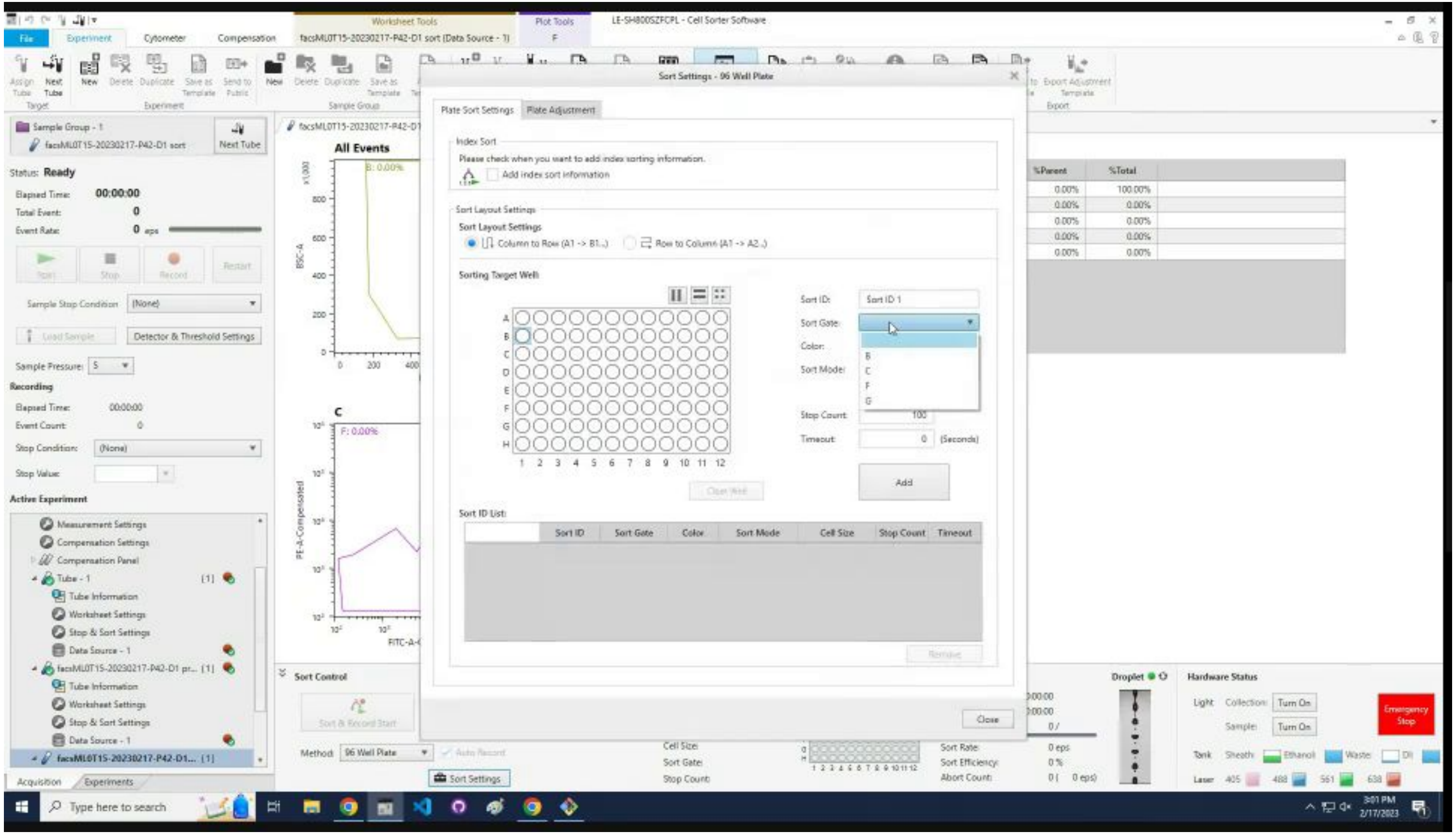

### Select the last gate.

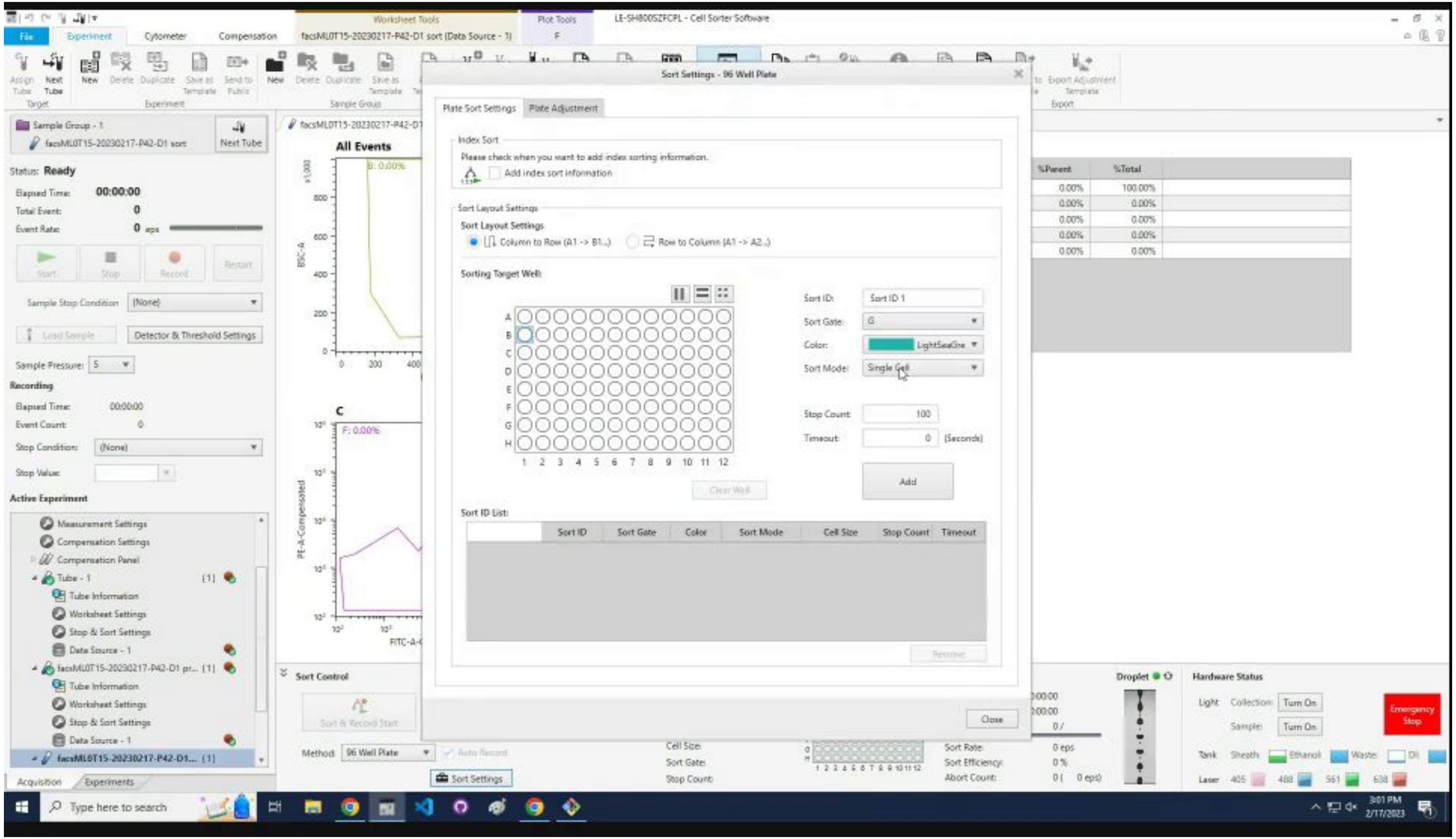

### Select the `Sort Mode` drop down.

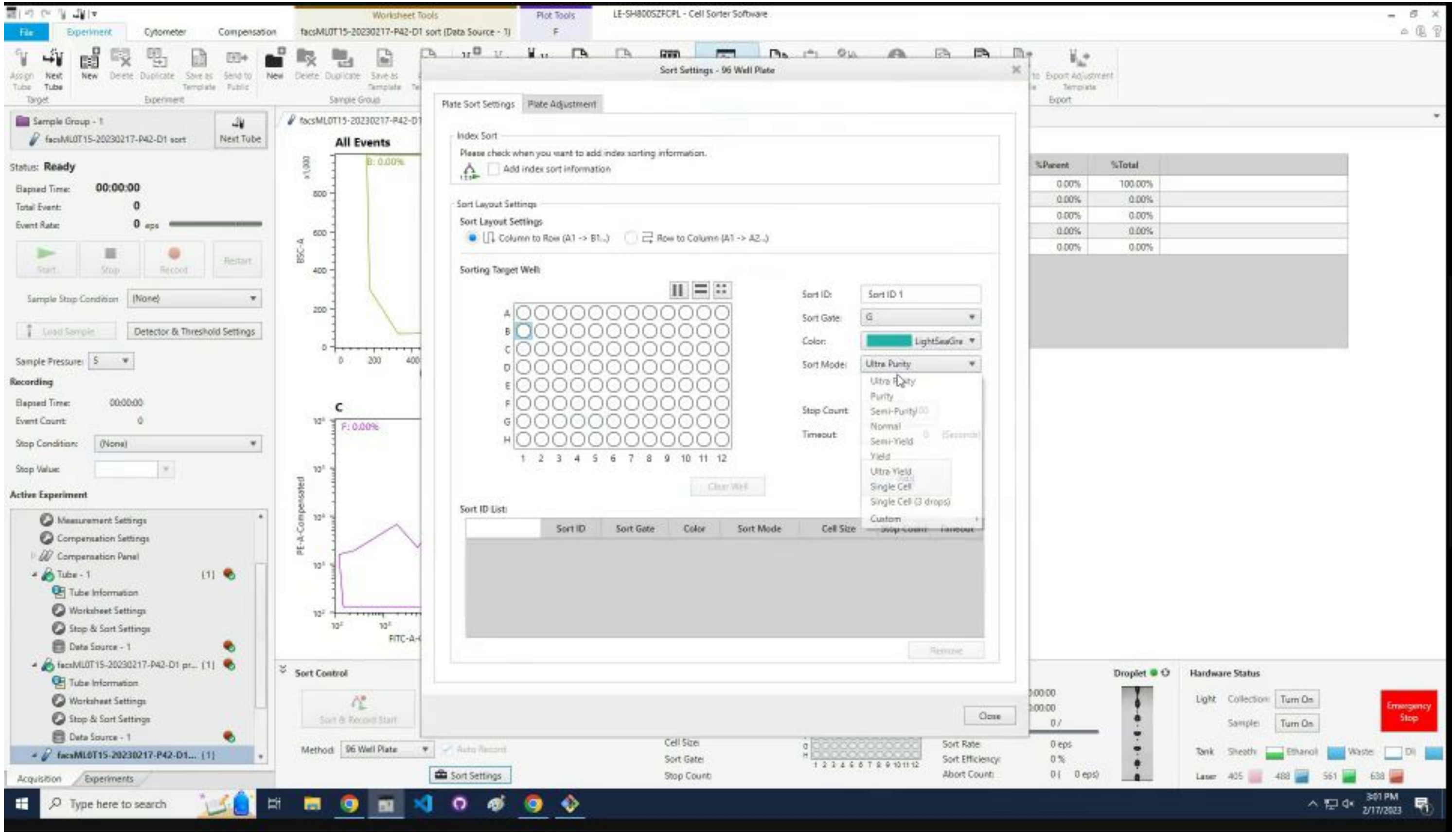

### Select `Ultra Purity`.

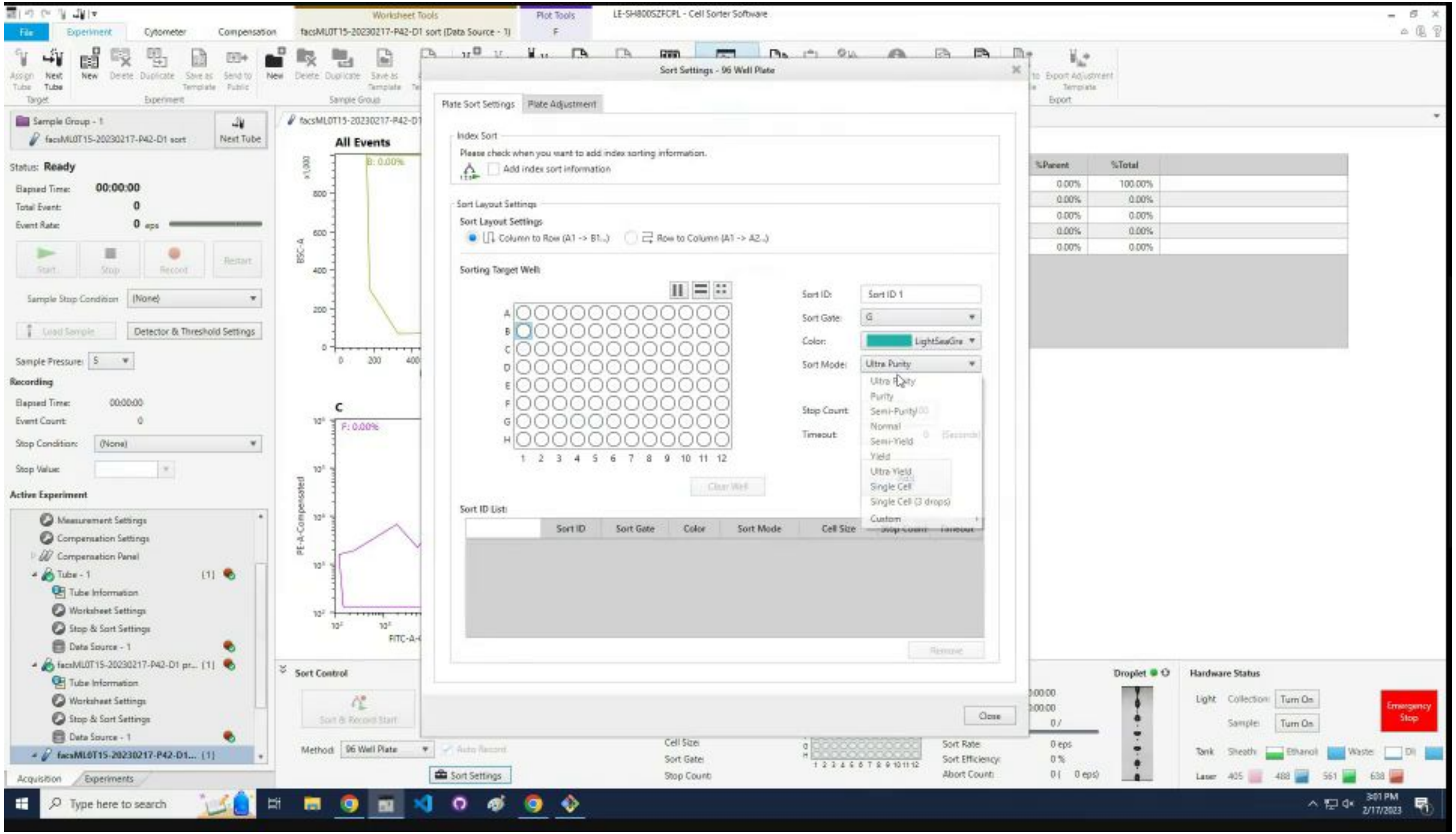

### Click in the `Stop Count`.

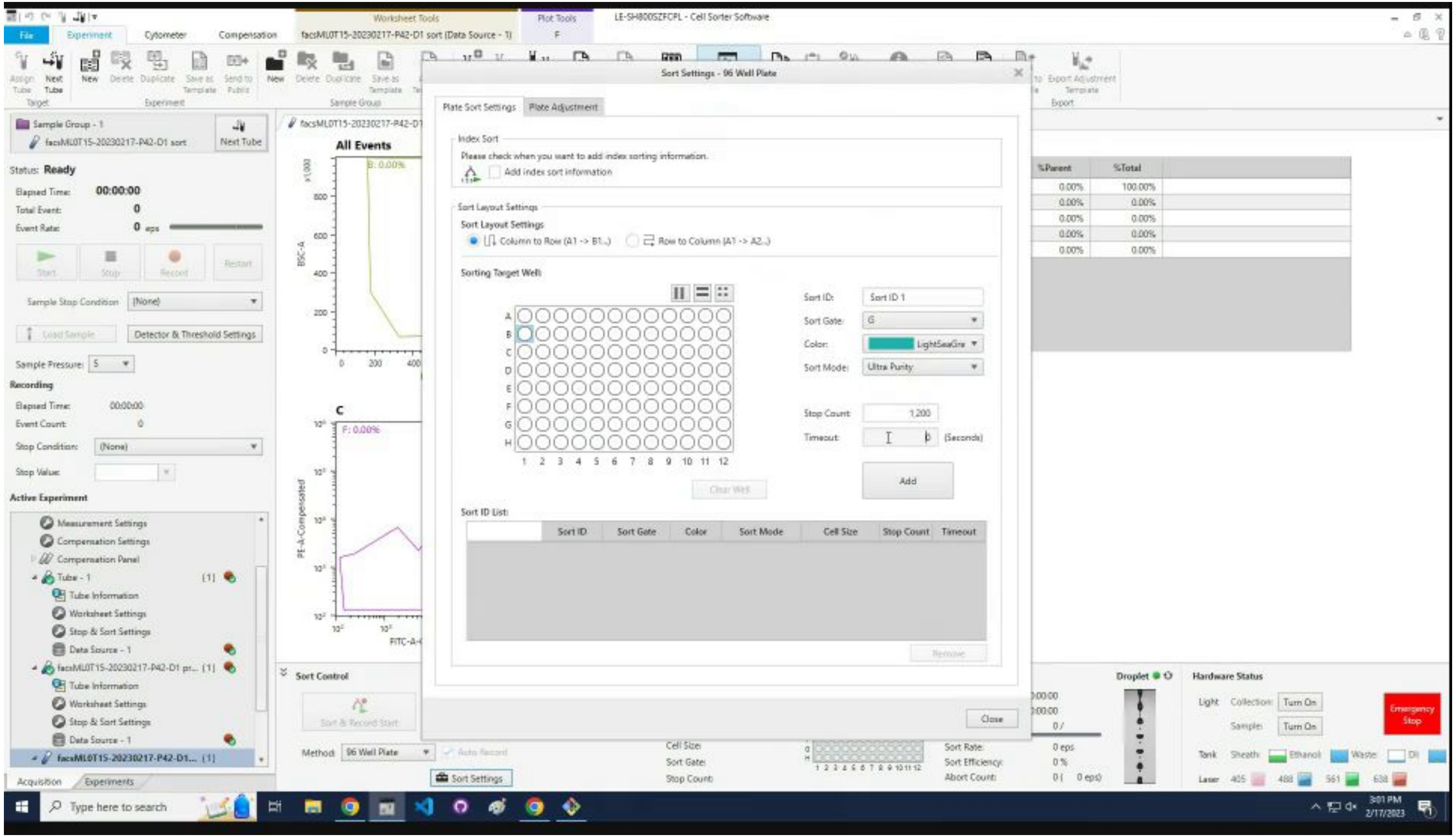

### Type in the desired count, likely `1200`.

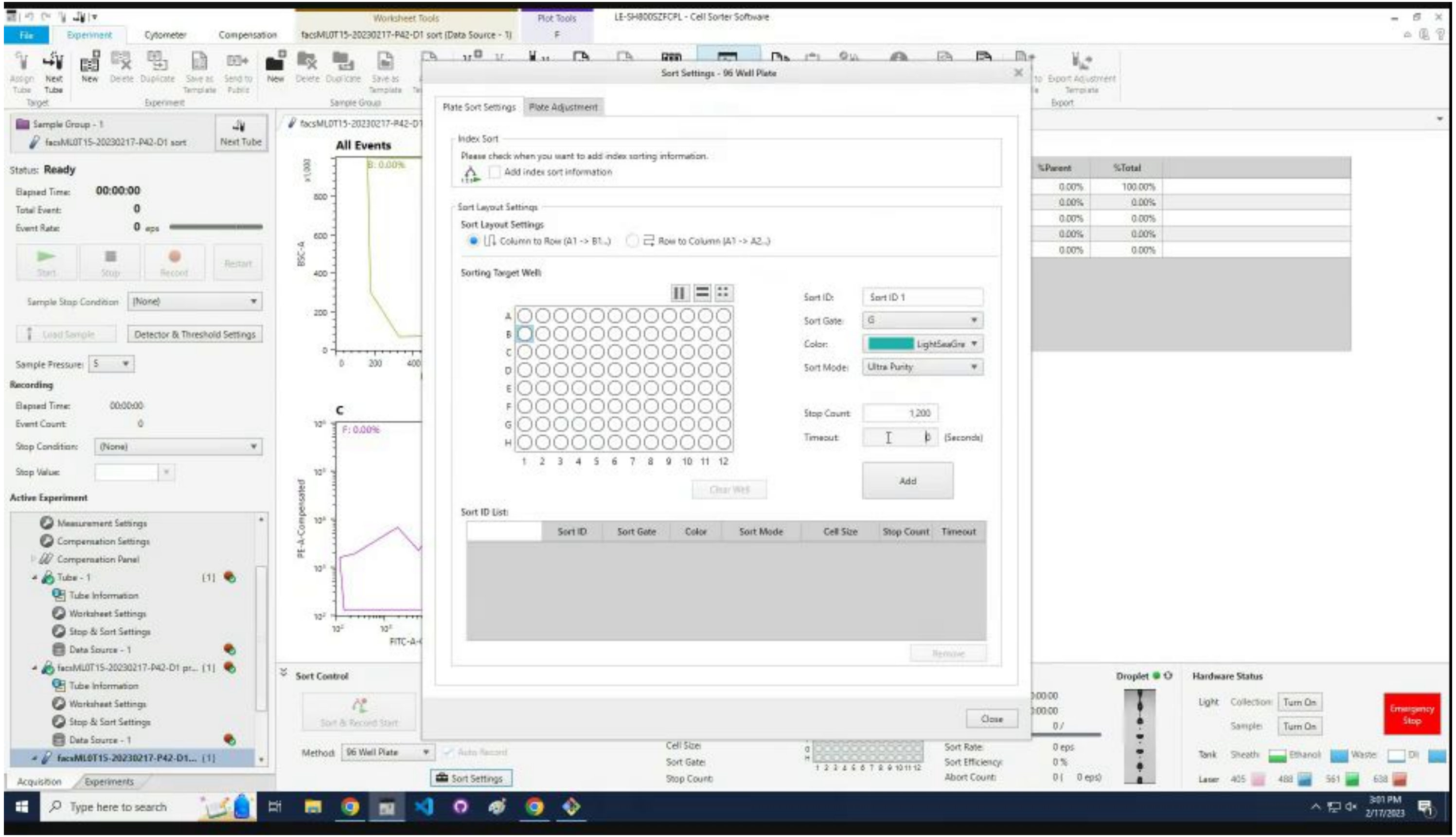

### Click in the `Timeout`.

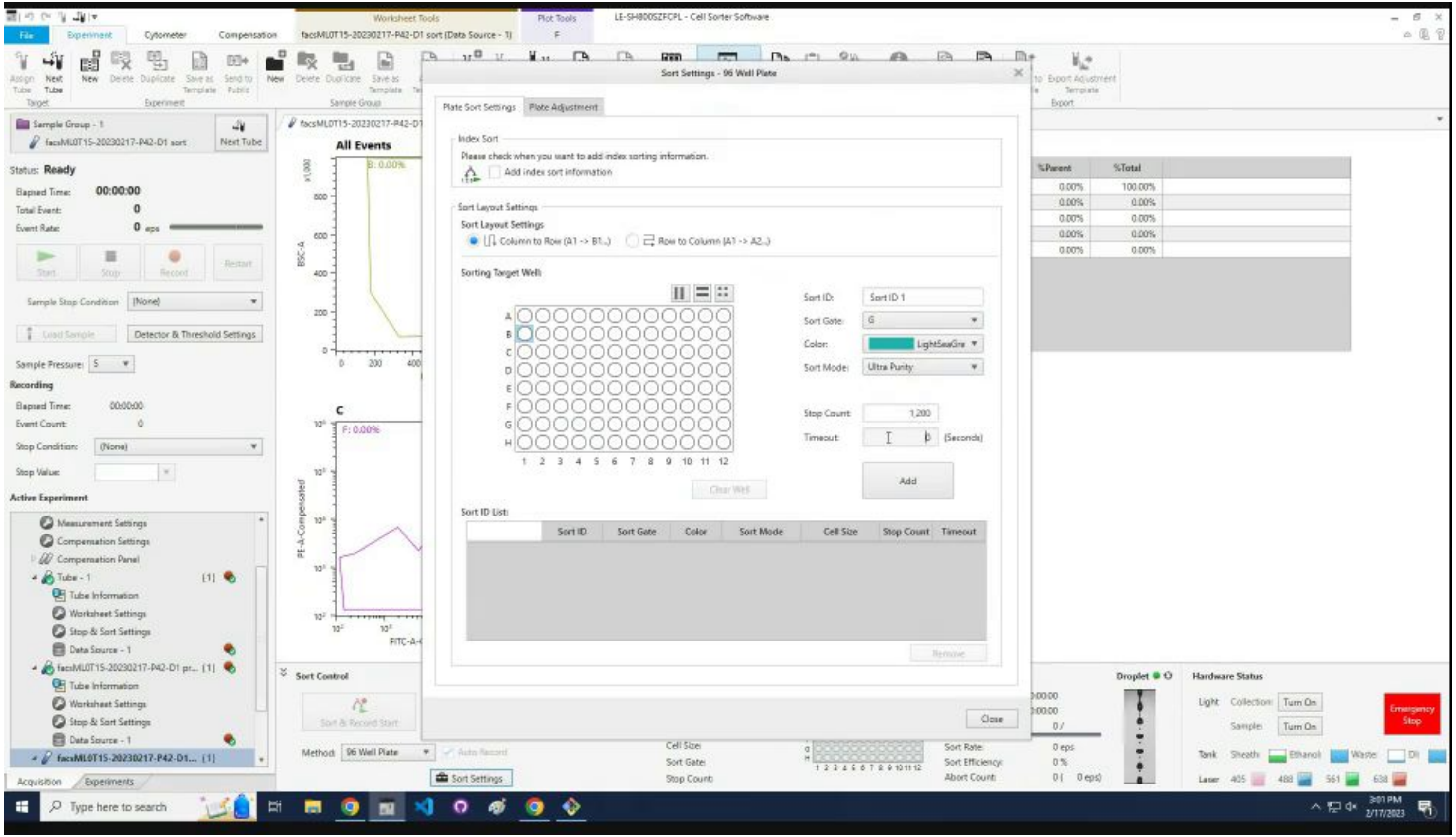

### Type in the time, likely `570`.

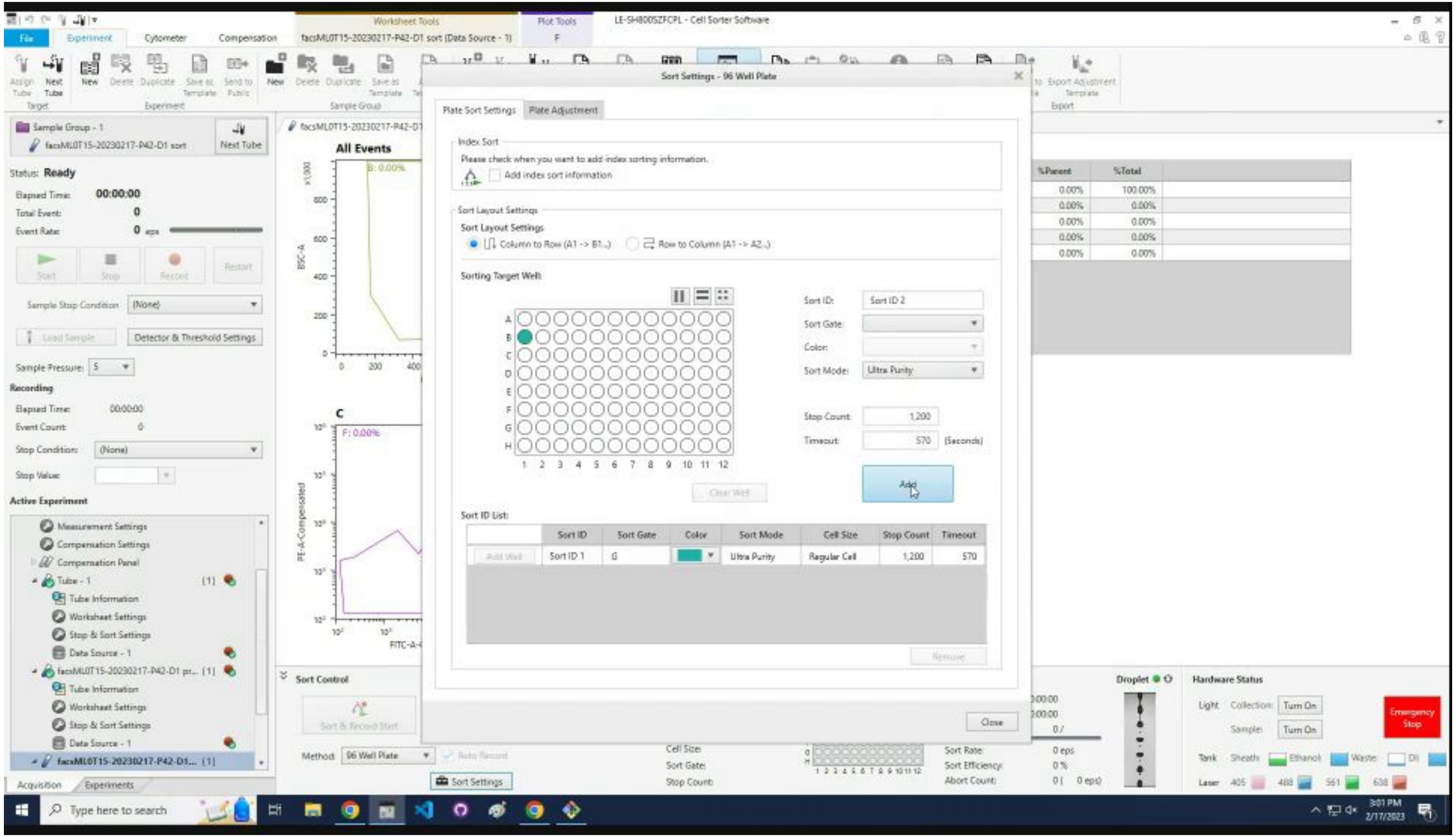

### Click Add.

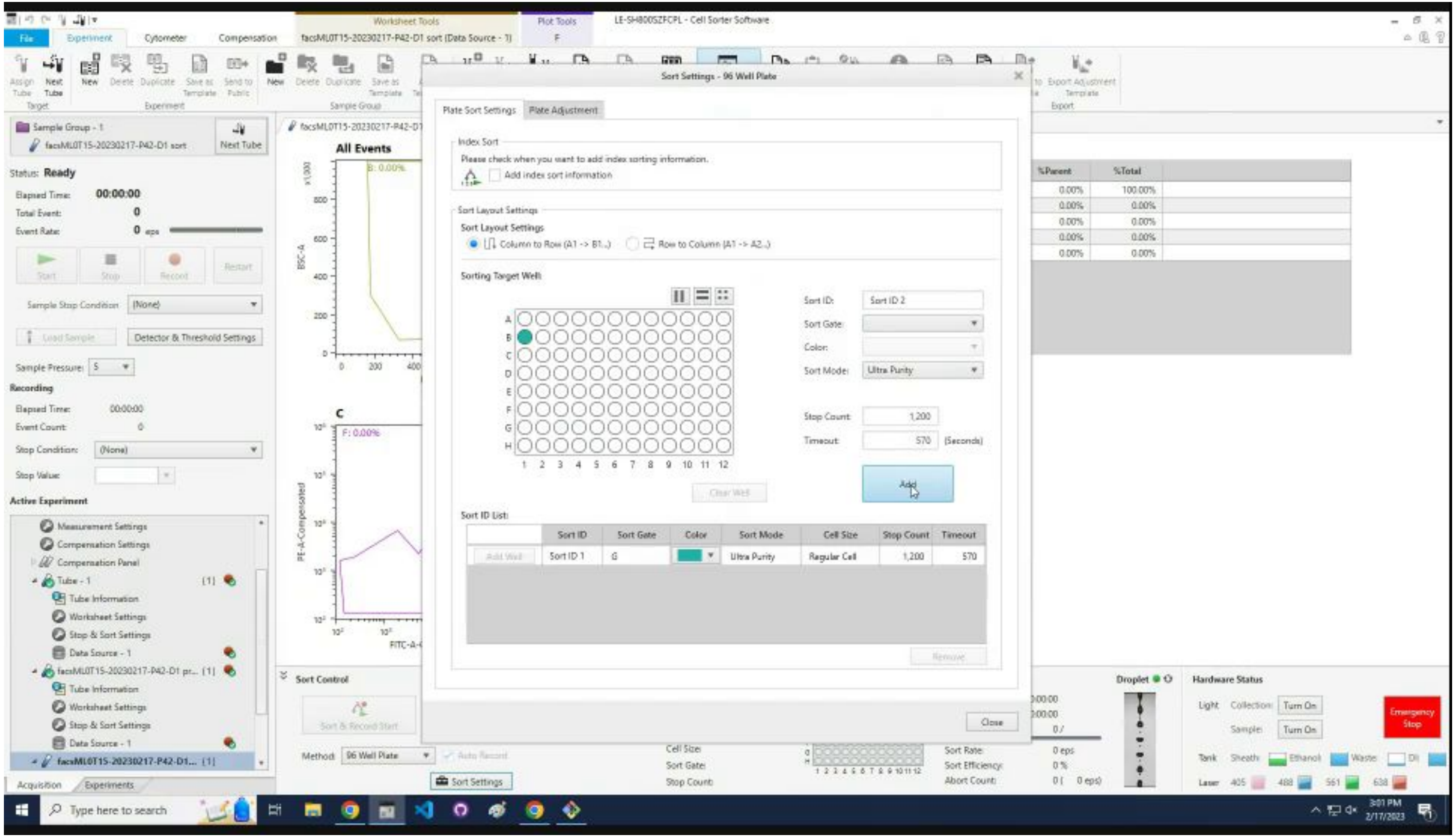
LE-SH800SZFCPL - Cell Sorter Software
File
Experiment
Cytometer
Compensation
Worksheet Tools
Plot Tools
Sort Settings - 96 Well Plate
Plate Sort Settings
Plate Adjustment
Index Sort
Please check when you want to add index sorting information.
Add index sort information
Sort Layout Settings
Column to Row (A1 -> B1...)
Row to Column (A1 -> A2...)
Sorting Target Well
Sort ID:
Sort ID 2
Sort Gate:
Color:
Sort Mode:
Ultra Purity
Stop Count:
1,200
Timeout:
570
(Seconds)
Add
Clear Well
Sort ID List
Sort ID
Sort Gate
Color
Sort Mode
Cell Size
Stop Count
Timeout
Add Well
Sort ID 1
G
Ultra Purity
Regular Cell
1,200
570
Remove
Close
All Events
Status: Ready
Elapsed Time:
00:00:00
Total Event:
Event Rate:
Start
Stop
Record
Restart
Sample Stop Condition
Detector & Threshold Settings
Sample Pressure:
Recording
Active Experiment
Measurement Settings
Compensation Settings
Compensation Panel
Tube - 1
Tube Information
Worksheet Settings
Stop & Sort Settings
Data Source - 1
Sort Control
Sort & Record Start
Method
96 Well Plate
Sort Settings
Droplet
Hardware Status
Emergency Stop
%Parent
%Total
Type here to search

If it is the first sort, go to completing Sort Settings slide.

If is not the first sort, go to next slide.

### Click the top entry in the `Sort ID List`.

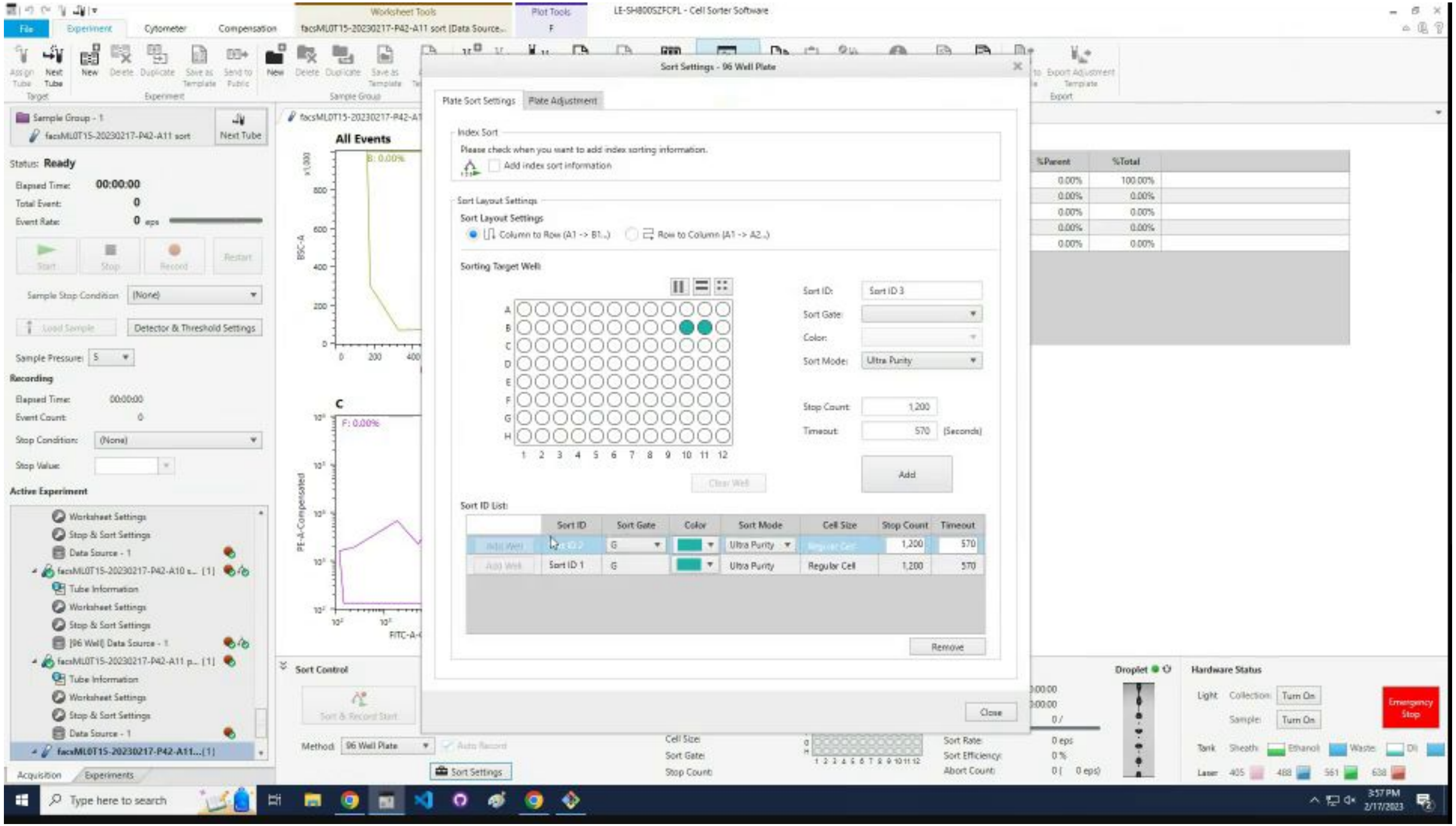

### Click `Remove`.

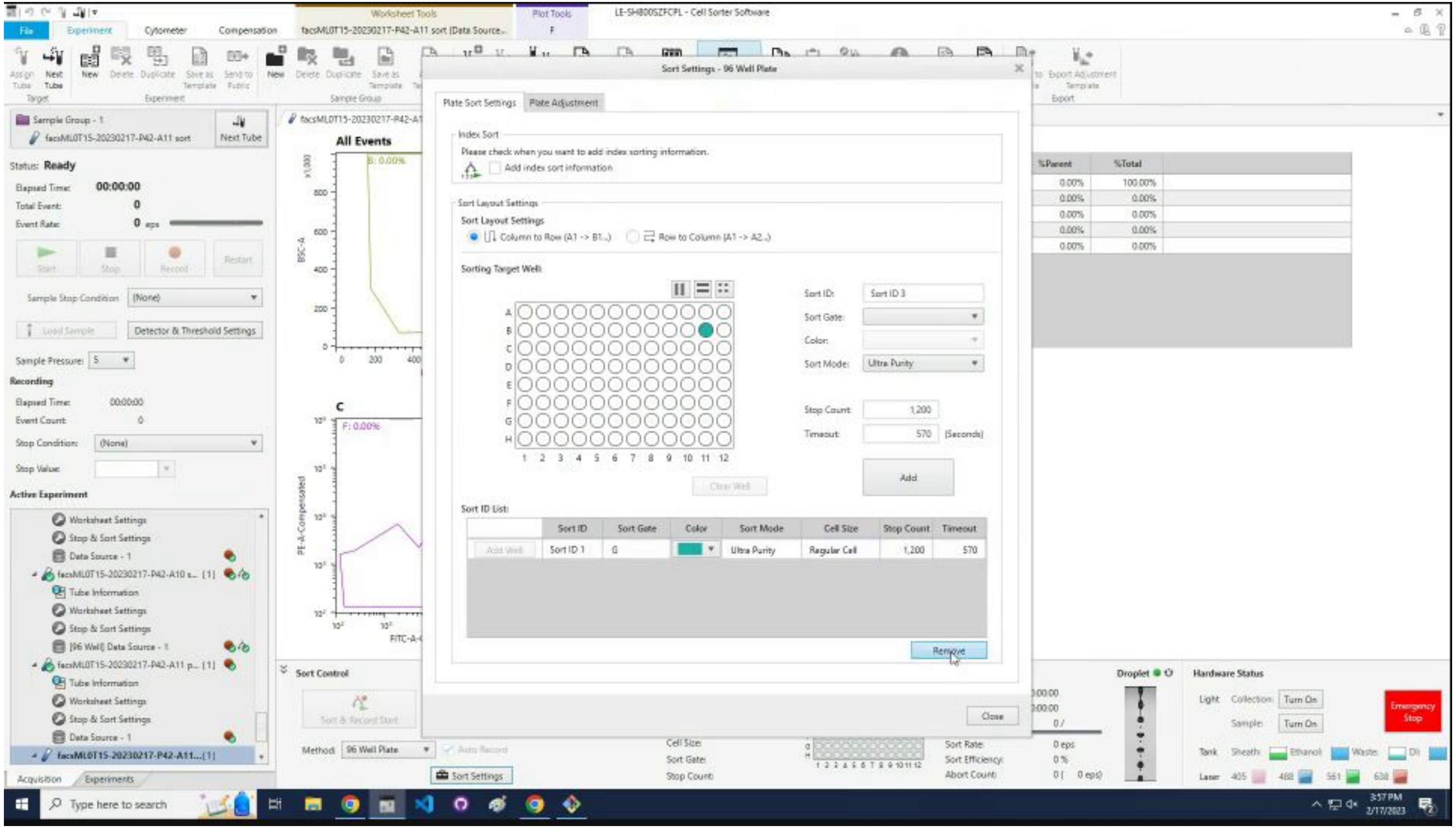

### Click `Close`.

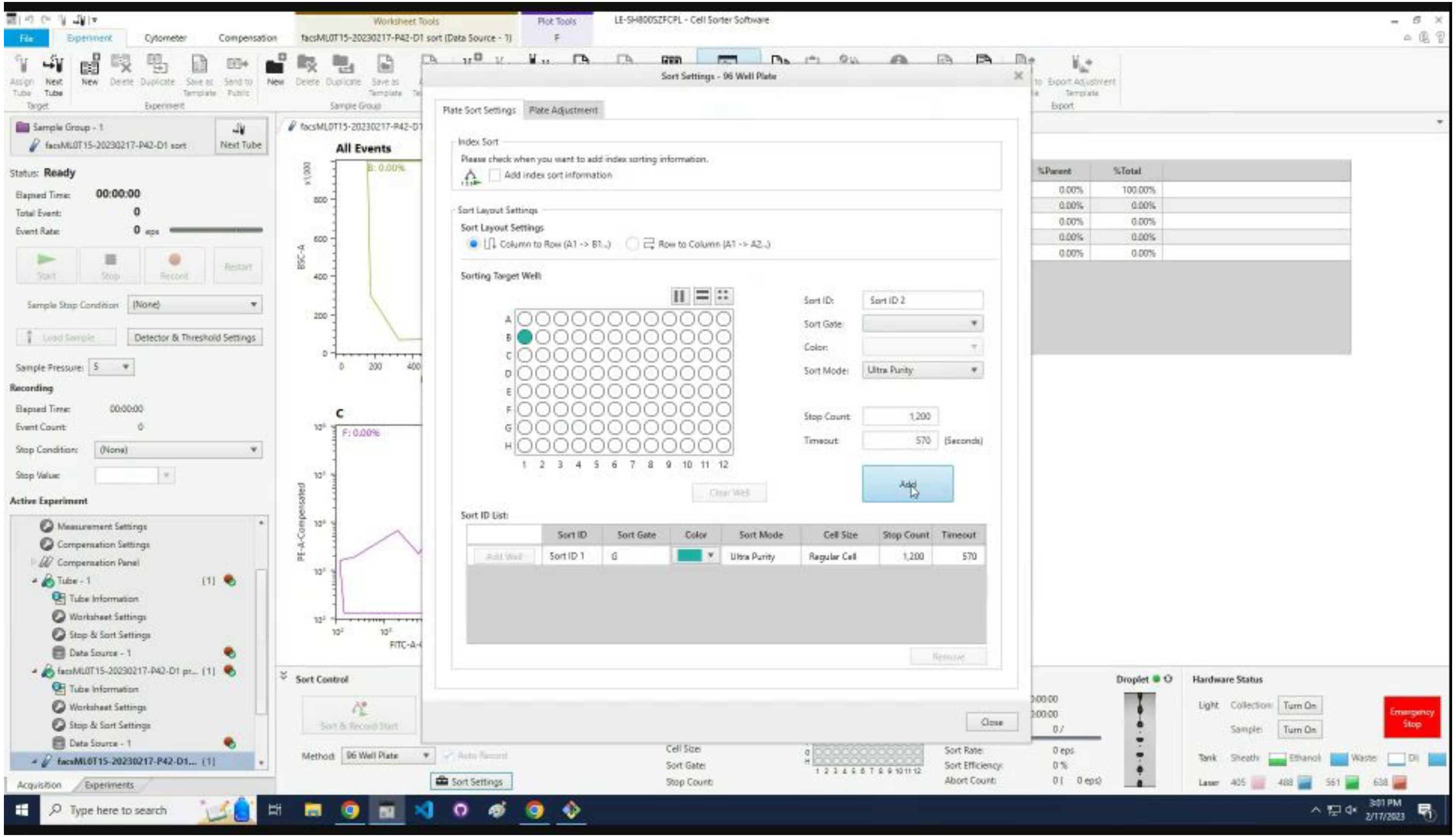
Sort Settings - 96 Well Plate
Plate Sort Settings
Plate Adjustment
Index Sort
Please check when you want to add index sorting information.
Add index sort information
Sort Layout Settings
Column to Row (A1 -> B1...)
Row to Column (A1 -> A2...)
Sorting Target Well:
Sort ID:
Sort ID 2
Sort Gate:
Color:
Sort Mode:
Ultra Purity
Stop Count:
1,200
Timeout:
570
(Seconds)
Add
Clear Well
Sort ID List:
Sort ID
Sort Gate
Color
Sort Mode
Cell Size
Stop Count
Timeout
Sort ID 1
G
Ultra Purity
Regular Cell
1,200
570
Remove
Close

### Click `Load Collection.`

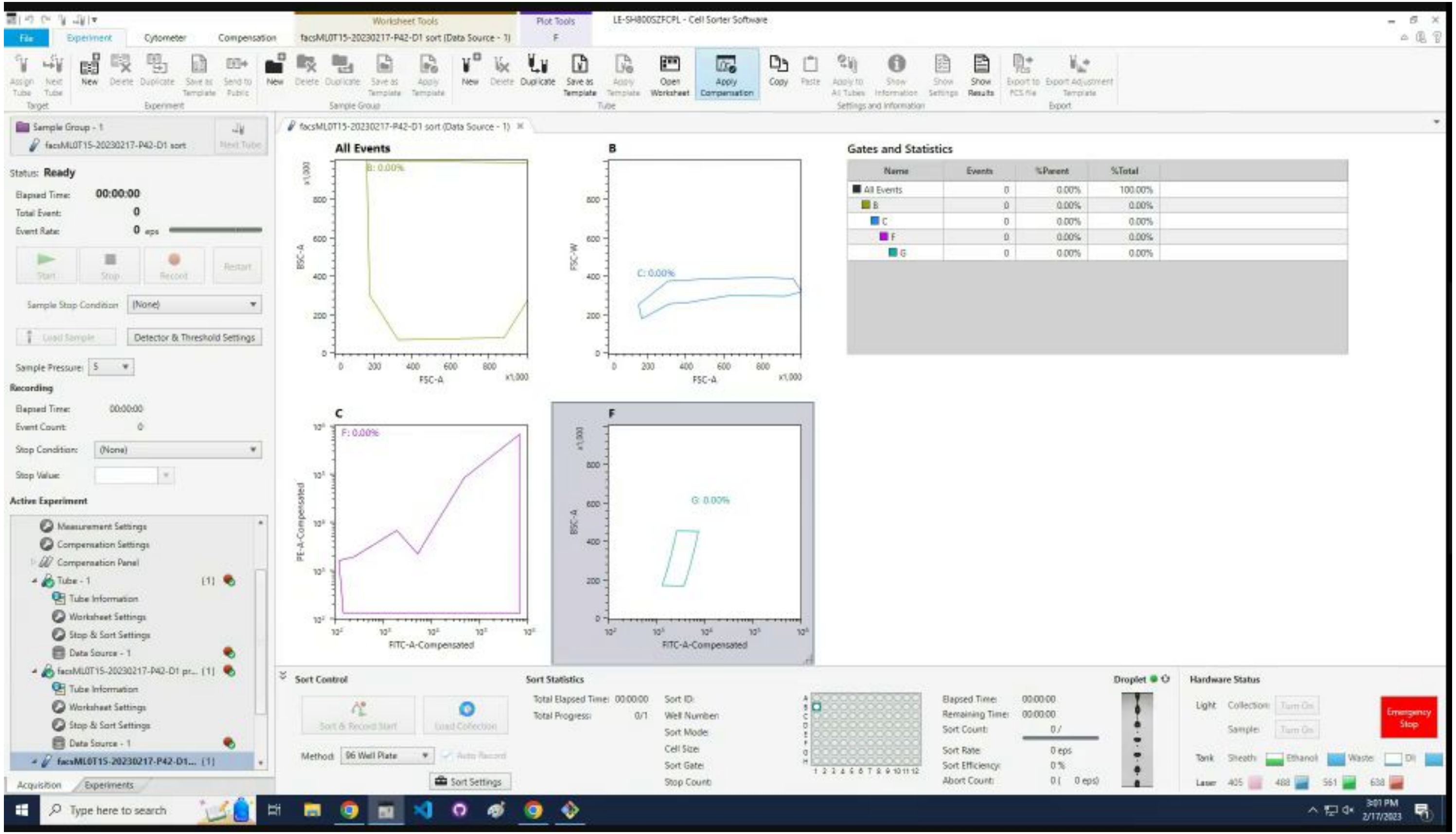

### Click `Start`.

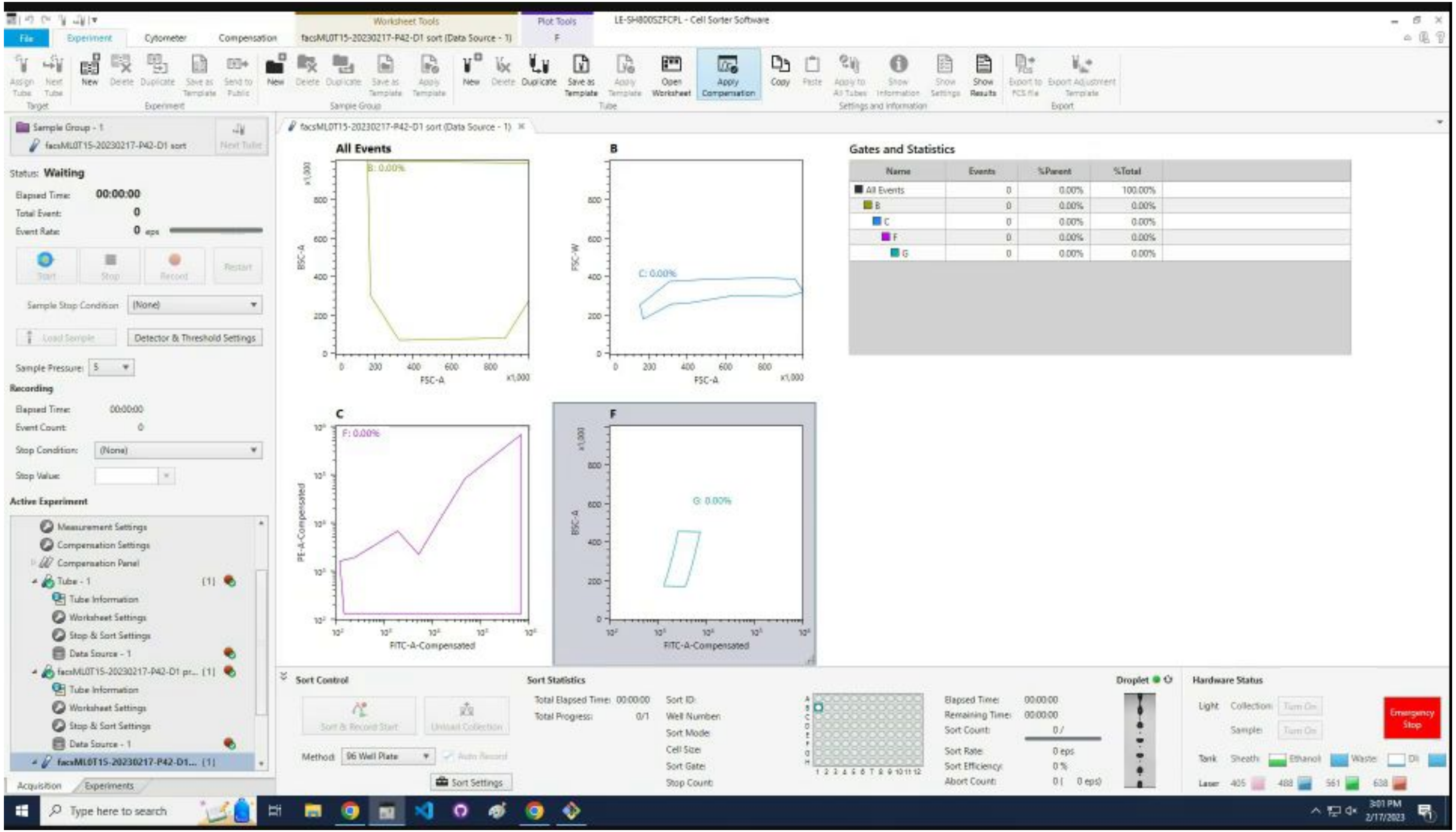
LE-SH800SZFCPL - Cell Sorter Software
Sample Group - 1
Status: Waiting
Elapsed Time: 00:00:00
Total Event: 0
Event Rate: 0
Start
Stop
Record
Restart
Sample Stop Condition
Detector & Threshold Settings
Sample Pressure: 5
Recording
Active Experiment
All Events
B
C
F
Gates and Statistics
Name
Events
%Parent
%Total
Sort Control
Sort Statistics
Droplet
Hardware Status
Emergency Stop

### Wait ~20 seconds for cells to appear.

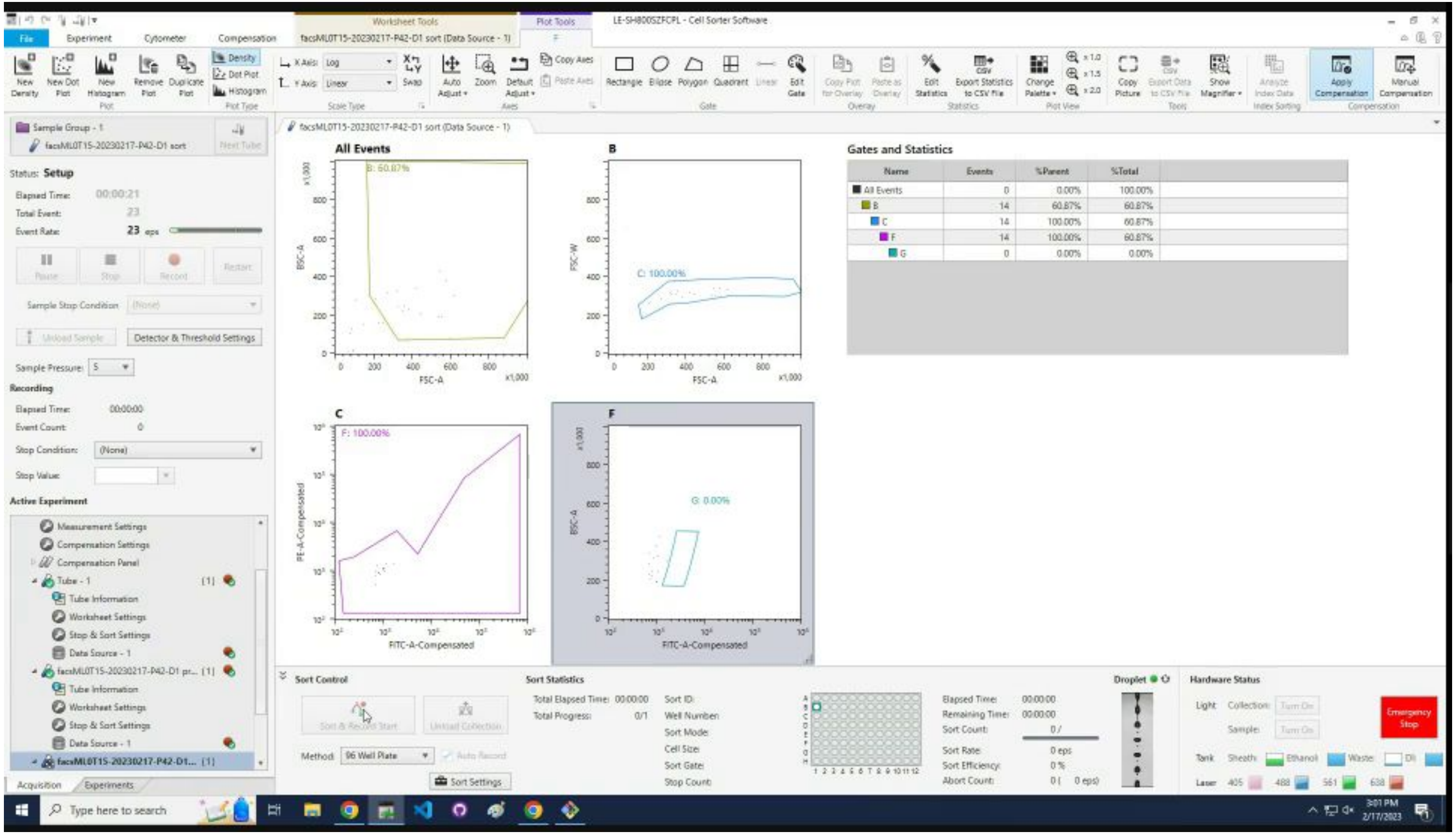

### Click `Sort & Record Start.`

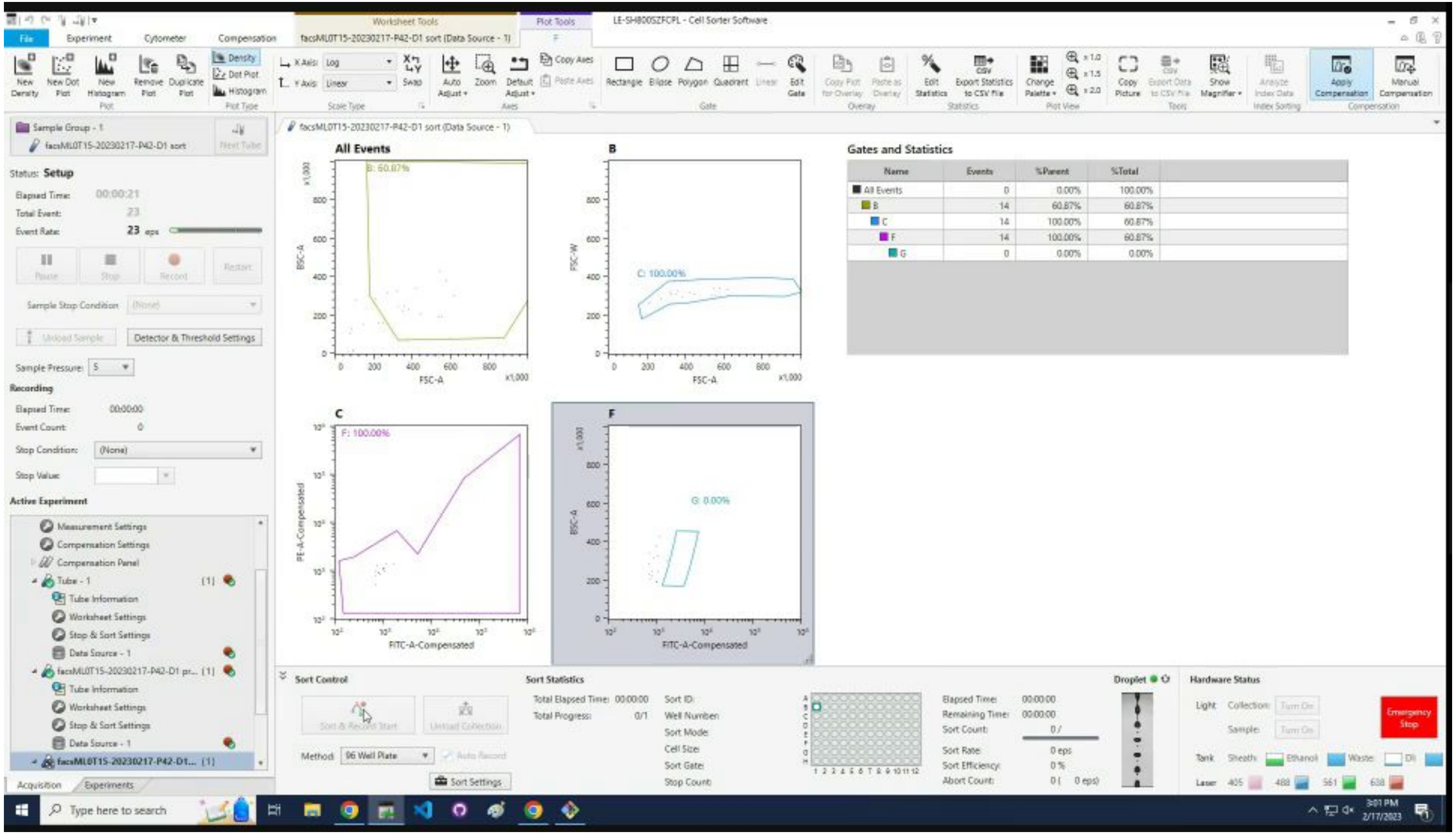

### Wait for the sort to finish.

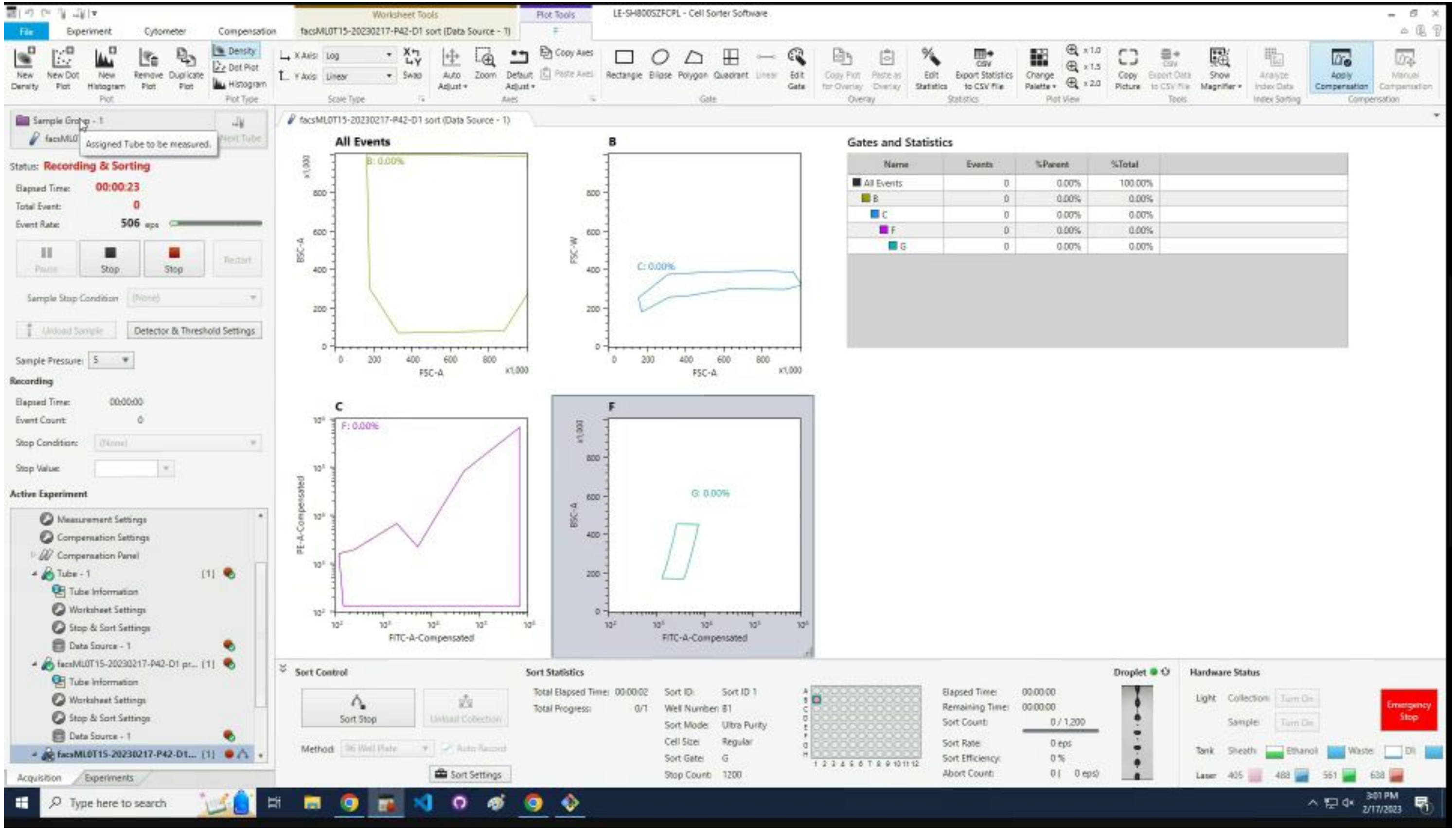

Click `Finish`.

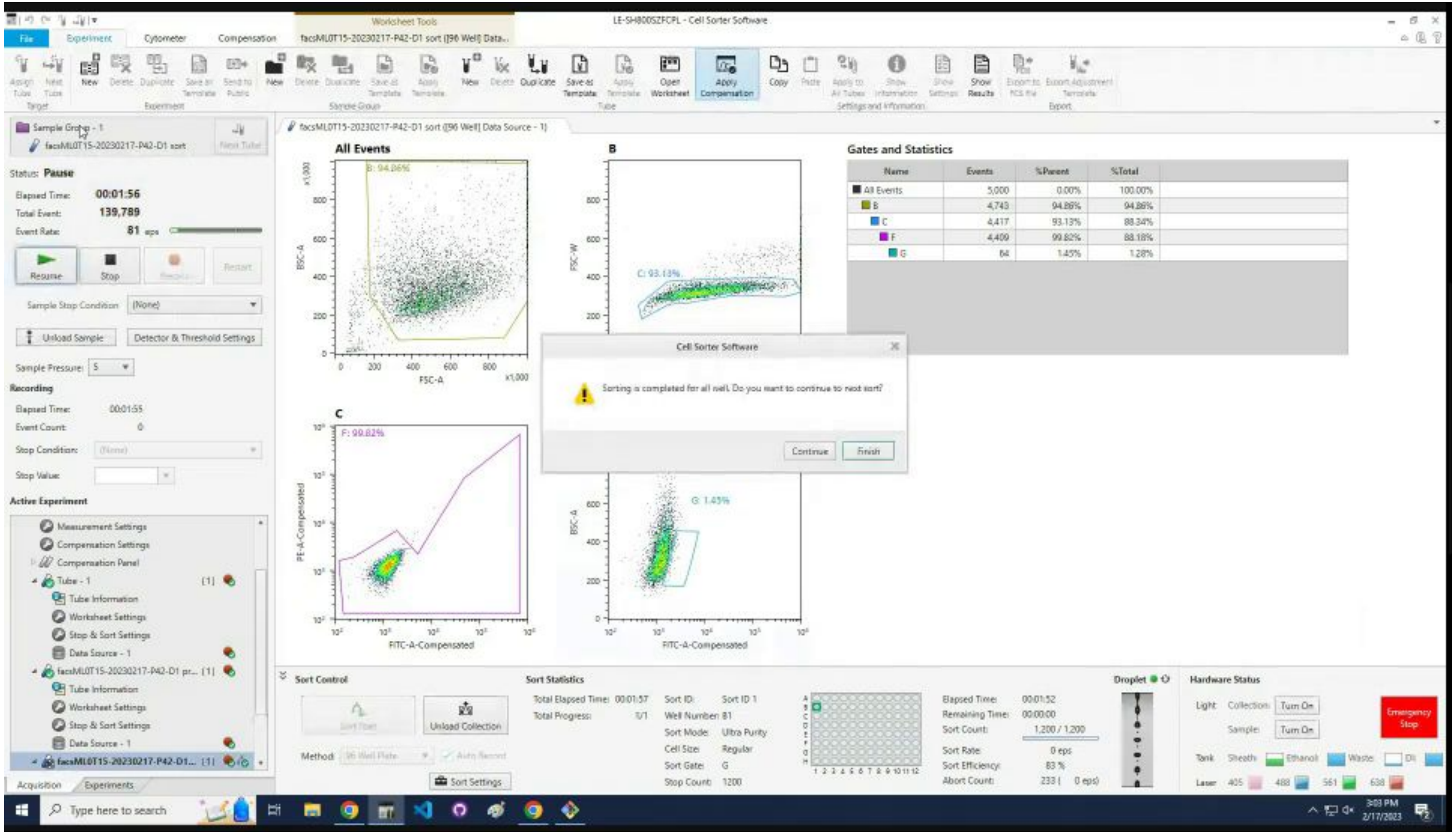

### Click the X to close the tube tab.

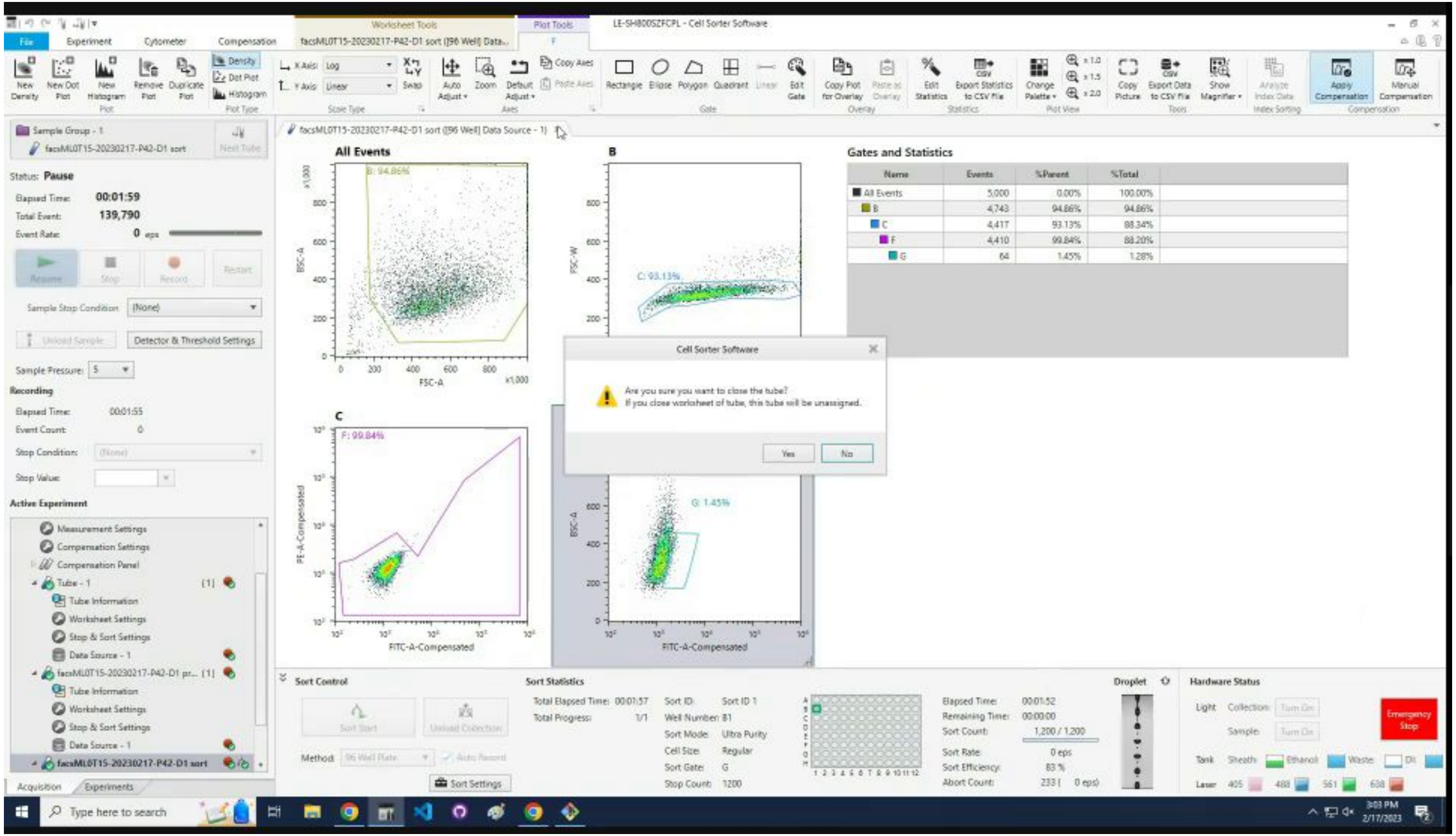

### Click `Yes` to confirm closing the tube tab.

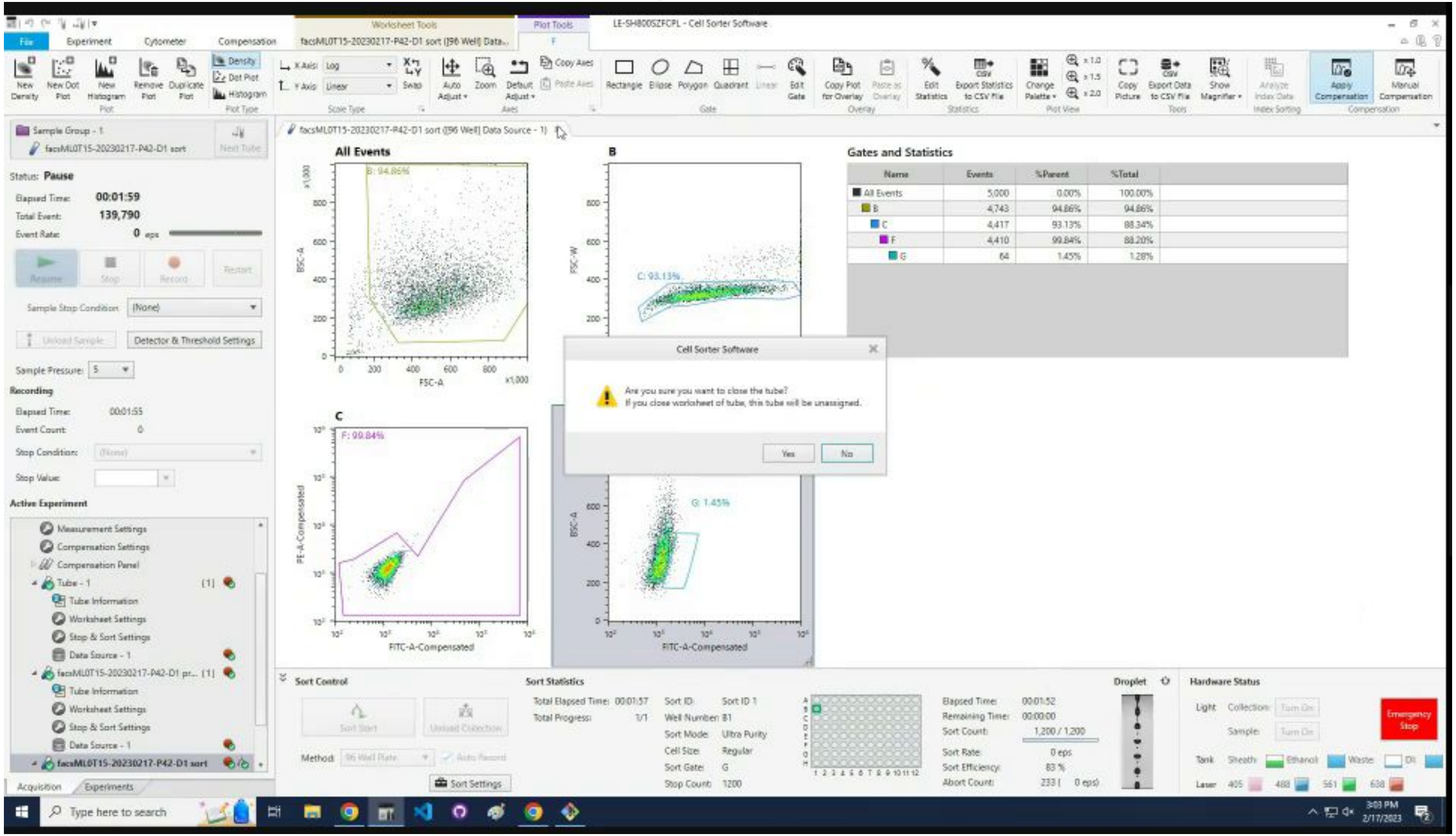

### Wait for the Sony to refresh the GUI after `Probe Wash`.

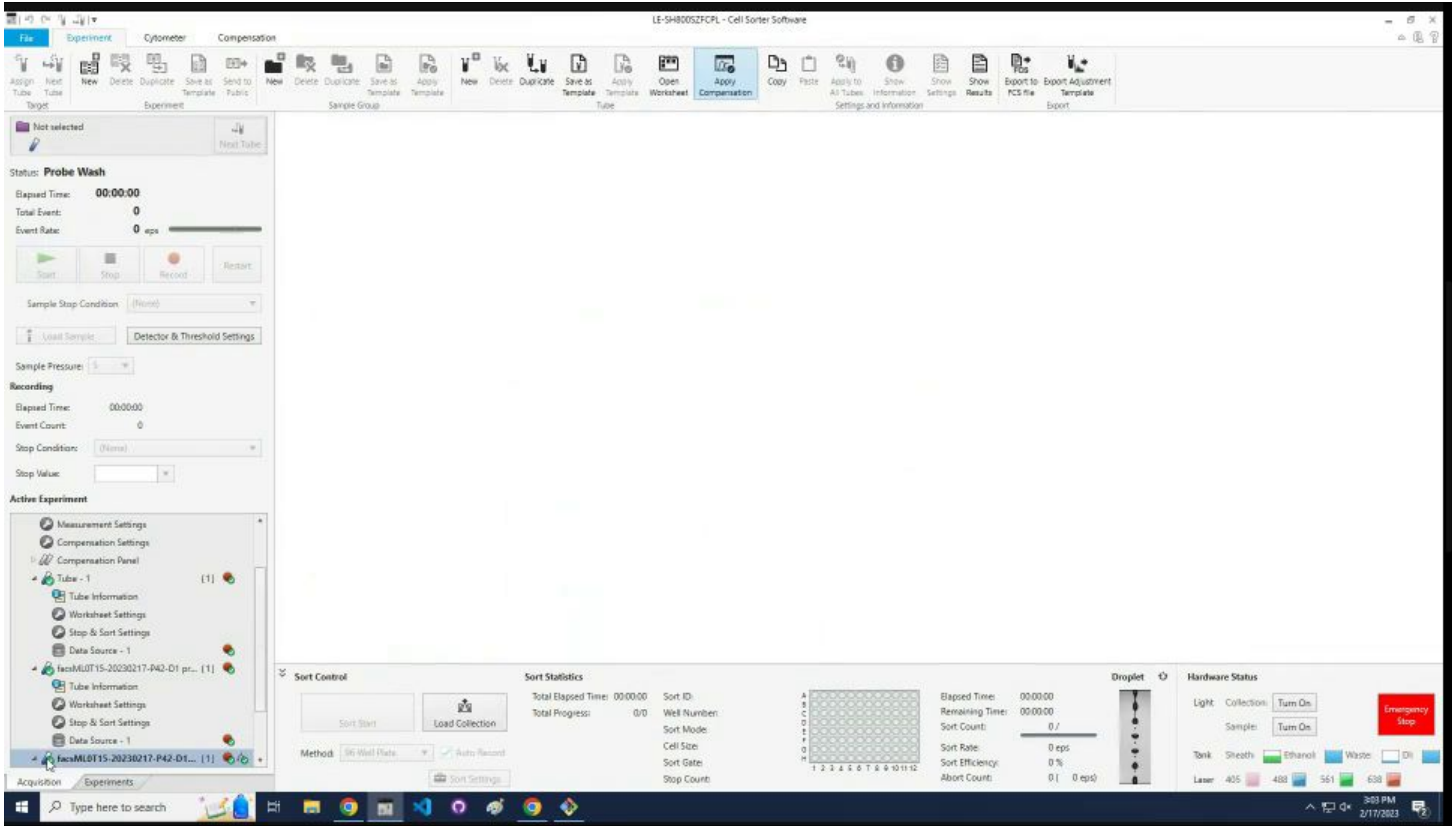

### Click on the finished sample tube.

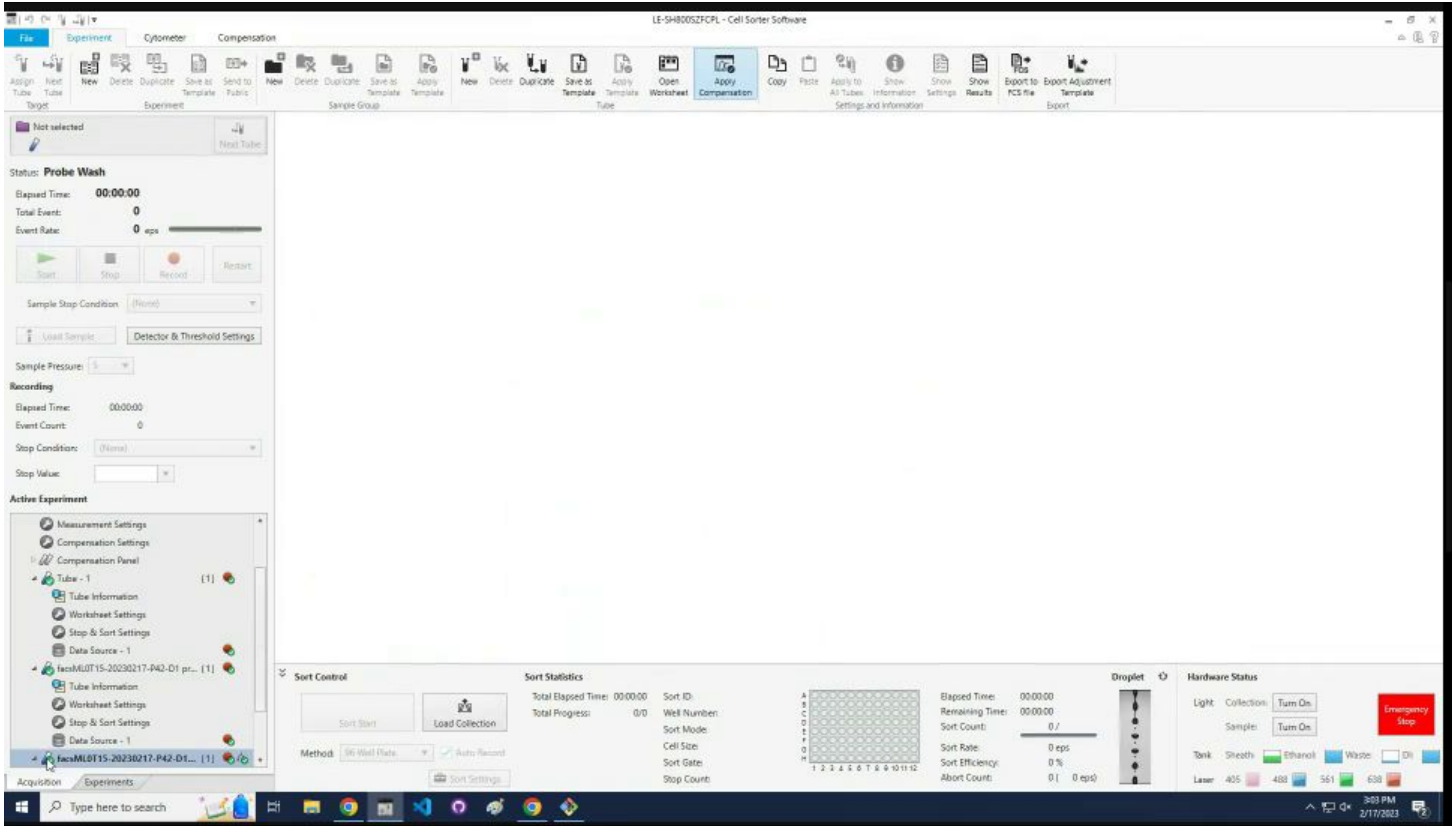

### Click the `Duplicate` icon.

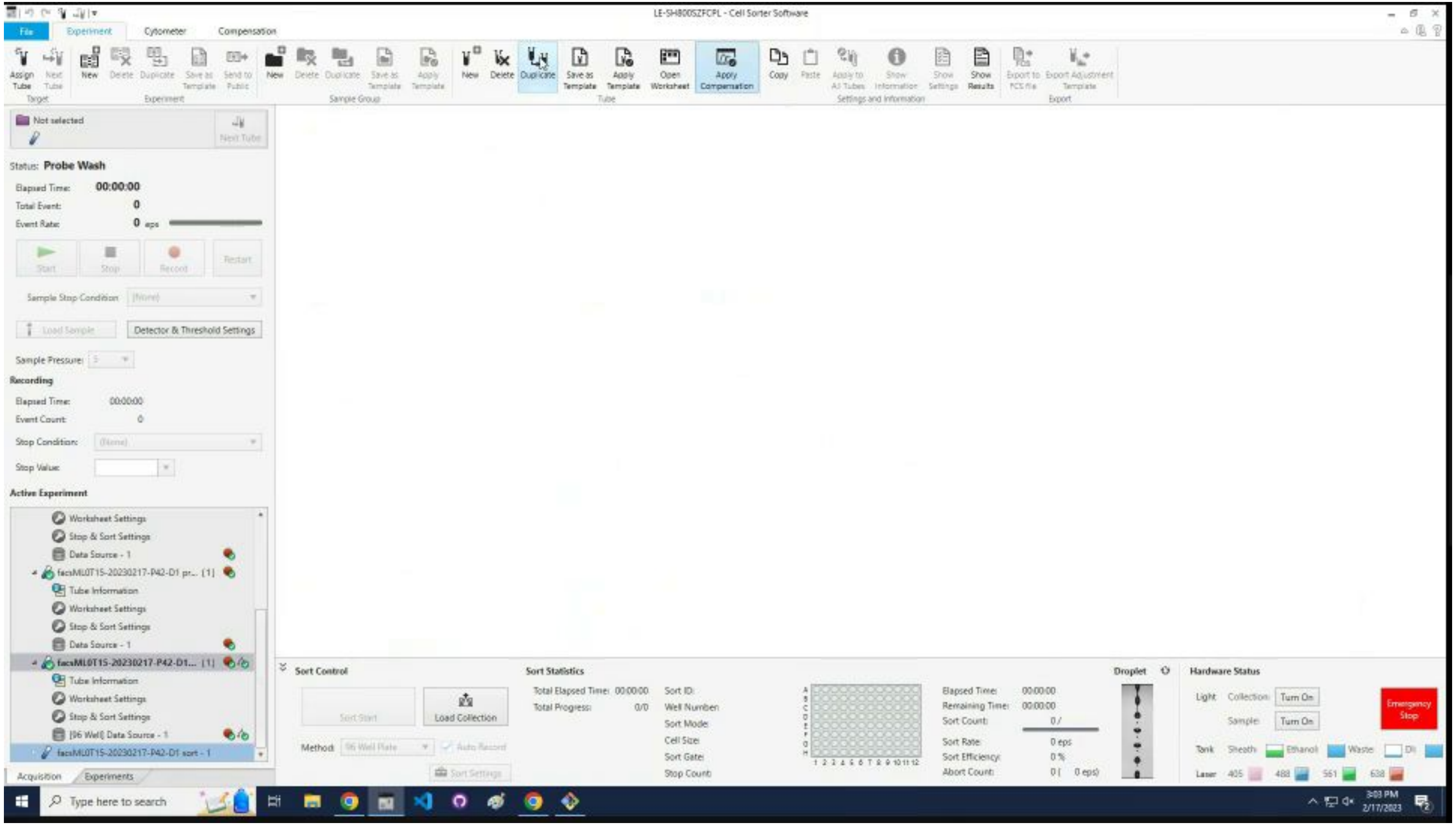

### Click the `Assign Tube` icon.

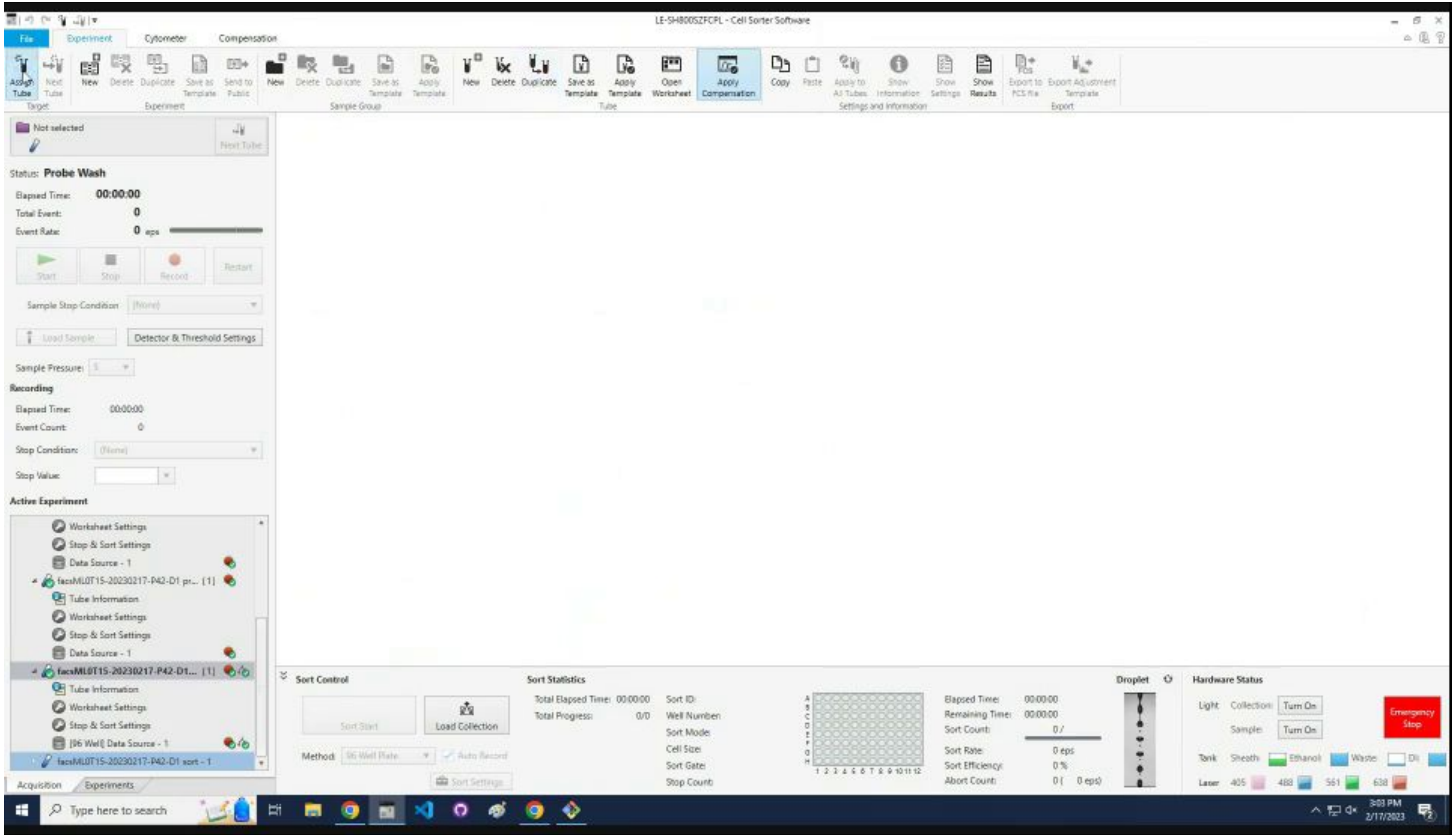

Repeat Iterate through samples section for more samples.

### Shutdown

### Answer `y/n` to close solenoid. (`No`, if sorting again.)

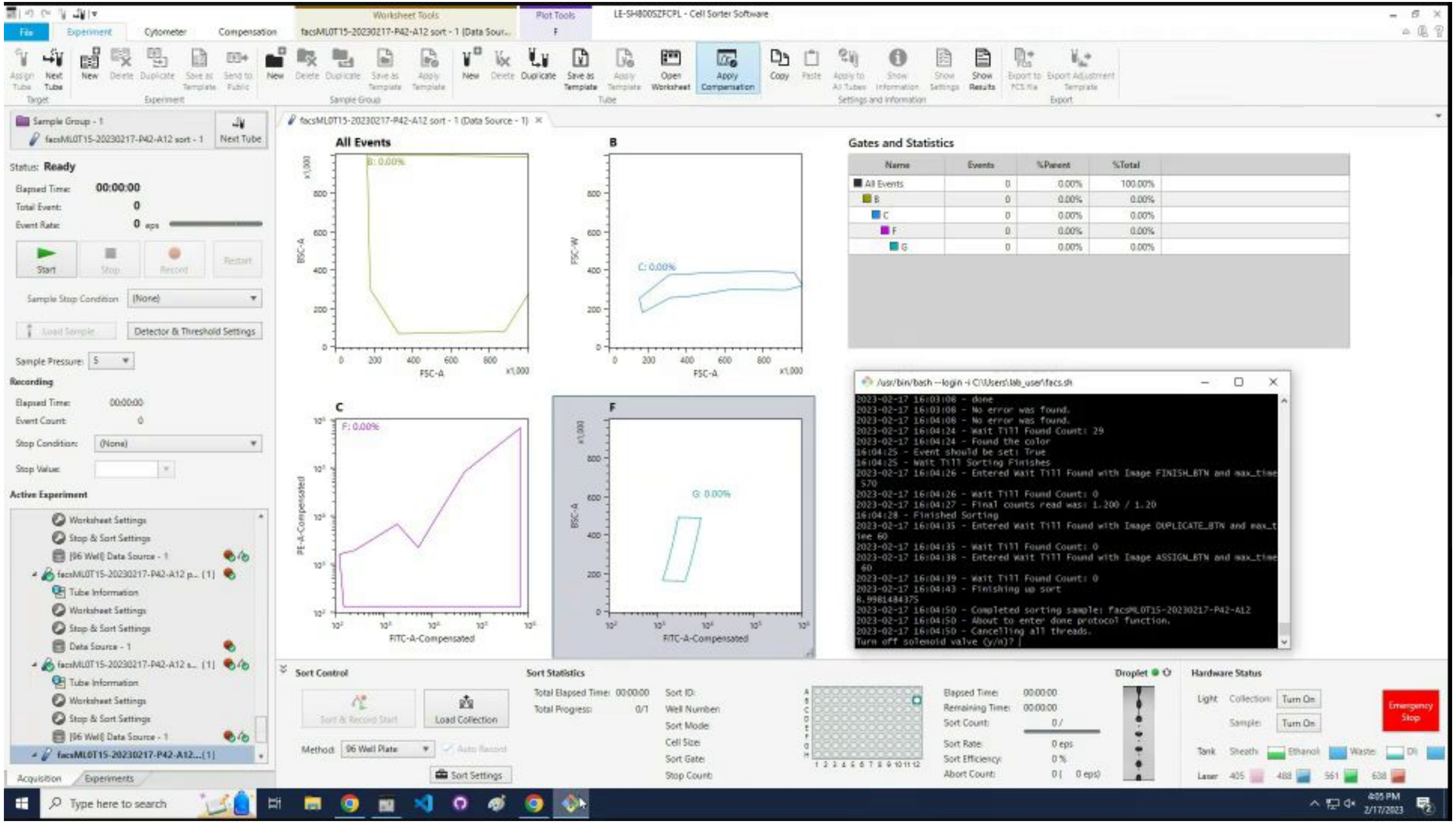

### An active blue tube remains for the next sorting run.

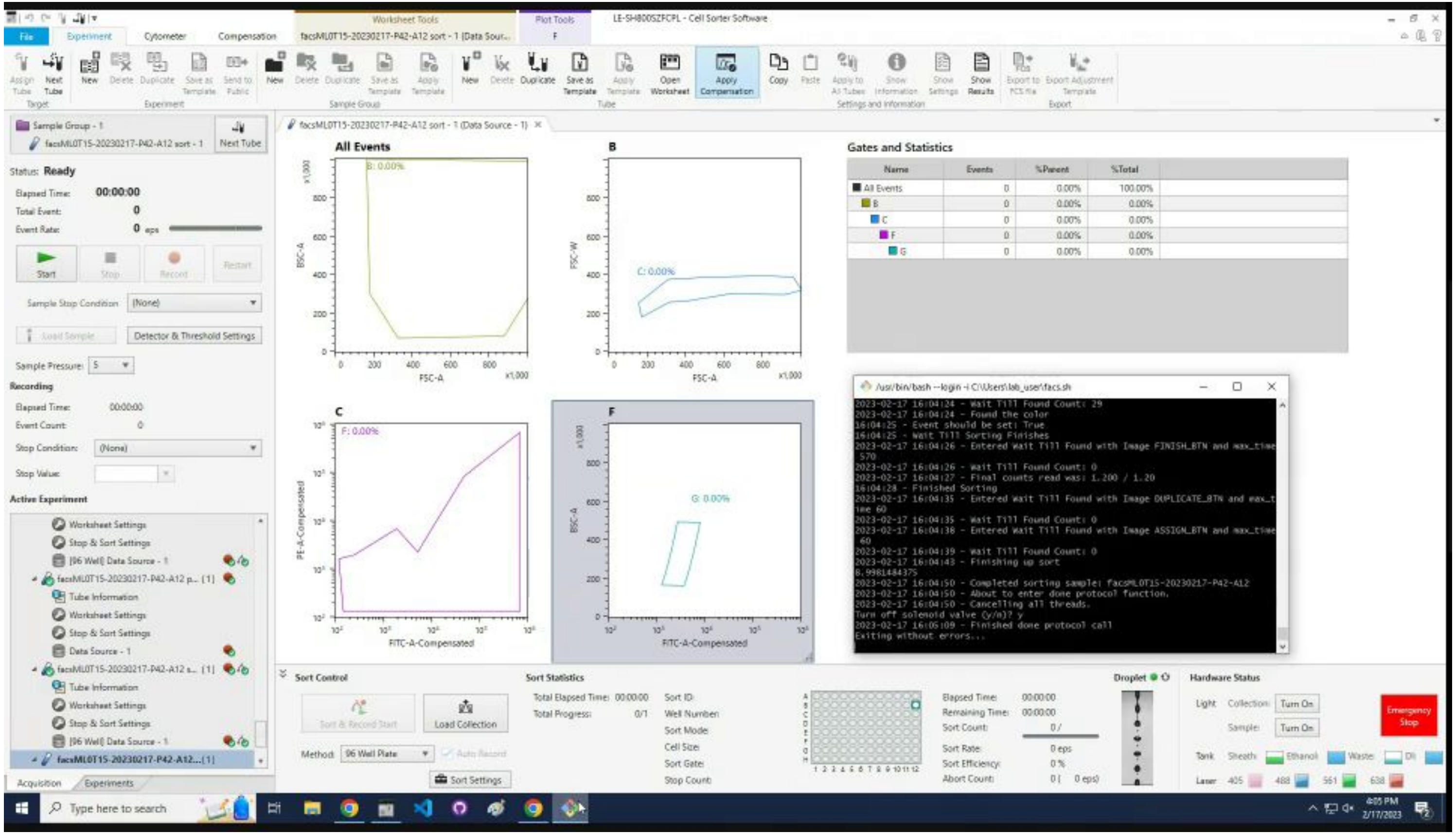

### Data Scraping

### Start the automated data scraping by clicking `scraper`.

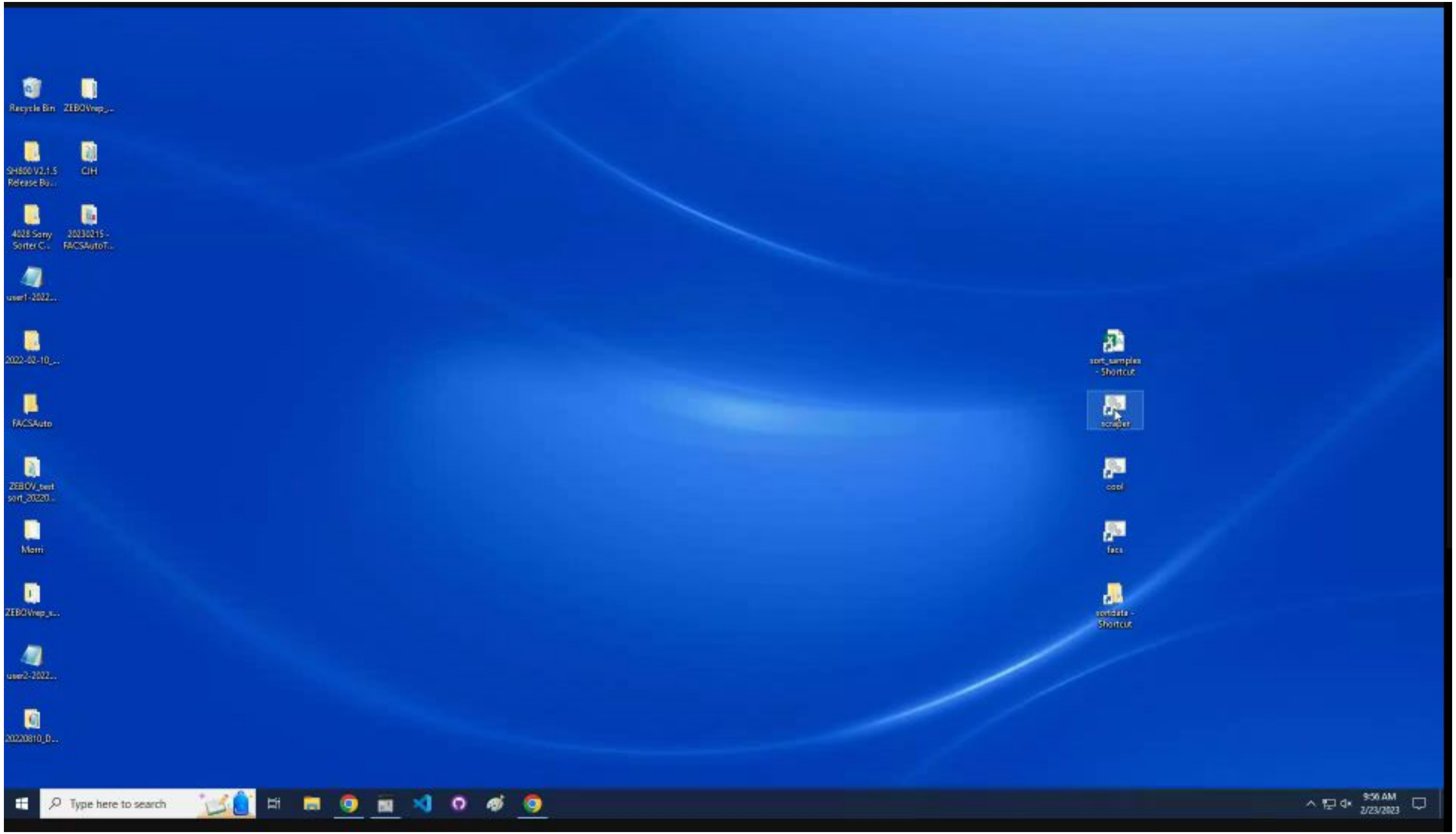

Enter `y/n` to export data from the Cell Sorter software.

If `yes`, skip to export.

If `no`, proceed to next slide.

### Enter `n` on startup, and skip to [plotting].

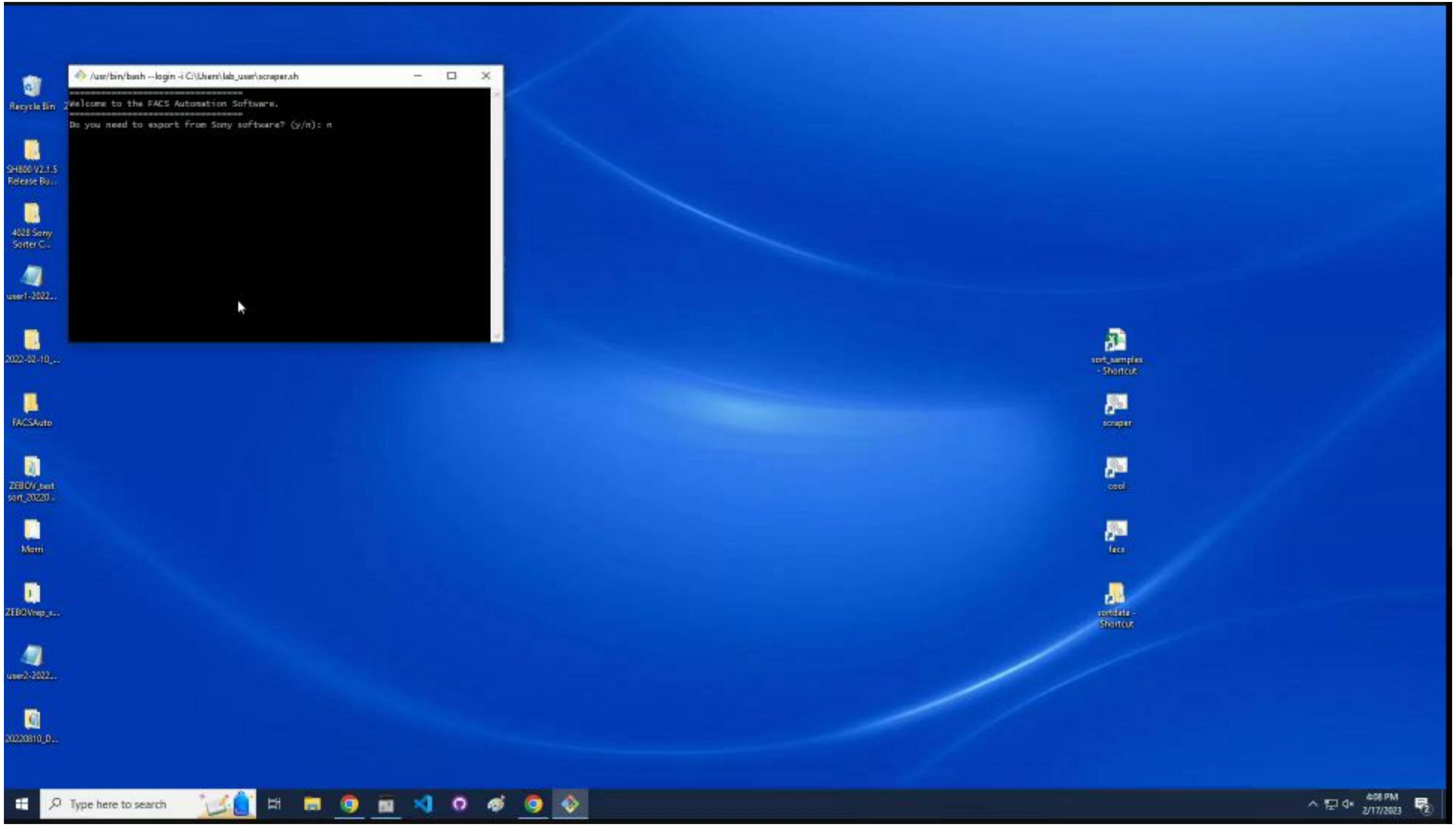

### Align the experiment to the top.

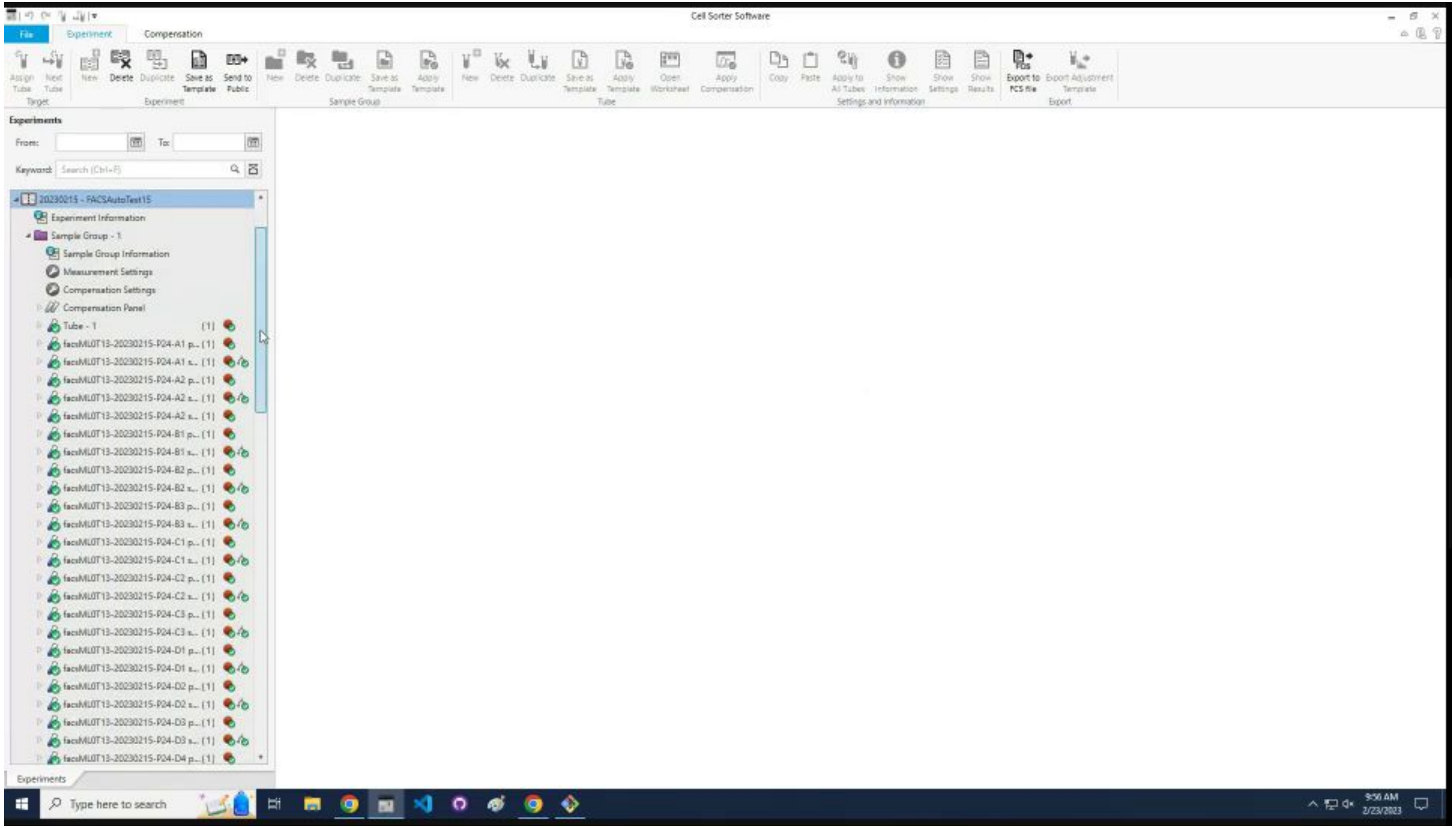

### Enter `y` to start the exports.

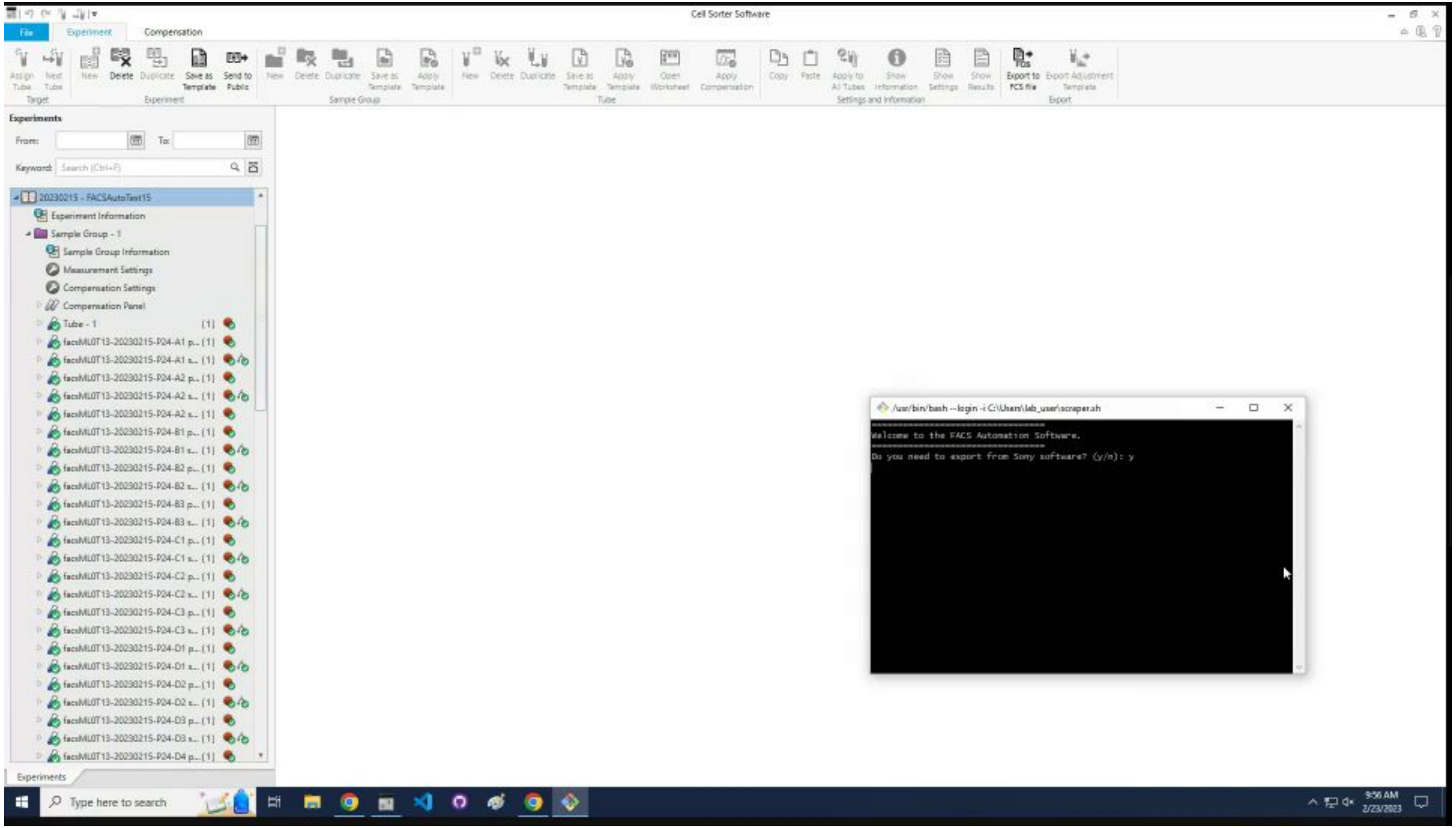

### The program finds the sorted samples.

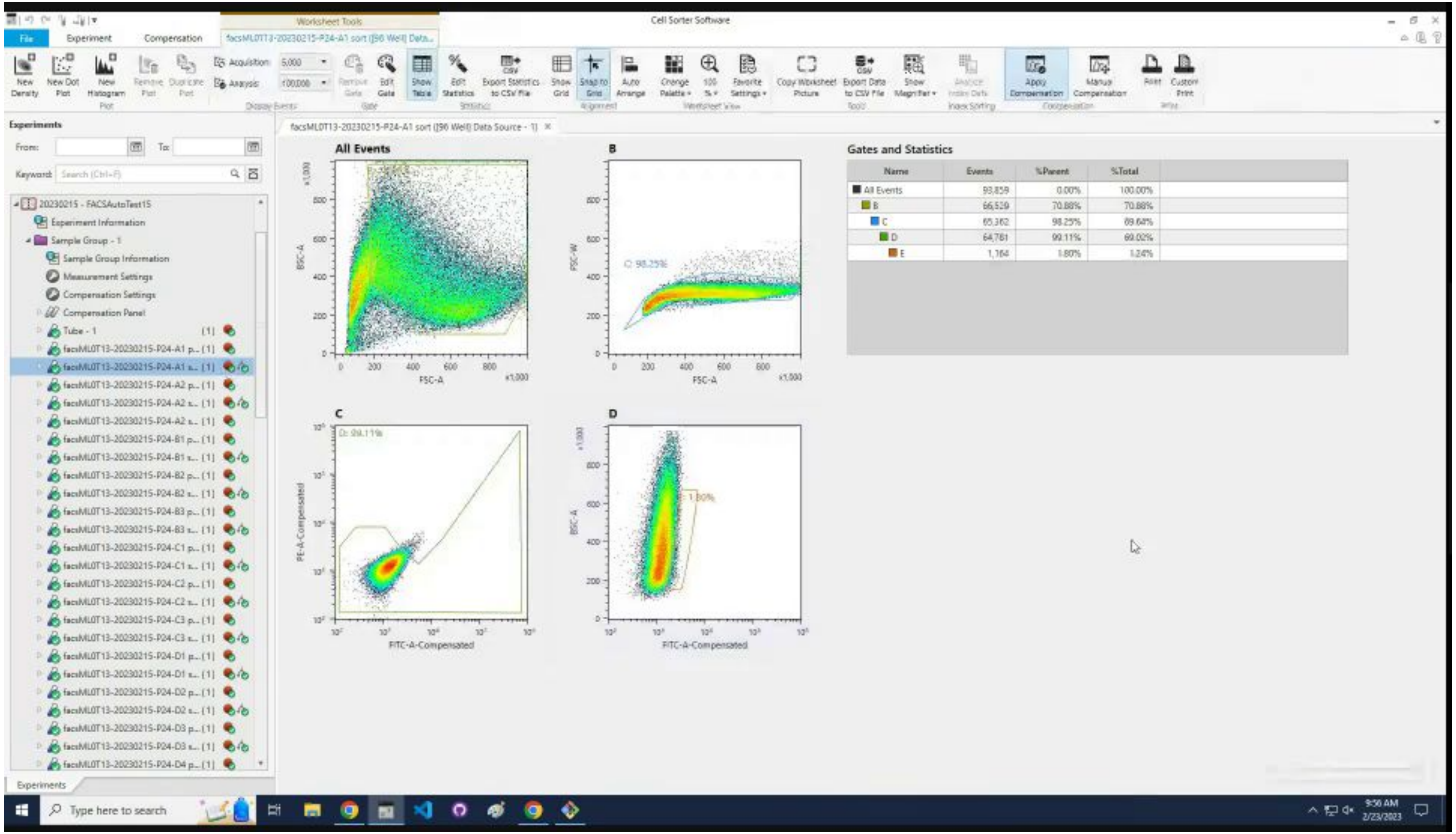


| Name | Events | %Parent | %Total |
|---|---|---|---|
| All Events | 93,859 | 0.00% | 100.00% |
| B | 66,529 | 70.88% | 70.88% |
| C | 65,362 | 98.25% | 69.64% |
| D | 64,761 | 99.11% | 69.02% |
| E | 1,164 | 1.80% | 1.24% |

### The filename is updated and the data is exported.

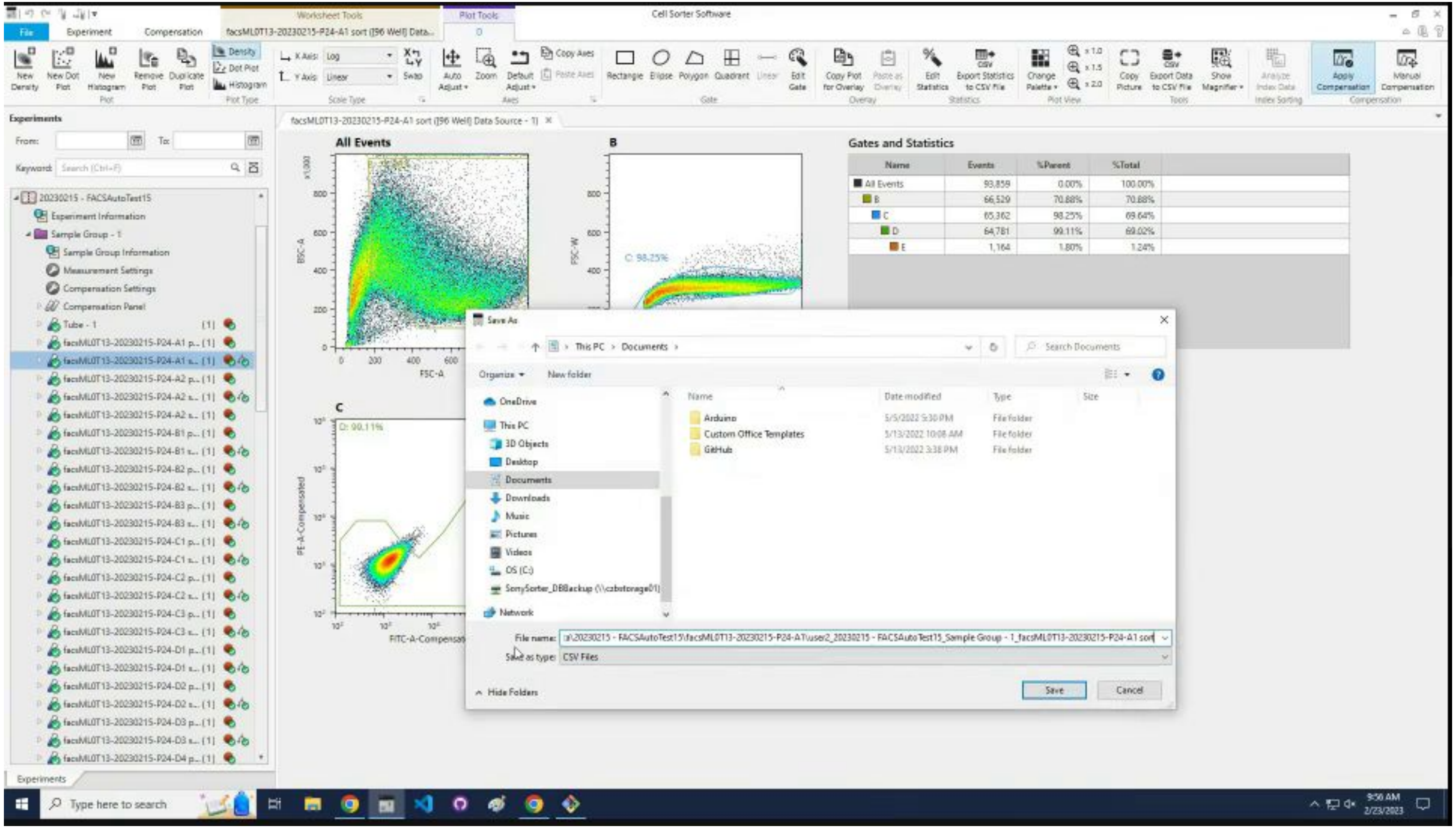

### The program scrolls through the full list of experiments.

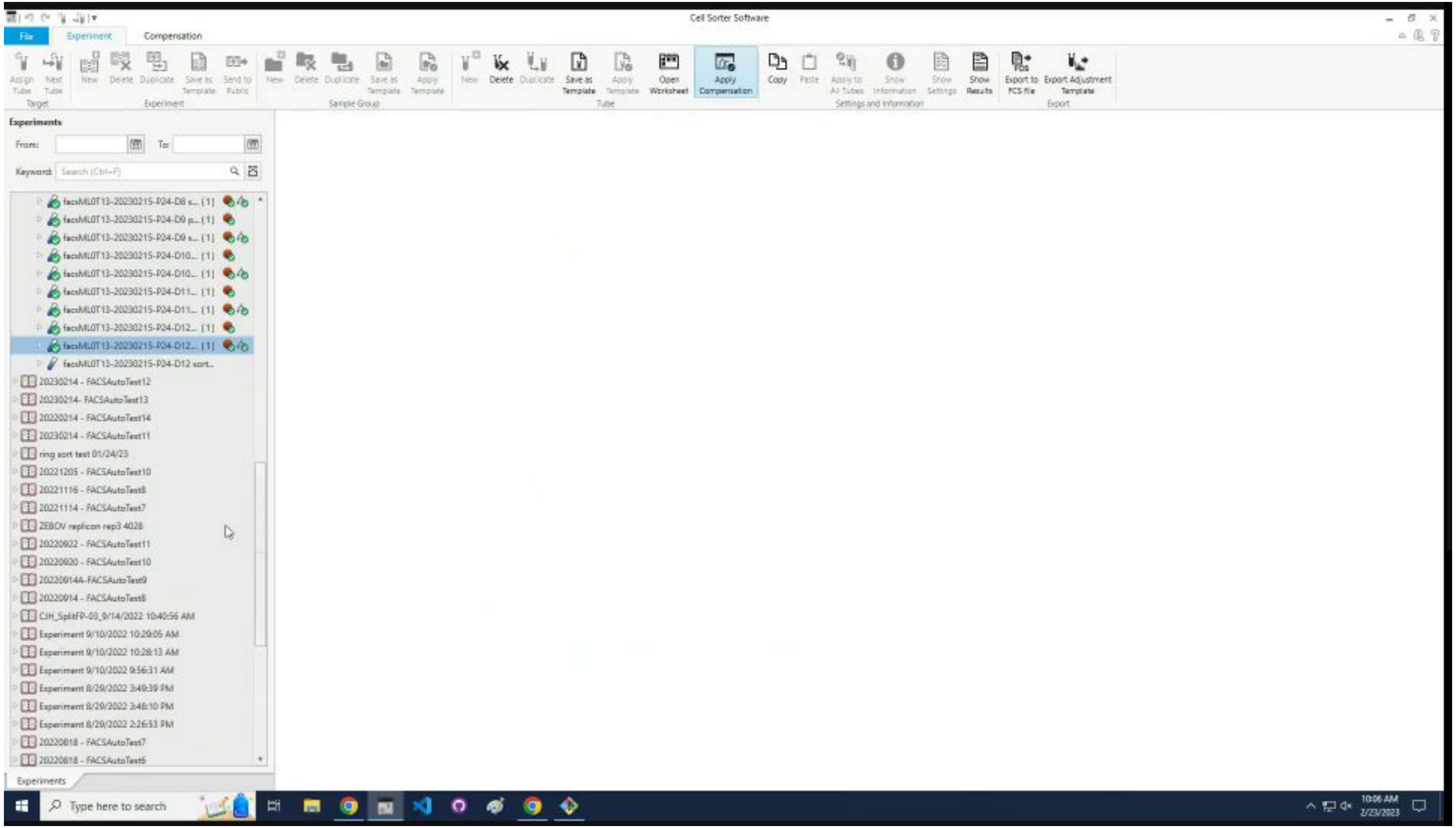

### The program asks for the folder with the data.

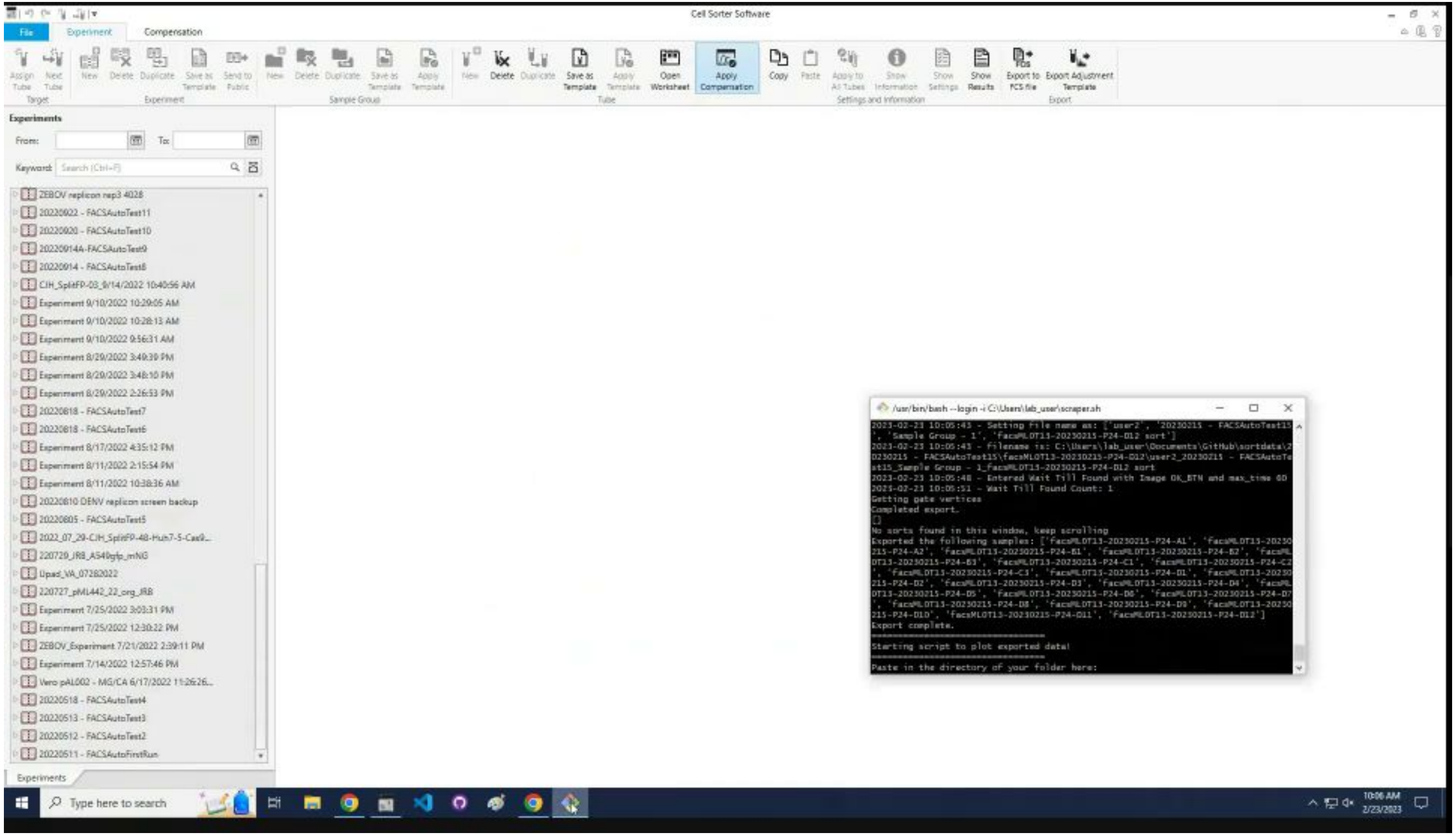

### Navigate to the `sortdata` folder.

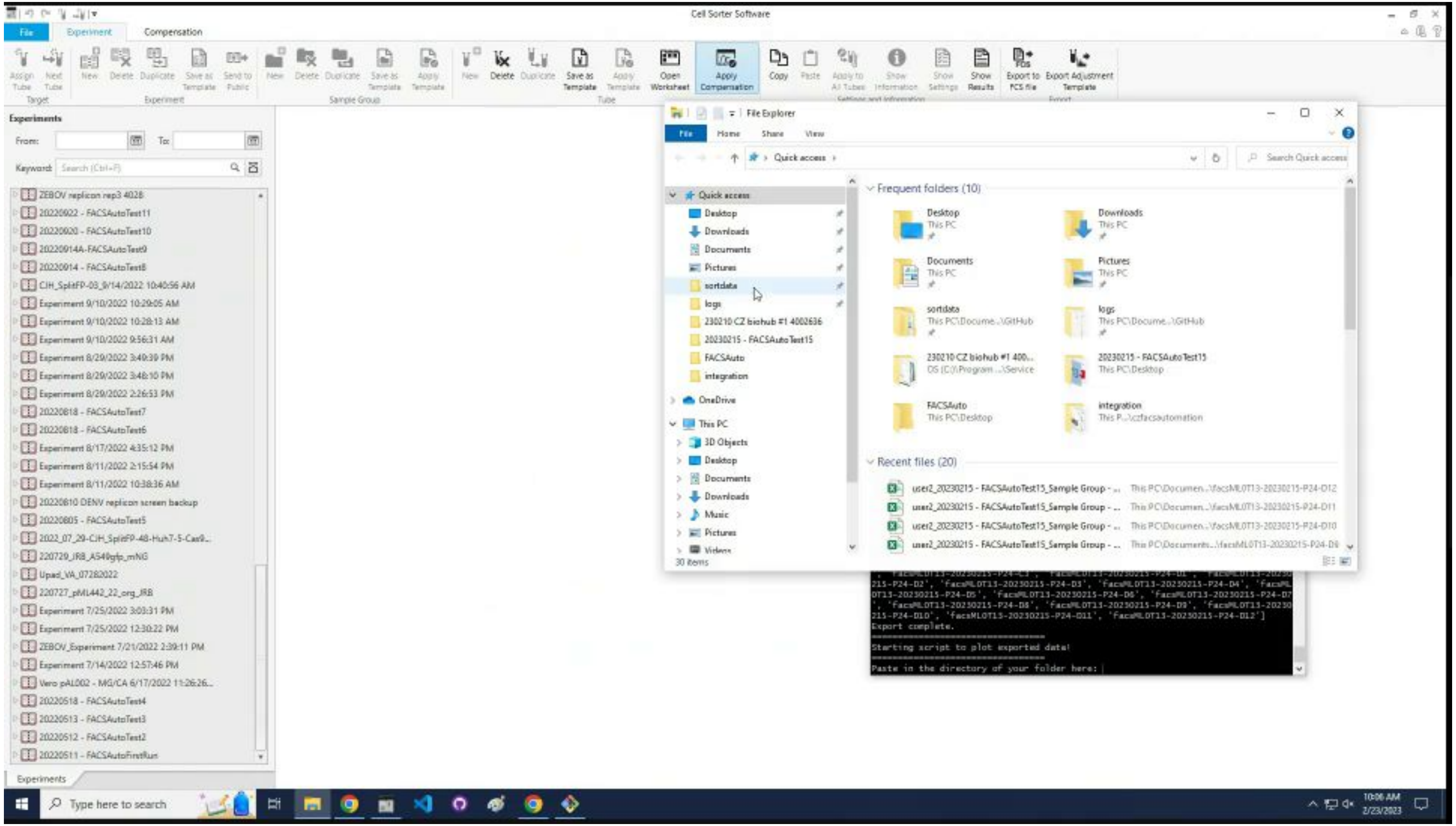

### Copy the file path for the folder containing the data.

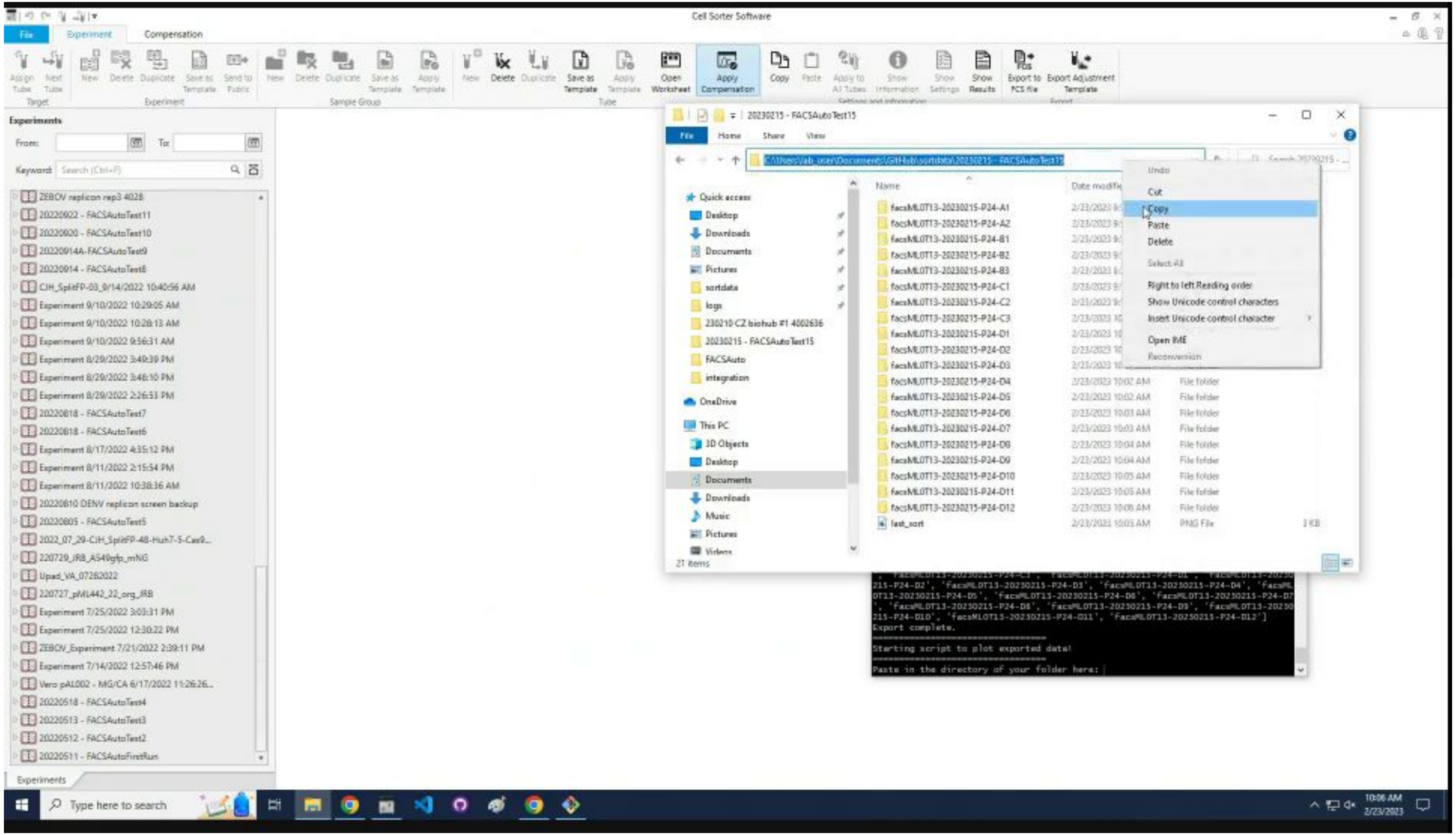

### Use `Right Click + Paste + Enter` to paste it.

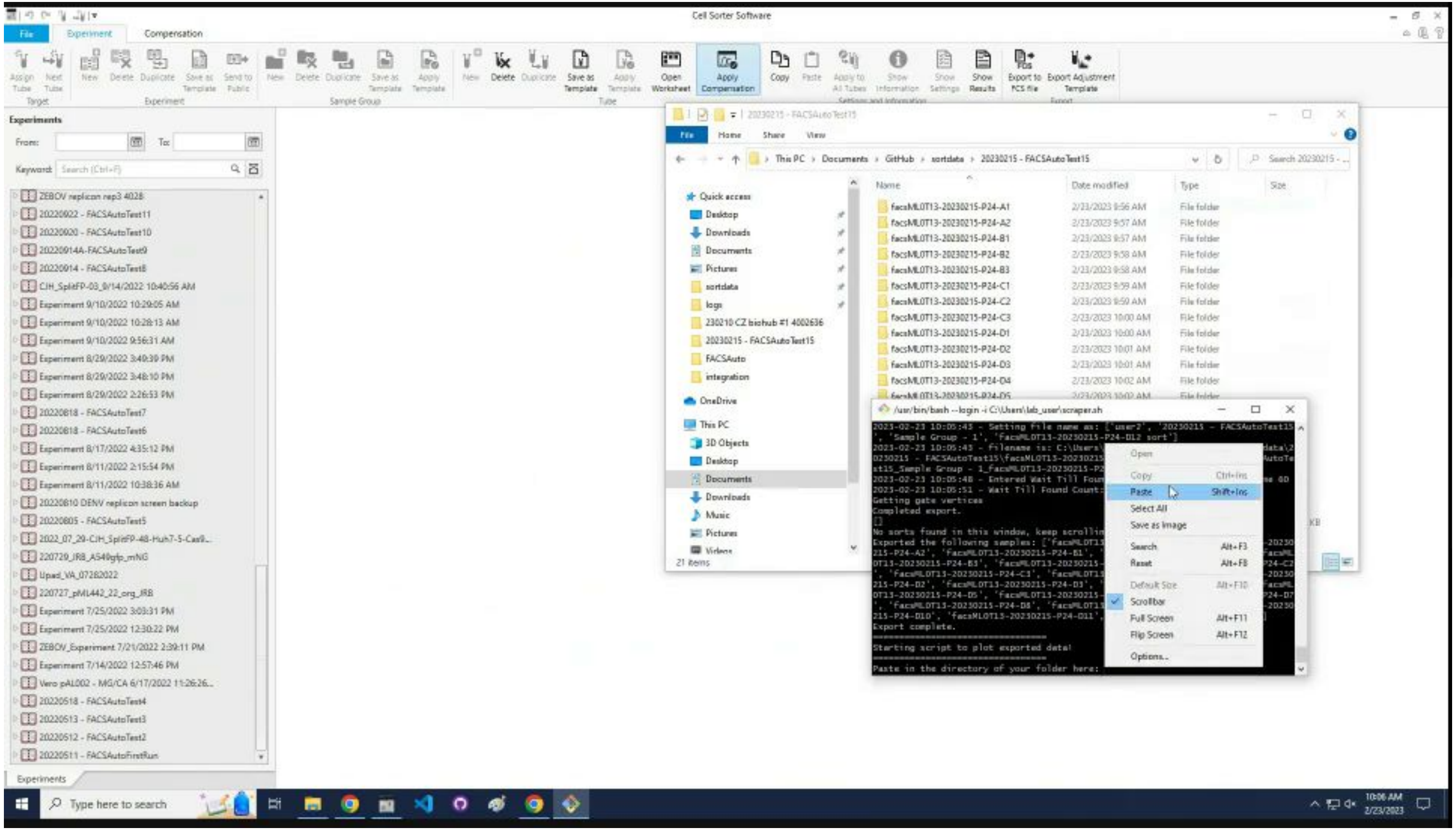

### Select which data to plot, `Enter` for all of it.

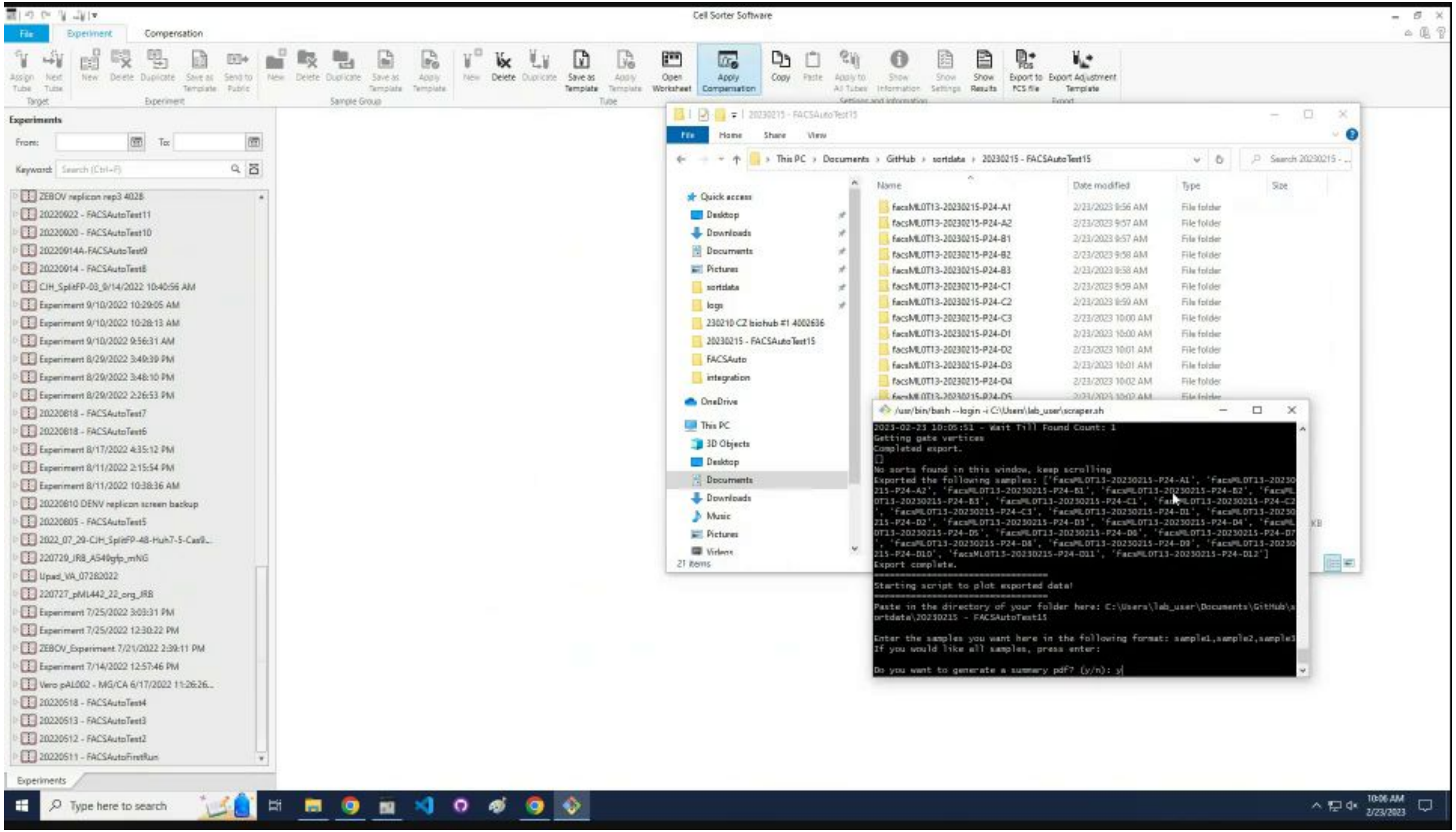

### Select `y/n` to generate a summary PDF.

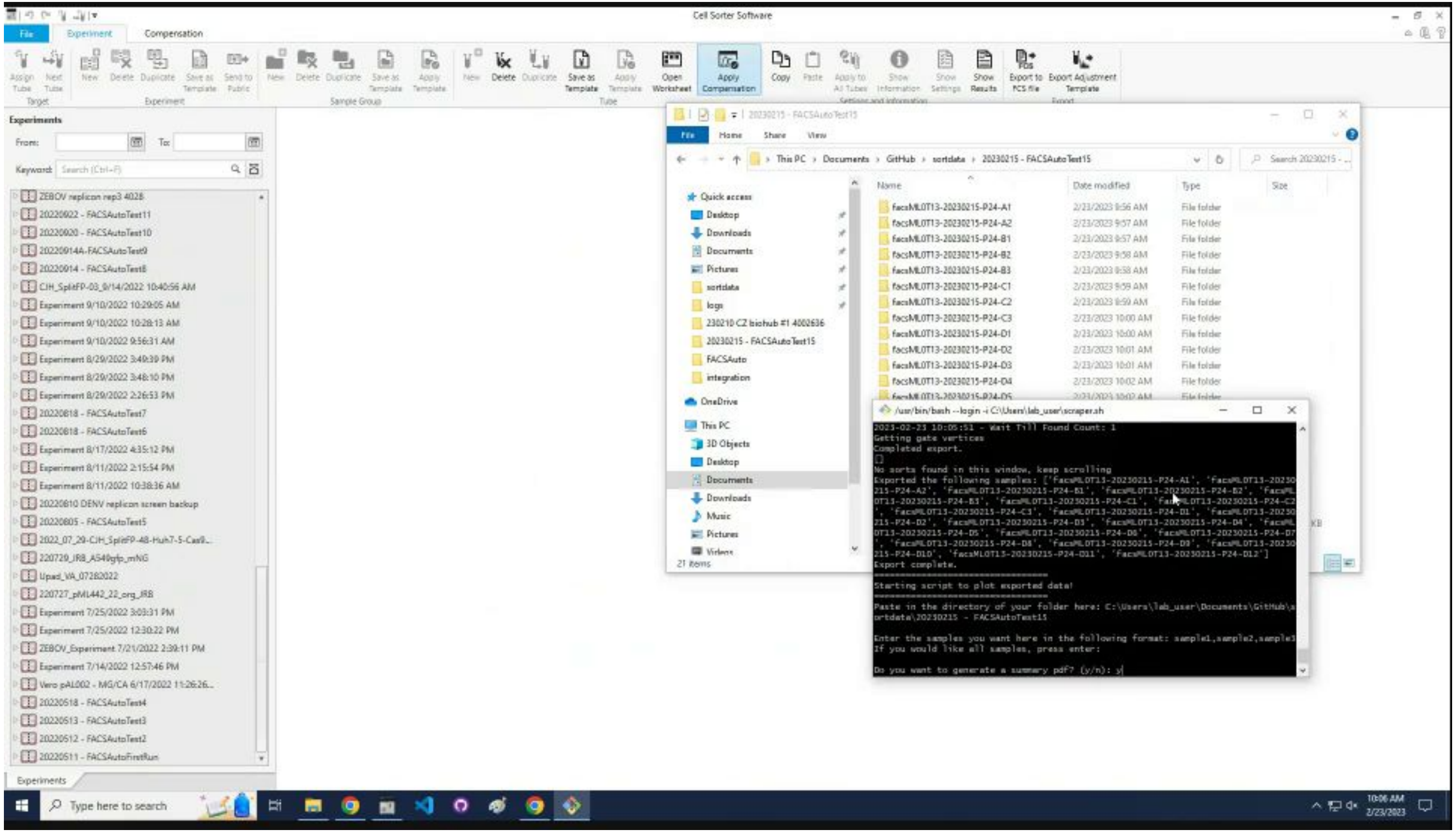

### The summary PDF is in the folder; the program exits.

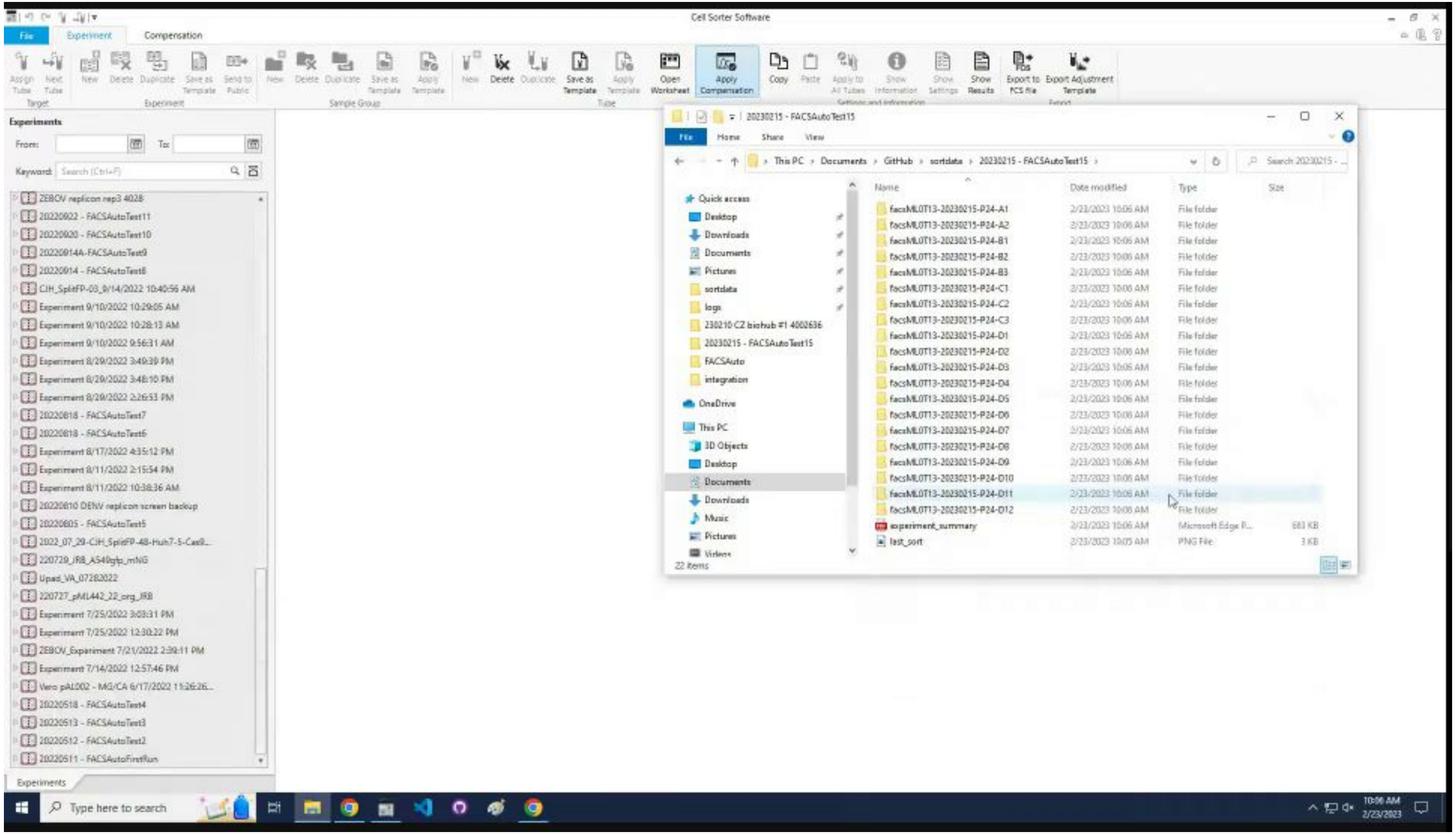
