## Supplemental Figures for "An open-source FACS automation system for high-throughput cell biology"

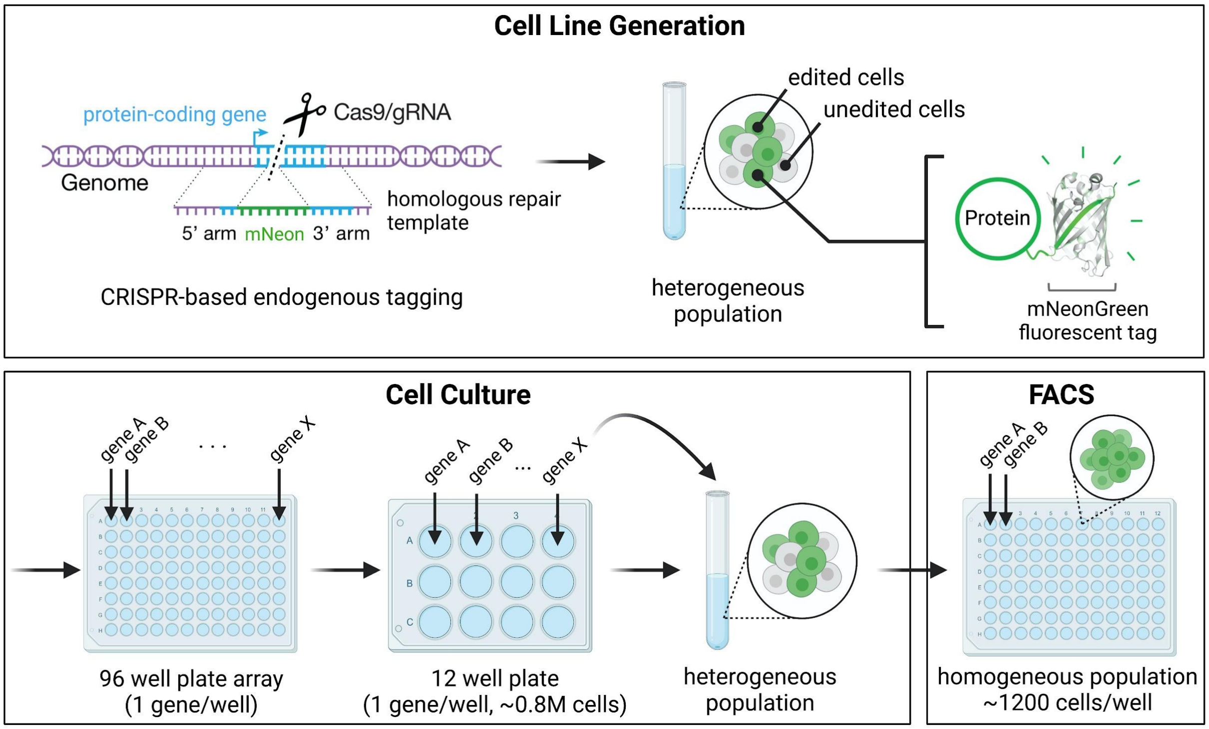


**S1 Fig. A CRISPR engineering workflow creates a library of fluorescently tagged protein-coding genes.**

CRISPR/Cas methods are used to tag endogenous protein-coding genes with a mNeonGreen fluorescent reporter. Corresponding cell pools contain a heterogeneous mixture of unedited (non-fluorescent) cells and cells with the successful edit. The gene edits are performed in a 96 well plate for high-throughput library generation. Prior to sorting, each cell population is expanded into 12-well plates and batches of 12 samples are processed for sorting with the FACS automation platform. In our case study, we aim to isolate 1,200 cells from the top 1% of the brightest fluorescent cell population for each sample.

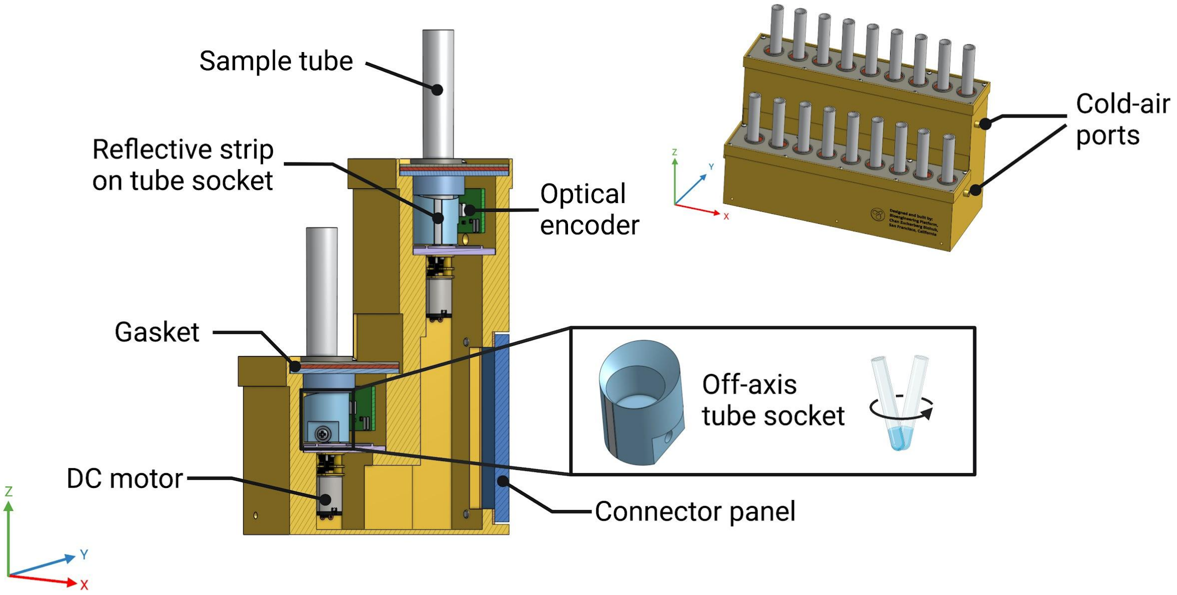


**S2 Fig. The tube housing stores, cools, and shakes the samples during the automation run.**

Samples are stored in a tube housing prior to starting the automation. Up to 18 samples can be housed at a time. The tube housing is cooled with a stream of cold air generated by a vortex tube, maintaining the sample temperature close to 4°C. Each sample sits inside of a tube socket attached to a DC motor. The motor rotates the tube off-axis such that it is shaken to maintain the cells in suspension. Optical encoders ensure that the sample tubes are presented to the robotic gripper in a controlled orientation.

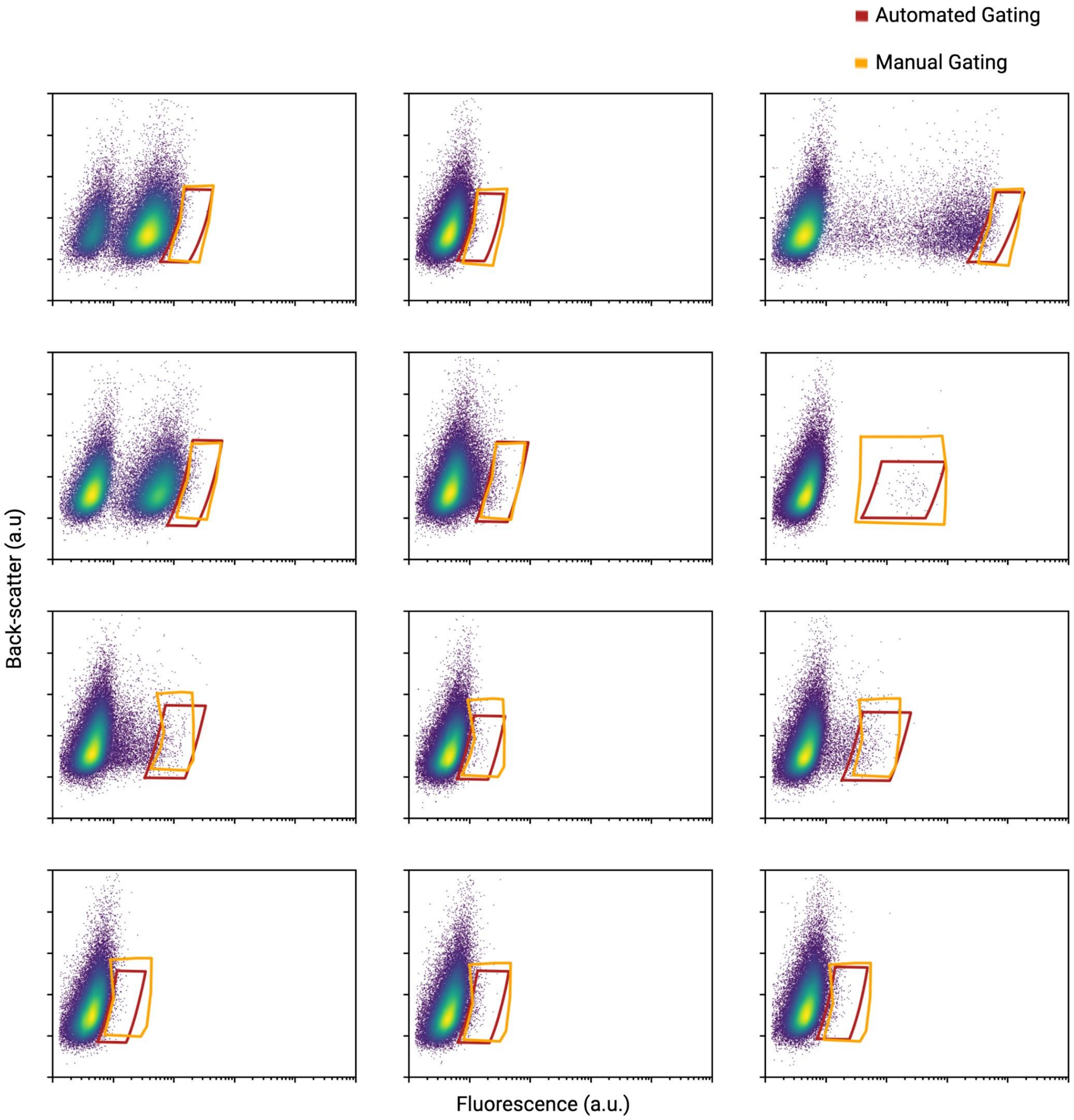


**S3 Fig. Overlays of the gates drawn for the same 12 samples sorted both by hand and with the automation show similar gated populations.**
Twelve samples were sorted both manually and with the automated system to compare the total time and active time to sort (Fig 4). In this figure, each graph represents the fluorescence profile from a cell population where a different gene was targeted (and will therefore display different fluorescent properties). The gates drawn by the algorithm (red) are overlaid with the gates

drawn manually by the user (orange) on the final population of fluorescent cells. The gated populations are similar. Importantly, the automated gating consistently draws the same gate from the distribution of cells, whereas the manual gates can vary from user-to-user and day-to-day.
